# Hybridization drives mitochondrial DNA degeneration and metabolic shift in a species with biparental mitochondrial inheritance

**DOI:** 10.1101/2022.04.29.490087

**Authors:** Mathieu Hénault, Souhir Marsit, Guillaume Charron, Christian R. Landry

**Author notes:** Current address: Département de biologie, chimie et géographie, Université du Québec à Rimouski, Rimouski, QC, Canada. Current address: Centre de foresterie des Laurentides, Ressources naturelles Canada, Québec, QC, Canada.

## Abstract

Mitochondrial DNA (mtDNA) is a cytoplasmic genome that is essential for respiratory metabolism. While uniparental mtDNA inheritance is most common in animals and plants, distinct mtDNA haplotypes can coexist in a state of heteroplasmy, either because of paternal leakage or de novo mutations. MtDNA integrity and the resolution of heteroplasmy have important implications, notably for mitochondrial genetic disorders, speciation and genome evolution in hybrids. However, the impact of genetic variation on the transition to homoplasmy from initially heteroplasmic backgrounds remains largely unknown. Here, we use *Saccharomyces* yeasts, fungi with constitutive biparental mtDNA inheritance, to investigate the resolution of mtDNA heteroplasmy in a variety of hybrid genotypes. We previously designed 11 crosses along a gradient of parental evolutionary divergence using undomesticated isolates of *Saccharomyces paradoxus* and *Saccharomyces cerevisiae*. Each cross was independently replicated 48 to 96 times, and the resulting 864 hybrids were evolved under relaxed selection for mitochondrial function. Genome sequencing of 446 MA lines revealed extensive mtDNA recombination, but recombination rate was not predicted by parental divergence level. We found a strong positive relationship between parental divergence and the rate of large-scale mtDNA deletions, which lead to the loss of respiratory metabolism. We also uncovered associations between mtDNA recombination, mtDNA deletion, and genome instability that were genotype-specific. Our results show that hybridization in yeast induces mtDNA degeneration through large-scale deletion and loss of function, with deep consequences for mtDNA evolution, metabolism and the emergence of reproductive isolation.

## INTRODUCTION

Mitochondrial DNAs (mtDNAs) are genomes that are maintained and expressed in mitochondria, which are ATP-producing organelles shared by virtually all Eukaryotes (Müller et al. 2012). Although mtDNAs vary greatly in terms of size and contents across Eukaryotes, all encode genes involved in respiration and protein synthesis (Adams and Palmer 2003). Most animals and plants have uniparental maternal mtDNA inheritance. Many exceptions exist to this rule, either because of constitutive uniparental or biparental inheritance (Breton et al. 2015; Wilson and Xu 2012; Wang et al. 2015), or leakage of paternal mtDNA (Parakatselaki and Ladoukakis 2021). Leakage and biparental mtDNA inheritance lead to a state of heteroplasmy, which is the intracellular coexistence of multiple mtDNA haplotypes. Evidence for paternal leakage is growing, even in species long considered to have strict uniparental maternal inheritance (Ladoukakis and Zouros 2017; Kondo et al. 1990; Payne et al. 2013; Gyllensten et al. 1991). Heteroplasmy may have important implications for mtDNA evolution, notably by enabling recombination between otherwise clonal mtDNAs (Rokas et al. 2003; Tsaousis et al. 2005; Ladoukakis et al. 2011). In humans, heteroplasmy is determinant in the onset of mitochondrial genetic disease, which are linked to inherited or de novo mtDNA variants (Stewart and Chinnery 2015; Wallace 2015; Stewart and Chinnery 2021). Additionally, when diverged populations or species hybridize, heteroplasmy can lead to the coexistence of more or less diverged mtDNAs which can then interact.

Genes encoded on mtDNAs are necessary, but not sufficient for carrying their essential metabolic function. The essential interactions between gene products encoded by mtDNAs and nuclear genomes require tight coevolution (Burton and Barreto 2012; Piccinini et al. 2021). Mitonuclear coevolution has important implications in the context of heteroplasmy in hybrids. Because mtDNA inheritance is non-mendelian, heteroplasmy is generally transient, with a single mtDNA haplotype eventually becoming fixed by vegetative segregation (Birky 2001). The return to homoplasmy can fix either parental haplotype, or a recombinant mtDNA molecule containing alleles from both parents (Rokas et al. 2003; Tsaousis et al. 2005). Heteroplasmy resolution can lead to the fixation of alleles involved in mitonuclear incompatibilities, thus compromising mitochondrial function and hybrid fitness. In addition, mitochondrial alleles can interact epistatically within recombinant mtDNAs (mito-mito epistasis) (Wolters et al. 2018).

The genotype of a hybrid may contribute to determine the outcome of heteroplasmy resolution. One general prediction is that the potential for epistatic interactions scales with the level of evolutionary divergence between the parents (Orr 1995). As such, mitonuclear incompatibilities are expected to be more frequent between distantly related genomes compared to closely related ones (Burton 2022). Under the same rationale, mito-mito incompatibilities are expected to show increased prevalence between more diverged mtDNAs. Nucleotide sequence divergence between mtDNAs is also expected to reduce the opportunities for homologous recombination. Little is known about the path to heteroplasmy resolution in hybrids and its consequences for hybrid evolution. Although the study of genomes sampled from natural populations yielded important insights into mtDNA introgression, recombination and horizontal transfer (Wu et al. 2015; Rice et al. 2013; Peris et al. 2017; Leducq et al. 2017), how parental divergence shapes the neutral landscape of mtDNA recombination and loss of function is largely unknown. Answering these questions requires a model system in which heteroplasmy is dominant and that can easily generate a large diversity of F1 hybrid genotypes.

Like many other fungi, yeasts of the *Saccharomyces* genus have biparental mtDNA inheritance (Solieri 2010; Wilson and Xu 2012). This enables the inheritance of either parental mtDNA in F1 hybrids, as well as recombination between parental haplotypes (Dujon et al. 1974). Because yeast F1 hybrids are initially heteroplasmic and quickly transition to homoplasmy by mitotic vegetative segregation (Solieri 2010), they are a powerful system to investigate the factors driving heteroplasmy resolution. In addition, respiratory metabolism in yeast is facultative, enabling the study of neutral mtDNA evolution without the selective pressure to maintain respiration. The undomesticated species *Saccharomyces paradoxus* is found in natural lineages with wide variation in genetic diversity and evolutionary divergence (Kuehne et al. 2007; Xia et al. 2017; Leducq et al. 2016, 2014), allowing laboratory crosses that span various levels of parental divergence.

Mutation accumulation (MA) is a type of experimental evolution that minimizes the power of natural selection. Combined with genome sequencing, MA experiments produce near-unbiased estimates of the spectrum of changes that spontaneously occur into genomes (Lynch et al. 2016). We previously performed a large-scale MA experiment on yeast hybrids (Charron et al. 2019; Hénault et al. 2020). Parental strains comprised undomesticated isolates of *S. paradoxus* (11 strains from three lineages) and *Saccharomyces cerevisiae* (two strains, Fig. 1A). Using these parental backgrounds, we designed 11 crosses spanning a range of evolutionary divergence from intra-lineage to interspecific. Each cross was replicated 48 to 96 times with independent matings, totalling 864 hybrid MA lines. We submitted each line to periodical extreme bottlenecks by streaking for single colonies on solid medium (Joseph and Hall 2004; Lynch et al. 2008) for ~770 mitotic generations (Fig. 1B). Here, we use this collection and the associated whole-genome sequencing datasets (Hénault et al. 2020; Marsit et al. 2021) to investigate the resolution of heteroplasmy in a variety of hybrid genotypes.

**Figure 1.**
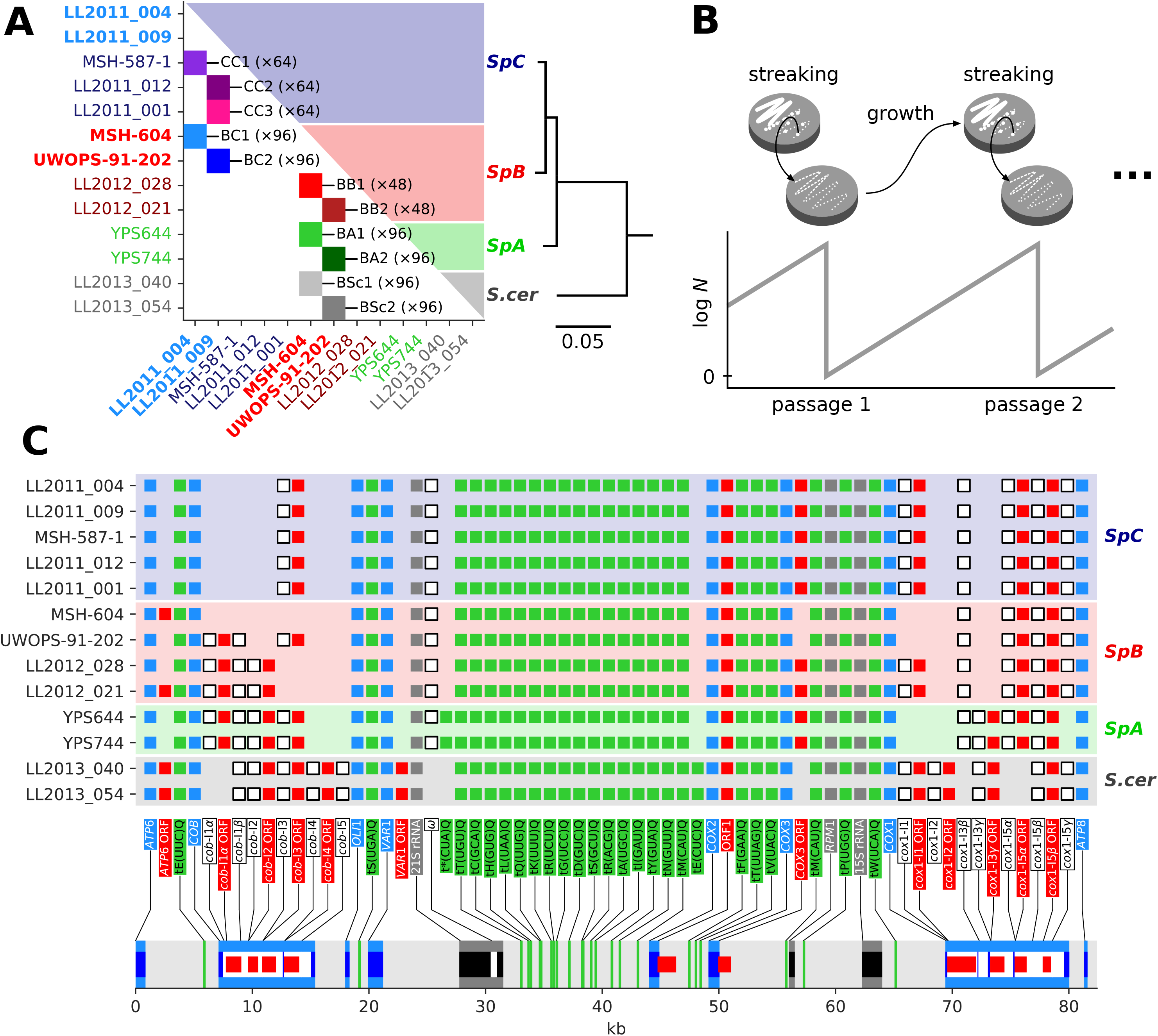
Design of the MA experiment on *Saccharomyces* hybrids and mtDNA contents of the parental strains. (*A*) Design of the MA crosses. Colored squares indicate crosses between the corresponding parental strains on both axes. Strains in bold are shared by multiple crosses. 864 independent MA lines were subdivided into 11 crosses spanning intra-lineage (*SpC*×*SpC*, CC; *SpB*×*SpB*, BB), intraspecific (*SpB*×*SpC*, BC; *SpB*×*SpA*, BA) and interspecific (*SpB*×*S. cerevisiae*, BSc) parental divergence. Each cross was replicated to initiate 48 to 96 independent evolution lines, all arising from distinct mating events. The phylogenetic tree summarizes the evolutionary divergence between parental strains (substitutions per site, based on nuclear genome-wide variants). (*B*) Microbial MA experiments minimize the efficiency of natural selection. Periodic extreme bottlenecks are achieved through streaking for single colonies on solid medium, which amplifies genetic drift. (*C*) Annotation summary of high-quality reference mtDNA assemblies for each parental strain. Colored squares denote the presence of each feature in the corresponding genome. Light blue, protein-coding genes; dark blue, protein-coding exons; white, introns; grey, RNA-coding genes; black, RNA-coding exons; red, intronic or free-standing ORFs; green, tRNAs.

## RESULTS

### Parental strains of the MA experiment display extensive variation in mtDNA content

We sequenced the genome of the 13 MA parental strains (Supplemental Table S1) using Oxford Nanopore long reads and produced high-quality mtDNA assemblies (Supplemental Fig. S1). Our long-read assemblies were consistent with a subset of five mtDNAs previously assembled from short reads (Leducq et al. 2017) (Supplemental Fig. S2). Assembly annotation revealed variation in mtDNA content, mostly in the presence of introns in the *COB* and *COX1* genes (Fig. 1C). MtDNA content was consistent within lineages, although substantial diversity was found in *SpB*. To enable comparative analyses based on a common reference sequence, the mtDNA with the largest feature set (LL2012_028) was used as a template to build an exhaustive artificial reference sequence by complementing it with introns found in other assemblies, but absent from LL2012_028.

### The landscape of mtDNA recombination is not predicted by parental divergence

For each cross, 36 to 47 hybrid MA lines were previously selected at random for short-read whole-genome sequencing, both at the initial timepoint (~60 generations post-mating) and final timepoint (after ~770 mitotic generations) of the MA experiment (Hénault et al. 2020; Marsit et al. 2021). We used this short-read data along with our mtDNA assemblies to identify a set of confident marker variants that discriminate the parental mtDNA haplotypes. We then used genotypes at marker positions to identify the recombination tracts for individual MA lines, yielding 482 independent mtDNA haplotypes. All samples were homoplasmic at both timepoints, with the vast majority of minor allele frequencies below 0.5% (Supplemental Fig. S3). However, for many lines, the mtDNA haplotypes sampled at the initial and final timepoints had incompatible genotypes, suggesting persistent segregation of distinct haplotypes in some MA lines. The frequency at which distinct haplotypes were sampled varied among crosses, reaching up to 33% of the lines in BA2 (Supplemental Table S3). All MA crosses experienced some extent of recombination, except BA1 which exhibited no recombinant mtDNA (Fig. 2A, Supplemental Fig. S4-14). Some crosses exhibited recombination hotspots, yielding high local similarity among haplotypes (Fig. 2B).

**Figure 2.**
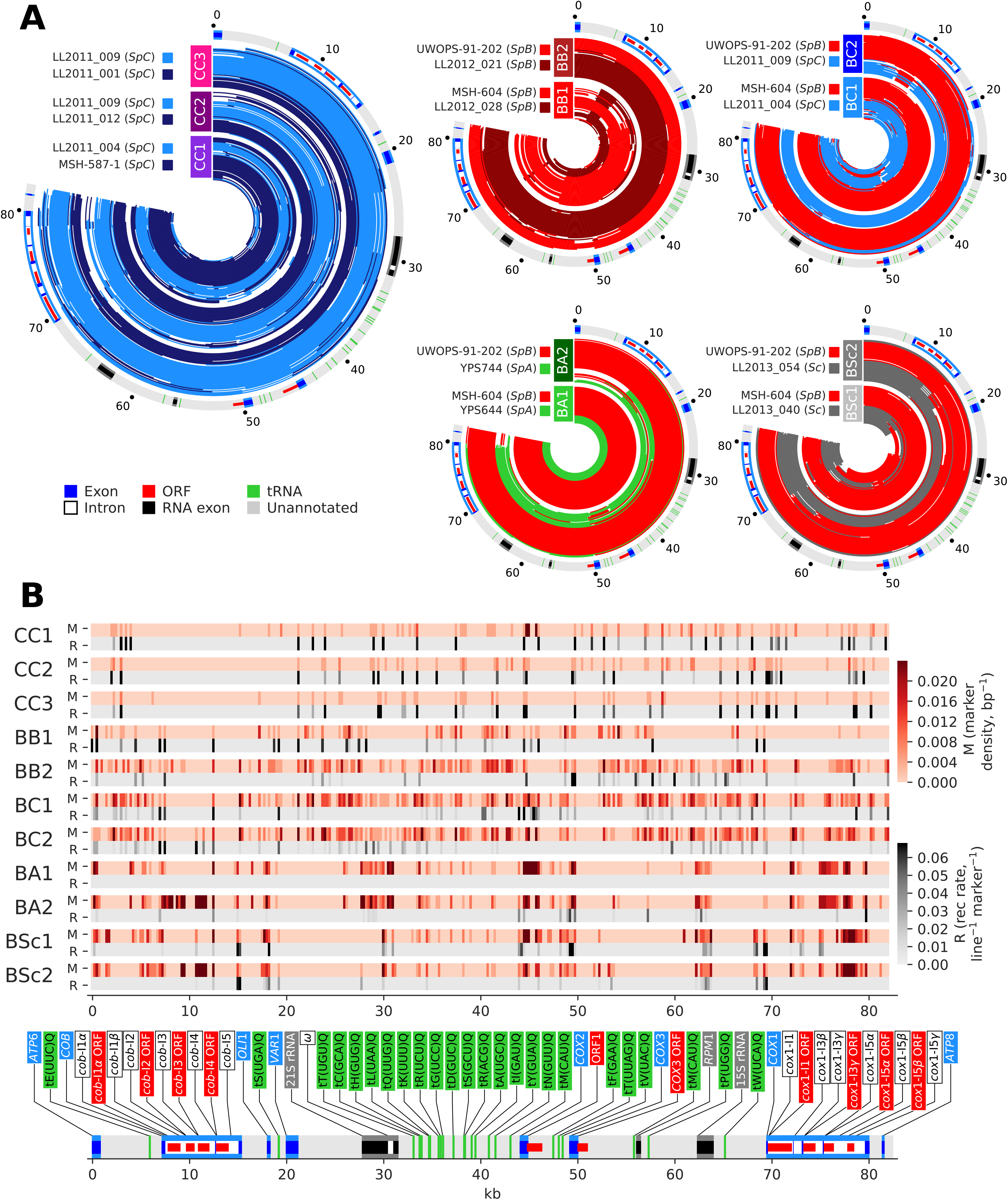
Recombination is pervasive between parental mtDNA haplotypes of hybrid MA lines. (*A*) Stacked concentric lines show recombination tracts of independent mtDNA haplotypes, colored by parental ancestry. Outer circles show annotation summary, with coordinates in kb. (*B*) Density of markers (red) and recombination junctions (black) in 250 bp windows. Recombination junction density is normalized by marker density.

With the exception of BA1, MA crosses exhibited frequencies of recombinant mtDNAs ranging from 28.9% to 60.0% (Fig. 3A). CC2 and BC1 crosses harbored the mtDNA haplotypes with the highest count of recombination tracts. The observation of strong contrasts in mtDNA recombination rates, notably between very similar hybrid genotypes, is consistent with many other aspects of genome evolution during MA (Hénault et al. 2020; Marsit et al. 2021; Fijarczyk et al. 2021). Recombination rate displayed no significant trend with evolutionary divergence between the parents of a cross (based on nuclear genome-wide variants; Fig. 3B, Supplemental Fig. S15A). Notably, the interspecific BSc crosses had recombination rates similar to intraspecific crosses. Excluding BSc crosses did not yield better correlations, suggesting an absence of relationship with evolutionary divergence, even at the intraspecific level. The average length of recombination tracts was higher for BSc crosses and had a significant positive correlation with parental divergence (Fig. 3C). Neither recombination rate nor tract length correlated with marker counts (Fig. 3B-C), suggesting that these estimates are largely unaffected by parental marker density. This indicates that mtDNA recombination rate is independent of the global evolutionary divergence level between the parents of a hybrid, contrary to the expectation that divergence should hamper recombination.

**Figure 3.**
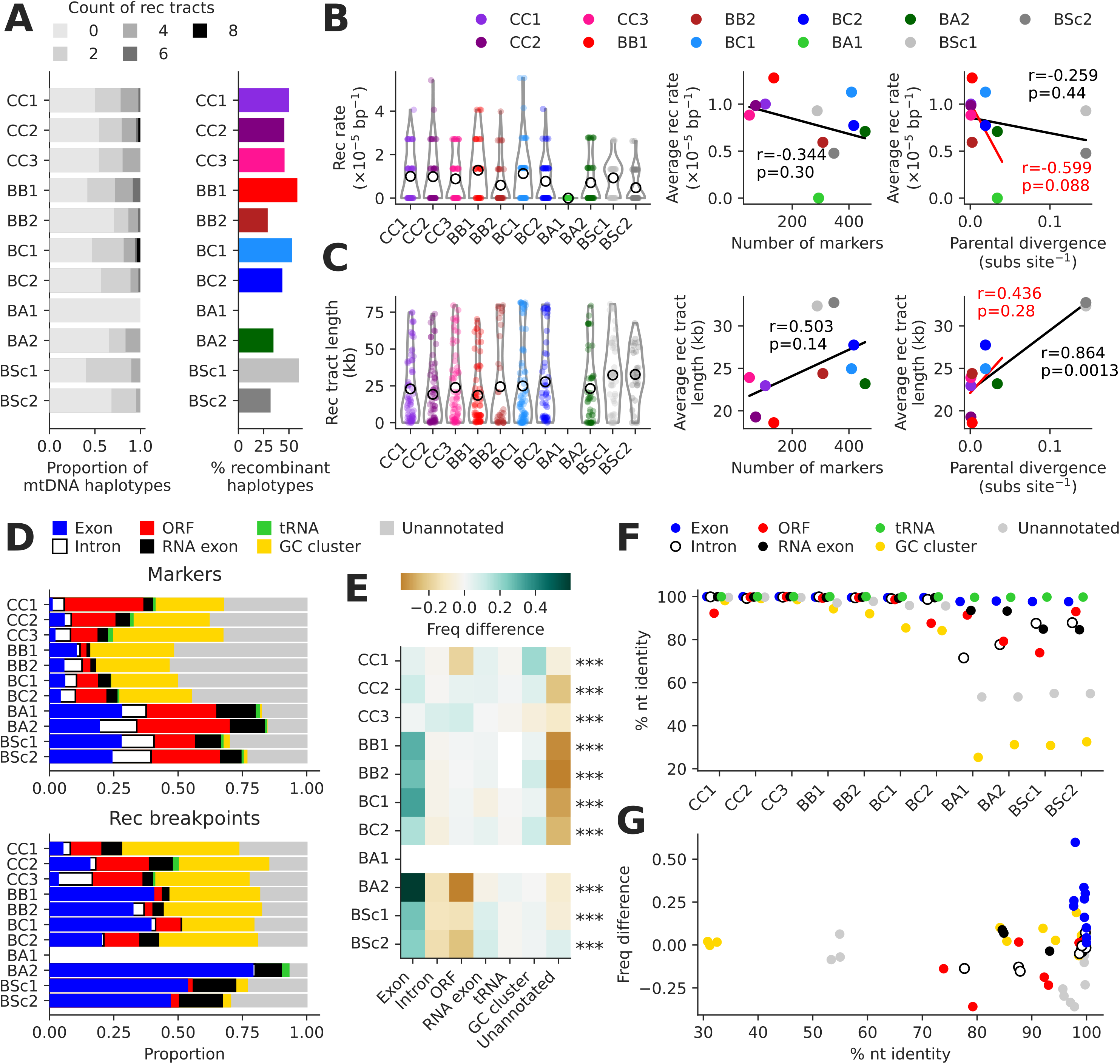
MtDNA protein-coding exons are enriched in recombination junctions. (*A*) Left: counts of recombination tracts per mtDNA haplotype. Right: percentages of recombinant haplotypes. (*B*) Left: recombination rate distributions for individual lines. Hollow circles show average recombination rates. Right: Pearson correlations between average recombination rate and number of markers or parental evolutionary divergence (substitutions per site, based on nuclear genome-wide variants). The correlation in red excludes BSc crosses. (*C*) Left: length distributions for individual recombination tracts. Hollow circles show average recombination tract lengths. Right: Pearson correlations between average recombination tract length and number of markers or parental evolutionary divergence (substitutions per site, based on nuclear genome-wide variants). The correlation in red excludes BSc crosses. (*D*) Distributions of annotation feature types for marker variants (top) and recombination breakpoints (bottom). (*E*) Overrepresentation of each annotation feature type in recombination junctions, calculated as the difference in frequency between breakpoints and markers for each feature type. FDR-corrected p-values of chi-squared tests for frequency deviations within each cross are shown (***: p≤0.001). (*F*) Percentage of nucleotide identity between parental mtDNAs for each annotation feature type. (*G*) Overrepresentation of each annotation feature type in recombination junctions against the percentage of nucleotide identity. Values for each cross are represented as separate points.

We classified recombination breakpoints by annotation feature type and contrasted the resulting distributions with parental markers (Fig. 3D). We found a strong overrepresentation of protein-coding exons in recombination breakpoints in most crosses (Fig. 3E). We next quantified the nucleotide identity between parental alleles of each cross for all mtDNA features, and asked whether it correlated with overrepresentation in recombination junctions. High sequence identity for some features in BSc crosses was consistent with inter-specific introgressions (Supplemental Fig. S16), many of which were previously described (Leducq et al. 2017; Peris et al. 2017). For instance, the introgression of *ORF1* from *S. cerevisiae* to *S. paradoxus SpB* strain UWOPS-91-202, but not to *SpB* strain MSH-604, explains the higher sequence similarity in BSc2 compared to BSc1 (Supplemental Fig. S16-17). Despite these introgressions, a phylogenetic tree built on mtDNA-encoded protein-coding DNA sequences showed the same global topology as the nuclear-encoded genes (Supplemental Fig. S15B). While protein-coding exons retain high nucleotide sequence identity in all crosses (Fig. 3F, Supplemental Fig. S15C-D), this does not solely explain their overrepresentation, as other annotation types displayed similar identity levels (Fig. 3G, Supplemental Fig. S17). In summary, mtDNA recombination occurs preferentially near protein-coding exon sequences and at a rate that is largely independent of parental evolutionary divergence.

We asked whether the overrepresentation of exons in recombination junctions could be explained by adjacent intron mobilization. Although intron mobility cannot be assessed strictly from genomic data, we looked for genomic signatures of putative mobilization. We focused on introns which show presence/absence polymorphism between the parents of each cross, making the conservative assumption that all polymorphic introns can mobilize. We examined the depth of coverage at intron-exon junctions, which shows distinctive profiles in the presence or absence of an intron. We identified eight putative mobility events not involving recombination, seven corresponding to *cox1*-I1 in BB1 and one corresponding to *cob*-I3 in BC1 (Supplemental Fig. S18). A total of 12 recombination junctions were consistent with mobilization-mediated recombination (Supplemental Fig. S19). Notably, *cox1*-I1 accounted for all the putative recombination-associated mobilization events in BB crosses. Although many intron presence/absence polymorphisms were consistent with mobilization-associated recombination, mobilization potentially explained only a minor fraction of the total recombination junctions (5/68 in BB1, 1/34 in BB2, 2/80 in BC1, 3/56 in BSc1 and 1/32 in BSc2).

Recombination may yield biases in the inheritance of mitochondrial alleles. MtDNA-wide inheritance ratios did not correlate with parental mtDNA copy number ratios estimated from depth of coverage of long-read sequencing libraries (Supplemental Fig. S20). However, copy number estimates may have low accuracy since mtDNA abundance depends on the growth phase (Galeota-Sprung et al. 2021) and we did not optimize the DNA extraction cultures for synchronicity. They nevertheless suggest that parental mtDNA copy numbers do not determine haplotype inheritance probability. We found significant deviations from mtDNA-wide inheritance ratios for some genes (Supplemental Fig. S21). Similar patterns were found in the two BSc crosses, with segments centered on *COX2* being mostly inherited from the *SpB* parent.

### Parental divergence predicts the frequency of loss of respiratory function and large-scale mtDNA deletions

We next characterized the loss of respiratory function in MA lines. The loss of respiration in yeast is spontaneous and most often explained by the loss of mtDNA integrity through large-scale deletions (Bernardi 1979). We measured the growth of all MA lines on media containing fermentable (glucose, YPD) or non-fermentable (glycerol and ethanol, YPEG) carbon sources using high-density colony array imaging. Since respiratory effects of mtDNA haplotypes or mitonuclear interactions are often revealed with exposure to higher temperatures (Li et al. 2019; Paliwal et al. 2014), growth was measured at both 25°C and 37°C. We classified lines as non-respiring if no growth was scored on YPEG at both temperatures. The frequency of respiration loss was higher for more divergent crosses (Fig. 4A). We quantified the depth of sequencing coverage along the mtDNA sequence and classified lines as harboring complete mtDNAs or mtDNAs with large-scale deletions (Supplemental Fig. S22-32). All parental strains had complete mtDNAs. The frequencies of deleted mtDNAs closely matched the frequencies of respiration loss (Fig. 4B), suggesting that the loss of mtDNA integrity is the main underlying cause. Frequencies of deleted mtDNAs had a significant positive correlation with parental divergence (Pearson correlation, r=0.88, p-value=3.5×10^−4^, Fig. 4C). For most crosses, deletions in mtDNAs had significantly negative effects on growth in YPD (Fig. 4D), similar to the effect of respiration loss in *S. cerevisiae* (Ephrussi et al. 1949; Vowinckel et al. 2021). In contrast, mtDNA deletions were associated with complete loss of growth in YPEG (Fig. 4D).

**Figure 4.**
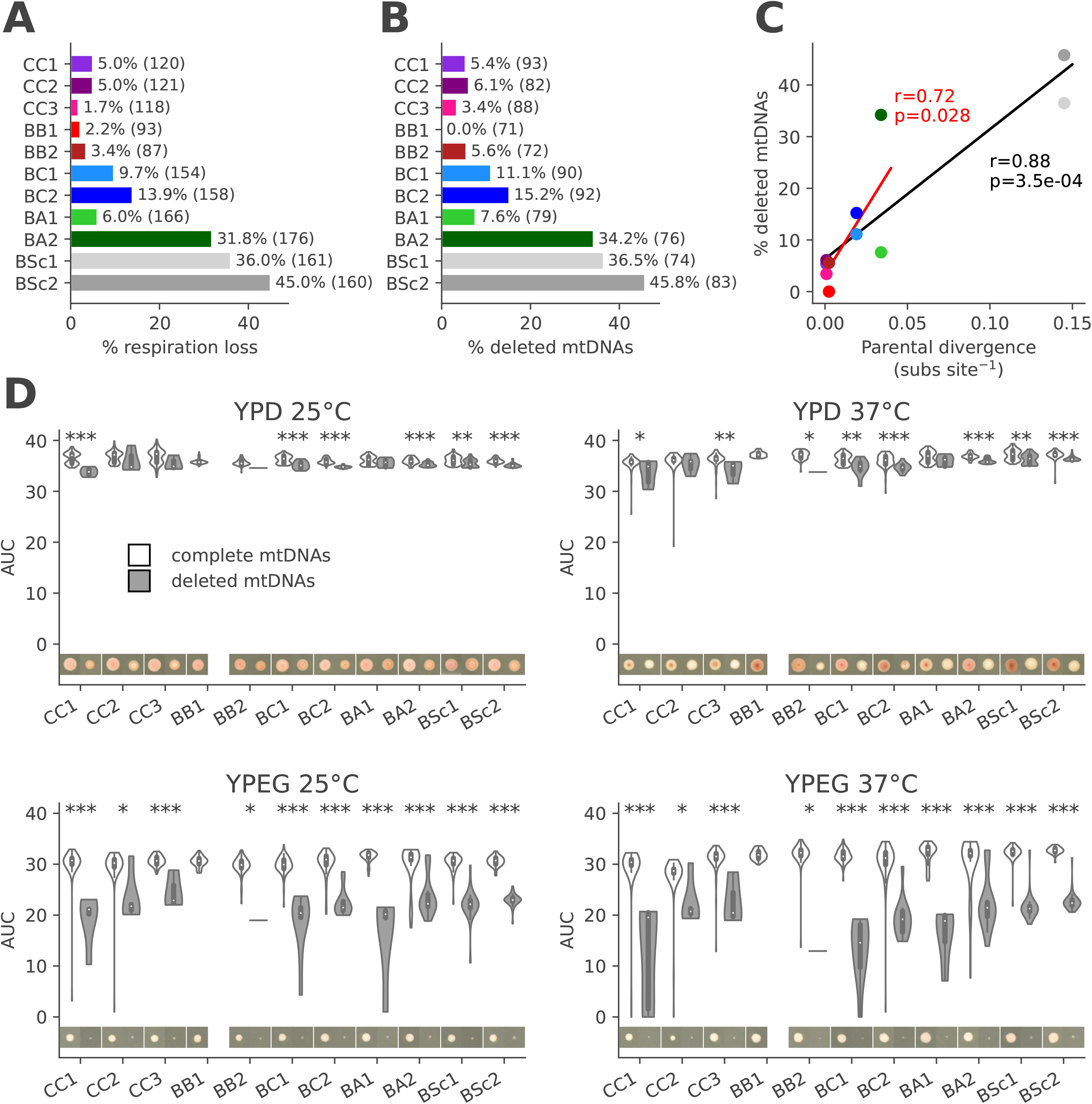
Frequent deletions in mtDNAs lead to the loss of respiratory function and occur preferentially in crosses with higher parental divergence. (*A*) Percentage of all MA lines with loss of respiration. Counts of lines (at initial and final timepoints combined) are shown. (*B*) Percentage of sequenced MA lines with mtDNA deletions. Counts of lines (initial and final timepoints combined) are shown. (*C*) Pearson’s correlation between percentage of mtDNAs with deletions and parental divergence for each cross (substitutions per site, based on nuclear genome-wide variants). The correlation shown in red excludes BSc crosses. (*D*) Area under growth curves (AUC) for lines with complete and deleted mtDNAs in the four tested conditions. FDR-corrected p-values for Mann-Whitney *U* tests between complete and deleted mtDNAs, *: p≤0.05, **: p≤0.01, ***: p≤0.001. Images show colonies of representative lines (i.e. the closest to the median) after four days.

We asked whether the frequency of respiration loss in each cross could be explained by additive parental effects. Spontaneous rates of respiration loss in parental strains were estimated by measuring the frequency of formation of non-respiring colonies, which display a characteristic small size phenotype (petite) on medium with limited glucose. Parental petite frequencies ranged from 0.2% to 4.9% (Supplemental Fig. S33A). However, the average parental petite frequency for each cross did not correlate with the percentage of lines that lost respiratory function (Pearson correlation, r=0.14, p-value=0.68; Supplemental Fig. S33B), indicating no additive parental effects. In addition, we observed that stocks of the haploid parental strains contained variable standing proportions of petite-forming cells (Supplemental Fig. S33C). The frequency of petite-forming cells did not correlate with the fraction of MA lines from the initial timepoint harboring non-recombinant, deleted mtDNAs of the corresponding parent (Pearson correlation, r=-0.29, p-value=0.36; Supplemental Fig. S33D). Thus, the frequency of respiration loss in MA crosses appears to be the product of hybridization, rather than additive parental effects or direct inheritance of defective mtDNAs.

A total of 25 MA lines had complete mtDNA haplotypes at the initial timepoint, but harbored mtDNAs with large deletions at the final timepoint. From these, 12 were consistent with partial or complete deletion of mtDNAs occurring during evolution (Supplemental Fig. S34). The remaining 13 lines showed incompatible genotypes between timepoints, indicating that distinct mtDNA haplotypes were still segregating in the lines at the initial timepoint. Likewise, ten lines gained complete mtDNAs after evolution, which is only consistent with the segregation of distinct haplotypes.

We asked if the genotype of non-deleted mtDNA haplotypes impacted the growth phenotypes of respiring lines. In many cases, growth for all mtDNA haplotypes collectively tended to be stronger at 37°C than at 25°C (Supplemental Fig. S35), in contrast to many other *Saccharomyces* hybrids (Baker et al. 2019; Li et al. 2019). However, overdominance for growth at high temperatures is not uncommon in hybrids of *S. paradoxus* (Charron and Landry 2017). The only significant difference between haplotypes was the growth advantage of non-recombinant *SpB* mtDNAs against non-recombinant *SpA* mtDNAs in the BA1 cross on YPEG at 37°C (Supplemental Fig. S35; Mann-Whitney *U* test p-value=1.56×10^−6^, FDR-corrected). Lines with *SpB* mtDNAs also tended to grow better on YPEG at 37°C in the BC2 cross, hinting towards a similar thermotolerance phenotype against *SpC* and recombinant mtDNAs (Supplemental Fig. S35-36). These results suggest that mitochondrial genetic variation is linked to thermal and respiratory adaptations in wild *S. paradoxus*.

We next investigated which mtDNA segments were mostly involved in deletions. We compared the depth of coverage profiles along mtDNAs and found recurring patterns among mtDNAs of non-respiring lines (Fig. 5). Notably, many BSc haplotypes comprised only *COX2* and *COX3*. This recurrent pattern emerged at two levels: among independently evolved lines, and in the independent crosses BSc1 and BSc2. These results suggest that mtDNA deletions may be predictable and repeatable in hybrids, and that high mtDNA deletion rates may be explained by deletion hotspots.

**Figure 5.**
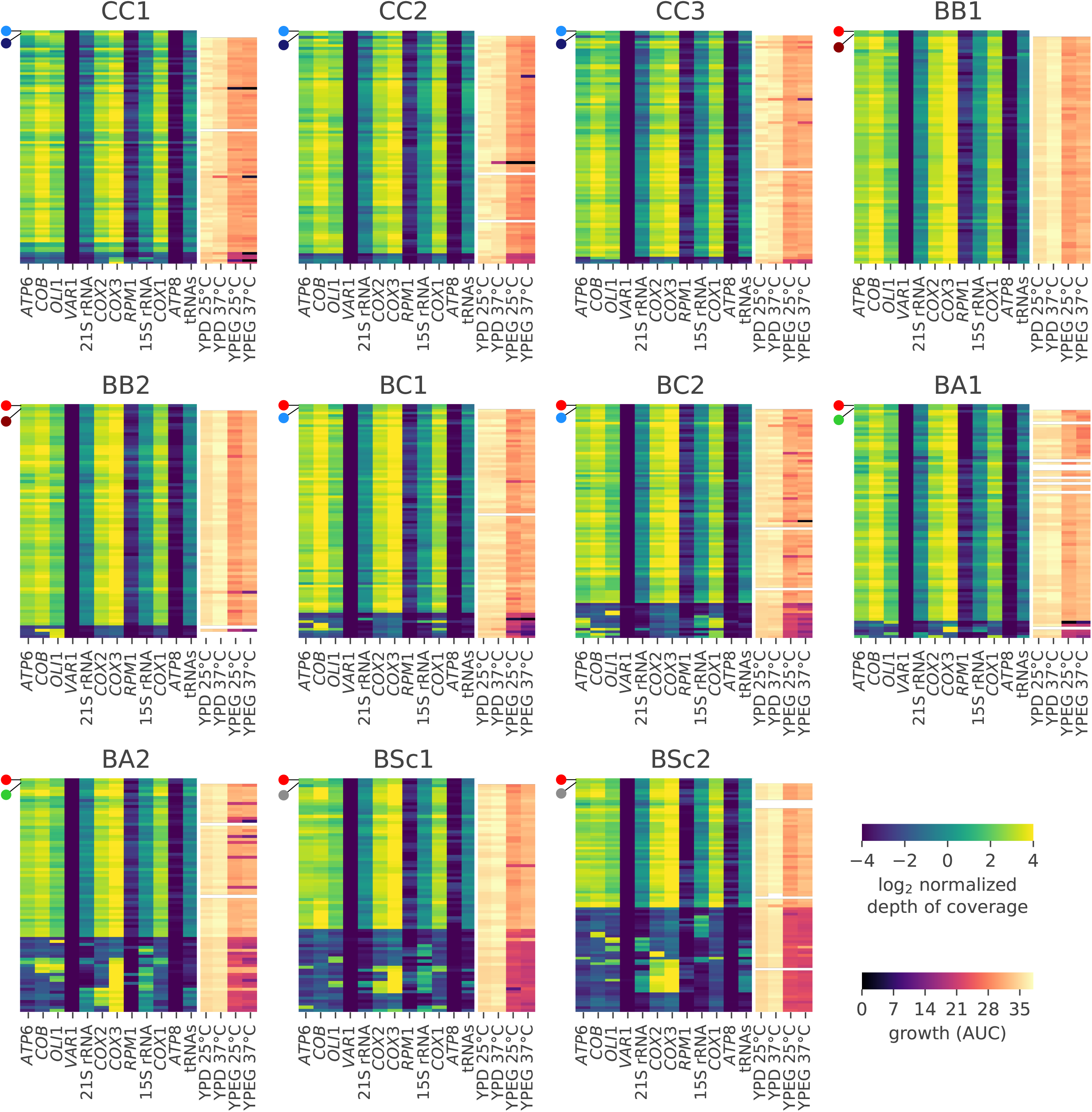
Recurrent mtDNA deletion patterns emerge in non-respiring lines. The left part of each heatmap shows the normalized depth of coverage of short-read sequencing libraries on the reference mtDNA sequence, grouped by mtDNA features. The right part of each heatmap shows the growth AUC in the four conditions tested. Rows represent individual MA lines (including initial and final timepoints). Lines are hierarchically clustered by depth of coverage profile similarity. The top lines represent the parental strains.

Depth of coverage profiles from short read mappings are highly heterogeneous along the mtDNA sequence, with some features displaying very low depth even in respiring lines (Fig. 5). To circumvent the difficulty of identifying deletion breakpoints from short reads, we randomly selected 122 lines from the final timepoint for long-read sequencing and assembled their mtDNAs. Fifteen haplotypes harbored large-scale deletions, and most deletion breakpoints were in unannotated regions (Supplemental Fig. S37, Supplemental Table S4). This result indicates that the types of sequences involved in mtDNA deletions are different from mtDNA recombination, suggesting distinct mechanisms.

### Aspects of mtDNA evolution show contrasted associations between MA crosses

We next characterized the associations between mtDNA recombination, mtDNA deletion and loss of respiration. All three were frequently observed at the initial timepoint, indicating that they can occur shortly after F1 zygotes are formed. We also observed distinct haplotypes sampled from the same line between timepoints, suggesting persistent heteroplasmy in some lines. We thus analyzed the initial and final timepoints separately. As expected, there was a strong negative relationship between mtDNA deletion and respiration (Fig. 6A,C). There were also significant associations between mtDNA deletion and recombination in BA2 and BSc1, although with opposite signs. While recombinant mtDNA haplotypes were more likely to harbor deletions in BA2, only one or two BSc1 haplotypes showed both recombination and deletion (Fig. 6B,D).

**Figure 6.**
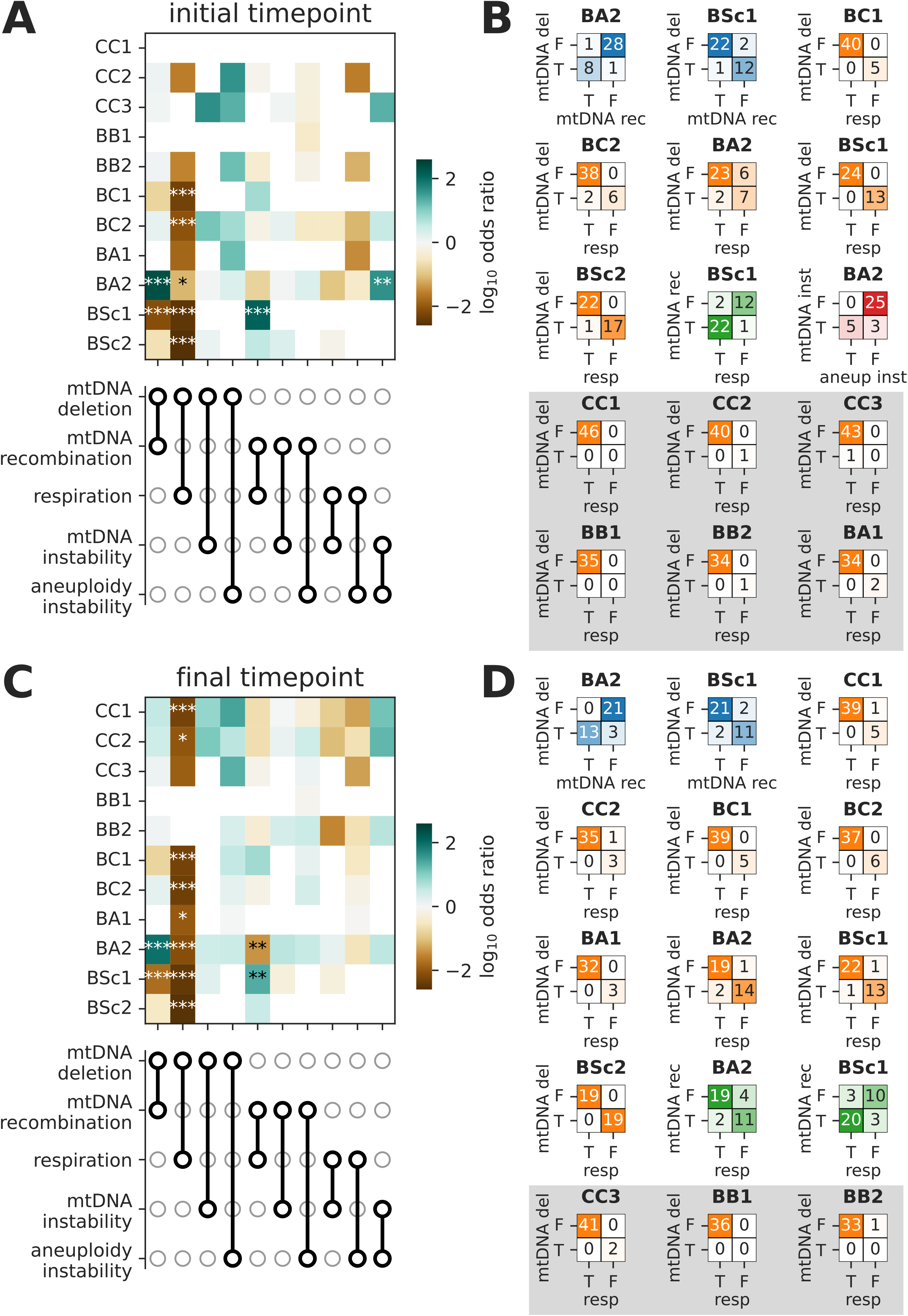
Associations between aspects of mtDNA evolution in MA lines. (*A*) Associations for lines at the initial timepoint. The heatmap shows the odds ratio for each pair of binary variables among mtDNA deletion, mtDNA recombination, respiration, mtDNA instability and aneuploidy instability. FDR-corrected p-values of Fisher’s exact test, *: p≤0.05, **: p≤0.01, ***: p≤0.001. (*B*) Counts of MA lines for each comparison with a statistically significant association, classified as true (T) or false (F) for the given variable. The shaded background comprises associations between mtDNA deletion and respiration that were not statistically significant. (*C*,*D*) Same as A and B, but for lines at the final timepoint.

While most MA lines had the expected negative association between mtDNA deletions and respiration, we observed many exceptions. Many non-respiring lines had non-deleted mtDNAs, which could be explained by other types of mutations (mitochondrial or nuclear) leading to respiration loss. However, mtDNAs with large deletions (which are incompatible with respiration) were found in six and three respiring lines at the initial and final timepoints, respectively (Fig. 6B,D). Moreover, loss of respiration without deletion was found in six lines at the initial timepoint, all in BA2, which are unlikely explained by de novo mutations given the short evolution time. We tested whether these inconsistencies were explained by differences between the cryopreserved archive stocks used for genome sequencing, and the copies of those stocks used for phenotyping. Spot assays in the four conditions revealed that both stocks exhibited the same phenotypes (Supplemental Fig. S38), thus ruling out an effect of stock propagation.

We looked for genomic changes other than mtDNA deletions that could have occurred rapidly in MA line evolution and explain respiration loss. We searched for de novo nucleotide variants in mtDNAs of the MA lines by looking for confident variants that were private to single lines. We found a handful of candidate de novo mutations at the initial timepoint but none were in coding regions (Supplemental Table S5), making them poor candidates to explain respiration loss. We next focused on aneuploidies in the nuclear genome, which were previously shown to arise early in MA (Fijarczyk et al. 2021; Marsit et al. 2021). Several inconsistent lines had aneuploidies, including five BA2 lines with monosomies for chromosomes I, III, VI and IX and/or trisomies for chromosomes IV and XII (Supplemental Fig. S39). However, parental allele frequencies on these chromosomes closely matched the diploid expectation of 50%. We quantified the deviation in depth of coverage from the value expected given the copy number call for each chromosome. Many lines with aneuploidy calls showed high deviations from the values expected from true aneuploidies (Supplemental Fig. S39A). We validated that these deviations were not artifacts from variation in library size (Supplemental Fig. S40). Continuous variation in depth of coverage and symmetrical parental allele frequencies are consistent with unbiased, segregating aneuploidies rather than fixed aneuploidies. These aneuploidies arose from single euploid clones in the population expansion leading to DNA extraction. We thus term this phenotype aneuploidy instability.

Focusing on BA2 at the initial timepoint, we assessed the presence of two mtDNA loci (*ATP6* and 21S rRNA) by PCR for all lines (Supplemental Table S6). While globally consistent with the sequencing libraries, both PCR amplicons were detected in the two respiring lines inferred to have mtDNA deletions from the genome sequencing (Supplemental Fig. S39C). Furthermore, at least one amplicon was missing for five out of six non-respiring lines with complete mtDNAs. This suggests that minor mtDNA variants segregating in the archive stocks were selected by chance when single clones were isolated for genome sequencing. We term the inconsistency between mitochondrial deletion and respiratory function mtDNA instability. BA2 had the highest proportion of lines with mtDNA instability, and those were primarily found at the initial timepoint (Supplemental Fig. S41A). We found a statistically significant association of mtDNA instability with aneuploidy instability in BA2 at the initial timepoint (Fig. 6B). Out of eight lines displaying mtDNA instability, five also were among the lines with the highest aneuploidy instability. There was a similar trend in other crosses, although only BA2 was statistically significant (Supplemental Fig. S41B). This suggests that a proportion of MA lines, specifically in the BA2 cross, suffered from high genomic instability, manifested by both frequent aneuploidy and mtDNA deletion.

## DISCUSSION

How distinct mtDNA haplotypes interact in hybrid backgrounds to shape the resolution of heteroplasmy and the evolution of mitochondrial functions remains understudied. In this study, we used *Saccharomyces* yeasts as a model to probe the neutral evolution of mtDNAs in initially heteroplasmic hybrids through experimental evolution by MA. We uncovered that recombination rates are independent of parental divergence, and recombination occurs preferentially near protein-coding exons. However, parental divergence is a strong predictor of the rate of large-scale mtDNA deletions, which is the primary mechanism for loss of respiratory metabolism. We also uncovered that some crosses have contrasted associations between mtDNA recombination and deletion, as well as shared genomic instability at the level of mtDNAs and nuclear aneuploidies.

Recombination between mtDNAs in heteroplasmic *S. cerevisiae* has been known for decades (Dujon et al. 1974). Recombination is likely central in the replication of mtDNAs, making it an inherent part of mitochondrial genetics rather than an accidental byproduct of heteroplasmy (Dujon 2020). Genomic studies brought further support for the idea that mtDNA recombination is pervasive (Fritsch et al. 2014; Bágeľová Poláková et al. 2021). Fritsch and collaborators characterized recombination in pools of diploid F1 hybrids from intraspecific crosses of *S. cerevisiae*, and found that recombination hotspots were concentrated in tRNA clusters and rRNA genes (Fritsch et al. 2014). Bágeľová Poláková and collaborators investigated three recombinant mtDNA haplotypes in F1 hybrids between *S. cerevisiae* and *S. paradoxus*, and attributed most of the observed recombination events to the mobilization of introns and free-standing ORFs (Bágeľová Poláková et al. 2021). Thus, our study is the first large-scale report of mtDNA recombination at the intra- and inter-specific levels, allowing the resolution of 482 independent mtDNA haplotypes. Our data confirms that mtDNA recombination is pervasive in a variety of hybrid genotypes. In contrast to previous studies, we found an association of recombination junctions with protein-coding exons and only a minor potential contribution for putative intron mobilization.

We observed associations between mtDNA recombination and deletion that suggest a mechanistic interaction, although this remains to be investigated. The most contrasting patterns were found between BA2 and BSc1, two crosses with frequent mtDNA deletions. In BA2, haplotypes harboring deletions were almost always recombinant, and non-deleted haplotypes rarely showed evidence for recombination. In contrast, in BSc1, mtDNA recombination and deletion were almost mutually exclusive. This is consistent with differences in mtDNA architecture between *S. paradoxus* lineages *SpB* and *SpA*. While *SpB* mtDNAs are collinear with other North American lineages and with *S. cerevisiae*, *SpA* harbors rearranged mtDNAs (Leducq et al. 2017; Yue et al. 2017). As such, homologous recombination between rearranged mtDNAs could lead to frequent deletions. However, deletion breakpoints were mostly located in unannotated regions, contrasting with recombination breakpoints and suggesting a distinct mechanism. The association between mtDNA instability and aneuploidy instability in BA2 is more challenging to interpret. BA2 was also the cross with the most frequent segregation of distinct mtDNA haplotypes between timepoints, suggesting rampant, persistent heteroplasmy. Nevertheless, it raises the possibility that both mtDNA and aneuploidy instability could be linked or share a common underlying cause. These types of genomic instability in BA2 add to many others previously characterized in our MA crosses (Marsit et al. 2021).

Experimental evolution by MA minimizes the efficiency of natural selection, enabling a selectively unbiased characterization of mtDNA evolution. The characterization of de novo mutation in nuclear genomes of our MA lines revealed signatures that were entirely consistent with neutral evolution (Fijarczyk et al. 2021). The frequent loss of respiratory function also suggests the absence of purifying selection. Although large-scale deletions leading to respiration mostly happened before the initial timepoint, a dozen MA lines had mtDNA haplotypes consistent with deletion occurring between the initial and final timepoints. While the MA design minimized the efficiency of natural selection at the level of individual cells, it did not alleviate it at the intracellular level. MtDNA haplotypes can outcompete others if they have a replication advantage, regardless of their effect on organismal fitness (Ma and O’Farrell 2016; Taylor et al. 2002). As such, some mtDNA haplotypes could have reached homoplasmy because of replication advantages.

Our results show that respiratory function is primarily lost through large-scale deletions within mtDNAs. The level of replication of our MA experiment yielded a high number of recombinant haplotypes. Nevertheless, cases of respiration loss in our experiment are likely entirely explained by mtDNA deletions. Additionally, we found no significant negative effect of non-deleted, recombinant mtDNA haplotypes on respiration. In BSc1, the non-significant growth difference between lines with *SpB* versus recombinant haplotypes on YPEG at 25°C was the second largest effect size, suggesting that some recombinant haplotypes might be disadvantaged. Despite this hypothetical case, we found no support for negative mito-mito epistasis, contrary to previous reports (Wolters et al. 2018; Leducq et al. 2017).

Mitonuclear incompatibilities are known to contribute to reproductive isolation between species of the *Saccharomyces* genus (Chou et al. 2010; Lee et al. 2008; Jhuang et al. 2017), along with other types of reproductive barriers (Hunter et al. 1996). Even at the intraspecific level in *S. cerevisiae* and *S. paradoxus*, crosses between diverged isolates often have low fertility (Greig et al. 2003; Charron et al. 2014; Leducq et al. 2016; Eberlein et al. 2019), including most of our MA crosses (Charron et al. 2019; Marsit et al. 2021). The loss of respiration adds another reproductive barrier, since meiosis (sporulation) is dependent on functioning respiratory metabolism (Jambhekar and Amon 2008). Indeed, our previous results showed a strong association between respiration loss and sporulation inability (Charron et al. 2019). Yeast can divide strictly mitotically in permissive laboratory conditions, and mitosis is dominant in the wild (Tsai et al. 2008). Sporulation is triggered in starvation conditions (Bautz Freese et al. 1982) and is likely to be essential in nature, thus selecting for mtDNA integrity. Nevertheless, some natural isolates of *S. cerevisiae* were reported to completely lack mtDNAs (De Chiara et al. 2020), suggesting that loss of sporulation ability can arise in nature. As such, mtDNA degeneration through large-scale deletion and loss of function in hybrids acts as an additional mechanism of reproductive isolation, both within and between species.

In conclusion, our study provides the first large-scale investigation of neutral heteroplasmy resolution and mtDNA evolution in hybrids. Our results show that distinct aspects of neutral mtDNA evolution can be either genotype-specific or, in contrast, highly predictable given the genomic composition of hybrids. We show that hybridization can trigger mtDNA degeneration with profound metabolic consequences and the emergence of reproductive isolation.

## METHODS

### Long read library preparation and sequencing

Stocks of the 13 haploid parental strains (Supplemental Table S1) and of 122 randomly selected MA lines from the final timepoint (Charron et al. 2019; Hénault et al. 2020) were thawed, streaked on YPD agar medium (1% yeast extract, 2% bio-tryptone, 2% glucose, 2% agar) and incubated at room temperature for three days. Cultures in YPD medium (1% yeast extract, 2% bio-tryptone, 2% glucose) were inoculated with single colonies and incubated overnight at room temperature without shaking. DNA was extracted from the cultures following a standard phenol-chloroform protocol. Oxford Nanopore native (PCR-free) genomic DNA libraries were prepared in multiplex with kits SQK-LSK109 and EXP-NBD104 (Oxford Nanopore Technologies, Oxford, UK). Libraries were sequenced with FLO-MIN106 (revC) flowcells on a MinION sequencer (MIN-101B). The sequencing and basecalling were run on a MinIT computer (MNT-001) with MinKNOW v3.3.2 and guppy v3.0.3. Reads were demultiplexed using guppy_basecaller v3.1.5.

### Long read mtDNA assemblies

Long reads matching mtDNA sequences were extracted from whole-genome libraries. For each strain, we made a compound reference genome comprising the nuclear genome of the corresponding *S. paradoxus* lineage(s) (*SpA*: LL2012_001, *SpB*: MSH-604, *SpC*: LL2011_012) (Eberlein et al. 2019), or *S. cerevisiae* (YPS128) (Yue et al. 2017), supplemented with a concatenated mitochondrial contig comprising *S. paradoxus SpB* (YPS138), *SpA* (CBS432) and (YPS138) and *S. cerevisiae* (S288c) (Yue et al. 2017) mtDNAs joined by 100 Ns. Nanopore libraries were mapped on the compound references using Minimap2 v2.20-r1061 (Li 2018a) with preset map-ont. Reads mapping to the mtDNA contig were extracted using SAMtools v1.15 (Li et al. 2009) and Seqtk v1.3-r106 (Li 2018b). Assembly of the extracted reads was performed using wtdbg2 v2.5 (Ruan and Li 2019) with options −AS 2 −g 72k −X 50 and a grid search of the following parameters: −k {23, 21, 19, 17, 15, 13, 11}, −p {8, 5, 2, 1, 0}, −l {4096 2048 1024}.

For parental strains, draft assemblies were aligned against the reference mtDNA assembly (Yue et al. 2017) of the closest species/lineage (*SpB* and *SpC*: YPS138, *SpA*: CBS432, *S. cerevisiae*: S288c) using MUMmer v3.23 (Kurtz et al. 2004). For each strain, the best assembly was selected on the basis of a quality score computed from the alignment of each query assembly using custom Python v3.9.10 (Van Rossum and Drake 2009) scripts. A genome coverage metric was first computed as the sum of absolute deviations from 1 of both query and reference coverage. A contiguity metric was computed as the inverse of the largest contig size in the assembly. Both coverage and continuity metrics were normalized to *Z*-scores and summed to yield the final score. Assemblies with the lowest score were circularized at the start of the *ATP6* gene and resulting duplications were manually trimmed. Circularized assemblies were polished by first aligning the basecalled long reads to draft assemblies using Minimap2 v2.20-r1061 with preset map-ont. Alignments were polished using the raw nanopore signal using nanopolish v0.13.3 (Loman et al. 2015) with parameter −–min-candidate-frequency 0.1. Short read libraries for each parental strain (Hénault et al. 2020; Marsit et al. 2021) were aligned on draft assemblies using BWA-MEM v0.7.17 and polishing was performed using Pilon v1.22 (Walker et al. 2014). Assemblies were aligned against the reference mtDNA sequence of *S. cerevisiae* S288c (Yue et al. 2017) using MUMmer v3.23. For strains YPS744, LL2011_012, MSH-587-1, MSH-604 and UWOPS-91-202, assemblies were compared to the corresponding mtDNA short-read assemblies from (Leducq et al. 2017) using the dnadiff tool from the MUMmer v3.23 suite.

For MA lines, draft assemblies were polished with basecalled long reads using medaka v1.4.4 (https://github.com/nanoporetech/medaka) with model r941_min_high_g303. Assemblies were circularized at the start of the *ATP6* gene (if present) and aligned to the artificial mtDNA reference (see below) using MUMmer v3.23.

### MtDNA assembly annotation and generation of mtDNA artificial reference

MtDNA assemblies of the 13 parental strains were annotated using the MFannot server (https://megasun.bch.umontreal.ca/cgi-bin/mfannot/mfannotInterface.pl) and annotations were manually curated. MFannot master files were converted to the GFF3 format using the mfannot2gff.pl script from the MITONOTATE pipeline (Seah 2016). We employed the nomenclature of *S. cerevisiae* S288c (Cherry et al. 2012) for genes and tRNAs. A reference library of intron sequences was assembled from the mtDNA assemblies and annotations of *S. paradoxus* strains CBS432 (GenBank accession JQ862335.1) (Procházka et al. 2012) and CBS7400 (GenBank accession KX657749.1) (Sulo et al. 2017), and *S. cerevisiae* S288c (GenBank accession AJ011856.1) (Foury et al. 1998). Introns annotated from our assemblies were renamed after their best hit from a search against the reference library using BLASTN v2.11.0 (Camacho et al. 2009), following the nomenclature of (Lambowitz and Belfort 1993). From the comparative analysis of mtDNA annotations, we built an artificial mtDNA reference comprising the largest set of features. We used the LL2012_028 genome as a template, and added introns *cob*-I3, *cob*-I4, *cob*-I5 and *cox1*-I3γ from YPS744 or LL2013_054 at the orthologous splice sites using custom Python v3.9.10 scripts. GC clusters on the artificial mtDNA reference were annotated using the four reference sequences presented in Table 1 of (Wu and Hao 2015) as queries for BLASTN v.2.11.0 with parameter word_size 7. The resulting hits were collapsed using BEDTools merge v2.30.0 (Quinlan and Hall 2010) and manually inspected to add or merge adjacent GC-rich segments. Parental mtDNAs were aligned to the artificial mtDNA contig using Mugsy v1r2.3 (Angiuoli and Salzberg 2011), and coordinate conversion tables were built using a custom Python v3.9.10 script.

### Phylogenetic analysis of protein-coding genes

Maximum likelihood phylogenetic trees were constructed using protein-coding genes of the nuclear and mitochondrial genomes. For mtDNAs, spliced protein-coding sequences of the eight canonical genes were aligned using MUSCLE v3.8.31 (Edgar 2004) and concatenated into a multiple sequence alignment of 6699 positions. For nuclear genomes, 81 orthogroups (each comprising a total of 26 *S. paradoxus* and *S. cerevisiae* sequences) were randomly chosen from (Eberlein et al. 2017), and protein-coding DNA sequences were concatenated to yield a multiple sequence alignment of 124,041 positions. Maximum likelihood phylogenetic trees were computed using RAxML-NG v1.1.0 (Kozlov et al. 2019) with the GTR+G model initiated with 10 parsimony trees and 200 bootstraps. For mitochondrial genes, the evolutionary divergence between the parents of each cross were extracted. For nuclear genes, the evolutionary divergence value of each cross was computed as the average of all pairwise distances within or between the corresponding parental lineage(s) or species. Phylogenetic trees were built for individual spliced protein-coding genes, introns and ORFs). DNA sequences were aligned using MUSCLE v3.8.31 and trees were inferred using the neighbor-joining algorithm implemented in the ape v5.6-2 library (Paradis and Schliep 2019) in R v4.2.1 (R Core Team 2016).

### Short read alignment and variant calling

Short read libraries corresponding to the 13 parental haploid strains and 447 hybrid yeast MA lines, sampled at both the initial (~60 mitotic generations after zygote formation) and final (after ~770 mitotic generations) timepoints, were downloaded from NCBI (BioProject PRJNA515073). Short reads were trimmed using Trimmomatic v0.36 (Bolger et al. 2014) with parameters ILLUMINACLIP:{custom adapters file}:2:30:10 TRAILING:3 SLIDINGWINDOW:4:15 MINLEN:36 (Hénault et al. 2020). We built a concatenated reference genome comprising nuclear contigs of *S. paradoxus SpB* strain MSH-604 (Eberlein et al. 2019) and *S. cerevisiae* strain YPS128 (Yue et al. 2017), to which we appended the artificial mtDNA contig. Reads were aligned to this reference using BWA-MEM v0.7.17 and mappings to the mtDNA contig were extracted using SAMtools v1.8. Variant calling was performed separately for each cross using FreeBayes v1.3.5 (Garrison and Marth 2012). Multiallelic variants were decomposed using vcfbreakmulti from vcflib v1.0.2 (Garrison 2018).

### Definition of mtDNA recombination tracts

The characterization of recombination in mtDNA haplotypes was performed using custom Python v3.9.10 scripts. For each cross, marker variants were identified for the parental genomes based on variant calls from short-read alignments on the artificial mtDNA reference. Candidate markers were filtered as follows. Each variant had to be supported by at least two read alignments. Parental alleles at each marker locus were validated using whole-genome alignments of the high-quality genome assemblies of each parental mtDNA and the associated coordinate conversion tables. Markers were required to have nucleotide calls and assembly alignments for both parents, thus excluding insertion and deletions.

For MA lines of each cross, variants corresponding to parental markers were filtered to exclude loci with low depth of coverage. For each line we defined a threshold as the highest value between the 10^th^ percentile of depth of coverage values for the nuclear genome of each line, or 20 reads. Variants passed the filter if supporting reads for the major allele were higher than the threshold. Since each MA line was sampled for sequencing at two timepoints, mtDNA haplotypes were compared across timepoints of individual lines to determine whether they were clonal or independent haplotypes that segregated from initially heteroplasmic lines. For each line, we computed the pairwise identity between mtDNA haplotypes as the proportion of identical calls at marker variants (excluding missing positions in either mtDNA). If the identity was 95% or higher, we kept only one mtDNA (preferentially the latest timepoint) as representative of the line. Independent haplotypes within each cross were clustered by identity using the nearest point algorithm from SciPy v1.8.0 (Virtanen et al. 2020).

Recombination tracts were defined by scanning and grouping adjacent marker variants of the same genotype along the genome, with a tolerance of one marker of the opposite genotype (i.e. two marker variants of the opposite genotype were required to break a tract). Tracts were circularized at the ends of the mtDNA contig. Recombination breakpoints were defined as the marker variants at the ends of each tract. Recombination breakpoint and marker variant densities were computed for 100 bp windows along the genome. Recombination rates were computed as the number of recombination tracts per haplotype divided by the combined length of parental mtDNAs. Average recombination rate and tract length for each cross were correlated with marker variant counts and parental evolutionary divergence values (see section *Phylogenetic analysis of protein-coding genes*).

### MtDNA inheritance biases

Recombination tracts were used to compute the proportion of each mtDNA haplotype that was inherited from each parent. The inheritance ratios were averaged over all independent haplotypes of a cross to yield genome-wide average inheritance ratios. The depth of coverage of long reads on each parental mtDNA assembly compared to nuclear chromosomes was used to compute a relative mtDNA copy number. For individual genes within haplotypes, discrete inheritance calls were generated by attributing either parental ancestry to annotations that had ratios of 10% or less, or 90% or more. Genes with ratios between those values were considered as recombinant and excluded. Binomial tests were used to test the deviation of non-recombinant genes from genome-wide inheritance ratios.

### Growth measurements of the MA lines collection

During the MA experiment, MA lines were sampled at regular intervals and archived in cryopreserved stocks in 96 well microplates. Archives stocks sampled at the initial and final timepoints were replicated on YPD medium and cryopreserved as stock copies. For high-throughput phenotyping, stock copies were thawed and spotted on YPD agar medium in OmniTray plates (Thermo Fisher Scientific, Waltham, USA) using a BM5-SC1 colony processing robot (S&P Robotics Inc., Toronto, Canada) and incubated at room temperature for four days. Colonies were replicated on YPD agar by condensing them into six 384-position source arrays and incubated at room temperature for three days. Source arrays were replicated twice on YPD agar medium and twice on YPEG agar medium (1% yeast extract, 2% bio-tryptone, 3% glycerol, 3% ethanol, 2% agar). Plates were incubated at 25°C or 37°C for the first round of selection during two days. Plates were replicated for the second round of selection on the same media and photographed after 24, 48 and 96 hours of incubation with the robotic platform. Colony sizes were quantified from the plate photos using gitter v1.1 (Wagih and Parts 2014) in R v3.3.2 (R Core Team 2016). Colony sizes in pixels were log-transformed and summed across imaging timepoints to estimate the area under growth curves (AUC). One colony (BSc2, line F53, initial timepoint, YPD 37°C) was removed due to visible contamination. Calls of growth absence based on pixel quantifications were manually curated with raw images.

### Measurement of parental petite frequencies

Parental rates of spontaneous petite formation were measured by streaking haploid stocks on YPD agar + NAT (100 μg ml^−1^ nourseothricin) or YPD agar + G418 (200 μg ml^−1^ geneticin), depending on the selection marker at the *HO* locus, and incubated at 25°C for six days. Two single large colonies per strain were resuspended in sterile water and diluted to an optical density at 600 nm (OD_600_) of 2.5×10^−4^, and 200 μl were plated on YPdG agar medium (1% yeast extract, 2% bio-tryptone, 3% glycerol, 0.1% glucose, 2% agar). Plates were incubated at 25°C for four days and photographed. The standing frequency of non-respiring cells within parental stock cultures was measured by spotting 20 μl of stocks on YPD agar and incubating for three days. Cells were resuspended in sterile water and diluted to an OD_600_ of 5×10^−4^, and 100 μl were plated on YPdG agar. Plates were incubated at 25°C for five days and photographed.

### Characterization of mtDNA deletions

Depth of coverage values along mtDNAs were extracted from short read alignments using SAMtools v1.15. Median depth values were computed for 100 bp windows and normalized by the depth of coverage on nuclear chromosomes. Depth values were grouped by annotation type and the fraction of positions with normalized depth values higher than 0.1 was extracted. For each cross, these fractions were normalized into *Z*-scores, and an mtDNA completion score was computed as the sum of *Z*-scores across annotation types for individual mtDNA haplotypes. Haplotypes were sorted by mtDNA completion score to classify them as complete or deleted, and the classification was manually curated by examination of the depth of coverage profiles. Haplotypes were clustered by Pearson correlation coefficient between normalized depth profiles using the nearest point algorithm from scipy v1.8.0.

### Detection of intron presence/absence polymorphisms

Presence or absence of introns in mtDNA haplotypes of MA lines was called by analyzing depth of coverage profiles at intron-exon junctions. For each polymorphic intron, the marker variants closest to both junctions (upstream or downstream) were identified. 100 bp windows centered around each junction were used to analyze depth of coverage profiles. Depth of coverage values were normalized by the 80^th^ percentile of depth values within a window. Euclidean distance and Pearson correlation coefficient were used to compare profiles of each line with both parental profiles. Evidence for intron presence was called at a junction if the euclidean distance and Pearson correlation coefficient were respectively lower and higher for comparisons to the parent harboring the intron. An intron was called present if its two junctions showed consistent evidence. Putative intron mobilization events not involving recombination were defined as intron presence with two flanking markers of the opposite ancestry. Putative recombination-associated mobilization events were defined as intron presence with one flanking marker of the opposite ancestry, consistent with neighboring recombination tracts.

### Confirmation of growth phenotypes by spot assays

A subset of lines classified as having mtDNA instability at the initial timepoint were selected for growth phenotypes validation by spot assays. Original archive stocks and copies were thawed, spotted on YPD agar medium and incubated at room temperature. Spots were used to inoculate precultures in YPD medium that were incubated overnight at 30°C. Precultures were diluted at 1 OD ml^−1^ in sterile water and five 5X serial dilutions were prepared. Aliquots of 5 μl of each dilution were spotted twice on YPD agar and YPEG agar. Plates were incubated at 25°C or 37°C for three days and photographed.

### De novo mutations identification

Candidate de novo single nucleotide mutations were identified from variant calls based on short read alignments. First, variant calls supported by at least five reads and which had an alternative allele ratio of at least 0.8 were extracted. Variants which were only observed in a single line were further checked for the absence of matching alleles in the parental variant calls from all crosses, and from lines of all the other crosses. For each remaining candidate, we extracted the proportion of lines in the cross with missing data at that locus. We defined a more stringent alignment criteria for supporting reads by requiring the length of the aligned reference segment to be between 141 and 161 bp (for 151 bp reads), and counted those high confidence alignments for each candidate variant. Finally, we counted the number of supporting reads for matching alleles in the whole dataset, excluding the line of interest. The final set of candidate de novo mutations consisted in variants for which the total number of supporting reads was at least twice the number of supporting reads in the whole dataset excluding the line, and had at least two supporting reads with high confidence alignments.

### Depth of coverage deviations on nuclear chromosomes

Variant calls from short-read alignments and aneuploidy calls were previously done by Fijarczyk and collaborators (Fijarczyk et al. 2021). BSc crosses were excluded from this analysis as the mapping strategy was different (using concatenated *SpB* and *S. cerevisiae* nuclear reference genomes). For each cross, 10000 nuclear variants were chosen at random. Allele ratios were examined along nuclear chromosomes. For individual lines, the median depth of coverage of each chromosome was normalized by the genome-wide median to yield the depth ratio. The copy number ratio was defined by dividing chromosome copy number calls by the global ploidy level of each line. Depth of coverage deviation scores were computed as (depth ratio – copy number ratio). Absolute deviation scores were averaged over all chromosomes and lines with an average absolute deviation score of at least 0.08 were classified as having the aneuploidy instability phenotype.

### MtDNA genotyping by PCR

Two loci were genotyped for presence or absence by PCR for all lines of the BA2 cross at the initial and final timepoints. DNA was extracted by resuspending colonies in 40 μl NaOH 20 mM and incubating at 95°C for 20 min. Two mtDNA loci in *ATP6* and 21S rRNA were amplified with primers and cycles detailed in Supplemental Table S6 (Leducq et al. 2017). Amplicons of the nuclear mating type loci were amplified as positive controls with primers and cycles detailed in Supplemental Table S6 (Huxley et al. 1990). Presence and absence were scored by agarose gel electrophoresis.

## DATA ACCESS

The long read data generated in this study have been submitted to the NCBI BioProject database (https://www.ncbi.nlm.nih.gov/bioproject/) under accession number PRJNA828354. The mtDNA assemblies generated in this study have been submitted to GitHub (https://github.com/Landrylab/mito_ma). Scripts and annotation data generated in this study are available in Supplemental Code and have been submitted to GitHub (https://github.com/Landrylab/mito_ma).

## ACKNOWLEDGEMENTS

We thank Anna Fijarczyk for comments on the manuscript and contributions to sequencing data preprocessing and analysis. We thank Hélène Martin for contributions to sequencing data preprocessing. We thank the members of the Landry laboratory for discussions on the project. We thank three anonymous reviewers for their useful comments on the manuscript. This project was supported by funding to C.R.L. from an NSERC discovery grant (RGPIN-2020-04844) and an FRQNT team grant (2019-PR-254415), and a NSERC Alexander Graham Bell doctoral scholarship to M.H. CRL holds the Canada Research Chair in Cellular Systems and Synthetic Biology.

## AUTHOR CONTRIBUTIONS

CRL and MH designed the project. MH performed the analyses and experiments. MH, SM and GC performed the MA evolution experiment and short read sequencing as cited. MH wrote the manuscript with inputs from all authors.

