## Supplemental Code for "Hybridization drives mitochondrial DNA degeneration and metabolic shift in a species with biparental mitochondrial inheritance": assembly_comparison.ipynb

```
# %% [code]
import pandas as pd
from Bio import SeqIO
import itertools
from collections import Counter
from matplotlib import pyplot as plt
import seaborn as sns
rc_sns = {'ytick_color':'k', 'xtick_color':'k', 'text_color':'k', 'font.sans-serif':'DejaVu Sans', 'figure.facecolor':(1,1,1,1)}
sns.set_style(style='ticks', rc=rc_sns)

# %% [markdown]
# Import metadata

# %% [code]
assemblies = pd.read_csv('/mnt/HDD3/mito_nanopore/leducq/assemblies.csv').set_index('strain')
assemblies['atp6'] = [f'{s}.atp6.fasta' for s in assemblies.index]

# %% [code]
assemblies_alias = dict()
assemblies_alias.update({g:g.split('.')[0]+'\nnanopore' for g in assemblies['nanopore']})
assemblies_alias.update({g:g.split('.')[0]+'\nillumina' for g in assemblies['atp6']})

# %% [markdown]
# Assembly comparisons with dnadiff

# %% [code]
#export comparisons to be made with dnadiff
comp = []
for (i,j) in itertools.combinations(list(assemblies['nanopore']) + list(assemblies['atp6']), 2):
    comp.append('\t'.join([i, j, f'{i[:-6]}_{j[:-6]}']))
#with open('/mnt/HDD3/mito_nanopore/leducq/comparisons.tsv', 'w') as handle:
#    handle.write('\n'.join(comp))

# %% [code]
comp = pd.read_csv('/mnt/HDD3/mito_nanopore/leducq/comparisons.tsv', sep='\t', header=None)

# %% [code]
#import genomes
genomes = {}
for g in assemblies['atp6'].values:
    genomes[g] = SeqIO.read(f'/mnt/HDD3/mito_nanopore/leducq/{g}', 'fasta')
for g in assemblies['nanopore'].values:
    genomes[g] = SeqIO.read(f'/mnt/HDD3/mito_nanopore/leducq/{g}', 'fasta')

# %% [code]
# compute statistics
comp_stats = []
for i in comp.index:
    genome1, genome2, name = comp.loc[i]
    snps = pd.read_csv(f'/mnt/HDD3/mito_nanopore/leducq/{name}.snps', sep='\t', header=None)
    snps['snp'] = snps.apply(lambda x: '.' not in x[[1,2]].values, axis=1)
    snps['ins'] = snps[1]=='.'
    snps['del'] = snps[2]=='.'
    snp_count = snps.value_counts('snp')[True]
    ins_count = snps.value_counts('ins')[True]
    del_count = snps.value_counts('del')[True]
    
    with open(f'/mnt/HDD3/mito_nanopore/leducq/{name}.rdiff', 'r') as handle:
        rdiff = Counter([line.split('\t')[1] for line in handle.read().splitlines()])
    gap_count = rdiff['GAP']
    
    comp_stats.append([genome1, genome2, name, snp_count, ins_count, del_count, gap_count])

comp_stats = pd.DataFrame(comp_stats, columns=['genome1', 'genome2', 'name', 'snp_count', 'ins_count', 'del_count', 'gap_count'])
comp_stats['size_diff'] = comp_stats.apply(lambda x: len(genomes[x['genome1']].seq)-len(genomes[x['genome2']].seq), axis=1)
comp_stats['indel_count'] = comp_stats['ins_count'] + comp_stats['del_count']

# %% [markdown]
# Fig S2

# %% [code]
fig = plt.figure(figsize=[8,8])
gs = plt.GridSpec(ncols=2, nrows=2, left=0.15, right=0.96, bottom=0.15, top=0.94,
                  hspace=0.6, wspace=0.6)
m_alias = {'snp_count':'single nucleotide variants',
           'indel_count': 'single nucleotide indels',
           'gap_count':'gaps',
           'size_diff':'size difference'}
for m, ax_idx in zip(['snp_count', 'indel_count', 'gap_count', 'size_diff'], 
                    itertools.product([0,1], [0,1])):
    ax = fig.add_subplot(gs[ax_idx])
    
    df = comp_stats.pivot_table(index='genome1', columns='genome2', values=m, aggfunc=lambda x: x)
    df = df.loc[assemblies['nanopore'], assemblies['atp6']]
    if m == 'size_diff':
        sns.heatmap(df, ax=ax, cmap='bwr', center=0, annot=True, fmt='.0f', cbar=False)
    else:
        sns.heatmap(df, ax=ax, cmap='Blues', annot=True, fmt='.0f', cbar=False)
    ax.set_yticklabels(assemblies['nanopore'].apply(lambda x: assemblies_alias[x]), rotation=45, va='top')
    ax.set_ylabel('')
    ax.set_xticklabels(assemblies['atp6'].apply(lambda x: assemblies_alias[x]), rotation=45, ha='right')
    ax.set_xlabel('')
    ax.set_title(m_alias[m], size=14)
    
fig.text(0.02, 0.95, 'A', size=24, weight='bold')
fig.text(0.52, 0.95, 'B', size=24, weight='bold')
fig.text(0.02, 0.47, 'C', size=24, weight='bold')
fig.text(0.52, 0.47, 'D', size=24, weight='bold')
    
#plt.show()
#plt.savefig('/home/mathieu/mhenault_landrylab/Publications/mito_ma/resubmission2_GRes/fig/assembly_comparison.svg')
plt.close()
```
