## Supplemental Code for "Hybridization drives mitochondrial DNA degeneration and metabolic shift in a species with biparental mitochondrial inheritance": divergence.ipynb

```
# %% [code]
import os
import re
import random
import pandas as pd
import numpy as np
from scipy import stats
import dendropy
from Bio import SeqIO
from Bio import SeqRecord
from Bio import Seq
from collections import Counter
import itertools
from matplotlib import pyplot as plt
import seaborn as sns
rc_sns = {'ytick_color':'k', 'xtick_color':'k', 'text_color':'k', 'font.sans-serif':'DejaVu Sans', 'figure.facecolor':(1,1,1,1)}
sns.set_style(style='ticks', rc=rc_sns)

# %% [markdown]
# Import metadata

# %% [code]
ma_strains = pd.read_csv('/mnt/HDD3/mito_ma/script/ma_strains_mthap.tsv', sep='\t', index_col=0)
parents_dict = {'CC1':('HI.4803.003.N712---N505.LL2011_004', 'NS.1250.002.N712---N505.MSH-587-1'),
               'CC2':('HI.4803.003.N711---N508.LL2011_009', 'NS.1250.002.N712---N507.LL2011_012'),
               'CC3':('HI.4803.003.N711---N508.LL2011_009', 'NS.1250.002.N712---N506.LL2011_001'),
               'BB1':('HI.4803.003.N712---N504.MSH-604', 'HI.4803.003.N712---N508.LL2012_028'),
               'BB2':('HI.4803.003.N711---N507.UWOPS-91-202', 'HI.4803.003.N712---N503.LL2012_021'),
               'BC1':('HI.4803.003.N712---N504.MSH-604', 'HI.4803.003.N712---N505.LL2011_004'),
               'BC2':('HI.4803.003.N711---N507.UWOPS-91-202', 'HI.4803.003.N711---N508.LL2011_009'),
               'BA1':('HI.4803.003.N712---N504.MSH-604', 'HI.4803.003.N712---N506.YPS644'),
               'BA2':('HI.4803.003.N711---N507.UWOPS-91-202', 'HI.4803.003.N712---N517.YPS744'),
               'BSc1':('HI.4803.003.N712---N504.MSH-604', 'HI.4803.003.N712---N507.LL2013_040'),
               'BSc2':('HI.4803.003.N711---N507.UWOPS-91-202', 'HI.4803.003.N712---N502.LL2013_054')}
cross_color = ma_strains.groupby('cross')['cross_color'].apply(lambda x: x.iloc[0]).to_dict()
cross_order = ['CC1','CC2','CC3','BB1','BB2','BC1','BC2','BA1','BA2','BSc1','BSc2']

taxa = pd.read_csv('/mnt/HDD3/mito_ma/divergence/taxa.csv', index_col=3)

# %% [markdown]
# Nuclear genome

# %% [code]
# import orthgroups list
all_orthogroups = [f for f in os.listdir('/mnt/HDD1/Dropbox/Spar_orthogroups/All_orthogroups_FASTA/') if re.match('Y.+\.fasta', f)]

# %% [code]
# Randomly select a subset of orthogroups
random.seed(42)
random_orthogroups = random.sample(all_orthogroups, 100)

# %% [code]
def get_strain_name(strain):
    
    m = re.search('/(.+)_200.fa.gtf', strain)
    if m:
        strain = m.group(1)
    else:
        m = re.search('/(.+).fa.gtf', strain)
        if m:
            strain = m.group(1)
    if strain+'.fasta' == f:
        strain = 'S288c'
    
    return strain

# %% [code]
# extract statistics on the selected orthogroups
og_summary = []
for f in random_orthogroups:
    SEQ = list(SeqIO.parse(f'/mnt/HDD1/Dropbox/Spar_orthogroups/All_orthogroups_FASTA/{f}', 'fasta'))
    for seq in SEQ:
        strain = get_strain_name(seq.id)
        length = len(seq.seq)
        og_summary.append([f, strain, length])
og_summary = pd.DataFrame(og_summary, columns=['gene', 'strain', 'length'])

og_summary = og_summary.pivot_table(index='gene', columns='strain', values='length')
og_summary.dropna(axis=0, inplace=True)
# exclude far-east strain and SpC*
og_summary.drop(['N-44','C03','A04','B03'], axis=1, inplace=True)

# %% [code]
# concatenate orthogroup sequences for nucelar genes
#initialize concatenated sequences
concat_nuc = {strain:SeqRecord.SeqRecord(seq='', id=strain, description='') for strain in og_summary.columns}
for f in og_summary.index:
    SEQ = list(SeqIO.parse(f'/mnt/HDD1/Dropbox/Spar_orthogroups/All_orthogroups_FASTA/{f}', 'fasta'))
    if len(SEQ) == 30:
        if len(set([len(seq.seq) for seq in SEQ])) == 1:
            for seq in SEQ:
                # parse strain name from fasta entries
                strain = get_strain_name(seq.id)
                if strain in concat_nuc:
                    #append sequence
                    concat_nuc[strain].seq += seq.seq

#with open('/mnt/HDD3/mito_ma/divergence/nuc_random_orthogroups.fasta', 'w') as handle:
#    SeqIO.write(concat_nuc.values(), handle, 'fasta')

# %% [code]
# parse the tree produced by RAxML
parents_tree = dendropy.Tree.get(path='/mnt/HDD3/mito_ma/divergence/raxml.support.renamed', schema='newick')
parents_tree.phylogenetic_distance_matrix().as_data_table().write_csv(out='/mnt/HDD3/mito_ma/divergence/nuc_pairwise_divergence.csv')
parents_dist_matrix = pd.read_csv('/mnt/HDD3/mito_ma/divergence/nuc_pairwise_divergence.csv', index_col=0)

# %% [markdown]
# Mitochondrial genome

# %% [code]
# list file names for the mitochondrial gene alignments
mito_genes = ['atp6.anchor.aln.fasta',
              'atp8.anchor.aln.fasta',
              'atp9.anchor.aln.fasta',
              'cob.spliced.aln.MSH-604.fasta',
              'cox1.spliced.aln.fasta',
              'cox2.anchor.aln.fasta',
              'cox3.anchor.aln.fasta',
              'rps3.anchor.aln.fasta']

# %% [code]
# concatenate orthogroup sequences for nucelar genes
#initialize concatenated sequences
concat_mt = {strain:SeqRecord.SeqRecord(seq='', id=strain, description='') for strain in ma_strains.loc[ma_strains['cross']=='P', 'strain'].values}
for f in mito_genes:
    SEQ = list(SeqIO.parse(f'/mnt/HDD3/mito_nanopore/muscle/aln/{f}', 'fasta'))
    if len(SEQ) == 13:
        if len(set([len(seq.seq) for seq in SEQ])) == 1:
            for seq in SEQ:
                strain = seq.id.split('.')[2]
                if strain in concat_mt:
                    #append sequence
                    concat_mt[strain].seq += seq.seq
                else:
                    print(f, ' no ', strain)

#with open('/mnt/HDD3/mito_ma/divergence/mt_orthogroups.fasta', 'w') as handle:
#    SeqIO.write(concat_mt.values(), handle, 'fasta')

# %% [code]
#parse tree
mt_tree = dendropy.Tree.get(path='/mnt/HDD3/mito_ma/divergence/mt_orthogroups.fasta.raxml.support', schema='newick')
mt_tree.phylogenetic_distance_matrix().as_data_table().write_csv(out='/mnt/HDD3/mito_ma/divergence/mt_pairwise_divergence.csv')
mt_dist_matrix = pd.read_csv('/mnt/HDD3/mito_ma/divergence/mt_pairwise_divergence.csv', index_col=0)
mt_dist_matrix.index = [i.replace(' ', '_') for i in mt_dist_matrix.index]
mt_dist_matrix.columns = [i.replace(' ', '_') for i in mt_dist_matrix.columns]

# %% [code]
#compute distances for crosses
dist_per_cross = []
#for cross in ['CC', 'BB', 'BC', 'BA', 'BSc']:
for cross, (p1, p2) in parents_dict.items():
    
    # mito distance
    p1, p2 = [p.split('.')[-1] for p in (p1, p2)]
    mt_dist = mt_dist_matrix.loc[p1, p2]
    
    # nuc distance
    nuc_dist = []
    Cross = cross[:-1]
    g1, g2 = Cross[0], Cross[1:]
    P1 = taxa.loc[g1, 'strain_id']
    P2 = taxa.loc[g2, 'strain_id']
    
    for i,j in itertools.product(P1, P2):
        if i != j and [i, j] == sorted([i,j]):
            nuc_dist.append(parents_dist_matrix.loc[i,j])
    nuc_dist = np.mean(nuc_dist)
    
    dist_per_cross.append([cross, Cross, mt_dist, nuc_dist])
dist_per_cross = pd.DataFrame(dist_per_cross, columns=['cross', 'Cross', 'mt_dist', 'nuc_dist'])
dist_per_cross['ratio'] = dist_per_cross['nuc_dist']/dist_per_cross['mt_dist']

# %% [markdown]
# Fig S15

# %% [code]
fig, axes = plt.subplots(nrows=2, figsize=[3,6], 
                         gridspec_kw=dict(hspace=0.4, height_ratios=[3,2], left=0.28, right=0.93, top=0.82, bottom=0.12))
ax = axes[0]
sns.scatterplot(x='nuc_dist', y='mt_dist', hue='cross', palette=cross_color, data=dist_per_cross, ax=ax)
ax.set_xlabel('nuc divergence (subs site$^{-1}$)')
ax.set_ylabel('mt divergence\n(subs site$^{-1}$)')
ax.legend(loc=6, bbox_to_anchor=[-0.05,1.25], ncol=3, frameon=False, handlelength=0.3)

ax = axes[1]
ax.bar(dist_per_cross['cross'], dist_per_cross['ratio'], color=[cross_color[cross] for cross in cross_order],  width=0.5)
ax.set_xlabel('')
ax.set_xticklabels(cross_order, rotation=90)
ax.set_ylabel('divergence ratio\n(nuc/mt)')

fig.text(0.01, 0.93, 'C', weight='bold', size=24)
fig.text(0.01, 0.35, 'D', weight='bold', size=24)

sns.despine()
#plt.savefig('/home/mathieu/mhenault_landrylab/Publications/mito_ma/resubmission2_GRes/fig/mito_nuc_divergence_CD.svg')
#plt.show()
plt.close()
```
