## Supplemental Code for "Hybridization drives mitochondrial DNA degeneration and metabolic shift in a species with biparental mitochondrial inheritance": make_liftover.py

```
#!/home/mathieu/miniconda3/bin/python

from Bio.AlignIO import parse
from sys import argv
from pandas import DataFrame, concat
from numpy import array, repeat

# parse maf alignments

def parse_maf(path, ref_name='artificial_genome.mt_art'):

    aln = list(parse(path, 'maf'))
    
    maf = {ai:{seq.id:seq for seq in a} for (ai,a) in enumerate(aln) if len(a)==2}
    # flip any alignment block for which the ref sequence is reversed
    for ai, a in maf.items():
        seq_names = a.keys()
        qry_name = [i for i in seq_names if i!=ref_name][0]
        ref = a[ref_name]
        qry = a[qry_name]

        ref_strand = ref.annotations['strand']
        qry_strand = qry.annotations['strand']
        
        if ref_strand == -1:
            ref.annotations['start'] = ref.annotations['srcSize'] - (ref.annotations['start'] + ref.annotations['size'])
            for seq in (ref, qry):
                seq.seq = seq.seq.reverse_complement()
            
                seq.annotations['strand'] = seq.annotations['strand']*(-2) # mark the flipped sequences as 2
                maf[ai][seq.id] = seq
            
        if qry_strand == -1:
            qry.annotations['start'] = qry.annotations['srcSize'] - (qry.annotations['start'] + qry.annotations['size'])
    
    return maf

def liftover_maf(maf):
    
    lift_list = []

    for ai, a in maf.items():

        ref = a['artificial_genome.mt_art']
        al = len(ref.seq)
        ref_len = ref.annotations['size']
        ref_start = ref.annotations['start']
        ref_offset = 0
        while ref.seq[ref_offset] == '-':
            ref_offset += 1

        lift = range(al)
        ref_pos = []
        for i in lift:
            # build ref positions
            if ref.seq[i] == '-':
                ref_pos.append(-1)
            else:
                ref_pos.append(ref_start)
                ref_start += 1

        ref_pos = array(ref_pos)

        lift_list.append(DataFrame([ref_pos, lift, repeat(ai, al)],
                                    index=['ref_pos','lift','aln']).T.astype(int))

    lift_list = concat(lift_list).reset_index(drop=True)
    lift_list.index = lift_list['ref_pos'].values
    
    return lift_list

in_path = argv[1]
out_path = argv[2]

lift = liftover_maf(parse_maf(in_path))
lift.to_csv(out_path)
```
