## Supplemental Code for "Hybridization drives mitochondrial DNA degeneration and metabolic shift in a species with biparental mitochondrial inheritance": mito_ma.ipynb

```
# %% [code]
import os
from os.path import getsize
from io import StringIO
import pandas as pd
import numpy as np
from scipy.cluster import hierarchy
from scipy import stats
from scipy.spatial.distance import euclidean
import statsmodels.api as sm
import statsmodels.formula.api as smf
from statsmodels.stats.multitest import multipletests
import intervaltree
import itertools
import random
import re
from Bio import SeqIO
from Bio import AlignIO
from Bio import Seq
from Bio import SeqRecord
import pysam
import gzip
from matplotlib import pyplot as plt
from matplotlib.colors import ListedColormap
from matplotlib.patches import FancyArrow
from matplotlib.patches import Arc
from matplotlib.patches import Rectangle
from matplotlib.patches import FancyBboxPatch
from matplotlib.patches import BoxStyle as bs
from matplotlib.patches import Polygon
from matplotlib.lines import Line2D
from matplotlib.colors import LinearSegmentedColormap
from matplotlib.offsetbox import OffsetImage, AnnotationBbox
from matplotlib.ticker import MultipleLocator
import seaborn as sns
from progressbar import ProgressBar
import pickle as pkl
from copy import copy
import colorcet
import PIL
rc_sns = {'ytick_color':'k', 'xtick_color':'k', 'text_color':'k', 'font.sans-serif':'DejaVu Sans', 'figure.facecolor':(1,1,1,1)}
sns.set_style(style='ticks', rc=rc_sns)

# %% [markdown]
# Import and process metadata tables

# %% [code]
# parse info table for MA lines
ma_strains = pd.read_csv('/mnt/HDD3/mito_ma/script/ma_strains.tsv', sep='\t', index_col=0)

#path on katak
path_ref = '/home/mahen44/mito_ma/ref/'
path_reads = {'HI':'/home/mahen44/MutationAccum/Helene/1_Trimm/Trimmo_1b/',
              'NS':'/home/mahen44/MA_lines_2019/1.Trimmo/Trimmo/'}

for cross, df in ma_strains.groupby('cross'):
    if cross in ['H1','H2']:
        ma_strains.loc[df.index, 'ref_map'] = f'{path_ref}ref_genome_BSc.mt_artSc.fasta'
    elif cross == 'P':
        for s, df1 in df.groupby('strain'):
            #if parent in BSc crosses, must map on both references, so that BAM headers are complete
            #for downstream variant calling
            if s in ['LL2013_040','LL2013_054']:
                ma_strains.loc[df1.index, 'ref_map'] = f'{path_ref}ref_genome_BSc.mt_artSc.fasta'
            else:
                ma_strains.loc[df1.index, 'ref_map'] = f'{path_ref}ref_genome_BSc.mt_art.fasta'
    else:
        ma_strains.loc[df.index, 'ref_map'] = f'{path_ref}ref_genome_BSc.mt_art.fasta'
        
for ins, df in ma_strains.groupby('instrument'):
    ma_strains.loc[df.index , 'R1'] = df['filename'].apply(lambda x: f'{path_reads[ins]}{x}_outR1P.fastq.gz')
    ma_strains.loc[df.index , 'R2'] = df['filename'].apply(lambda x: f'{path_reads[ins]}{x}_outR2P.fastq.gz')
    
# re-export ma_strains
#ma_strains.to_csv('/mnt/HDD3/mito_ma/script/ma_strains_map.tsv', sep='\t')

# %% [code]
# parse metadata table for MA lines and edit
ma_strains = pd.read_csv('/mnt/HDD3/mito_ma/script/ma_strains_mthap.tsv', sep='\t', index_col=0)

cross_alias = {'VL3':'CC1',
              'VL4':'CC2',
              'VL5':'CC3',
              'VL1':'BB1',
              'VL2':'BB2',
              'L1':'BC1',
              'L2':'BC2',
              'M1':'BA1',
              'M2':'BA2',
              'H1':'BSc1',
              'H2':'BSc2'}
cross_order = {j:i for i,j in enumerate(['CC1', 'CC2', 'CC3', 'BB1', 'BB2', 'BC1', 'BC2', 'BA1', 'BA2', 'BSc1', 'BSc2'])}

ma_strains['cross'] = ma_strains['cross'].replace(cross_alias)
ma_strains['strain_passage'] = ma_strains.apply(lambda x: f'{x["strain"]}_P{x["passage"]:.0f}', axis=1)

# %% [code]
# define general dictionaries
parents_dict = {'CC1':('HI.4803.003.N712---N505.LL2011_004', 'NS.1250.002.N712---N505.MSH-587-1'),
               'CC2':('HI.4803.003.N711---N508.LL2011_009', 'NS.1250.002.N712---N507.LL2011_012'),
               'CC3':('HI.4803.003.N711---N508.LL2011_009', 'NS.1250.002.N712---N506.LL2011_001'),
               'BB1':('HI.4803.003.N712---N504.MSH-604', 'HI.4803.003.N712---N508.LL2012_028'),
               'BB2':('HI.4803.003.N711---N507.UWOPS-91-202', 'HI.4803.003.N712---N503.LL2012_021'),
               'BC1':('HI.4803.003.N712---N504.MSH-604', 'HI.4803.003.N712---N505.LL2011_004'),
               'BC2':('HI.4803.003.N711---N507.UWOPS-91-202', 'HI.4803.003.N711---N508.LL2011_009'),
               'BA1':('HI.4803.003.N712---N504.MSH-604', 'HI.4803.003.N712---N506.YPS644'),
               'BA2':('HI.4803.003.N711---N507.UWOPS-91-202', 'HI.4803.003.N712---N517.YPS744'),
               'BSc1':('HI.4803.003.N712---N504.MSH-604', 'HI.4803.003.N712---N507.LL2013_040'),
               'BSc2':('HI.4803.003.N711---N507.UWOPS-91-202', 'HI.4803.003.N712---N502.LL2013_054')}

parents_filename = {'MSH-604':'HI.4803.003.N712---N504.MSH-604',
                   'UWOPS-91-202':'HI.4803.003.N711---N507.UWOPS-91-202',
                   'LL2012_021':'HI.4803.003.N712---N503.LL2012_021',
                   'LL2012_028':'HI.4803.003.N712---N508.LL2012_028',
                   'LL2011_004':'HI.4803.003.N712---N505.LL2011_004',
                   'LL2011_009':'HI.4803.003.N711---N508.LL2011_009',
                   'MSH-587-1':'NS.1250.002.N712---N505.MSH-587-1',
                   'LL2011_012':'NS.1250.002.N712---N507.LL2011_012',
                   'LL2011_001':'NS.1250.002.N712---N506.LL2011_001',
                   'YPS644':'HI.4803.003.N712---N506.YPS644',
                   'YPS744':'HI.4803.003.N712---N517.YPS744',
                   'LL2013_040':'HI.4803.003.N712---N507.LL2013_040',
                   'LL2013_054':'HI.4803.003.N712---N502.LL2013_054'}

parents_group = {'MSH-604':'SpB',
                   'UWOPS-91-202':'SpB',
                   'LL2012_021':'SpB',
                   'LL2012_028':'SpB',
                   'LL2011_004':'SpC',
                   'LL2011_009':'SpC',
                   'MSH-587-1':'SpC',
                   'LL2011_012':'SpC',
                   'LL2011_001':'SpC',
                   'YPS644':'SpA',
                   'YPS744':'SpA',
                   'LL2013_040':'Sc',
                   'LL2013_054':'Sc'}

parents_color = {'MSH-604':'red',
                   'UWOPS-91-202':'red',
                   'LL2012_021':'darkred',
                   'LL2012_028':'darkred',
                   'LL2011_004':'dodgerblue',
                   'LL2011_009':'dodgerblue',
                   'MSH-587-1':'midnightblue',
                   'LL2011_012':'midnightblue',
                   'LL2011_001':'midnightblue',
                   'YPS644':'limegreen',
                   'YPS744':'limegreen',
                   'LL2013_040':'dimgrey',
                   'LL2013_054':'dimgrey'}

cross_color = ma_strains.groupby('cross')['cross_color'].apply(lambda x: x.iloc[0]).to_dict()

# %% [code]
# parse list of features that show presence/absence polymorphisms
with open('/mnt/HDD3/mito_nanopore/artificial_genome/pres_abs_poly.txt', 'r') as handle:
    pres_abs_poly = handle.read().splitlines()

# %% [code]
# export lists of bam files for freebayes
for cross, df in ma_strains.sort_values(by=['strain','passage']).groupby('cross'):
    if cross in cross_order:
        filenames = list(parents_dict[cross]) + list(df['filename'].values)
        #with open(f'/mnt/HDD3/mito_ma/freebayes/bam_files_{cross}.txt', 'w') as handle:
        #    handle.write('\n'.join([f'/mnt/HDD3/mito_ma/bam/{i}.mt.rmdup.rg.bam' for i in filenames]))

# %% [markdown]
# Define general functions

# %% [code]
def plot_pval_symbol(p):
    if p<=0.001:
        pdisplay = '***'
    elif p<=0.01:
        pdisplay = '**'
    elif p<=0.05:
        pdisplay = '*'
    else:
        pdisplay = None
    return pdisplay

def plot_pval_text(p):
    if p<=0.001:
        pdisplay = f'{p:.1e}'
    elif p<=0.01:
        pdisplay = f'{p:.4f}'
    elif p<=0.1:
        pdisplay = f'{p:.3f}'
    elif p<=1:
        pdisplay = f'{p:.2f}'
    elif p>1:
        pdisplay = f'{p:.1f}'
    else:
        pdisplay = None
    return pdisplay

# %% [code]
def nan_distance(u,v):
    U = u[(~np.isnan(u)) & (~np.isnan(v))]
    V = v[(~np.isnan(u)) & (~np.isnan(v))]
    return (U != V).sum()/U.shape[0]

# %% [markdown]
# Import genome assemblies and annotations

# %% [code]
GFF = pd.read_csv('/mnt/HDD3/mito_nanopore/mfannot/gff_edit.csv', sep='\t')
GFF.columns = list(range(9)) + list(GFF.columns)[9:]
GFF = GFF.astype({3:int,4:int,'strain_order':int,'annot_width':float,'annot_plot_order':int})

artificial_genome_feat = pd.read_csv('/mnt/HDD3/mito_nanopore/artificial_genome/artificial_genome.features.csv', sep=',')
artificial_genome_feat.columns = list(range(9)) + list(artificial_genome_feat.columns)[9:]

# %% [code]
with open('/mnt/HDD3/mito_nanopore/artificial_genome/artificial_genome.fasta') as handle:
    artificial_genome = SeqIO.read(handle, 'fasta')
ref_genome_length = len(artificial_genome.seq)

# %% [code]
parents_order = GFF.groupby('strain').apply(lambda x: x.iloc[0]['strain_order']).sort_values()

at_dict = GFF.groupby('Name').apply(lambda x: x.iloc[0]['annot_type']).to_dict()
at_dict.update(artificial_genome_feat.groupby('Name').apply(lambda x: x.iloc[0]['annot_type']).to_dict())

genes_to_display = ['atp6',
                    'cob',
                    'atp9',
                    'rps3',
                    'rnl',
                    'cox2',
                    'cox3',
                    'rnpB',
                    'rns',
                    'cox1',
                    'atp8',
                    'trna']

genes_alias = dict(zip(genes_to_display, ['$ATP6$',
                                          '$COB$',
                                          '$OLI1$',
                                          '$VAR1$',
                                          '21S rRNA',
                                          '$COX2$',
                                          '$COX3$',
                                          '$RPM1$',
                                          '15S rRNA',
                                          '$COX1$',
                                          '$ATP8$',
                                          'tRNAs']))
cond_alias = {'YPD_25':'YPD 25°C', 'YPD_37':'YPD 37°C', 'YPEG_25':'YPEG 25°C', 'YPEG_37':'YPEG 37°C'}

# %% [code]
# define aliases for plotting feature names
pres_abs_poly_alias = {}
intron_gene_alias = {'a':'$cox1$', 'b':'$cob$'}
intron_i_alias = {'alpha':r'$\alpha$', 'beta':r'$\beta$', 'gamma':r'$\gamma$'}
for a in pres_abs_poly:
    #genes
    if a in genes_alias:
        pres_abs_poly_alias[a] = genes_alias[a]
    
    #trna
    elif a[:3] == 'trn':
        m = re.match('(?:trn)([ACDEFGHIKLMNPQRSTVWY*]{1})\(([aucg]{3,4})\)', a)
        AA, cd =  m.groups()
        pres_abs_poly_alias[a] = f't{AA}({cd.upper()})Q'
    
    #orf
    elif '_orf' in a:
        a1 = a.split('_')[0]
        if a1 in genes_alias:
            pres_abs_poly_alias[a] = f'{genes_alias[a1]} ORF'
        else:
            m = re.match('([ab])i([1-5])(alpha|beta|gamma){0,1}', a1)
            g,n,i = m.groups()
            if i:
                i = intron_i_alias[i]
            else:
                i = ''
            pres_abs_poly_alias[a] = fr'{intron_gene_alias[g]}-I{n}{i} ORF'
            
    elif a == 'omega':
        pres_abs_poly_alias[a] = '$\omega$'
        
    elif a == 'ORF1':
        pres_abs_poly_alias[a] = 'ORF1'
    
    else:
        m = re.match('([ab])i([1-5])(alpha|beta|gamma){0,1}', a)
        g,n,i = m.groups()
        if i:
            i = intron_i_alias[i]
        else:
            i = ''
        pres_abs_poly_alias[a] = f'{intron_gene_alias[g]}-I{n}{i}'

# %% [code]
#add 500-bp wide windows to artificial genome gff for unannotated regions
other_idx = artificial_genome_feat.shape[0]
i = 1
annot_other = []
while i+500 < ref_genome_length:
    new = pd.Series(['mt_art', 'manual', 'intergenic', i, i+499, f'other{other_idx}', '0.8', 'other', 0],
                   index=[0, 1, 2, 3, 4, 'Name', 'annot_color', 'annot_type', 'annot_plot_order'], name=other_idx)
    annot_other.append(new)
    i += 500
    other_idx += 1
#add final line
new = pd.Series(['mt_art', 'manual', 'intergenic', i, ref_genome_length, f'other{other_idx}', '0.8', 'other', 0],
               index=[0, 1, 2, 3, 4, 'Name', 'annot_color', 'annot_type', 'annot_plot_order'], name=other_idx)
annot_other.append(new)

# %% [code]
artificial_genome_feat = pd.concat([artificial_genome_feat, pd.DataFrame(annot_other)])

# %% [code]
annot_type_order = {'exon':0, 'intron':1, 'orf':2, 'rna_exon':3, 'tRNA':4, 'gc_cluster':5, 'other':6}
annot_type_color = artificial_genome_feat.groupby('annot_type').apply(lambda x: x.iloc[0]['annot_color']).loc[annot_type_order].to_dict()
annot_type_color['other'] = '0.8'
annot_type_alias = dict(zip(['exon','intron','orf','rna_exon','tRNA','gc_cluster','other'],
                           ['Exon','Intron','ORF','RNA exon','tRNA','GC cluster','Unannotated']))

# %% [code]
# Import assemblies
CORR = {}

for s in ma_strains.loc[ma_strains['cross']=='P', 'strain']:
    with open(f'/mnt/HDD3/mito_nanopore/polishing/{s}/{s}.nanopolish.pilon.fasta') as handle:
        CORR[s] = SeqIO.read(handle, 'fasta')

# %% [code]
#export gff annotations for 13 genomes

for s, gff in GFF.groupby('strain'):

    new_col8 = []
    for i in gff['Name'].values:
        if i in pres_abs_poly_alias:
            new = pres_abs_poly_alias[i].replace('\\','').replace('$','')
        else:
            new = i
        new_col8.append(f'ID={new};Name={new};transl_table=3;gene={new}')
    gff[8] = new_col8
    gff[2] = gff['annot_type']
    gff[0] = f'{s}.mt_genome'
    gff = gff[range(9)].astype(str).values
    
    if 'LL2013' in s:
        species = 'Saccharomyces cerevisiae'
    else:
        species = 'Saccharomyces paradoxus'
          
    #parse genome length and add to the file header
    
    seq_id = f'{s}.mt_genome [organism={species}] complete mitochondrial genome of strain {s}'
    seq = SeqRecord.SeqRecord(seq=CORR[s].seq, id=seq_id, description='')
    
    #with open(f'/mnt/HDD3/mito_nanopore/final_assemblies/{s}.mt_genome.fasta', 'w') as handle:
    #    SeqIO.write(seq, handle, 'fasta')
    
    header = f'##gff-version 3\n##sequence-region 1 {len(seq.seq)}\n'
    content = '\n'.join(['\t'.join(line) for line in gff])
    
    #write
    #with open(f'/mnt/HDD3/mito_nanopore/final_assemblies/{s}.mt_genome.gff', 'w') as handle:
    #    handle.write(header+content)

# %% [markdown]
# Import vcf files for mtDNAs 

# %% [code]
# import vcf files
def parse_vcf(vcf_file, split_multi=True):

    skiprows = 0
    with gzip.open(vcf_file, 'rb') as handle:
        line = handle.readline()
        while line.decode('utf-8')[:2] == '##':
            skiprows += 1
            line = handle.readline()

    vcf = pd.read_csv(vcf_file, sep='\t', compression='gzip', skiprows=skiprows)
    samples = vcf.columns[9:]

    # attribute unique IDs to variants
    vcf['var_uid'] = [f'v{i}' for i in range(vcf.shape[0])]
    vma_idx = 0
    for vma, df in vcf.groupby(['#CHROM','POS','REF']):
        vcf.loc[df.index, 'vma_uid'] = f'vma{vma_idx}'
        vma_idx += 1

    vcf.index = vcf['var_uid'].values
    # melt vcf
    vcf_melt = pd.melt(vcf, id_vars=['var_uid','vma_uid'], value_vars=samples, var_name='filename', value_name='GT')

    Formats = set(vcf['FORMAT'])
    if len(Formats) == 1:
        Format = list(Formats)[0].split(':')
        for i,tag in enumerate(Format):
            new = vcf_melt['GT'].apply(lambda x: x.split(':')[i]).replace('.', np.nan)
            if tag in ['GT','DP','RO','AO','QR','QA']:
                new = new.astype(float)
            vcf_melt[tag.lower()] = new

    # get counts of supporting observations, and drop genotypes called with <=1 obs
    for a, df in vcf_melt.groupby('gt'):
        if a == 0:
            vcf_melt.loc[df.index, 'obs'] = df['ro']
            vcf_melt.loc[df.index, 'ratio'] = df['ro']/df['dp']
        elif a == 1:
            vcf_melt.loc[df.index, 'obs'] = df['ao']
            vcf_melt.loc[df.index, 'ratio'] = df['ao']/df['dp']
    vcf_melt.loc[vcf_melt['obs']<=1, 'gt'] = np.nan

    vcf_melt['a_ratio'] = vcf_melt['ao']/vcf_melt['dp']
    vcf_melt['r_ratio'] = vcf_melt['ro']/vcf_melt['dp']
    vcf_melt['#CHROM'] = vcf.loc[vcf_melt['var_uid'], '#CHROM'].values
    vcf_melt['POS'] = vcf.loc[vcf_melt['var_uid'], 'POS'].values
    vcf_melt['REF'] = vcf.loc[vcf_melt['var_uid'], 'REF'].values
    vcf_melt['ALT'] = vcf.loc[vcf_melt['var_uid'], 'ALT'].values

    vcf = vcf.drop(samples, axis=1)

    return vcf, vcf_melt

# %% [code]
VCF = {}
VCF_MELT = {}

for cross in cross_order:

    vcf, vcf_melt = parse_vcf(f'/mnt/HDD3/mito_ma/freebayes/freebayes.{cross}.breakmulti.vcf.gz')
    VCF[cross] = vcf
    VCF_MELT[cross] = vcf_melt

# %% [markdown]
# Liftover procedure from whole genome alignments

# %% [code]
# parse maf alignments

def parse_maf(s, path):

    #s_mugsy = f'{s.replace("-","_")}.mt_pilon'
    aln = list(AlignIO.parse(path, 'maf'))
    
    maf = {ai:{seq.id:seq for seq in a} for (ai,a) in enumerate(aln) if len(a)==2}
    # flip any alignment block for which the ref sequence is reversed
    for ai, a in maf.items():
        seq_names = a.keys()
        ref_name = 'artificial_genome.mt_art'
        qry_name = [i for i in seq_names if i!=ref_name][0]
        ref = a[ref_name]
        qry = a[qry_name]
        ref_strand = ref.annotations['strand']
        qry_strand = qry.annotations['strand']
        
        if ref_strand == -1:
            ref.annotations['start'] = ref.annotations['srcSize'] - (ref.annotations['start'] + ref.annotations['size'])
            for seq in (ref, qry):
                seq.seq = seq.seq.reverse_complement()
            
                seq.annotations['strand'] = seq.annotations['strand']*(-2) # mark the flipped sequences as 2
                maf[ai][seq.id] = seq
            
        if qry_strand == -1:
            qry.annotations['start'] = qry.annotations['srcSize'] - (qry.annotations['start'] + qry.annotations['size'])
    
    return maf

def liftover_maf(maf):
    
    lift_list = []

    for ai, a in maf.items():

        ref = a['artificial_genome.mt_art']
        al = len(ref.seq)
        ref_len = ref.annotations['size']
        ref_start = ref.annotations['start']
        ref_offset = 0
        while ref.seq[ref_offset] == '-':
            ref_offset += 1

        lift = range(al)
        ref_pos = []
        for i in lift:
            # build ref positions
            if ref.seq[i] == '-':
                ref_pos.append(-1)
            else:
                ref_pos.append(ref_start)
                ref_start += 1

        ref_pos = np.array(ref_pos)

        lift_list.append(pd.DataFrame([ref_pos,lift,np.repeat(ai, al)],
                                      index=['ref_pos','lift','aln']).T.astype(np.int32))

    lift_list = pd.concat(lift_list).reset_index(drop=True)
    lift_list.index = lift_list['ref_pos'].values
    
    return lift_list

# %% [code]
MAF = {}
for s in ma_strains.loc[ma_strains['cross']=='P', 'strain']:
    MAF[s] = parse_maf(s, f'/mnt/HDD3/mito_nanopore/mugsy/{s}/mt_art.maf')

# %% [code]
LIFT = {}
for s in ma_strains.loc[ma_strains['cross']=='P', 'strain']:
    LIFT[s] = liftover_maf(MAF[s])

# %% [code]
# check the lift procedure
for s, lift in LIFT.items():
    for a, df in lift.groupby('aln'):
        df1 = df.loc[df['ref_pos']!=-1]
        test1 = np.array([artificial_genome.seq[i] for i in df1['ref_pos']])
        test2 = np.array([MAF[s][a]['artificial_genome.mt_art'][i] for i in df1['lift']])
        if np.all(test1 == test2):
            print(f'pass {s} {a}')
        else:
            print(f'error {s} {a}')

# %% [markdown]
## Compute identity levels among parental strains per annotation type

# %% [code]
#define interval tree to query positions for genomic features
gft = intervaltree.IntervalTree()
for i in artificial_genome_feat.index:
    start, end = artificial_genome_feat.loc[i, [3,4]].values
    gft[start:end+1] = i

# %% [code]
# categorize genome positions and compute identity levels

ref_genome_func = []
idx = 0
with ProgressBar(max_value=ref_genome_length) as bar:
    for pos in np.arange(ref_genome_length)+1:
        F = gft[pos]
        if F:
            at, name = artificial_genome_feat.loc[[f[2] for f in F]].sort_values(by='annot_plot_order', ascending=False).iloc[0][['annot_type','Name']].values
            ref_genome_func.append([pos, at, name])
        else:
            ref_genome_func.append([pos, 'other', np.nan])
        idx += 1
        bar.update(idx)

ref_genome_func = pd.DataFrame(ref_genome_func, columns=['pos', 'annot_type', 'name'])
ref_genome_func.set_index('pos', inplace=True, drop=False)
ref_genome_func.loc[0] = np.nan

# %% [code]
MAF_IDENTITY = []

for s, maf in MAF.items():
    lift = LIFT[s].copy()
    lift['annot_type'] = ref_genome_func.loc[lift['ref_pos']+1, 'annot_type'].values
    lift['name'] = ref_genome_func.loc[lift['ref_pos']+1, 'name'].values
    s_mugsy = f'{s.replace("-","_")}.mt_pilon'
    for (at, name, ai), df in lift.groupby(['annot_type', 'name', 'aln']):
        ref_pos = df.index
        alt_pos = lift.loc[ref_pos, 'lift']
        alleles = [maf[ai][s_mugsy].seq[i] for i in alt_pos]
        MAF_IDENTITY.extend([[s, i, at, name, seq] for (i, seq) in zip(ref_pos, alleles)])

MAF_IDENTITY = pd.DataFrame(MAF_IDENTITY, columns=['strain','pos','annot_type','name','allele'])

# %% [code]
# exclude missing introns and ORFs from the MAF_IDENTITY
intron_presence_dict = GFF.loc[GFF['annot_type'].isin(['orf','intron'])].groupby('strain').apply(lambda x: set(x['Name'])).to_dict()

for (s, n), df in MAF_IDENTITY.loc[MAF_IDENTITY['annot_type'].isin(['orf','intron'])].groupby(['strain', 'name']):
    #eliminate bi3, because the non-orf portion of the intron is very small in all but SpA and Sc
    if (n not in intron_presence_dict[s]) or (n == 'bi3'):
        MAF_IDENTITY.drop(df.index, axis=0, inplace=True)

# %% [code]
# compute sequence identity per annotation type
at_ident = []
for (at, name), df in MAF_IDENTITY.groupby(['annot_type','name']):
    df = df.pivot_table(index='pos', columns='strain', values='allele', aggfunc=lambda x: x)
    for cross, parents in parents_dict.items():
        parents = [p.split('.')[-1] for p in parents]
        p1, p2 = parents
        if all([p in df.columns for p in parents]):
            equal = df.loc[(~df[p1].isna()) & (~df[p2].isna())].apply(lambda x: x[p1]==x[p2], axis=1).value_counts()
            equal.index = [str(x) for x in equal.index]
            for b in ('True', 'False'):
                if b not in equal.index:
                    equal.loc[b] = 0
            at_ident.append([at, name, cross, equal.loc['True'], equal.loc['False']])

at_ident = pd.DataFrame(at_ident, columns=['annot_type','name','cross','true','false'])
at_ident['identity'] = at_ident['true']/(at_ident['true']+at_ident['false'])

# %% [markdown]
# Annotation summary
## Fig 1C

# %% [code]
fig = plt.figure(figsize=[10,5])
gs = plt.GridSpec(ncols=1, nrows=2, height_ratios=[7,1], hspace=0.7,
                 left=0.12, bottom=0.1, right=0.94, top=0.98)

at_color_hm = artificial_genome_feat.groupby('annot_type').apply(lambda x: x.iloc[0]['annot_color']).to_dict()

ax1 = fig.add_subplot(gs[0])
pap_pos = {j:i for i,j in enumerate(pres_abs_poly)}
for (s, name, at), df in GFF.loc[GFF['Name'].isin(pres_abs_poly)].groupby(['strain','Name','annot_type']):
    #if at in plot_features:
    mfc = at_color_hm[at]
    mec = 'white'
    lw = 0
    if mfc == 'white':
        mec = 'black'
        lw = 1

    ax1.scatter(pap_pos[name], parents_order[s], marker='s', s=54, color=mfc, edgecolors=mec, linewidths=lw, zorder=1)

for group, (y, dy), fc in zip(['SpC','SpB','SpA','S.cer'], [(0,5), (5,4), (9,2), (11,2)], ['#00008B','#EE0000','#00CD00','0.25']):
    rect_spacing = 0.05
    Rect = Rectangle((-1, y-0.5+rect_spacing), 65, dy-2*rect_spacing, fc=fc, lw=0, zorder=0, clip_on=False, alpha=0.15)
    ax1.add_patch(Rect)
    mid = np.mean([y, dy+y])
    ax1.text(64.5, mid-0.5, group, color=fc, size=11, fontweight='semibold', fontstyle='italic',
             va='center', ha='left', clip_on=False, zorder=1)

# add labels to ax1
for i, a in enumerate(pres_abs_poly):
    fc = at_color_hm[at_dict[a]]
    ec = 'white'
    tc = 'white'
    if fc == 'white':
        tc = 'black'
        ec = 'black'
    if fc == 'limegreen':
        tc = 'black'

    t = ax1.text(i, 13, pres_abs_poly_alias[a], rotation=90, ha='center', va='top', color=tc, size=7,
                 bbox=dict(ec=ec, fc=fc, lw=0.5, boxstyle='square,pad=0.12'), zorder=1)
    if a in artificial_genome_feat['Name'].values:
        ax1.plot([i,i], [13,16.5], lw=0.5, color='black', zorder=-1, clip_on=False)

ax1.set_xticks([])
ax1.set_xlim(-1,64)   

ax1.set_ylim(-0.5, 12.5)
ax1.set_yticks(parents_order)
ax1.set_yticklabels(parents_order.index)
ax1.invert_yaxis()

for i in ['left','bottom','right','top']:
    ax1.spines[i].set_visible(False)

# plot GFF entries
ax2 = fig.add_subplot(gs[1])
ax2.arrow(0, 1, ref_genome_length, 0, color='0.9', width=1, head_length=0, head_width=0, zorder=-1)

for i in artificial_genome_feat.loc[~artificial_genome_feat['annot_type'].isin(['gc_cluster','other'])].sort_values(by='annot_plot_order', ascending=False).index:
    start, end, color, w, z = artificial_genome_feat.loc[i, [3,4,'annot_color','annot_width','annot_plot_order']]
    ax2.arrow(start, 1, end-start, 0, color=color, width=w, head_length=0, head_width=0, zorder=z)

ax2.set_xlim(0,ref_genome_length)
ax2.set_xticks(np.arange(0,83e3,10e3))
ax2.set_xticklabels(np.arange(0,83,10))
ax2.set_xlabel('kb')
ax2.set_ylim(0.5, 1.5)
ax2.set_yticks([])

for i in ['left','right','top']:
    ax2.spines[i].set_visible(False)

#link items from one panel to the other
for i, a in enumerate(pres_abs_poly):
    if a in artificial_genome_feat['Name'].values:
        pos = artificial_genome_feat.loc[artificial_genome_feat['Name']==a].iloc[0,3]#[[3,4]].mean()
        (x_ax1, y1_ax1), (x_ax1, y2_ax1)  = ax1.transData.transform([(i, 16.5), (i, 13)])
        (x_ax1, y1_ax1), (x_ax1, y2_ax1) = ax2.transData.inverted().transform([(x_ax1, y1_ax1), (x_ax1, y2_ax1)])

        ax2.plot([pos, x_ax1], [1.5, y1_ax1], lw=0.5, color='black', zorder=-1, clip_on=False)


#plt.savefig(f'/home/mathieu/mhenault_landrylab/Publications/mito_ma/resubmission2_GRes/fig/Fig1C.svg', dpi=300)
#plt.show()
plt.close()

# %% [markdown]
## Fig 1A

# %% [code]
#plot crosses scheme
fig = plt.figure(figsize=[4.5,4])
gs = plt.GridSpec(ncols=1, nrows=1, left=0.2, right=0.77, top=0.98, bottom=0.3)
ax = fig.add_subplot(gs[0])

cm_hm_cross = ListedColormap([cross_color[cross] for cross in cross_order], 11)
hm = np.ndarray([13,13])
hm[:] = np.nan

for cross in parents_dict:
    p1, p2 = [p.split('.')[-1] for p in parents_dict[cross]]
    x, y = parents_order[p1], parents_order[p2]
    
    if cross in ['CC1', 'CC2', 'CC3']:
        nrep = 64
    elif cross in ['BB1', 'BB2']:
        nrep = 48
    else:
        nrep = 96
        
    if x > y:
        hm[x][y] = cross_order[cross]
        ax.text(y+1.6, x+0.5, fr'{cross} ($\times${nrep})', size=9, color='k', ha='left', va='center', zorder=3)
        ax.plot([y+1, y+1.5], [x+0.5, x+0.5], color='k', lw=1, zorder=3)

    else:
        hm[y][x] = cross_order[cross]
        ax.text(x+1.6, y+0.5, fr'{cross} ($\times${nrep})', size=9, color='k', ha='left', va='center', zorder=3)
        ax.plot([x+1, x+1.5], [y+0.5, y+0.5], color='k', lw=1, zorder=2)
    
P = Polygon(np.array([[0,13,0,0], [13,13,0,13]]).T, fc='white', zorder=1)
ax.add_patch(P)

for group, (y, dy), fc in zip(['SpC','SpB','SpA','S.cer'], [(0,5), (5,4), (9,2), (11,2)], ['#00008B','#EE0000','#00CD00','0.25']):
    rect_spacing = 0.05
    
    Rect = Rectangle((0, y+rect_spacing), 13, dy-2*rect_spacing, fc=fc, lw=0, zorder=0, clip_on=False, alpha=0.3)
    ax.add_patch(Rect)
    mid = np.mean([y, dy+y])
    ax.text(13.1, mid, group, color=fc, size=11, fontweight='semibold', fontstyle='italic',
             va='center', ha='left', clip_on=False, zorder=0)


ax.pcolormesh(hm, cmap=cm_hm_cross, zorder=3)
ax.set_xlim(0,13)
ax.set_xticks(np.arange(13)+0.5)
ax.set_xticklabels([])
ax.set_yticks(np.arange(13)+0.5)
ax.set_yticklabels([])

for p in parents_order.index:
    if p in ['LL2011_004', 'LL2011_009', 'MSH-604', 'UWOPS-91-202']:
        fw = 'bold'
    else:
        fw = 'normal'
    ax.text(parents_order.loc[p]+0.5, 13.5, p, color=parents_color[p], rotation=45, size=10, ha='right', va='top', weight=fw)
    ax.text(-0.5, parents_order.loc[p]+0.5, p, color=parents_color[p], ha='right', va='center', weight=fw)

ax.set_xlim(0,13)
ax.set_ylim(0,13)
ax.invert_yaxis()

for i in ['right','top']:
    ax.spines[i].set_visible(False)


#plt.savefig(f'/home/mathieu/mhenault_landrylab/Publications/mito_ma/resubmission2_GRes/fig/Fig1A.svg', dpi=300)
#plt.show()
plt.close()

# %% [markdown]
# MtDNA Recombination
## Definition of parental marker variants

# %% [code]
def test_marker_lift(cross, vma, ps, variant_type):
    p_alt, p_ref = np.repeat(np.nan, 2)
    
    if variant_type == 'single':
        vcf = VCF[cross].set_index(VCF[cross]['vma_uid'].values)
        pos1, ref, alt = vcf.loc[vma, ['POS','REF','ALT']].values
        # correct pos for 0-based
        pos1 -= 1    
        pos2 = pos1 + len(ref)
        # parse which parent has the ref and alt alleles
        p_alt, p_ref = np.repeat(np.nan, 2)
        for s in parents_dict[cross]:
            if ps[1] == s:
                p_alt = s.split('.')[-1]
            elif ps[0] == s:
                p_ref = s.split('.')[-1]
                
    elif variant_type == 'multi':
        vcf = VCF[cross].set_index(VCF[cross]['var_uid'].values)
        p_ref = ps[0][2].split('.')[-1]
        pos1, real_ref, ref = vcf.loc[ps[0][0], ['POS','REF','ALT']].values
        
        p_alt = ps[1][2].split('.')[-1]
        alt = vcf.loc[ps[1][0], 'ALT']
        # correct pos for 0-based
        pos1 -= 1    
        pos2 = pos1 + len(real_ref)
        
    else:
        raise ValueError('illegal variant type specified')

    #if vma position is not present in both liftovers, return np.nan
    if all([pos1 in LIFT[p_ref].index,
            pos1 in LIFT[p_alt].index,
            pos2 in LIFT[p_ref].index,
            pos2 in LIFT[p_alt].index]):
        
        variants = {}
        
        for s, allele in zip([p_ref, p_alt], [ref, alt]):
            
            s_mugsy = f'{s.replace("-","_")}.mt_pilon'
            maf = MAF[s]
            lift = LIFT[s]
            ai1, ai2 = lift.loc[[pos1, pos2], 'aln']
            if ai1 == ai2:
                lift = lift.loc[lift['aln']==ai1]
                lift_pos1 = lift.loc[pos1, 'lift']
                lift_pos2 = lift.loc[pos2, 'lift']
                lift = lift.set_index(lift['lift'])

                allele_seq = str(maf[ai1][s_mugsy].seq[lift_pos1:lift_pos2]).replace('-','')
                variants[s] = allele_seq == allele
        if len(variants) == 2:
            return all(variants.values())
        else:
            return False
    
    else:
        return False

# %% [code]
PARENTAL_SINGLE = {}
PARENTAL_MULTI = {}

min_depth = 2

for cross in cross_order:
    
    vcf = VCF[cross]
    vcf_melt = VCF_MELT[cross]
    p1, p2 = parents_dict[cross]
    
    # classify parental snps if they can be used to discrimitate parental haplotypes
    vcf_melt_p1 = vcf_melt.loc[vcf_melt['filename']==p1]
    vcf_melt_p1.index = vcf_melt_p1['var_uid']
    vcf_melt_p2 = vcf_melt.loc[vcf_melt['filename']==p2]
    vcf_melt_p2.index = vcf_melt_p2['var_uid']

    parental_single = {}
    parental_multi = {}
    for vma, df in vcf.groupby('vma_uid'):
        # biallelic snp case
        if df.shape[0] == 1:
            v = df.index[0]
            d1, d2 = np.repeat(np.nan, 2)
            

            a1 = vcf_melt_p1.loc[v, 'gt']
            if a1 == 1:
                d1 = vcf_melt_p1.loc[v, 'ao']
            if a1 == 0:
                d1 = vcf_melt_p1.loc[v, 'ro']

            a2 = vcf_melt_p2.loc[v, 'gt']
            if a2 == 1:
                d2 = vcf_melt_p2.loc[v, 'ao']
            if a2 == 0:
                d2 = vcf_melt_p2.loc[v, 'ro']

            if all([a1 != a2,
                    not np.isnan(a1), 
                    not np.isnan(a2), 
                    d1>=min_depth, 
                    d2>=min_depth]) :
                parental_single[vma] = {a1:p1, a2:p2}

        # multiallelic snp case
        else:
            v = df.index
            A1 = vcf_melt_p1.loc[v, 'gt'].values
            A2 = vcf_melt_p2.loc[v, 'gt'].values
            if (not np.all(A1 == A2)) and (1 in A1) and (1 in A2):
                sub1 = vcf_melt_p1.loc[v]
                v1 = sub1.loc[sub1['gt']==1].index[0]
                a1 = vcf_melt_p1.loc[v1, 'gt']
                d1 = vcf_melt_p1.loc[v1, 'ao']

                sub2 = vcf_melt_p2.loc[v]
                v2 = sub2.loc[sub2['gt']==1].index[0]
                a2 = vcf_melt_p2.loc[v2, 'gt']
                d2 = vcf_melt_p2.loc[v2, 'ao']

                if all([a1 != a2 or v1 != v2,
                       d1>=min_depth,
                       d2>=min_depth]):
                    parental_multi[vma] = [(v1,a1,p1), (v2,a2,p2)]
    
    PARENTAL_SINGLE[cross] = parental_single
    PARENTAL_MULTI[cross] = parental_multi

# %% [code]
remove_variants = {}
for cross in cross_order:
    remove_variants[cross] = {}
    for vma, ps in PARENTAL_SINGLE[cross].items():
        test = test_marker_lift(cross, vma, ps, 'single')
        remove_variants[cross][vma] = test
        
    for vma, ps in PARENTAL_MULTI[cross].items():
        test = test_marker_lift(cross, vma, ps, 'multi')
        remove_variants[cross][vma] = test

    # analyze parental markers in presence absence polymorphisms

    not_conserved = []
    parents = [p.split('.')[-1] for p in parents_dict[cross]]
    gff = GFF.loc[GFF['strain'].isin(parents)]
    for name, df in gff.groupby('Name'):
        if not all([p in df['strain'].values for p in parents]):
            not_conserved.append(name)
    
    vcf = VCF[cross].set_index('vma_uid')
    for vma in list(PARENTAL_SINGLE[cross])+list(PARENTAL_MULTI[cross]):
        pos = vcf.loc[[vma]].iloc[0]['POS']
        f = gft[pos]
        if f:
            name = list(f)[0][2]
            name = artificial_genome_feat.loc[name, 'Name']
            if name in not_conserved:
                remove_variants[cross][vma] = False

# %% [code]
for cross in cross_order:
    for vma in remove_variants[cross]:
        if remove_variants[cross][vma] == False:
            if vma in PARENTAL_SINGLE[cross]:
                del PARENTAL_SINGLE[cross][vma]
            if vma in PARENTAL_MULTI[cross]:
                del PARENTAL_MULTI[cross][vma]

# %% [markdown]
## Classify snps

# %% [code]
# input the coverage depth on nuclear chromosomes
idx = 0
depth_nuclear = {}
with ProgressBar(max_value=ma_strains.loc[ma_strains['cross'].isin(cross_order)].shape[0]) as bar:
    
    for s in ma_strains.loc[ma_strains['cross'].isin(cross_order)].index:
        depth = pd.read_csv(f'/mnt/HDD2/seqdata/depth/{s}/{s}_SpB.depth', sep='\t', header=None)
        depth_median = []
        for tig, df in depth.groupby(0):
            if tig[:3]=='utg':
                depth_median.append(df[2])
        depth_median = np.concatenate(depth_median)
        depth_nuclear[s] = [np.median(depth_median), np.mean(depth_median), np.std(depth_median), np.quantile(depth_median, 0.1)]
        idx += 1
        bar.update(idx)

depth_nuclear = pd.DataFrame(depth_nuclear).T
depth_nuclear.columns = ['median','mean','std','q10']
depth_nuclear['thres'] = np.where(depth_nuclear['q10']>=20, depth_nuclear['q10'], 20)

# %% [code]
#add parental values computed separately
for s in ma_strains.loc[ma_strains['cross']=='P'].index:
    depth = pd.read_csv(f'/mnt/HDD3/mito_ma/depth_parents/{s.split(".")[-1]}.depth', sep='\t', header=None)
    depth_nuclear.loc[s, 'median'] = depth[2].median()

# %% [code]
for cross in cross_order:

    vcf_melt = VCF_MELT[cross]
    vcf_melt['marker'] = False
    vcf_melt['parent'] = 'none'

    parental_single = PARENTAL_SINGLE[cross]
    parental_multi = PARENTAL_MULTI[cross]
    
    for vma, df in vcf_melt.groupby('vma_uid'):
        if vma in parental_single:
            for gt,p in parental_single[vma].items():
                idx = df.loc[df['gt'] == gt].index
                vcf_melt.loc[idx, 'marker'] = True
                vcf_melt.loc[idx, 'parent'] = p
        if vma in parental_multi:
            for (v, gt, p) in parental_multi[vma]:
                df1 = df.loc[df['var_uid']==v]
                idx = df1.loc[df1['gt'] == gt].index
                vcf_melt.loc[idx, 'marker'] = True
                vcf_melt.loc[idx, 'parent'] = p
                
    # add whole-genome depth information
    for s, df in vcf_melt.groupby('filename'):
        # exclude parents
        if ma_strains.loc[s, 'cross'] == cross:
            vcf_melt.loc[df.index, 'depth_nuclear'] = depth_nuclear.loc[s,'thres']
            
    #add binary filter for whether support is higher than nuclear depth
    for gt, df in vcf_melt.groupby('gt'):
        if gt == 1:
            vcf_melt.loc[df.index, 'higher'] = df['ao'] >= df['depth_nuclear']
        if gt == 0:
            vcf_melt.loc[df.index, 'higher'] = df['ro'] >= df['depth_nuclear']
    
    vcf_melt['line'] = ma_strains.loc[vcf_melt['filename'], 'strain'].values
    vcf_melt['passage'] = ma_strains.loc[vcf_melt['filename'], 'passage'].values
    
    VCF_MELT[cross] = vcf_melt

# %% [code]
# make a dictionary of the markers for each cross
MARKERS = {}

for cross in cross_order:
    vcf_melt = VCF_MELT[cross]
    vcf_melt = vcf_melt.loc[vcf_melt['marker']]
    MARKERS[cross] = sorted(set(vcf_melt['vma_uid']))

# %% [markdown]
## Determine clonal mt haplotypes

# %% [code]
# compute distance between haplotypes within lines based on markers
distance = []
for cross in cross_order:
    parents_mapping_dict = {j:i for i,j in enumerate(parents_dict[cross])}
    vcf_melt = VCF_MELT[cross]
    vcf_melt = vcf_melt.loc[(vcf_melt['marker']) & (vcf_melt['higher'])]
    vcf_melt = vcf_melt.pivot_table(index='var_uid', columns='filename', values='parent', aggfunc=lambda x: x)
    for line, df in ma_strains.loc[(ma_strains['cross']==cross) &
                                   (ma_strains['filename'].isin(vcf_melt.columns)) & 
                                   (ma_strains['identity_filter'])].sort_values(by='passage').groupby('strain'):

        profiles = vcf_melt.loc[:, df['filename']]
        if profiles.shape[1] > 1:
            for (i,j) in itertools.combinations(profiles.columns, 2):
                u = profiles[i].replace(parents_mapping_dict).values
                v = profiles[j].replace(parents_mapping_dict).values

                d = nan_distance(u,v)
                distance.append([cross, line, i, j, 1-d])
        elif profiles.shape[1] == 1:
            fn = profiles.columns[0]
            distance.append([cross, line, fn, fn, 1])

distance = pd.DataFrame(distance, columns=['cross','line','filename1','filename2','dist'])

# %% [code]
# get lines to examine manually
mthap_manual_examination = []
ma_strains['mt_haplotype'] = 'none'
for s, df in distance.groupby('line'):
    if df.shape[0] == 1:
        fn1 = df.iloc[0]['filename1']
        fn2 = df.iloc[0]['filename2']
        if df.iloc[0]['dist'] >= 0.95:
            ma_strains.loc[fn1, 'mt_haplotype'] = 'clone'
            ma_strains.loc[fn2, 'mt_haplotype'] = s
        else:
            ma_strains.loc[fn1, 'mt_haplotype'] = fn1.split('.')[-1]
            ma_strains.loc[fn2, 'mt_haplotype'] = fn2.split('.')[-1]
    else:
        mthap_manual_examination.append(s)

# %% [code]
# manual examination
fig, axes = plt.subplots(ncols=3, nrows=3, figsize=[10,10])
for (i,j), s in zip(itertools.product(range(3), range(3)), mthap_manual_examination):
    ax = axes[i,j]
    dat = distance.loc[distance['line']==s].copy()
    dat['s1'] = dat['filename1'].apply(lambda x: ' '.join([x.split('.')[i] for i in (0,-1)]))
    dat['s2'] = dat['filename2'].apply(lambda x: ' '.join([x.split('.')[i] for i in (0,-1)]))
    sns.heatmap(dat.pivot_table(index='s1', columns='s2', values='dist'), vmin=0.9, vmax=1, cmap='bwr', ax=ax)
    ax.set_title(s)

plt.tight_layout()
#plt.show()
plt.close()

# %% [code]
# manual examination of allele ratios
cross = 'BA1'
vcf_melt = VCF_MELT[cross]
test = {}

for s, dat in vcf_melt.groupby('filename'):
    
    ratios = np.where(dat['gt']==0, dat['r_ratio'], dat['a_ratio'])
    test[s] = ratios[(ratios>0) & (ratios<0.9)].shape[0]/ratios.shape[0]

    c = 'black'
    z = 1
    if s == 'HI.4803.002.N703---N508.B2_P35':
        c = 'red'
        z = 10

    plt.plot(np.sort(ratios), np.linspace(0,1,ratios.shape[0]), label=s, c=c, alpha=0.3, zorder=z)

plt.title(cross)
plt.xlim(0.8,1)
plt.ylim(0,0.4)
#plt.show()
plt.close()

# %% [code]
# low allelic ratios to exclude
low_ratio = ['NS.1250.001.N704---N507.J40_P35',
             'HI.4802.003.N711---N503.I7_P1',
             'HI.4803.001.N704---N504.I25_P35',
             'HI.4803.001.N708---N504.D85_P35',
             'HI.4803.002.N703---N508.B2_P35',
             'HI.4803.001.N708---N506.B40_P35',
             'HI.4803.001.N706---N506.F81_P35']

# %% [code]
# manually process lines for which there is more than P1 and P35
ma_strains.loc[ma_strains['strain'].isin(mthap_manual_examination), 'mt_haplotype'] = 'clone'
ma_strains.loc['NS.1250.002.N704---N504.B32_P35', 'mt_haplotype'] = 'B32'
ma_strains.loc['NS.1249.002.N712---N508.B40_P16', 'mt_haplotype'] = 'B40_P16'
ma_strains.loc['HI.4803.001.N708---N506.B40_P35', 'mt_haplotype'] = 'B40_P35_HI'
ma_strains.loc['NS.1250.002.N704---N507.B40_P35', 'mt_haplotype'] = 'B40_P35_NS'
ma_strains.loc['HI.4803.003.N701---N502.B49_P35', 'mt_haplotype'] = 'B49'
ma_strains.loc['NS.1249.002.N710---N504.B55_P1', 'mt_haplotype'] = 'B55_P1'
ma_strains.loc['NS.1250.002.N705---N508.B55_P35', 'mt_haplotype'] = 'B55_P35'
ma_strains.loc['HI.4803.002.N712---N504.D36_P35', 'mt_haplotype'] = 'D36'
ma_strains.loc['HI.4803.003.N703---N508.F38_P25', 'mt_haplotype'] = 'F38'
ma_strains.loc['NS.1250.001.N711---N503.F43_P35', 'mt_haplotype'] = 'F43'
ma_strains.loc['NS.1250.002.N709---N508.F57_P35', 'mt_haplotype'] = 'F57'
# manually switch clonal haplotypes
ma_strains.loc['HI.4802.002.N712---N517.A62_P1', 'mt_haplotype'] = 'A62'
ma_strains.loc['HI.4803.003.N703---N502.A62_P35', 'mt_haplotype'] = 'clone'
ma_strains.loc['HI.4802.002.N702---N508.F3_P1', 'mt_haplotype'] = 'F3'
ma_strains.loc['HI.4803.003.N705---N505.F3_P35', 'mt_haplotype'] = 'clone'
ma_strains.loc['HI.4802.002.N707---N502.C46_P1', 'mt_haplotype'] = 'C46'
ma_strains.loc['HI.4803.003.N711---N505.C46_P35', 'mt_haplotype'] = 'clone'
ma_strains.loc['NS.1249.002.N701---N506.J29_P1', 'mt_haplotype'] = 'J29'
ma_strains.loc['NS.1250.001.N703---N503.J29_P35', 'mt_haplotype'] = 'clone'
ma_strains.loc['NS.1249.001.N707---N505.J15_P1', 'mt_haplotype'] = 'J15'
ma_strains.loc['NS.1250.002.N703---N503.J15_P35', 'mt_haplotype'] = 'clone'
ma_strains.loc['NS.1249.002.N703---N508.J54_P1', 'mt_haplotype'] = 'J54'
ma_strains.loc['NS.1250.001.N712---N504.J54_P35', 'mt_haplotype'] = 'clone'
ma_strains.loc['NS.1249.001.N712---N517.J64_P1', 'mt_haplotype'] = 'J64'
ma_strains.loc['NS.1250.002.N712---N502.J64_P35', 'mt_haplotype'] = 'clone'
ma_strains.loc['NS.1249.001.N712---N507.L5_P1', 'mt_haplotype'] = 'L5'
ma_strains.loc['NS.1250.002.N706---N503.L5_P35', 'mt_haplotype'] = 'clone'
ma_strains.loc['NS.1249.002.N705---N504.L46_P1', 'mt_haplotype'] = 'L46'
ma_strains.loc['NS.1250.002.N711---N517.L46_P35', 'mt_haplotype'] = 'clone'
ma_strains.loc['HI.4802.001.N707---N507.B54_P1', 'mt_haplotype'] = 'B54'
ma_strains.loc['HI.4803.003.N703---N505.B54_P35', 'mt_haplotype'] = 'clone'
ma_strains.loc['HI.4802.001.N711---N504.E34_P1', 'mt_haplotype'] = 'E34'
ma_strains.loc['HI.4803.001.N705---N505.E34_P35', 'mt_haplotype'] = 'clone'
ma_strains.loc['HI.4802.003.N705---N505.E37_P1', 'mt_haplotype'] = 'E37'
ma_strains.loc['HI.4803.002.N707---N508.E37_P35', 'mt_haplotype'] = 'clone'
ma_strains.loc['HI.4802.001.N703---N503.C87_P1', 'mt_haplotype'] = 'C87'
ma_strains.loc['HI.4803.003.N707---N502.C87_P35', 'mt_haplotype'] = 'clone'
ma_strains.loc['HI.4802.001.N707---N508.F15_P1', 'mt_haplotype'] = 'F15'
ma_strains.loc['HI.4803.001.N710---N503.F15_P35', 'mt_haplotype'] = 'clone'

# %% [code]
# combine info of clonal mthap with ones with low allele support ratio
ma_strains['mthap_filter'] = np.where((ma_strains['mt_haplotype'].isin(['clone','none']))
                                      | (ma_strains['filename'].isin(low_ratio)), False, True)
ma_strains['mtdel_filter'] = np.where((ma_strains['mt_haplotype']=='none')
                                      | (ma_strains['filename'].isin(low_ratio)), False, True)
ma_strains['mt_hap_del'] = np.nan
for (s,p), df in ma_strains.loc[ma_strains['mtdel_filter']].groupby(['strain','passage']):
    if df.shape[0] == 1:
        ma_strains.loc[df.index, 'mt_hap_del'] = f'{s}_P{p}'
    else:
        ma_strains.loc[df.index, 'mt_hap_del'] = np.where(df['mt_haplotype']=='clone', df['filename'].apply(lambda x: x.split('.')[-1]), df['mt_haplotype'])

# %% [code]
# add mt_hap_del ids to lines with no mtDNA variants ('none' in mt_haplotype)
df = ma_strains.loc[(ma_strains['mt_haplotype']=='none')
               & (ma_strains['identity_filter']==True)
               & (ma_strains['cross'].isin(cross_order))]
ma_strains.loc[df.index, 'mt_hap_del'] = df['strain_passage']
ma_strains.loc[df.index, 'mtdel_filter'] = True

df = ma_strains.loc[ma_strains['cross']=='P']
ma_strains.loc[df.index, 'mt_hap_del'] = df['strain']

# %% [markdown]
## Final export of MA strains metadata with haplotype info

# %% [code]
#final export
#ma_strains.to_csv('/mnt/HDD3/mito_ma/script/ma_strains_mthap.tsv', sep='\t')
ma_strains = pd.read_csv('/mnt/HDD3/mito_ma/script/ma_strains_mthap.tsv', sep='\t', index_col=0)

# %% [code]
strains_mthap = ma_strains.loc[ma_strains['mthap_filter'], 'filename']
strains_mtdel = ma_strains.loc[ma_strains['mtdel_filter'], 'filename']

# %% [markdown]
## Sort lines by similarity of marker variants

# %% [code]
# sort lines by marker variants profile similarity
STRAIN_ORDER_MARKERS = {}

for cross in cross_order:

    strains_to_include = ma_strains.loc[(ma_strains['mthap_filter']) & 
                                        (ma_strains['cross']==cross), 'filename']
    parents_mapping_dict = {j:i for i,j in enumerate(parents_dict[cross])}
    vcf_melt = VCF_MELT[cross]
    vcf_melt = vcf_melt.loc[(vcf_melt['marker']) & 
                            (vcf_melt['vma_uid'].isin(MARKERS[cross])) & 
                            (vcf_melt['filename'].isin(strains_to_include))]
    vcf_melt = vcf_melt.pivot_table(index='var_uid', columns='filename', values='parent', aggfunc=lambda x: x).replace(parents_mapping_dict)

    dendro = hierarchy.dendrogram(hierarchy.linkage(vcf_melt.T.values, metric=nan_distance), no_plot=True)
    strain_order = [vcf_melt.columns[i] for i in dendro['leaves']]

    STRAIN_ORDER_MARKERS[cross] = strain_order

# %% [code]
#manually flip the order of some crosses
for cross in ['CC2','CC3','BB1','BC2','BA1','BSc1']:
    STRAIN_ORDER_MARKERS[cross] = STRAIN_ORDER_MARKERS[cross][::-1]

# %% [markdown]
## Table S3

# %% [code]
consistent_hap = []
for (cross, s), df in ma_strains.loc[(ma_strains['passage'].isin([1, 35])) &
                                    (ma_strains['cross'].isin(cross_order))].sort_values(by='passage').groupby(['cross','strain']):
    if df.shape[0] == 2:
        if np.all(df['passage'].values == [1, 35]):
            if 'clone' in df['mt_haplotype'].values:
                cst = 'consistent'
            else:
                cst = 'inconsistent'
            consistent_hap.append([cross, s, cst])
consistent_hap = pd.DataFrame(consistent_hap, columns=['cross', 'strain', 'consistent'])
cons_hap_pivot = consistent_hap.value_counts(['cross','consistent'], sort=False).rename('count').reset_index().\
pivot_table(index='cross', columns='consistent', values='count', aggfunc=lambda x: x).loc[cross_order]
cons_hap_pivot['percent_inconsistent'] = cons_hap_pivot['inconsistent']/(cons_hap_pivot['consistent']+cons_hap_pivot['inconsistent'])*100
#cons_hap_pivot.to_csv('/mnt/HDD3/mito_ma/results/cons_hap_pivot.csv')

# %% [markdown]
## Fig S3

# %% [code]
#bins = np.linspace(0, 0.5, 101)
bins = [0, 0.005, 0.05, 0.5]

minor_af_hist = []

for (cross, p), df in ma_strains.loc[(ma_strains['identity_filter']==True) &
                                     (ma_strains['cross'].isin(cross_order)) &
                                     (ma_strains['passage'].isin([1,35]))].groupby(['cross', 'passage']):


    vcf_melt = VCF_MELT[cross]
    vcf_melt = vcf_melt[(vcf_melt['higher']==True) &
                        (~vcf_melt['ratio'].isna())].set_index('filename')

    S = df.index
    S = [s for s in S if s in vcf_melt.index]
    

    ratio = vcf_melt.loc[S, ['a_ratio', 'r_ratio']].min(axis=1)
    h = np.histogram(ratio, bins=bins)[0]/ratio.shape[0]

    h = pd.Series(h, index=[f'[{bins[i]}, {bins[i+1]})' for i in range(3)], name=cross).rename('freq').reset_index()
    h['cross'] = cross
    h['passage'] = p
    minor_af_hist.append(h)
        
minor_af_hist = pd.concat(minor_af_hist)

# %% [code]
fig = plt.figure(figsize=[7,7])
gs = plt.GridSpec(ncols=4, nrows=3, hspace=0.8, wspace=1,
                  bottom=0.12, right=0.97, top=0.96, left=0.12)
cross_ax = dict(zip(cross_order, itertools.product(range(3), range(4))))
passage_color_dict={1:'0.6', 35:'k'}

for cross in cross_order:
    
    ax = fig.add_subplot(gs[cross_ax[cross]])
    df = minor_af_hist.loc[minor_af_hist['cross']==cross]
    
    sns.barplot(x='index', y='freq', hue='passage', palette=passage_color_dict, data=df)
    ax.legend_.remove()
    plt.xticks(rotation=45, ha='right')
    
    ax.set_xlabel('')
    ax.set_ylim(0, 0.85)
    ax.set_ylabel('proportion of minor\nallele frequencies')
    ax.set_title(cross)
    
ax = fig.add_axes([0.75, 0.18, 0.15, 0.08])
ax.axis('off')
legend_elms = [Line2D([0], [0], color='white', marker='s', ms=9, mfc=mfc, label=l) for (l, mfc) in zip(['initial timepoint', 'final timepoint'], ['0.6', 'k'])]
ax.legend(handles=legend_elms, loc=3, bbox_to_anchor=[0,0], frameon=False)

sns.despine()
#plt.savefig('/home/mathieu/mhenault_landrylab/Publications/mito_ma/resubmission2_GRes/fig/allele_freq.svg', dpi=300)
#plt.show()
plt.close()

# %% [markdown]
## Define tracts of parental ancestry

# %% [code]
# find tracts of parental ancestry
def make_tracts(cross, markers):

    TRACTS = []

    vcf = VCF[cross]
    vcf_melt = VCF_MELT[cross]
    vcf_melt = vcf_melt.loc[(vcf_melt['marker']) & 
                            (vcf_melt['higher']) &
                            (vcf_melt['vma_uid'].isin(markers))]

    for s, df in vcf_melt.sort_values(by='POS').groupby('filename'):

        tracts = []
        latest = 'init'
        new_tract = []

        for i,idx in enumerate(df.index):
            call = df.loc[idx, 'parent']
            if call != latest:
                #first, go one further and see if it matches the latest
                if i < df.shape[0]-1:
                    next_idx = df.index[i+1]
                    next_call = df.loc[next_idx, 'parent']
                    if next_call == latest:
                        continue

                #if there is an active new tract, dump it first
                if len(new_tract) > 0:
                    tracts.append(new_tract)
                #define new tract with the first index in
                new_tract = [idx]
                #update latest 
                latest = call

            else:
                new_tract.append(idx)
                latest = call
                if i == df.shape[0]-1:
                    tracts.append(new_tract)

        tracts_format = []
        for t in tracts:
            if len(t) > 1:
                pos = df.loc[t, 'POS'].values
                call = list(set(df.loc[t, 'parent']))[0]
                tracts_format.append([cross, s, pos.min(), pos.max(), call])

        TRACTS.extend(tracts_format)

    TRACTS = pd.DataFrame(TRACTS, columns=['cross','strain','start','end','parent'])

    # circularize tracts
    for s, df in TRACTS.sort_values(by='start').groupby('strain'):
        if df.shape[0] == 1:
            TRACTS.loc[df.index[0], 'start'] = 0
            TRACTS.loc[df.index[0], 'end'] = ref_genome_length
        if df.shape[0] > 2:
            first = df.index[0]
            last = df.index[-1]

            if df.loc[first, 'parent'] == df.loc[last, 'parent']:
                TRACTS.loc[first, 'start'] = df.loc[last, 'start']
                TRACTS.drop(last, inplace=True)
    return TRACTS

# %% [code]
TRACTS = []
for cross in cross_order:
    tracts = make_tracts(cross, MARKERS[cross])
    TRACTS.append(tracts)
TRACTS = pd.concat(TRACTS).reset_index(drop=True)

# %% [markdown]
## Haplotype paintings

# %% [code]
def plot_rearrangement_SpA():
    arcs = [(21e3,47e3), (57.5e3,65.5e3), (60.5e3,26e3), (46.5e3,69e3)]
    y_offset = -0.1
    for a in arcs:
        al = np.abs(a[0]-a[1])
        a = Arc((np.mean(a), 1.7+y_offset), al, 5e-5*al, theta1=0, theta2=180, lw=1, alpha=0.5, color='k', clip_on=False)
        ax.add_artist(a)
        y_offset += 0.05

    segments = [(0,21e3), (47e3,57.5e3), (65.5e3, 60.5e3), (26e3,46.5e3), (69e3, len(artificial_genome.seq))]
    y_offset = -0.1
    for seg in segments:
        if seg[0] < seg[1]:
            color = 'k'
            start = min(seg)
            end = max(seg)
        else:
            color='r'
            start = max(seg)
            end = min(seg)
        

        ax.arrow(start, 1.7+y_offset, end-start, 0, color=color, width=0.1, head_length=1e3, length_includes_head=True, head_width=0.2, zorder=z)
        y_offset += 0.05

# %% [code]
def transform_rad(x, padding_rad=1.28):
    return (x/(ref_genome_length*padding_rad))*2*np.pi

# %% [markdown]
## Fig 2A

# %% [code]
fig = plt.figure(figsize=[10,6])
gs = plt.GridSpec(ncols=3, nrows=2)

inc = 0.05
lw = 0.8
pad_y = 0.015
pad_x = 4/7*pad_y
pad_lines = 5
plot_data = True

for cross_type, rect in zip([['CC1','CC2','CC3'],
                             ['BB1','BB2'],
                             ['BC1','BC2'],
                             ['BA1','BA2'],
                             ['BSc1','BSc2']],
                                    
                            [(0, 1/4, 3/7, 3/4),
                             (3/7, 1/2, 2/7,1/2),
                             (5/7, 1/2, 2/7,1/2),
                             (3/7, 0, 2/7,1/2),
                             (5/7, 0, 2/7,1/2)]):
    
    #apply padding
    rect = np.array(rect)+np.array([pad_x, pad_y, -2*pad_x, -2*pad_y])
    
    ax = fig.add_axes(rect, projection='polar')
    

    ax2 = fig.add_axes(rect)
    ax2.axis('off')
    ax2.set_xlim(0,10)
    ax2.set_ylim(0,10)
    

    cross_ybar = {}
    y = 0
    
    total_s = sum([len(STRAIN_ORDER_MARKERS[cross]) for cross in cross_type]) + pad_lines*(len(cross_type)-1)
    

    for cross in cross_type:
        
        y_start_cross = y
        
        for s in STRAIN_ORDER_MARKERS[cross]:
            if plot_data:
                tracts = TRACTS.loc[TRACTS['strain']==s]

                for i in tracts.index:
                    (start, end, parent) = tracts.loc[i, ['start','end','parent']]
                    start, end = [transform_rad(x) for x in (start, end)]
                    parent = parent.split('.')[-1]
                    c = parents_color[parent]

                    if start < end:
                        X = np.arange(start, end, inc)
                        Y = np.repeat(y, X.shape[0])
                        ax.plot(X, Y, color=c, lw=lw)
                    elif start > end:
                        X = np.arange(start, transform_rad(ref_genome_length), inc)
                        Y = np.repeat(y, X.shape[0])
                        ax.plot(X, Y, color=c, lw=lw)
                        X = np.arange(0, end, inc)
                        Y = np.repeat(y, X.shape[0])
                        ax.plot(X, Y, color=c, lw=lw)
            y += 1
        
        cross_ybar[cross] = [y_start_cross, y]
        
        y += pad_lines

    # plot GFF entries
    
    R = Rectangle((0, y), transform_rad(ref_genome_length), 10, fc='0.9', lw=0, zorder=-1)
    ax.add_patch(R)
    
    for i in artificial_genome_feat.loc[~artificial_genome_feat['annot_type'].isin(['gc_cluster','other'])].sort_values(by='annot_plot_order', ascending=False).index:
        start, end, color, w, z = artificial_genome_feat.loc[i, [3,4,'annot_color','annot_width','annot_plot_order']]
        start, end = [transform_rad(x) for x in (start, end)]
        w *= 10
        R = Rectangle((start, y+(10-w)/2), end-start, w, fc=color, lw=0, zorder=z)
        ax.add_patch(R)
        
    y += 3*pad_lines
    
    for X in np.arange(0,ref_genome_length, 1e4):
        x = transform_rad(X)
        ax.scatter(x, y, marker='o', s=4, color='k')
        if x <= np.pi:
            ha='left'
        else:
            ha='right'
        if -1*np.pi/2 <= x < np.pi/2:
            va = 'bottom'
        else:
            va = 'top'
        ax.text(x, y+pad_lines, f'{X*1e-3:.0f}', size=8, va=va, ha=ha, color='k')
    
    ax.set_theta_offset(.5*np.pi)
    ax.set_theta_direction(-1)
    ax.grid(False)
    
    ax.set_yticks([])
    ax.set_ylim(np.array([0, y]) + np.repeat(total_s,2)*np.array([-0.3, 0.1]))
    
    ax.spines['polar'].set_visible(False)
    ax.set_xticks([])
    
    
    T1 = ax.transData.transform
    T2 = ax2.transData.inverted().transform
    
    for cross in cross_type:
        p1, p2 = [p.split('.')[-1] for p in parents_dict[cross]]
        ybar = np.array(cross_ybar[cross])*0.93
        (x1, y1), (x2, y2) = T2(T1([(0, yi) for yi in ybar]))
        dy = y2-y1
        if cross in ['CC1', 'CC2', 'CC3']:
            dx = -0.5
        else:
            dx = -0.7
        R = Rectangle((x1+0.1*dx, y1), dx, dy, fc=cross_color[cross], lw=0)
        ax2.add_patch(R)
        ax2.text(x1+0.6*dx, y1+0.5*dy, cross, color='w', rotation=90, ha='center', va='center')
        
        ax2.scatter([3.9, 3.9], [y1+0.7*dy, y1+0.3*dy], marker='s', s=42, color=[parents_color[p] for p in [p1, p2]])
        ax2.text(3.6, y1+0.7*dy, fr'{p1} (${parents_group[p1]}$)', size=8, ha='right', va='center')
        ax2.text(3.6, y1+0.3*dy, fr'{p2} (${parents_group[p2]}$)', size=8, ha='right', va='center')

# plot legend
ax = fig.add_axes((0, 0, 3/7, 1/4))
ax.axis('off')
legend_elms = []
for at in annot_type_order:
    if at != 'gc_cluster':
        l = annot_type_alias[at]
        fc = annot_type_color[at]
        lw = 0
        if fc == 'white':
            lw = 1
        legend_elms.append(Line2D([0], [0], color='w', marker='s', ms=9, mfc=fc, mec='black', mew=lw, label=l))
ax.legend(handles=legend_elms, ncol=3, loc=9, bbox_to_anchor=(0.5, 0.8), frameon=False,
         borderpad=0.0, labelspacing=0.2, handlelength=0.3, columnspacing=1.5)


#plt.savefig(f'/home/mathieu/mhenault_landrylab/Publications/mito_ma/resubmission2_GRes/fig/Fig2A.svg', dpi=300)
#plt.show()
plt.close()

# %% [markdown]
## Recombination breakpoints

# %% [code]
bins = pd.interval_range(start=0, end=ref_genome_length+1, freq=250, closed='left')

breakpoints = []

for cross, tracts in TRACTS.loc[TRACTS['strain'].isin(strains_mthap)].groupby('cross'):

    markers = MARKERS[cross]
    count_markers = pd.cut(VCF[cross].loc[VCF[cross]['vma_uid'].isin(markers)].groupby('vma_uid')['POS'].apply(lambda x: x.iloc[0]), bins=bins).value_counts().rename('marker')
    tracts = tracts.copy()
    # filter out haplotypes with no recombination breakpoint
    for s, df in tracts.groupby('strain'):
        if df.shape[0] == 1:
            tracts.drop(df.index, inplace=True)
    count_bkp = pd.cut(tracts[['start','end']].values.flatten(), bins=bins).value_counts().rename('bkp')
    dat = pd.concat([count_markers, count_bkp], axis=1)
    dat['mid'] = [i.mid for i in dat.index]
    dat['cross'] = cross
    dat['bin'] = dat.index
    breakpoints.append(dat)

breakpoints = pd.concat(breakpoints).reset_index(drop=True)
        
# compute recombination rate per window per strain
for cross, df in breakpoints.groupby('cross'):
    n = ma_strains.loc[(ma_strains['filename'].isin(strains_mthap)) & (ma_strains['cross']==cross)].shape[0]
    breakpoints.loc[df.index, 'rate'] = np.where(df['bkp']==0, 0, df['bkp']/(df['marker']*n))

# %% [markdown]
## Fig 2B

# %% [code]
fig = plt.figure(figsize=[10,6])
gs = plt.GridSpec(ncols=1, nrows=2, height_ratios=[3,1], hspace=0.15, left=0.09, right=0.89, top=0.97, bottom=0.08)
ax = fig.add_subplot(gs[0])
wdw = 250
xlim = (-500/wdw, (ref_genome_length+500)/wdw)

dat = [np.nan]*22

for cross, idx in cross_order.items():
    
    bp = breakpoints.loc[breakpoints['cross']==cross].sort_values(by='mid').set_index('mid')
    m = (bp['marker']/wdw).rename(f'{cross}.mark')
    if cross != 'BA1':
        r = bp['rate'].rename(f'{cross}.rec')
    else:
        r = pd.Series(np.repeat(0, m.shape[0]), index=m.index, name=f'{cross}.rec')
    
    dat[2*idx] = m
    dat[2*idx+1] = r

dat = pd.concat(dat, axis=1).T
vmax_mark = np.quantile(dat.iloc[np.arange(0,22,2)].values.flatten(), 0.98)
vmax_rec = np.quantile(dat.iloc[np.arange(0,22,2)+1].values.flatten(), 0.98)

mask = np.ndarray(dat.shape)
mask[np.arange(0,22,2), :] = False
mask[np.arange(0,22,2)+1, :] = True
hm = np.ma.MaskedArray(dat, mask)

HM1 = ax.imshow(hm, aspect='auto', interpolation='nearest', cmap='Reds',
                vmin=-0.2*vmax_mark, vmax=vmax_mark)

mask = np.ndarray(dat.shape)
mask[np.arange(0,22,2), :] = True
mask[np.arange(0,22,2)+1, :] = False
hm = np.ma.MaskedArray(dat, mask)
HM2 = ax.imshow(hm, aspect='auto', interpolation='nearest', cmap='Greys',
                vmin=-0.2*vmax_rec, vmax=vmax_rec)

ax.set_xlim(xlim)
ax.set_xticks(np.arange(0, dat.shape[1], 1e4/wdw))
ax.set_xticklabels(range(0,90,10))

ax.set_ylim(21.6, -0.6)
ax.set_yticks(range(22))
ax.set_yticklabels(np.tile(['M','R'], 11), size=9)


for i in ['left','right','top']:
    ax.spines[i].set_visible(False)
for y in np.arange(-0.5, 23, 2):
    ax.axhline(y, color='white', lw=5, zorder=1)
for cross, y in cross_order.items():
    ax.text(-4e3/wdw, 2*y+0.5, cross, va='center', ha='right', size=14)

#plot color bars

ax_cbar = fig.add_axes([0.89, 0.7, 0.01, 0.2])
cb = plt.colorbar(HM1, cax=ax_cbar, orientation='vertical', ticks=np.arange(0, vmax_mark, float(f'{vmax_mark/6:.0e}')), 
                  label=' M (marker\ndensity, bp$^{-1}$)')
cb.outline.set_visible(False)
ax_cbar.set_ylim(0, vmax_mark)

ax_cbar = fig.add_axes([0.89, 0.4, 0.01, 0.2])
cb = plt.colorbar(HM2, cax=ax_cbar, orientation='vertical', ticks=np.arange(0, vmax_rec, float(f'{vmax_rec/6:.0e}')),
                  label='R (rec rate,\nline$^{-1}$ marker$^{-1}$)')
cb.outline.set_visible(False)
ax_cbar.set_ylim(0, vmax_rec)


# plot GFF entries

ax = fig.add_subplot(gs[1])
xlim = (-500, ref_genome_length+500)
ax.arrow(0, 1, ref_genome_length, 0, color='0.9', width=1, head_length=0, head_width=0, zorder=-1)

for i in artificial_genome_feat.loc[~artificial_genome_feat['annot_type'].isin(['gc_cluster','other'])].sort_values(by='annot_plot_order', ascending=False).index:
    start, end, color, w, z = artificial_genome_feat.loc[i, [3,4,'annot_color','annot_width','annot_plot_order']]
    ax.arrow(start, 1, end-start, 0, color=color, width=w, head_length=0, head_width=0, zorder=z)

#create linked labels
pap_sub = [a for a in pres_abs_poly if a in artificial_genome_feat['Name'].values]
for x, a in zip(np.linspace(-6e3, ref_genome_length+6e3, len(pap_sub)),
                pap_sub):

    fc = at_color_hm[at_dict[a]]
    ec = 'white'
    tc = 'white'
    if fc == 'white':
        tc = 'black'
        ec = 'black'
    if fc == 'limegreen':
        tc = 'black'

    t = ax.text(x, 6, pres_abs_poly_alias[a], rotation=90, ha='center', va='top', color=tc, size=8,
                 bbox=dict(ec=ec, fc=fc, lw=0.5, boxstyle='square,pad=0.18'), zorder=1)
    pos = artificial_genome_feat.loc[artificial_genome_feat['Name']==a].iloc[0,3]
    ax.plot([pos, x, x], [1.5, 3, 6], lw=0.5, color='black', zorder=-1, clip_on=False)

ax.set_xticks(np.arange(0,83e3,10e3))
ax.set_xticklabels(np.arange(0,83,10))
ax.set_xlabel('kb')
ax.set_ylim(0.5, 6)
ax.set_yticks([])
ax.set_xlim(xlim)
for i in ['left','right','top']:
    ax.spines[i].set_visible(False)

plt.savefig(f'/home/mathieu/mhenault_landrylab/Publications/mito_ma/resubmission2_GRes/fig/Fig2B.svg', dpi=300)
#plt.show()
plt.close()

# %% [markdown]
## Fig S4-14

# %% [code]
xlim = (-500,ref_genome_length+500)

for cross, tracts in TRACTS.groupby('cross'):

    fig = plt.figure(figsize=[8,9])
    gs = plt.GridSpec(nrows=4, ncols=1, height_ratios=[10,1.5,1,1], hspace=0.3,
                     top=0.92, bottom=0.07, right=0.96, left=0.12)

    # plot haplotype paintings
    ax = fig.add_subplot(gs[0])

    strain_order = STRAIN_ORDER_MARKERS[cross]
    strain_order_plot = dict([(j,i) for (i,j) in enumerate(strain_order)])
    markers = MARKERS[cross]
    vcf_melt = VCF_MELT[cross]
    vcf_melt = vcf_melt.loc[(vcf_melt['vma_uid'].isin(markers)) &
                            (vcf_melt['higher']==True) &
                            (vcf_melt['parent'].isin(parents_dict[cross]))]

    alpha = 0.3
    for s, df in tracts.groupby('strain'):
        if s in strain_order:
            y = strain_order_plot[s]
            for i in df.index:
                (start, end, parent) = df.loc[i, ['start','end','parent']]
                c = parents_color[parent.split('.')[-1]]
                if start < end:
                    fa = FancyArrow(start, y, end-start, 0, width=0.8, head_length=0, head_width=0, color=c, lw=0, alpha=alpha)
                    ax.add_patch(fa)
                elif start > end:
                    fa1 = FancyArrow(0, y, end, 0, width=0.8, head_length=0, head_width=0, color=c, lw=0, alpha=alpha)
                    fa2 = FancyArrow(start, y, ref_genome_length-start, 0, width=0.8, head_length=0, head_width=0, color=c, lw=0, alpha=alpha)
                    ax.add_patch(fa1)
                    ax.add_patch(fa2)
            ax.text(-500, y, ma_strains.loc[s, 'mt_haplotype'], ha='right', va='center', size=6)

    for (s, parent), df in vcf_melt.groupby(['filename','parent']):
        if s in strain_order:
            y = strain_order_plot[s]
            c = parents_color[parent.split('.')[-1]]
            ax.scatter(df['POS'], np.repeat(y, df.shape[0]), s=6, color=c)

    ax.set_ylim(-3, len(strain_order_plot))
    ax.set_xlim(xlim)
    ax.set_xticks(np.arange(0, 81e3, 1e4))
    ax.set_xticklabels(range(0,90,10))
    for i in ['left','right','top']:
        ax.spines[i].set_visible(False)
    ax.set_yticks([])
    ax.legend(handles=[Line2D([0], [0], color='white', marker='s', ms=12, mfc=parents_color[parent.split('.')[-1]],
                              label=fr'{parent.split(".")[-1]} (${parents_group[parent.split(".")[-1]]}$)', ) for parent in parents_dict[cross]],
              ncol=2, frameon=False, bbox_to_anchor=[0.5,0.99], loc=8)

    #set title of the main axes
    fig.text(0.5, 0.99, f'{cross} ({len(markers)} markers, {len(strain_order)} haplotypes)', size=14, ha='center', va='top')

    
    # plot GFF entries
    ax = fig.add_subplot(gs[1])
    for i in ['left','right','top']:
        ax.spines[i].set_visible(False)
    ax.arrow(0, 1, ref_genome_length, 0, color='0.9', lw=0, width=1, head_length=0, head_width=0, zorder=-1)

    legend_elms = []
    for at in annot_type_order:
        if at != 'gc_cluster':
            l = annot_type_alias[at]
            fc = annot_type_color[at]
            lw = 0
            if fc == 'white':
                lw = 1
            legend_elms.append(Line2D([0], [0], color='w', marker='s', ms=9, mfc=fc, mec='black', mew=lw, label=l))
    ax.legend(handles=legend_elms, ncol=6, loc=9, bbox_to_anchor=(0.5, 1.1), frameon=False,
             borderpad=0.0, labelspacing=0.2, handlelength=0.3, columnspacing=1.5)

    for i in artificial_genome_feat.loc[~artificial_genome_feat['annot_type'].isin(['gc_cluster','other'])]\
.sort_values(by='annot_plot_order', ascending=False).index:
        start, end, color, w, z = artificial_genome_feat.loc[i, [3,4,'annot_color','annot_width','annot_plot_order']]
        ax.arrow(start, 1, end-start, 0, color=color, lw=0, width=w, head_length=0, head_width=0, zorder=z)
    ax.set_xlim(xlim)
    ax.set_ylim(0.45, 4)
    ax.set_xticks(np.arange(0, 81e3, 1e4))
    ax.set_xticklabels(range(0,90,10))
    ax.set_yticks([])

    if cross in ['BA1', 'BA2']:
        plot_rearrangement_SpA()

    # plot recombination rate
    bp = breakpoints.loc[breakpoints['cross']==cross].sort_values(by='mid')

    ax = fig.add_subplot(gs[2])
    ax.plot(bp['mid'], bp['rate'], color='k', zorder=1, clip_on=False)
    ax.set_ylabel('rec rate\n(line$^{-1}$ marker$^{-1}$)', size=8)
    ax.set_xlim(xlim)
    ax.set_xticks(np.arange(0, 81e3, 1e4))
    ax.set_xticklabels(range(0,90,10))
    sns.despine(ax=ax, trim=True)

    #plot marker density
    ax = fig.add_subplot(gs[3])
    ax.plot(bp['mid'], bp['marker']*0.01, color='red', zorder=0)
    ax.set_ylabel('marker density\n(bp$^{-1}$)', size=8)
    ax.set_xlim(xlim)
    ax.set_xticks(np.arange(0, 81e3, 1e4))
    ax.set_xticklabels(range(0,90,10))
    ax.set_xlabel('kb')
    sns.despine(ax=ax, trim=True)

    #plt.savefig(f'/home/mathieu/mhenault_landrylab/Publications/mito_ma/resubmission2_GRes/fig/detailed_haplotypes_paintings_{cross}.svg')
    #plt.show()
    plt.close()

# %% [markdown]
## Recombination rate, tract length and feature enrichment

# %% [code]
#import growth rates in liquid media from Fijarczyk et al. 2021
growth = pd.read_csv('/home/mathieu/mhenault_landrylab/Experiments/generation_time/analysis/max_yield.csv', index_col=0)
growth = growth.loc[(growth['sp_high_t0']) & (growth['sp_growth'])].groupby('strain')['max_yield'].mean()

# %% [code]
#compute the sum of parental mtDNA sizes
parents_genome_size = {cross: sum([len(CORR[p.split('.')[-1]].seq) for p in parents_dict[cross]]) for cross in cross_order}

# %% [code]
parents_dist_matrix = pd.read_csv('/mnt/HDD3/mito_ma/divergence/nuc_pairwise_divergence.csv', index_col=0)
taxa = pd.read_csv('/mnt/HDD3/mito_ma/divergence/taxa.csv', index_col=3)
parents_dist = {}
for cross in cross_order:
    
    nuc_dist = []
    Cross = cross[:-1]
    g1, g2 = Cross[0], Cross[1:]
    P1 = taxa.loc[g1, 'strain_id']
    P2 = taxa.loc[g2, 'strain_id']
    
    for i,j in itertools.product(P1, P2):
        if i != j and [i, j] == sorted([i,j]):
            nuc_dist.append(parents_dist_matrix.loc[i,j])
    
    parents_dist[cross] = np.mean(nuc_dist)

# %% [code]
#build recombination summary tables:
# count of rec tracts per line
REC = []
# rec tract lengths
RECL = []
# features of rec breakpoints
RECF = []
for (cross, s), df in TRACTS.groupby(['cross','strain']):
    if s in strains_mthap:
        if df.shape[0] > 1:
            bp = df[['start','end']].values.flatten()
            REC.append([cross, s, bp.shape[0]/2]) #divide by two, because each event has 2 junctions

            for b in bp:
                F = gft[b]
                if F:
                    F = list(F)
                    at, name = artificial_genome_feat.loc[[f[2] for f in F]].sort_values(by='annot_plot_order', ascending=False).iloc[0][['annot_type','Name']].values
                    RECF.append([cross, s, b, at, name])
                else:
                    RECF.append([cross, s, b, 'other', 'none'])
            for i in df.index:
                start, end = df.loc[i, ['start','end']].values
                if start < end:
                    RECL.append([cross, s, i, end-start+1])
                if start > end:
                    RECL.append([cross, s, i, end+ref_genome_length-start+1])
        else:
            REC.append([cross, s, 0])

REC = pd.DataFrame(REC, columns=['cross','strain','count'])
RECF = pd.DataFrame(RECF, columns=['cross','strain','breakpoint','annot_type','name'])
RECL = pd.DataFrame(RECL, columns=['cross','strain','tract','len'])
#transform tract lengths in kb
RECL['len'] *= 1e-3
# add bins to RECL
RECL['bin'] = pd.cut(RECL['len'], bins=pd.interval_range(start=0, end=83000, freq=100, closed='left'))

# %% [code]
# add variables to the REC table
# growth-adjusted recombination rate
for s, df in REC.groupby('strain'):
    for f in ['passage', 'identity_filter', 'diploid_filter', 'timepoint_filter','mt_haplotype']:
        REC.loc[df.index, f] = ma_strains.loc[s, f]

    s = s.split('.')[-1].split('_')[0]
    if s in growth.index:
        REC.loc[df.index, 'growth'] = growth.loc[s]

# compute rates and growth-adjusted rates
median_growth = np.median(REC.loc[~np.isnan(REC['growth']), 'growth'])

for (s, cross), df in REC.groupby(['strain','cross']):
    df = df.iloc[0]

    rate = df['count']/parents_genome_size[cross]
    REC.loc[df.name, 'rate'] = rate*1e5

# %% [code]
# relative abundance of feature types in rec breakpoints
RECF_ABD = RECF.value_counts(['cross','name','annot_type']).rename('abd').reset_index()
for cross, df in RECF_ABD.groupby('cross'):
    RECF_ABD.loc[df.index, 'abd_rel'] = df['abd']/df['abd'].sum()
RECF_ABD['at_order'] = RECF_ABD['annot_type'].replace(annot_type_order)

# %% [code]
RECF_ABD_AGG = RECF_ABD.groupby(['cross', 'annot_type', 'at_order'])['abd'].sum().reset_index()
for cross, df in RECF_ABD_AGG.groupby('cross'):
    RECF_ABD_AGG.loc[df.index, 'abd_rel'] = df['abd']/df['abd'].sum()

# %% [code]
# feature classification of markers
MARKF = []
for cross, markers in MARKERS.items():
    vcf = VCF[cross].groupby('vma_uid').apply(lambda x: x.iloc[0]['POS'])
    for m in markers:
        pos = vcf.loc[m]
        F = gft[pos]
        if F:
            F = list(F)
            at, name = artificial_genome_feat.loc[[f[2] for f in F]].sort_values(by='annot_plot_order', ascending=False).iloc[0][['annot_type','Name']].values
            if at == 'gene':
                print(pos, at)
                break
            MARKF.append([cross, m, at, name])
        else:
            MARKF.append([cross, m, 'other', 'none'])
MARKF = pd.DataFrame(MARKF, columns=['cross','vma_uid','annot_type','name'])

# %% [code]
# relative abundance of feature types in markers
MARKF_ABD = MARKF.value_counts(['cross','name','annot_type']).rename('abd').reset_index()
for cross, df in MARKF_ABD.groupby('cross'):
    MARKF_ABD.loc[df.index, 'abd_rel'] = df['abd']/df['abd'].sum()
MARKF_ABD['at_order'] = MARKF_ABD['annot_type'].replace(annot_type_order)

# %% [code]
MARKF_ABD_AGG = MARKF_ABD.groupby(['cross', 'annot_type', 'at_order'])['abd'].sum().reset_index()
for cross, df in MARKF_ABD_AGG.groupby('cross'):
    MARKF_ABD_AGG.loc[df.index, 'abd_rel'] = df['abd']/df['abd'].sum()

# %% [code]
#average variables for crosses
AVERAGE_REC = []

for cross, df in REC.groupby('cross'):
    
    mean_rate = df['rate'].mean()
    #mean_adj_rate = df['rate_adj'].mean()
    mean_len = RECL.loc[RECL['cross']==cross, 'len'].mean()
    n_markers = len(MARKERS[cross])
    p1, p2 = [p.split('.')[-1] for p in parents_dict[cross]]
    dist = parents_dist[cross]
    
    AVERAGE_REC.append([cross, mean_rate, mean_len, n_markers, dist])
AVERAGE_REC = pd.DataFrame(AVERAGE_REC, columns=['cross','rate','len','n_markers','dist'])
AVERAGE_REC.set_index('cross', drop=False, inplace=True)

# %% [code]
def regression_plot(data, x, y, c, px, py, xplot=None):
    reg = stats.linregress(data[x], data[y])
    if xplot == None:
        X = np.array([data[x].min(), data[x].max()])
    else:
        X = np.array(xplot)
    ax.plot(X, X*reg.slope+reg.intercept, color=c, zorder=0)
    ax.text(px, py, f'r={reg.rvalue:.3f}\np={plot_pval_text(reg.pvalue)}', size=10, transform=ax.transAxes, color=c)

# %% [markdown]
## Fig 3ABC

# %% [code]
for (pis, pis_alias) in zip([True,False], ['', '.no_is']):

    plot_intraspecific = pis

    fig = plt.figure(figsize=[10, 4.5])

    gs = plt.GridSpec(ncols=5, nrows=2, width_ratios=[1.5,1,4,2,2], wspace=0.7, hspace=0.7,
                      left=0.05, right=0.97, bottom=0.15, top=0.85)

    ### counts of breakpoints
    
    ax1 = fig.add_subplot(gs[:, 0])
    if pis == True:
        ax2 = fig.add_subplot(gs[:, 1])
    else:
        ax2 = fig.add_subplot(gs[:, :2])

    fig.text(0.01, 0.93, 'A', size=24, weight='bold')

    bkp_count_color = dict(zip(range(0,9,2), 
                              [f'{1-(i/10):.2f}' for i in np.logspace(0,1,5)]))

    y_labels = pd.Series(dtype=str)
    for cross, df in REC.groupby('cross'):
        y = (cross_order[cross]-10)*-1
        y_labels.loc[y] = cross
        df1 = df.value_counts('count').reset_index()
        df1.columns = ['bkp','count']
        df1['freq'] = df1['count']/df1['count'].sum()
        df1 = df1.sort_values(by='bkp')
        df1['cumsum'] = np.cumsum(df1['freq'])

        for i in df1.index:
            ct, cs = df1.loc[i, ['bkp', 'cumsum']]
            ax1.barh(y, cs, color=bkp_count_color[ct], lw=0, zorder=(ct/2-4)*-1)

        perc_rec = (1-df1.iloc[0,2])*100
        ax2.barh(y, perc_rec, color=cross_color[cross], lw=0)

    
    ax1.set_yticks(range(11))
    ax1.set_yticklabels(y_labels.loc[range(11)])
    ax1.set_xticks([0,0.5,1])
    ax1.set_xlabel('Proportion of\nmtDNA haplotypes')

    ax2.set_yticks(range(11))
    ax2.set_yticklabels(y_labels.loc[range(11)])
    ax2.set_xticks([0, 25, 50])
    ax2.set_xlabel('% recombinant\nhaplotypes')

    legend_elms = [Line2D([0], [0], color='w', marker='s', ms=10, label=l, mfc=c) for l,c in bkp_count_color.items()]
    ax1.legend(handles=legend_elms, loc=3, ncol=3, bbox_to_anchor=[-0.15,1], title='Count of rec tracts', frameon=False,
              columnspacing=1, borderaxespad=0.2)
    
    if pis == False:
        ax1.remove()
        ax2.set_xticks(np.arange(0,61,10))

    ### recombination rate

    fig.text(0.29, 0.93, 'B', size=24, weight='bold')

    ax = fig.add_subplot(gs[0,2])


    sns.violinplot(x='cross', y='rate', data=REC, order=cross_order, color='w', ax=ax, zorder=0,
                  cut=0, scale='width', inner=None, lw=1)
    sns.stripplot(x='cross', y='rate', data=REC, order=cross_order, ax=ax,
                  palette=cross_color, size=4, jitter=0.15, alpha=0.3, zorder=3)

    ax.set_xticklabels(cross_order, rotation=45, ha='right')
    ax.set_xlabel('')
    ax.set_ylabel('Rec rate\n'+r'($\times10^{-5}$ bp$^{-1}$)')

    ax.scatter(np.arange(11), AVERAGE_REC.loc[cross_order, 'rate'], color=(1,1,1,0), edgecolor='k', zorder=4)

    ax = fig.add_subplot(gs[0,3])
    for cross in cross_order:
        mean = AVERAGE_REC.loc[cross, 'rate']
        n_markers = AVERAGE_REC.loc[cross, 'n_markers']
        ax.scatter(n_markers, mean, color=cross_color[cross])
    ax.set_xlabel('Number of markers')
    ax.set_ylabel('Average rec rate\n'+r'($\times10^{-5}$ bp$^{-1}$)')

    # regression
    regression_plot(AVERAGE_REC, 'n_markers', 'rate', 'k', 0.1, 0.2)

    ax = fig.add_subplot(gs[0,4])
    for cross in cross_order:
        mean = AVERAGE_REC.loc[cross, 'rate']
        dist = AVERAGE_REC.loc[cross, 'dist']
        ax.scatter(dist, mean, color=cross_color[cross])
    ax.set_xlabel('Parental divergence\n(subs site$^{-1}$)')
    ax.set_ylabel('Average rec rate\n'+r'($\times10^{-5}$ bp$^{-1}$)')

    # regression
    regression_plot(AVERAGE_REC, 'dist', 'rate', 'k', 0.5, 0.8, xplot=None)
    if plot_intraspecific:
        regression_plot(AVERAGE_REC.loc[AVERAGE_REC['cross'].apply(lambda x: x not in ['BSc1','BSc2'])],
                        'dist', 'rate', 'red', 0.45, 0.1, xplot=(0,0.04))

    ### recombination tract length

    fig.text(0.29, 0.46, 'C', size=24, weight='bold')

    ax = fig.add_subplot(gs[1,2])

    sns.violinplot(x='cross', y='len', data=RECL, order=cross_order, color='w', ax=ax, zorder=0,
                  cut=0, scale='width', inner=None, lw=1)
    sns.stripplot(x='cross', y='len', data=RECL, order=cross_order, ax=ax,
                  palette=cross_color, size=4, jitter=0.15, alpha=0.3, zorder=3)

    ax.set_xticklabels(cross_order, rotation=45, ha='right')
    ax.set_xlabel('')
    ax.set_ylabel('Rec tract length\n(kb)')

    ax.scatter(np.arange(11), AVERAGE_REC.loc[cross_order, 'len'], color=(1,1,1,0), edgecolor='k', zorder=4)

    ax = fig.add_subplot(gs[1,3])
    for cross in cross_order:
        mean = AVERAGE_REC.loc[cross, 'len']
        n_markers = AVERAGE_REC.loc[cross, 'n_markers']
        ax.scatter(n_markers, mean, color=cross_color[cross])
    ax.set_xlabel('Number of markers')
    ax.set_ylabel('Average rec tract\nlength (kb)')

    # regression
    regression_plot(AVERAGE_REC.loc[AVERAGE_REC['cross']!='BA1'], 'n_markers', 'len', 'k', 0.2, 0.6)

    ax = fig.add_subplot(gs[1,4])
    for cross in cross_order:
        mean = AVERAGE_REC.loc[cross, 'len']
        dist = AVERAGE_REC.loc[cross, 'dist']
        ax.scatter(dist, mean, color=cross_color[cross])
    ax.set_xlabel('Parental divergence\n(subs site$^{-1}$)')
    ax.set_ylabel('Average rec tract\nlength (kb)')

    # regression
    regression_plot(AVERAGE_REC.loc[AVERAGE_REC['cross']!='BA1'], 'dist', 'len', 'k', 0.6, 0.4)
    if plot_intraspecific:
        regression_plot(AVERAGE_REC.loc[~AVERAGE_REC['cross'].isin(['BA1','BSc1','BSc2'])],
                        'dist', 'len', 'red', 0.1, 0.85, xplot=(0,0.04))


    ax = fig.add_subplot(gs[0, 2:])
    ax.axis('off')
    legend_elms = [Line2D([0], [0], color='white', marker='o', ms=8, mfc=cross_color[cross], label=cross) for cross in cross_order]
    ax.legend(handles=legend_elms, ncol=6, loc=3, bbox_to_anchor=[0.08, 1.05])

    sns.despine()
    #if pis == True:
        #plt.savefig(f'/home/mathieu/mhenault_landrylab/Publications/mito_ma/resubmission2_GRes/fig/Fig3ABC{pis_alias}.svg', dpi=300)
        #plt.show()
    plt.close()

# %% [code]
#compute frequency deviations (i.e. overrepresentation score) 
# for aggregated and expanded  datasets


freq_dev_agg = {}
chi2_fdagg_pval = []
for cross in cross_order:
    
    freq_obs = RECF_ABD_AGG.loc[RECF_ABD_AGG['cross']==cross].set_index('annot_type')['abd'].rename('obs')
    freq_exp = MARKF_ABD_AGG.loc[MARKF_ABD_AGG['cross']==cross].set_index('annot_type')['abd'].rename('exp')
    
    freq = pd.concat([freq_obs, freq_exp], axis=1).fillna(0)
    chi2 = stats.chisquare(freq['obs'], freq['exp']/freq['exp'].sum()*freq['obs'].sum())
    chi2_fdagg_pval.append([cross, chi2[1]])
    freq_dev_agg[cross] = (freq['obs']/freq['obs'].sum())-(freq['exp']/freq['exp'].sum())
    
freq_dev_agg = pd.DataFrame(freq_dev_agg)
chi2_fdagg_pval = pd.DataFrame(chi2_fdagg_pval, columns=['cross', 'pval']).set_index('cross')
chi2_fdagg_pval.drop('BA1', inplace=True)
chi2_fdagg_pval['pval_corr'] = multipletests(chi2_fdagg_pval['pval'], method='fdr_bh')[1]

# %% [code]
freq_dev = pd.concat([RECF_ABD.set_index(['cross','annot_type','name'])['abd_rel'].rename('rec_abd_rel'),
                      MARKF_ABD.set_index(['cross','annot_type','name'])['abd_rel'].rename('mark_abd_rel'),
                      at_ident.set_index(['cross','annot_type','name'])['identity']], axis=1).fillna(0)
freq_dev['freq_dev'] = freq_dev['rec_abd_rel'] - freq_dev['mark_abd_rel']
freq_dev = freq_dev.loc[[c for c in cross_order if c!='BA1']]

# %% [markdown]
## Fig 3DEFG

# %% [code]
fig = plt.figure(figsize=[10,5])
gs = plt.GridSpec(ncols=3, nrows=2, width_ratios=[2,1,3], wspace=0.45, hspace=0.35,
                  left=0.05, right=0.98, bottom=0.1, top=0.85)

fig.text(0.01, 0.92, 'D', size=24, weight='bold')

for ax_idx, dat, alias in zip([0,1], [MARKF_ABD_AGG, RECF_ABD_AGG], ['Markers','Rec breakpoints']):
    
    ax = fig.add_subplot(gs[ax_idx, 0])
    y_label = pd.Series(dtype=str)
    for cross, df in dat.sort_values(by='at_order', ascending=True).groupby('cross'):
        df = df.copy()
        df['cumsum'] = np.cumsum(df['abd_rel'])
        y = (cross_order[cross]-10)*-1
        y_label.loc[y] = cross
        for i in df.sort_values(by='cumsum', ascending=True).index:
            at = df.loc[i, 'annot_type']
            fc = annot_type_color[at]
            if at == 'intron':
                ec = 'k'
            else:
                ec = fc
            z = (annot_type_order[at]-5)*-1
            if cross == 'CC1' and alias == 'MARK':
                l = annot_type_alias[at]
            else:
                l = None
            ax.barh(y, df.loc[i, 'cumsum'], color=fc, lw=1, edgecolor=ec, zorder=z)
    if 3 not in y_label.index:
        y_label[3] = 'BA1'
    ax.set_title(alias)
    ax.set_yticks(range(11))
    ax.set_yticklabels(y_label.loc[range(11)])
    
    if ax_idx == 1:
        ax.set_xlabel('Proportion')
        
# legend squares
ax = fig.add_subplot(gs[0,:])
ax.axis('off')
legend_elms = []
for at in annot_type_order:
    l = annot_type_alias[at]
    fc = annot_type_color[at]
    lw = 0
    if fc == 'white':
        lw = 1
    legend_elms.append(Line2D([0], [0], color='w', marker='s', ms=9, mfc=fc, mec='black', mew=lw, label=l))
ax.legend(handles=legend_elms, ncol=4, loc=3, bbox_to_anchor=(0, 1.15), frameon=False,
         borderpad=0.0, labelspacing=0.2, handlelength=0.3, columnspacing=1.5)

    
# heatmap of frequency deviations between markers and recombination breakpoints
ax = fig.add_axes([0.36, 0.2, 0.15, 0.55])
cbar_ax = fig.add_axes([0.36, 0.85, 0.15, 0.03])
fig.text(0.31, 0.83, 'E', size=24, weight='bold')

sns.heatmap(freq_dev_agg.T.loc[cross_order, annot_type_order], cmap='BrBG', center=0, ax=ax, 
            cbar_ax=cbar_ax, cbar_kws=dict(orientation='horizontal', ticks=np.arange(-0.2,0.5,0.2), label='Freq difference'))


for cross in chi2_fdagg_pval.index:
    ax.text(7.2, cross_order[cross]+0.7, plot_pval_symbol(chi2_fdagg_pval.loc[cross, 'pval_corr']),
            size=12, ha='left', va='center')
ax.set_xlabel('')
ax.set_xticks(np.arange(7)+0.5)
ax.set_xticklabels([annot_type_alias[at] for at in annot_type_order], rotation=45, ha='right')

## F panel - plot identity levels
ax = fig.add_subplot(gs[0, 2])
fig.text(0.55, 0.92, 'F', size=24, weight='bold')
legend_elms = [np.nan]*7

for (at, cross), df in at_ident.loc[at_ident['annot_type'].isin(annot_type_order)].groupby(['annot_type','cross']):
    x = cross_order[cross]-0.35+annot_type_order[at]*0.1
    fc = annot_type_color[at]
    lw = 0
    if fc == 'white':
        lw = 1
    
    ax.scatter(x, df['identity'].mean(), marker='o', color=fc, edgecolors='k', linewidths=lw)
    if cross == 'CC1':
        legend_elms[annot_type_order[at]] = Line2D([0], [0], color='w', marker='o', mfc=fc, mec='k', mew=lw, label=annot_type_alias[at])
    
ax.legend(handles=legend_elms, ncol=4, loc=3, bbox_to_anchor=(-0.08, 1.15), frameon=False,
         borderpad=0.0, labelspacing=0.2, handlelength=0.3, columnspacing=1.5)

ax.set_xlim(-0.6, 10.6)
ax.set_xticks(range(11))
ax.set_xticklabels(cross_order, rotation=45, ha='right')

ax.set_ylim(0.19, 1.03)
ax.set_yticks(np.arange(0.2, 1.1, 0.2))
ax.set_yticklabels(np.arange(20, 101, 20))
ax.set_ylabel('% nt identity')

## G panel - identity vs freq deviation
ax = fig.add_subplot(gs[1, 2])
fig.text(0.55, 0.42, 'G', size=24, weight='bold')

for (at, cross), df in at_ident.groupby(['annot_type', 'cross']):
    if at in annot_type_order:
        fd = freq_dev_agg.loc[at, cross]
        fc = annot_type_color[at]
        lw = 0
        if fc == 'white':
            lw = 1
        ax.scatter(df['identity'].mean(), fd, c=fc, edgecolor='black', lw=lw, zorder=(annot_type_order[at]*-1)+6)
        
ax.set_xlim(0.29, 1.03)
ax.set_xticks(np.arange(0.3, 1.1, 0.1))
ax.set_xticklabels(np.arange(30,101,10))

ax.set_xlabel('% nt identity')
ax.set_ylabel('Freq difference')

sns.despine()
for i in ['bottom', 'left']:
    cbar_ax.spines[i].set_visible(False)

#plt.savefig(f'/home/mathieu/mhenault_landrylab/Publications/mito_ma/resubmission2_GRes/fig/Fig3DEFG.svg', dpi=300)
#plt.show()
plt.close()

# %% [markdown]
## Fig S17

# %% [code]
fig = plt.figure(figsize=[8,8])
gs = plt.GridSpec(ncols=4, nrows=4, hspace=0.5, wspace=0.6, left=0.1, top=0.95, right=0.98, bottom=0.08)
cross_ax = dict(zip(['CC1','CC2','CC3','BB1','BB2','BC1','BC2','BA2','BSc1','BSc2'], itertools.product(range(3), range(4))))

fig.text(0.01, 0.96, 'A', weight='bold', size=24)

legend_elms = [np.nan]*7
for cross, df in freq_dev.groupby('cross'):
    ax = fig.add_subplot(gs[cross_ax[cross]])
    for at, df1 in df.groupby('annot_type'):
        if at in annot_type_order:
            
            fc = annot_type_color[at]
            lw = 0
            if fc == 'white':
                lw = 1
            ax.scatter(df1['identity'], df1['freq_dev'], c=fc, edgecolor='black', lw=lw, alpha=0.5, zorder=(annot_type_order[at]*-1)+6)
            
            if cross == 'CC1':
                legend_elms[annot_type_order[at]] = Line2D([0], [0], color='w', marker='o', mfc=fc, mec='k', mew=lw, label=annot_type_alias[at])
        
    ax.text(0.5, 1, cross, ha='center', va='top', transform=ax.transAxes, size=14)
    ax.set_xlim(-0.05, 1.05)
    ax.set_xticks(np.arange(0, 1.1, 0.25))
    ax.set_xticklabels(np.arange(0,101,25))
    ax.set_xlabel('% nt identity')
    ax.set_ylim(-0.3, 0.3)
    ax.set_ylabel('freq difference', labelpad=1)
    
ax = fig.add_subplot(gs[2, 2:])
ax.axis('off')
ax.legend(handles=legend_elms, loc=10, ncol=3, handlelength=0.3)
   
fig.text(0.01, 0.25, 'B', weight='bold', size=24)
ax = fig.add_subplot(gs[3, :2])
dat = freq_dev.reset_index()
dat = dat.loc[dat['annot_type']=='exon']
name_order = np.sort(dat['name'].unique())
sns.stripplot(x='name', y='freq_dev', hue='cross', order=name_order, palette=cross_color, 
              hue_order=['CC1','CC2','CC3','BB1','BB2','BC1','BC2','BA2','BSc1','BSc2'], data=dat, ax=ax)
ax.legend_.remove()
ax.set_xticklabels(name_order, rotation=45, ha='right')
ax.set_xlabel('')
ax.set_ylim(-0.1, 0.5)
ax.set_ylabel('freq difference')


ax = fig.add_subplot(gs[3, 2:])
ax.axis('off')
legend_elms = [Line2D([0], [0], color='w', marker='o', mfc=cross_color[cross], label=cross)
               for cross in ['CC1','CC2','CC3','BB1','BB2','BC1','BC2','BA2','BSc1','BSc2']]
ax.legend(handles=legend_elms, loc=10, ncol=3, handlelength=0.3)
    
sns.despine()
#plt.savefig('/home/mathieu/mhenault_landrylab/Publications/mito_ma/resubmission2_GRes/fig/freq_diff_ident.svg')
#plt.show()
plt.close()

# %% [markdown]
## Inheritance biases

# %% [code]
INHERIT = []
for (cross,s), df in TRACTS.loc[TRACTS['strain'].isin(strains_mthap)].groupby(['cross','strain']):
    p1, p2 = parents_dict[cross]
    
    tracts_data = pd.Series(np.repeat(np.nan, ref_genome_length), index=np.arange(ref_genome_length)+1)
    for i in df.index:
        start, end, p = df.loc[i, ['start','end', 'parent']]
        if start < end:
            tracts_data.loc[start:end] = p
        if start > end:
            tracts_data.loc[start:] = p
            tracts_data.loc[:end] = p
    
    #score inheritance of specific features
    for (at, f, start, end), df1 in artificial_genome_feat.groupby(['annot_type', 'Name',3,4]):
        inherit = tracts_data.loc[start:end].value_counts()
        length = end-start+1
        if p1 in inherit.index:
            cp1 = inherit.loc[p1]
        else:
            cp1 = 0
        
        if p2 in inherit.index:
            cp2 = inherit.loc[p2]
        else:
            cp2 = 0
        
        cp_sum = cp1+cp2
        if cp_sum > 0 and cp_sum/length >= 0.8:
            ratio = cp1/cp_sum
        else:
            ratio = np.nan
        
        INHERIT.append([cross, s, at, f, p1, p2, cp1, cp2, length, ratio])
        
    # score global inheritance
    inherit = tracts_data.value_counts()
        
    if p1 in inherit.index:
        cp1 = inherit.loc[p1]
    else:
        cp1 = 0

    if p2 in inherit.index:
        cp2 = inherit.loc[p2]
    else:
        cp2 = 0

    cp_sum = cp1+cp2    
    ratio = cp1/cp_sum
    
    INHERIT.append([cross, s, 'whole_genome', 'whole_genome', p1, p2, cp1, cp2, ref_genome_length, ratio])

INHERIT = pd.DataFrame(INHERIT, columns=['cross','strain','annot_type','Name','p1','p2','cp1','cp2','len','ratio'])

# %% [code]
#produce discrete counts
def generate_discrete_call(x, threshold=0.1):
    
    p1, p2, ratio = x[['p1','p2','ratio']]
    if np.isnan(ratio):
        call = 'none'
    else:
        if ratio >= 1-threshold:
            call = p1
        elif ratio <= threshold:
            call = p2
        else:
            call = 'rec'

    return call

# %% [code]
INHERIT['call'] = INHERIT.apply(lambda x: generate_discrete_call(x), axis=1)

# %% [code]
# compute mean inheritance across all mtDNA
INH_WHOLE_GENOME = INHERIT.loc[INHERIT['annot_type']=='whole_genome'].groupby('cross')['ratio'].mean()

# %% [code]
#import mito CNs 
mito_cn = []
for cross, P in parents_dict.items():
    p1, p2 = [p.split('.')[-1] for p in P]
    cn = {}
    for p in (p1, p2): 
        with open(f'/mnt/HDD3/mito_nanopore/mito_cn/mito_cn_{p}.txt') as handle:
            cn[p] = [p] + [float(i) for i in handle.read().split(',')]
    
    ratio = cn[p1][2]/(cn[p1][2]+cn[p2][2])
    mito_cn.append([cross, ratio] + cn[p1] + cn[p2])

mito_cn = pd.DataFrame(mito_cn, columns=['cross','ratio','p1','mt_p1','nuc_p1','cn_p1','p2','mt_p2','nuc_p2','cn_p2']).set_index('cross', drop=False)

# %% [markdown]
## Fig S20

# %% [code]
X = mito_cn.loc[cross_order, 'ratio']
Y = INH_WHOLE_GENOME.loc[cross_order]
lr = stats.linregress(X, Y)

fig, ax = plt.subplots(figsize=(5,4))
ax.scatter(X, Y, color=[cross_color[cross] for cross in cross_order])

X = np.array([X.min(), X.max()])
ax.plot(X, X*lr.slope+lr.intercept, color='black', zorder=-1)
ax.text(0.3, 0.3, f'r={lr.rvalue:.3f}\np={lr.pvalue:.3f}', transform=ax.transAxes)
ax.set_xlabel('mt CN ratio')
ax.set_ylabel('mt inheritance ratio')

legend_elms = [Line2D([0], [0], color='white', marker='o', ms=8, mfc=cross_color[cross], label=cross) for cross in cross_order]
ax.legend(handles=legend_elms, ncol=1, loc=3, bbox_to_anchor=[1.02,0.12])

sns.despine()
plt.tight_layout()

#plt.savefig(f'/home/mathieu/mhenault_landrylab/Publications/mito_ma/resubmission2_GRes/fig/FigS4_mito_cn.svg')
#plt.show()
plt.close()

# %% [markdown]
## Binomial tests for deviation from mtDNA-wide inheritance ratios

# %% [code]
INH_BIN = []
for (cross, f, p1, p2), df in INHERIT.loc[(INHERIT['annot_type'].isin(['gene','rna'])) & 
                                          (~INHERIT['call'].isin(['rec','none']))].groupby(['cross','Name','p1','p2']):
    ref_ratio = INH_WHOLE_GENOME.loc[cross]
    inherit = df.value_counts('call')
    if p1 in inherit.index:
        cp1 = inherit.loc[p1]
    else:
        cp1 = 0
    if p2 in inherit.index:
        cp2 = inherit.loc[p2]
    else:
        cp2 = 0
    bin_pval = stats.binom_test(cp1, cp1+cp2, ref_ratio)
    INH_BIN.append([cross, f, ref_ratio, cp1/(cp1+cp2), bin_pval])
INH_BIN = pd.DataFrame(INH_BIN, columns=['cross','feature','ref_ratio','ratio','pval'])

# %% [code]
INH_BIN['ratio_corr'] = INH_BIN['ratio']-INH_BIN['ref_ratio']
INH_BIN['pval_corr'] = multipletests(INH_BIN['pval'], method='fdr_bh')[1]

# %% [markdown]
## Fig S21

# %% [code]
fig = plt.figure(figsize=[9,6])
gs = plt.GridSpec(ncols=5, nrows=2, hspace=0.5, wspace=0.4,
                 left=0.08, bottom=0.13, top=0.95, right=0.97)

feat_order = dict(zip(artificial_genome_feat.sort_values(by=3).loc[artificial_genome_feat['annot_type'].isin(['gene','rna']), 'Name'].values,
                     range(11)))
cross_ax = dict(zip(['CC1','CC2','CC3','BB1','BB2','BC1','BC2','BA2','BSc1','BSc2'], itertools.product(range(2), range(5))))

for cross, df in INH_BIN.groupby('cross'):

    if cross != 'BA1':
        x,y = cross_ax[cross]
        #plot corrected ratios

        ref_ratio = INH_WHOLE_GENOME[cross]
        bound = np.array([-0.5, 0.5]) + ref_ratio
        bound_ticks_labels = np.array([bound[0], ref_ratio, bound[1]])
        if bound_ticks_labels[0] < 0:
            bound_ticks_labels[0] = 0
        if bound_ticks_labels[2] > 1:
            bound_ticks_labels[2] = 1
        bound_ticks = bound_ticks_labels - ref_ratio

        ax = fig.add_subplot(gs[x,y])
        sns.barplot(x='feature', y='ratio_corr', order=feat_order, color=cross_color[cross], data=df,
                    ax=ax, zorder=1, linewidth=0)

        ax.set_facecolor((0,0,0,0))
        ax.set_xticklabels([pres_abs_poly_alias[x] for x in feat_order], size=8, rotation=90)
        ax.set_xlabel('')

        ax.set_ylim(-0.5, 0.5)
        ax.set_yticks(bound_ticks)
        ax.set_yticklabels([f'{y:.2f}' for y in bound_ticks_labels])
        if y == 0:
            ax.set_ylabel('Inheritance ratio')
        else:
            ax.set_ylabel('')

        ax.set_title(cross, size=14)

        # add p-values
        for i in df.index:
            feat, ratio, pval = df.loc[i, ['feature','ratio_corr','pval_corr']]
            if ratio < 0:
                y = ratio-0.05
            elif ratio > 0:
                y = ratio+0.05
            ax.text(feat_order[feat], y-0.02, plot_pval_symbol(pval), ha='center', va='center', size=12, color='k', zorder=2)
        
        #add background patches
        p1, p2 = parents_dict[cross]
        for p, sign, va1, va2 in zip([p1, p2], [1, -1], ['bottom','top'], ['top', 'bottom']):
            p = p.split('.')[-1]
            fa = Rectangle((-0.5, 0), 11, 0.5*sign, fc=parents_color[p], ec='white', alpha=0.15, zorder=0)
            ax.add_patch(fa)
            ax.text(0.5, 0.5+(sign*0.48), p, va=va2, ha='center', size=10, color='k', transform=ax.transAxes)

sns.despine()

#plt.savefig('/home/mathieu/mhenault_landrylab/Publications/mito_ma/resubmission2_GRes/fig/inheritance_ratios.svg')
#plt.show()
plt.close()

# %% [markdown]
# Growth data

# %% [code]
#define dictionaries for conditions
cond_order = ['YPD_25','YPD_37','YPEG_25','YPEG_37']
cond_color = dict(zip(cond_order, ['1.0', '0.8', '0.6', '0.4']))

# %% [code]
#import growth data from gitter
gitter = pd.read_csv('/home/mathieu/mhenault_landrylab/Experiments/screen_glycerol/results/gitter_mean.tsv', sep='\t', index_col=0)

gitter['strain_passage'] = gitter.apply(lambda x: f'{x["strain"]}_P{x["passage"]:.0f}', axis=1)

gitter_pivot = gitter.loc[gitter['contamination_filter']].pivot_table(index='strain_passage', columns='condition', values='auc', aggfunc=lambda x: x)
# exclude strain with no growth
gitter_pivot.loc[gitter_pivot['YPD_25']<10] = np.nan
# nan-out strains which had no cells printed
for sp in ['J41_P35','J9_P35','K40_P35','L32_P35','I41_P35','A15_P35','D74_P35','B89_P35','F80_P35','F16_P35','F24_P35']:
    gitter_pivot.loc[sp] = np.nan

# %% [code]
#integrate growth data with deletion and score respiration
GDR = gitter_pivot.copy()

for sp in GDR.index:
    if sp in ma_strains.index:
        df = ma_strains.loc[sp]
        if df.shape[0] == 1:
            if df.loc['mt_hap_del'].isna():
                print(sp)
    s, p = sp.split('_')
    GDR.loc[sp, 'strain'] = s
    GDR.loc[sp, 'passage'] = p

for (sp, cross, mthd), df in ma_strains.loc[~ma_strains['mt_hap_del'].isna()].groupby(['strain_passage','cross','mt_hap_del']):
    if sp in GDR.index:
        if df.shape[0] == 1:
            GDR.loc[sp, 'cross'] = cross
            GDR.loc[sp, 'mt_hap_del'] = mthd
        else:
            GDR.loc[sp, 'cross'] = cross
            GDR.loc[sp, 'mt_hap_del'] = sp
for sp, df in ma_strains.groupby('strain_passage'):
    if sp in GDR.index:
        idf = set(df['identity_filter'])
        if False in idf:
            GDR.loc[sp, 'identity_filter'] = False
        else:
            GDR.loc[sp, 'identity_filter'] = True
# add cross info for all lines, regardless of sequencing data
GDR['Cross'] = gitter.groupby('strain_passage').apply(lambda x: x['cross'].iloc[0]).loc[GDR.index]
#thermotolerance
gd_tests = {'therm_ypd':['YPD_37','YPD_25'],
           'therm_ypeg':['YPEG_37','YPEG_25'],
           'resp_25':['YPEG_25','YPD_25'],
           'resp_37':['YPEG_37','YPD_37']}
gd_tests_alias = {'therm_ypd':'thermotolerance YPD',
                  'therm_ypeg':'thermotolerance YPEG',
                  'resp_25':'respiration 25°C',
                  'resp_37':'respiration 37°C'}
for t, (c1, c2) in gd_tests.items():
    GDR[t] = GDR[c1]/GDR[c2]
GDR['YPEG'] = GDR[['YPEG_25','YPEG_37']].mean(axis=1)

GDR['resp'] = GDR['YPEG'] > 25
GDR.loc[GDR['YPEG'].isna(), 'resp'] = np.nan

#manually correct respiration data
resp_manual_correction = ['D83_P1', 'D87_P35', 'D93_P35', 'D4_P35', 'A8_P35',
                          'J32_P35', 'K30_P35', 'L41_P35', 'J8_P35', 'J49_P35', 'L47_P35']
for sp in resp_manual_correction:
    GDR.loc[sp, 'resp'] = True

# %% [markdown]
# mtDNA deletions
## Input depth of coverage files on mtDNA

# %% [code]
DEPTH = {}
idx = 0
with ProgressBar(max_value=ma_strains.shape[0]) as bar:
    for s in ma_strains.index:
        depth = pd.read_csv(f'/mnt/HDD3/mito_ma/depth/{s}.mt.rmdup.rg.depth', sep='\t', header=None)

        depth.columns = ['chrom','pos','depth']
        DEPTH[s] = depth
        idx += 1
        bar.update(idx)

# %% [markdown]
## Bin depth in 100 bp windows

# %% [code]
DEPTH_BIN = {}
idx = 0
with ProgressBar(max_value=len(DEPTH)) as bar:
    for s, depth in DEPTH.items():
        if s in depth_nuclear.index:
            depth['bin'] = pd.cut(depth['pos'], bins=pd.interval_range(start=0, end=82500, freq=100))
            depth['Bin'] = depth['bin'].apply(lambda x: x.mid)
            depth = depth.groupby('Bin')['depth'].median().rename('depth_median').reset_index()
            depth['depth_median_norm'] = depth['depth_median']/depth_nuclear.loc[s, 'median']
            depth['strain'] = s
            DEPTH_BIN[s] = depth
        idx += 1
        bar.update(idx)

# %% [code]
# add functional annotations to midpoints of bins used to score coverage depth
bin_func_annot = {}
for b in set(DEPTH_BIN['HI.4802.001.N701---N502.C68_P1']['Bin']):
    F = list(gft[b])
    if len(F) > 0:
        F = artificial_genome_feat.loc[[f[2] for f in F]]
        at = F.sort_values(by=['annot_plot_order'], ascending=False).iloc[0,16]
        bin_func_annot[b] = at
    else:
        bin_func_annot[b] = 'other'
bin_func_annot = pd.Series(bin_func_annot)

# %% [code]
# add functional annotations to individual positions
pos_func_annot = {}
for p in range(1, len(artificial_genome.seq)+1):
    F = list(gft[p])
    if len(F) > 0:
        F = artificial_genome_feat.loc[[f[2] for f in F]]
        idx = F.sort_values(by=['annot_plot_order'], ascending=False).index[0]
        pos_func_annot[p] = idx
    else:
        pos_func_annot[p] = -1
pos_func_annot = pd.Series(pos_func_annot)

# %% [code]
for s, db in DEPTH_BIN.items():
    db['func'] = bin_func_annot.loc[db['Bin']].values
    DEPTH_BIN[s] = db

# %% [markdown]
## Compute coverage statistics to identify deleted haplotypes

# %% [code]
DBF = []
for s, db in DEPTH_BIN.items():
    if ma_strains.loc[s, 'mtdel_filter']:
        # define 3 floor threshold for relative depth of coverage (0.1 is retained)
        for t in 0.01, 0.05, 0.1:
            p = db.groupby('func').apply(lambda x: np.mean(x['depth_median_norm']>t))
            DBF.append(pd.concat([pd.Series([s,t], index=['filename','threshold']), p]))

DBF = pd.concat(DBF, axis=1).T

for c in ['cross','strain','passage','strain_passage']:
    DBF[c] = ma_strains.loc[DBF['filename'], c].values

ft_list = ['exon','intron','orf','other','rna_exon','tRNA']
for (cross,t), df in DBF.groupby(['cross','threshold']):
    for ft in ft_list:
        DBF.loc[df.index, ft] = (df[ft]-df[ft].mean())/df[ft].std()

DBF['ft_sum'] = DBF[ft_list].sum(axis=1)

# %% [code]
# manualy define thresholds corresponding to deleted haplotypes
ft_thres = {'CC1':-11,
           'CC2':-9,
           'CC3':-10,
           'BB1':-10,
           'BB2':-5,
           'BC1':-4.5,
           'BC2':-4,
           'BA1':-5,
           'BA2':0,
           'BSc1':1,
           'BSc2':0.5}

# %% [code]
for cross, df in DBF.loc[DBF['threshold']==0.1].groupby('cross'):
    t = ft_thres[cross]
    DBF.loc[df.index, 'complete'] = df['ft_sum'] > t

# %% [code]
#manually correct some complete profiles
DBF.loc[(DBF['filename']=='HI.4802.002.N703---N504.B7_P1') & (DBF['threshold']==0.1), 'complete'] = True
DBF.loc[(DBF['filename']=='NS.1249.002.N703---N503.E19_P1') & (DBF['threshold']==0.1), 'complete'] = True

# %% [markdown]
## Fig 4

# %% [code]
for (pis, pis_alias) in zip([True,False], ['', '.no_is']):

    plot_intraspecific = pis

    fig = plt.figure(figsize=[9,9])
    gs = plt.GridSpec(ncols=3, nrows=3, width_ratios=[4,4,5], wspace=0.65, hspace=0.45,
                      left=0.08, right=0.97, top=0.95, bottom=0.06)

    ### Respiration loss ###

    ax = fig.add_subplot(gs[0,0])

    df = GDR.loc[GDR['identity_filter']!=False].value_counts(['Cross','resp']).rename('count').reset_index()\
    .pivot_table(index='Cross', columns='resp', values='count', aggfunc=lambda x: x)
    df['ratio'] = df[False]/(df[True] + df[False])*100
    ax.barh(range(11), df.loc[cross_order, 'ratio'], color=[cross_color[cross] for cross in cross_order])
    for (cross,y) in cross_order.items():
        x = df.loc[cross, 'ratio']
        total = df.loc[cross, True] + df.loc[cross, False]
        ax.text(x+1, y, f'{x:.1f}% ({total:.0f})', size=9, ha='left', va='center')
    ax.set_yticks(range(11))
    ax.set_yticklabels(cross_order)
    ax.invert_yaxis()
    ax.set_xlim(0,49)
    ax.set_xlabel('% respiration loss')

    ### mtDNA deletion ###

    ax = fig.add_subplot(gs[0,1])

    df = DBF.loc[DBF['threshold']==0.1].copy()
    S = list(set(df['filename']))

    df1 = df.value_counts(['cross','complete']).rename('count').reset_index().pivot_table(index='cross', columns='complete', values='count', aggfunc=lambda x: x).fillna(0)
    df1['ratio'] = df1[False]/(df1[True] + df1[False])*100

    ax.barh(range(11), df1.loc[cross_order, 'ratio'], color=[cross_color[cross] for cross in cross_order])
    for (cross,y) in cross_order.items():
        x = df1.loc[cross, 'ratio']
        total = df1.loc[cross, True] + df1.loc[cross, False]
        ax.text(x+1, y, f'{x:.1f}% ({total:.0f})', size=9, ha='left', va='center')
    ax.set_yticks(range(11))
    ax.set_yticklabels(cross_order)
    ax.invert_yaxis()
    ax.set_xlim(0,49)
    ax.set_xlabel('% deleted mtDNAs')

    ### Correlation of deletion with genetic distance ###

    ax = fig.add_subplot(gs[0,2])
    dist = []
    for cross in cross_order:
        dist.append(parents_dist[cross])
    ax.scatter(dist, df1.loc[cross_order, 'ratio'], color=[cross_color[cross] for cross in cross_order])

    lr = stats.linregress(dist, df1.loc[cross_order, 'ratio'])
    X = (0,0.15)
    ax.plot(X, [(x*lr.slope)+lr.intercept for x in X], c='k', zorder=-1)
    ax.text(0.65, 0.45, f'r={lr.rvalue:.2f}\np={plot_pval_text(lr.pvalue)}', color='k', transform=ax.transAxes)

    if plot_intraspecific:
        lr = stats.linregress(dist[:-2], df1.loc[list(cross_order)[:-2], 'ratio'])
        X = (0,0.04)
        ax.plot(X, [(x*lr.slope)+lr.intercept for x in X], c='red', zorder=-1)
        ax.text(0.3, 0.7, f'r={lr.rvalue:.2f}\np={plot_pval_text(lr.pvalue)}', color='red', transform=ax.transAxes)

    #ax.set_xlabel('parental divergence')
    ax.set_xlabel('Parental divergence\n(subs site$^{-1}$)')
    ax.set_ylabel('% deleted mtDNAs')

    ### Relation between mtDNA completeness and respiration ###

    growth_dat = []
    for s in S:
        s = s.split('.')[-1]
        if s in GDR.index:
            growth_dat.append(GDR.loc[s, cond_order])
        else:
            growth_dat.append(np.repeat(np.nan, 4))
    growth_dat = pd.DataFrame(growth_dat, index=S, columns=cond_order)
    df = df.set_index('filename').loc[S]
    df = pd.concat([df, growth_dat], axis=1)
    df = pd.melt(df, id_vars=['strain','passage','strain_passage','cross','complete'],
                 value_vars=cond_order, var_name='cond', value_name='AUC').set_index('strain_passage')

    mwu_growth_complete = []
    for (cross, cond), df1 in df.loc[~df['AUC'].isna()].groupby(['cross','cond']):
        s_u = np.nan
        s_v = np.nan
        u = df1.loc[df1['complete']==True, 'AUC']
        if u.shape[0] > 0:
            s_u = (u-u.median()).abs().sort_values().index[0]
        v = df1.loc[df1['complete']==False, 'AUC']
        if v.shape[0] > 0:
            s_v = (v-v.median()).abs().sort_values().index[0]
        if u.shape[0] > 0 and v.shape[0] > 0:
            U, pval = stats.mannwhitneyu(u, v)
            mwu_growth_complete.append([cond, cross, U, pval, s_u, s_v])
        else:
            mwu_growth_complete.append([cond, cross, np.nan, np.nan, s_u, s_v])
    mwu_growth_complete = pd.DataFrame(mwu_growth_complete, columns=['cond', 'cross', 'U', 'pval', 's_u', 's_v'])
    mwu_growth_complete.loc[~mwu_growth_complete['pval'].isna(), 'pval_corr'] = \
    multipletests(mwu_growth_complete.loc[~mwu_growth_complete['pval'].isna(), 'pval'], method='fdr_bh')[1]
    mwu_growth_complete.set_index(['cond','cross'], inplace=True)

    gs2 = plt.GridSpec(ncols=2, nrows=3, height_ratios=[2.2,2,2], wspace=0.2, hspace=0.45,
                      left=0.07, right=0.97, top=0.96, bottom=0.06)

    cond_ax = dict(zip(cond_order, [(1,0), (1,1), (2,0), (2,1)]))
    for cond, df1 in df.groupby('cond'):
        ax = fig.add_subplot(gs2[cond_ax[cond]])
        sns.violinplot(x='cross', y='AUC', hue='complete', scale='width', cut=0, palette={True:'white', False:'grey'}, linewidth=1,
                       data=df1, ax=ax, order=cross_order, hue_order=[True, False])
        for cross, x in cross_order.items():
            pval, s_u, s_v = mwu_growth_complete.loc[(cond, cross), ['pval_corr', 's_u', 's_v']]
            ax.text(x, 40, plot_pval_symbol(pval), size=12, ha='center', va='center')
            for s, x in zip([s_u, s_v], [x-0.24, x+0.24]):
                if not pd.isna(s):
                    img = plt.imread(f'/mnt/HDD3/mito_ma/results/spot_crops/singles/{s}_{cond}.png')
                    imagebox = OffsetImage(img, zoom=0.08)
                    ab = AnnotationBbox(imagebox, (x, -4), pad=0, frameon=False)
                    ax.add_artist(ab)

        ax.set_title(cond_alias[cond], size=14)
        ax.legend_.remove()
        ax.set_ylim(-8,42)
        ax.set_xlabel('')
        ax.set_xticklabels(cross_order, rotation=30, ha='right')
        if cond == 'YPD_25':
            legend_elms = [Line2D([0], [0], color='w', marker='s', mfc=mfc, mec='k', mew=1, ms=12, label=l) 
                           for (l, mfc) in zip(['complete mtDNAs', 'deleted mtDNAs'], ['white','grey'])]
            ax.legend(handles=legend_elms, loc=2, bbox_to_anchor=(0.1,0.6), frameon=False)

    sns.despine()

    fig.text(0.02, 0.96, 'A', size=24, weight='bold')
    fig.text(0.35, 0.96, 'B', size=24, weight='bold')
    fig.text(0.68, 0.96, 'C', size=24, weight='bold')
    fig.text(0.02, 0.63, 'D', size=24, weight='bold')

    #if pis == True:
    #    for ext in ['png','pdf','svg']:
    #        plt.savefig(f'/home/mathieu/mhenault_landrylab/Publications/mito_ma/resubmission2_GRes/fig/Fig4{pis_alias}.{ext}', dpi=300)
        #plt.show()
    plt.close()

# %% [code]
# sort lines by deletion profile similarity
STRAIN_ORDER_DELETION = {}

for cross in cross_order:

    strain_order = {}
    for c, df in DBF.loc[(DBF['cross']==cross) & (DBF['threshold']==0.1)].groupby('complete'):
        S = set(df['filename'])
        if len(S) > 2:
            dat = pd.concat([DEPTH_BIN[s] for s in S]).pivot_table(index='strain', columns='Bin', values='depth_median_norm', aggfunc=lambda x: x)
            dendro = hierarchy.dendrogram(hierarchy.linkage(dat.values, metric=lambda u,v: 1-stats.pearsonr(u,v)[0]), no_plot=True)
            strain_order[c] = [dat.index[i] for i in dendro['leaves']]
        else:
            strain_order[c] = list(S)
    STRAIN_ORDER_DELETION[cross] = strain_order

# %% [code]
#with open('/mnt/HDD3/mito_ma/results/STRAIN_ORDER_DELETION.pkl', 'wb') as handle:
#    pkl.dump(STRAIN_ORDER_DELETION, handle)
with open('/mnt/HDD3/mito_ma/results/STRAIN_ORDER_DELETION.pkl', 'rb') as handle:
    STRAIN_ORDER_DELETION = pkl.load(handle)

# %% [markdown]
## Fig S22-32

# %% [code]
for cross in cross_order:

    fig = plt.figure(figsize=[12,12])
    gs = plt.GridSpec(nrows=3, ncols=3, height_ratios=[15,1,1], width_ratios=[5, 1, 0.4], wspace=0.5, hspace=0.2,
                     top=0.93, bottom=0.03, right=0.98, left=0.09)

    strain_order = STRAIN_ORDER_DELETION[cross]

    if False in strain_order.keys():
        S = strain_order[True]+strain_order[False]
    else:
        S = strain_order[True]

    S = list(parents_dict[cross]) + S
    dat = pd.concat([DEPTH_BIN[s] for s in S]).pivot_table(index='strain', columns='Bin', values='depth_median_norm', aggfunc=lambda x: np.log2(x+0.05))

    ax = fig.add_subplot(gs[0,0])
    HM1 = ax.imshow(dat.loc[S], cmap='viridis', vmin=-4, vmax=4,
                    aspect='auto', interpolation='nearest')
    
    ax.set_title(cross, size=16)
    ax.set_xticks(range(0,830,100))
    ax.set_xticklabels(range(0,83,10))
    ax.set_xlim(-0.5, 825-0.5)
    ax.set_yticks(range(len(S)))
    ax.set_yticklabels(ma_strains.loc[S, 'mt_hap_del'], size=7)

    #deletion data
    for y, s in enumerate(S):
        if (s in strain_order[True]) or (s in parents_dict[cross]):
            mec = (1,1,1,0)
            mfc = 'k'
        else:
            mec = 'k'
            mfc = 'w'
        ax.scatter(860, y, marker='s', color=mfc, edgecolor=mec, lw=1, s=24, clip_on=False)
    ax.text(860, len(S)+1, 'complete', rotation=90, ha='center', va='top')

    # plot growth data
    ax = fig.add_subplot(gs[0,1])
    growth_dat = []
    resp_dat = {}
    for s in S:
        sp = s.split('.')[-1]

        if sp in GDR.index:
            growth_dat.append(GDR.loc[sp, cond_order])
            resp_dat[s] = GDR.loc[sp, 'resp']
        else:
            growth_dat.append(np.repeat(np.nan, 4))
            resp_dat[s] = np.nan

    growth_dat = pd.DataFrame(growth_dat, index=S, columns=gitter_pivot.columns)
    
    HM2 = ax.imshow(growth_dat.loc[S], cmap='magma', vmin=0, vmax=38,
                    aspect='auto', interpolation='nearest')
    
    ax.set_xticks(range(4))
    ax.set_xticklabels([cond_alias[c] for c in growth_dat.columns], rotation=90)
    ax.set_xlim(-0.5, 3.5)
    ax.set_yticks(range(len(S)))
    ax.set_yticklabels(ma_strains.loc[S, 'mt_hap_del'], size=7)

    #respiration data
    for y, s in enumerate(S):
        if (resp_dat[s] == True) or (s in parents_dict[cross]):
            mec = (1,1,1,0)
            mfc = 'k'
        elif resp_dat[s] == False:
            mec = 'k'
            mfc = 'w'
        else:
            mec = (1,1,1,0)
            mfc = (1,1,1,0)
        ax.scatter(4.5, y, marker='s', color=mfc, edgecolor=mec, lw=1, s=24, clip_on=False)
    ax.text(4.5, len(S)+1, 'respiration', rotation=90, ha='center', va='top')

    #colony images
    ax = fig.add_subplot(gs[0,2])
    img = PIL.Image.open(f'/mnt/HDD3/mito_ma/results/spot_crops/{cross}_assembled.png')
    ax.imshow(np.asarray(img))
    
    off_yticks = img.size[1]/len(S)
    ax.set_yticks((np.arange(len(S))*off_yticks) + 0.5*off_yticks)
    ax.set_yticklabels(ma_strains.loc[S, 'mt_hap_del'], size=7)

    off_xticks = img.size[0]/4
    ax.set_xticks((np.arange(4)*off_xticks) + 0.5*off_xticks)
    ax.set_xticklabels([cond_alias[c] for c in growth_dat.columns], rotation=90, size=7)

    # plot GFF entries
    ax = fig.add_subplot(gs[1,0])

    for i in ['left','right','top']:
        ax.spines[i].set_visible(False)
    ax.arrow(0, 1, ref_genome_length, 0, color='0.9', lw=0, width=1, head_length=0, head_width=0, zorder=-1)

    legend_elms = []
    for at in annot_type_order:
        if at != 'gc_cluster':
            l = annot_type_alias[at]
            fc = annot_type_color[at]
            lw = 0
            if fc == 'white':
                lw = 1
            legend_elms.append(Line2D([0], [0], color='w', marker='s', ms=9, mfc=fc, mec='black', mew=lw, label=l))
    ax.legend(handles=legend_elms, ncol=6, loc=9, bbox_to_anchor=(0.5, 1.4), frameon=False,
             borderpad=0.0, labelspacing=0.2, handlelength=0.3, columnspacing=1.5)

    for i in artificial_genome_feat.loc[~artificial_genome_feat['annot_type'].isin(['gc_cluster','other'])].\
    sort_values(by='annot_plot_order', ascending=False).index:
        start, end, color, w, z = artificial_genome_feat.loc[i, [3,4,'annot_color','annot_width','annot_plot_order']]
        ax.arrow(start, 1, end-start, 0, color=color, lw=0, width=w, head_length=0, head_width=0, zorder=z)
    ax.set_xlim(xlim)
    ax.set_ylim(0.45, 3)
    ax.set_xticks(np.arange(0, 81e3, 1e4))
    ax.set_xticklabels(range(0,90,10))
    ax.set_xlabel('kb')
    ax.set_yticks([])
    
    if cross in ['BA1', 'BA2']:
        plot_rearrangement_SpA()

    ax_cbar = fig.add_axes([0.28, 0.07, 0.15, 0.01])
    cb = plt.colorbar(HM1, cax=ax_cbar, orientation='horizontal', ticks=np.linspace(-4,4,5), label='log$_2$ normalized depth of coverage')
    cb.outline.set_visible(False)

    ax_cbar = fig.add_axes([0.7, 0.07, 0.15, 0.01])
    cb = plt.colorbar(HM2, cax=ax_cbar, orientation='horizontal', ticks=np.linspace(0,35,6), label='growth (AUC)')
    cb.outline.set_visible(False)

    fig.text(0.02, 0.95, 'A', weight='bold', size=24)
    fig.text(0.66, 0.95, 'B', weight='bold', size=24)
    fig.text(0.88, 0.95, 'C', weight='bold', size=24)
    
    #plt.savefig(f'/home/mathieu/mhenault_landrylab/Publications/mito_ma/resubmission2_GRes/fig/detailed_depth_hm_{cross}.svg', dpi=300)
    #plt.show()
    plt.close()

# %% [markdown]
## Compute median depth per gene

# %% [code]
mean_depth_genes = {}

idx = 0
with ProgressBar(max_value=len(DEPTH)) as bar:
    for s, db in DEPTH.items():

        cross = ma_strains.loc[s, 'cross']
        if (cross in cross_order) or (cross == 'P'):

            mdg = {}

            db = db.copy()
            db['func_idx'] = pos_func_annot.loc[db['pos']].values
            db['Name'] = artificial_genome_feat.loc[db['func_idx'], 'Name'].values
            #collapse them gene annotations
            db['Name'] = np.where(db['Name'].apply(lambda x: x[:5]=='cob-E'), 'cob', db['Name'])
            db['Name'] = np.where(db['Name'].apply(lambda x: x[:6]=='cox1-E'), 'cox1', db['Name'])
            db['Name'] = np.where(db['Name'].apply(lambda x: x[:5]=='rnl-e'), 'rnl', db['Name'])
            db['Name'] = np.where(db['Name'].apply(lambda x: x[:3]=='trn'), 'trna', db['Name'])

            for name, df in db.groupby('Name'):
                mdg[name] = df['depth'].median()/depth_nuclear.loc[s, 'median']

            mean_depth_genes[s] = pd.Series(mdg)

        idx += 1
        bar.update(idx)

# %% [markdown]
## Fig 5

# %% [code]
MDG_GITTER = {}

fig = plt.figure(figsize=[9,9])
gs = plt.GridSpec(ncols=4, nrows=3, hspace=0.6, wspace=0.05, left=0.02, right=0.98, bottom=0.1, top=0.96)

cross_ax_dict = dict(zip(cross_order, itertools.product(range(3), range(4))))

for cross, df in ma_strains.loc[ma_strains['mtdel_filter']].groupby('cross'):
    strain_order = STRAIN_ORDER_DELETION[cross]
    
    if False in strain_order:
        S = strain_order[True] + strain_order[False]
    else:
        S = strain_order[True]
    
    S = list(parents_dict[cross]) + S
    
    # coverage depth data
    depth_dat = pd.concat([mean_depth_genes[s] for s in S], axis=1).T
    depth_dat.index = S
    depth_dat = depth_dat.loc[S, genes_to_display]

    depth_dat = depth_dat.applymap(lambda x: np.log2(x+0.05))
    
    # growth data
    growth_dat = []
    for s in S:
        s = s.split('.')[-1]
        if s in gitter_pivot.index:
            growth_dat.append(gitter_pivot.loc[s])
        else:
            growth_dat.append(np.repeat(np.nan, 4))
        
    growth_dat = pd.DataFrame(growth_dat, index=S, columns=gitter_pivot.columns)
    
    dat = pd.concat([depth_dat, growth_dat], axis=1)
    
    MDG_GITTER[cross] = dat
    
    #plot

    ax = fig.add_subplot(gs[cross_ax_dict[cross]])
    
    mask = np.ndarray(dat.shape)
    mask[:, :-4] = False
    mask[:, -4:] = True
    hm = np.ma.MaskedArray(dat, mask)
    HM1 = ax.imshow(hm, aspect='auto', interpolation='nearest', cmap='viridis', vmin=-4, vmax=4)
    
    mask = np.ndarray(dat.shape)
    mask[:, :-4] = True
    mask[:, -4:] = False
    hm = np.ma.MaskedArray(dat, mask)
    HM2 = ax.imshow(hm, aspect='auto', interpolation='nearest', cmap='magma', vmin=0, vmax=38)

    ax.axvline(11.5, color='white', lw=2, zorder=2)
    
    ax.set_xticks(range(16))
    ax.set_xticklabels([genes_alias[i] for i in genes_to_display] + [cond_alias[i] for i in growth_dat.columns], rotation=90, size=9)
    ax.set_yticks([])
    
    ax.spines['top'].set_visible(False)
    ax.spines['left'].set_visible(False)
    ax.spines['right'].set_visible(False)
    ax.spines['bottom'].set_visible(False)
    
    ax.set_title(cross, size=14)
    
    for y, s in enumerate(S[:2]):
        ax.plot([-0.5, -1.5], [y, 5*y], color='k', clip_on=False, lw=0.5, zorder=0.5)
        ax.scatter(-1.5, 5*y, s=24, color=parents_color[s.split('.')[-1]], clip_on=False, zorder=1)

ax_cbar = fig.add_axes([0.78, 0.25, 0.15, 0.015])
cb = plt.colorbar(HM1, cax=ax_cbar, orientation='horizontal', ticks=np.linspace(-4,4,5), label='log$_2$ normalized\ndepth of coverage')
cb.outline.set_visible(False)

ax_cbar = fig.add_axes([0.78, 0.12, 0.15, 0.015])
cb = plt.colorbar(HM2, cax=ax_cbar, orientation='horizontal', ticks=np.linspace(0,35,6), label='growth (AUC)')
cb.outline.set_visible(False)

#for ext in ['svg','pdf','png']:
#    plt.savefig(f'/home/mathieu/mhenault_landrylab/Publications/mito_ma/resubmission2_GRes/fig/Fig5.{ext}', dpi=300)

#plt.show()
plt.close()

# %% [markdown]
# Associations between aspects of mtDNA evolution
## Parse nuclear vcf files from Fijarczyk et al. 2021

# %% [code]
# import raw vcf files
VCF_MELT_NUC = {}

for cross in cross_order:
    path = f'/mnt/HDD1/Dropbox/denovo_mut/data/denovo_vcf/var_{cross}.fil.vcf.gz'
    vcf = pd.read_csv(path, comment='#', compression='gzip', sep='\t')
    with gzip.open(path) as handle:
        for line in handle.readlines():
            if b'#CHROM' in line:
                header = line.decode().strip('\n').split('\t')
                break
    vcf.columns = header
    samples = vcf.columns[9:]
    #randomly subsample 10k variants
    variants = np.random.choice(vcf.index, 10000)
    vcf = vcf.loc[variants]
    vcf['var_uid'] = [f'v{i}' for i in range(vcf.shape[0])]
    vcf_melt = pd.melt(vcf, id_vars=['var_uid','FORMAT'], value_vars=vcf.columns[9:], var_name='filename', value_name='GT')
    vcf_melt['#CHROM'] = vcf.set_index('var_uid').loc[vcf_melt['var_uid'], '#CHROM'].values
    vcf_melt['#POS'] = vcf.set_index('var_uid').loc[vcf_melt['var_uid'], 'POS'].values
    
    for f, df in vcf_melt.groupby('FORMAT'):
        Format = f.split(':')
        for i,tag in enumerate(Format):
            new = df['GT'].apply(lambda x: x.split(':')[i]).replace('.', np.nan)
            if tag in ['DP','AO','RO']:
                new = new.astype(float)
            vcf_melt.loc[df.index, tag.lower()] = new
    
    VCF_MELT_NUC[cross] = vcf_melt
    print(cross)

# %% [code]
for cross in cross_order:
    if cross not in ['BSc1', 'BSc2']:
        vcf_melt = VCF_MELT_NUC[cross]
        p1, p2 = [p.split('.')[-1] for p in parents_dict[cross]]
        idx = 0

        with ProgressBar(max_value=10000) as bar:
            for v, df in vcf_melt.groupby('var_uid'):
                a1 = set(df.set_index('filename').loc[p1, 'gt'].split('/'))
                a2 = set(df.set_index('filename').loc[p2, 'gt'].split('/'))

                allele_order = np.nan
                if a1 == {'0'} and a2 == {'1'}:
                    allele_order = 0
                elif a1 == {'1'} and a2 == {'0'}:
                    allele_order = 1

                vcf_melt.loc[df.index, 'allele_order'] = allele_order

                idx += 1
                bar.update(idx)

        for o, df in vcf_melt.groupby('allele_order'):
            if o == 0:
                vcf_melt.loc[df.index, p1] = df['ad'].apply(lambda x: x.split(',')[0])
                vcf_melt.loc[df.index, p2] = df['ad'].apply(lambda x: x.split(',')[1])
            elif o == 1:
                vcf_melt.loc[df.index, p1] = df['ad'].apply(lambda x: x.split(',')[1])
                vcf_melt.loc[df.index, p2] = df['ad'].apply(lambda x: x.split(',')[0])
        VCF_MELT_NUC[cross] = vcf_melt

# %% [code]
#parse chromosome offset values for reference nuclear genomes
tig_offset = {cross:{} for cross in cross_order}

offset = 0
with open('/home/mathieu/paradoxus_nanopore/paradoxus4/assemblies/paradoxus4_bc02_smartdenovo_pilon-nanopolish_reordered.fasta') as handle:
    for seq in SeqIO.parse(handle, 'fasta'):
        for cross in ['BB1','BB2','BC1','BC2','BA1','BA2']:
            tig_offset[cross][seq.id] = offset
        offset += len(seq.seq)
            
offset = 0
with open('/home/mathieu/paradoxus_nanopore/paradoxus4/assemblies/paradoxus4_bc03_smartdenovo_pilon-nanopolish_reordered.fasta') as handle:
    for seq in SeqIO.parse(handle, 'fasta'):
        for cross in ['CC1','CC2','CC3']:
            tig_offset[cross][seq.id] = offset
        offset += len(seq.seq)

# %% [markdown]
## Inspect allelic ratios along nuclear chromosomes

# %% [code]
depth_per_chrom = []
for cross in cross_order:
    
    vcf_melt = VCF_MELT_NUC[cross]
    dpc = pd.DataFrame(vcf_melt.groupby(['filename','#CHROM'])['dp'].median())
    for s, df in vcf_melt.groupby('filename'):
        dpc.loc[s, 'median_dp'] = df['dp'].median()
    dpc = dpc.reset_index()
    dpc['cross'] = cross
    depth_per_chrom.append(dpc)
    
depth_per_chrom = pd.concat(depth_per_chrom).reset_index(drop=True)

depth_per_chrom['line'] = depth_per_chrom['filename'].apply(lambda x: x.split('_')[0])
for s, df in depth_per_chrom.groupby('filename'):
    if len(s.split('_')) == 2:
        depth_per_chrom.loc[df.index, 'passage'] = int(s.split('_')[1].strip('P'))

# %% [code]
#import table with copy number calls per chromosome from Fijarczyk et al. 2021
fijarczyk_S4 = pd.read_csv('/home/mathieu/mhenault_landrylab/papiers/fijarczyk_gbe2021_SUPP/TableS4_fijarczyk_gbe2021.csv', skiprows=3, header=None)
fijarczyk_S4.columns = ['cross','line','tig','chrom','n_mut','gen','cn_Tini','cn_Tend','mean_cn','fil_len','fil_len*cn']
for s, df in fijarczyk_S4.groupby('line'):
    fijarczyk_S4.loc[df.index, 'major_cn_Tini'] = np.round((df['fil_len']*df['cn_Tini']).sum()/df['fil_len'].sum())
    fijarczyk_S4.loc[df.index, 'major_cn_Tend'] = np.round((df['fil_len']*df['cn_Tend']).sum()/df['fil_len'].sum())
    

fijarczyk_S4.set_index(['line','tig'], inplace=True)

# %% [code]
tig_chrom = {}
for cross, df in fijarczyk_S4.groupby('cross'):
    tig_chrom[cross] = df.groupby('tig').apply(lambda x: x.iloc[0]['chrom'])

# %% [code]
for (s, p, tig), df in depth_per_chrom.groupby(['line','passage','#CHROM']):
    if s in fijarczyk_S4.index:
        if p == 1:
            depth_per_chrom.loc[df.index, 'cn'] = fijarczyk_S4.loc[(s,tig), 'cn_Tini']
            depth_per_chrom.loc[df.index, 'major_cn'] = fijarczyk_S4.loc[(s,tig), 'major_cn_Tini']
        if p > 16:
            depth_per_chrom.loc[df.index, 'cn'] = fijarczyk_S4.loc[(s,tig), 'cn_Tend']
            depth_per_chrom.loc[df.index, 'major_cn'] = fijarczyk_S4.loc[(s,tig), 'major_cn_Tend']

# %% [code]
depth_per_chrom['cn_ratio'] = depth_per_chrom['cn']/depth_per_chrom['major_cn']
depth_per_chrom['dp_ratio'] = depth_per_chrom['dp']/depth_per_chrom['median_dp']
depth_per_chrom['ratio_dev'] = depth_per_chrom['dp_ratio']-depth_per_chrom['cn_ratio']
for cross, df in depth_per_chrom.groupby('cross'):
    depth_per_chrom.loc[df.index, 'chrom'] = tig_chrom[cross].loc[df['#CHROM']].values


# %% [code]
dpc_mean = pd.concat([depth_per_chrom.groupby(['filename','cross','line','passage']).apply(lambda x: np.mean(np.abs(x['ratio_dev']))).rename('mean_ratio_dev'),
                      depth_per_chrom.groupby(['filename','cross','line','passage']).apply(lambda x: np.median(x['median_dp'])).rename('median_dp')], axis=1).reset_index()

# %% [code]
for [cross, p], df in dpc_mean.loc[~dpc_mean['mean_ratio_dev'].isna()].groupby(['cross','passage']):
    #t = np.quantile(df['mean_ratio_dev'], 0.9)
    dpc_mean.loc[df.index, 'high_ratio_dev'] = df['mean_ratio_dev'] >= 0.08

# %% [code]
#import pcr data
pcr_markers = pd.read_csv('/home/mathieu/mhenault_landrylab/Experiments/screen_glycerol/pcr_validation/pcr_markers.csv')

pcr_markers['strain_passage'] = pcr_markers.apply(lambda x: f'{x["strain"]}_P{x["passage"]}', axis=1)
pcr_markers.set_index('strain_passage', inplace=True)

for sp in pcr_markers.index:
    if np.all(pcr_markers.loc[sp, ['atp6','rnl','mat']] == 0):
        pcr_markers.loc[sp, ['atp6','rnl','mat']] = np.repeat(np.nan, 3)

# %% [markdown]
## Fig S39

# %% [code]
chrom_order = ['chrI','chrII','chrIII','chrIV','chrV','chrVI','chrVII','chrVIII','chrIX','chrX','chrXI','chrXII','chrXIII','chrXIV','chrXV','chrXVI']
pcr_marker_color = {0:'red', 1:'white'}


cross = 'BA2'
    
fig = plt.figure(figsize=[10,7])
gs = plt.GridSpec(ncols=4, nrows=1, width_ratios=[16,16,12,4], wspace=0.4, left=0.08, right=0.96, top=0.93, bottom=0.25)

dat_cross = depth_per_chrom.loc[(depth_per_chrom['passage']==1) & 
                                (depth_per_chrom['cross']==cross) & 
                                ~(depth_per_chrom['cn'].isna())]
S = ma_strains.loc[(ma_strains['cross']==cross) & (ma_strains['passage']==1) & (ma_strains['identity_filter']==True)]
S = S.groupby('strain').apply(lambda x: x.iloc[0]['filename'])
strain_order = dat_cross.groupby('line').apply(lambda x: np.mean(np.abs(x['ratio_dev']))).sort_values().index
strain_order = [s for s in S.index if s not in strain_order] + [s for s in strain_order if s in S]

ax = fig.add_subplot(gs[0])

dat = dat_cross.pivot_table(index='line', columns='chrom', values='ratio_dev', aggfunc=lambda x: np.mean(x))
for s in strain_order:
    if s not in dat.index:
        dat.loc[s] = np.nan

HM1 = ax.imshow(dat.loc[strain_order, chrom_order], cmap='bwr', aspect='auto', interpolation='nearest', vmin=-0.5, vmax=0.5)
ax.set_yticks(range(len(strain_order)))
ax.set_yticklabels(strain_order)
ax.set_xticks(range(16))
ax.set_xticklabels(chrom_order, rotation=90)


ax = fig.add_subplot(gs[1])

dat = dat_cross.pivot_table(index='line', columns='chrom', values='cn', aggfunc=lambda x: np.mean(x))
for s in strain_order:
    if s not in dat.index:
        dat.loc[s] = np.nan
HM2 = ax.imshow(dat.loc[strain_order, chrom_order], cmap='binary', aspect='auto', interpolation='nearest', vmin=0, vmax=4)

ax.set_yticks(range(len(strain_order)))
ax.set_yticklabels(strain_order)
ax.set_xticks(range(16))
ax.set_xticklabels(chrom_order, rotation=90)   

# add mito variables 
# coverage depth data
ax = fig.add_subplot(gs[2])

depth_dat = []
for s in strain_order:
    if S.loc[s] in mean_depth_genes:
        depth_dat.append(mean_depth_genes[S.loc[s]])
    else:
        depth_dat.append(pd.Series(np.repeat(np.nan, 12), index=genes_to_display))
depth_dat = pd.concat(depth_dat, axis=1).T
#pd.concat([mean_depth_genes[S.loc[s]] for s in strain_order], axis=1).T
depth_dat.index = strain_order
depth_dat = depth_dat.loc[strain_order, genes_to_display]
depth_dat = depth_dat.applymap(lambda x: np.log2(x+0.05))

HM3 = ax.imshow(depth_dat, aspect='auto', interpolation='nearest', cmap='viridis',
         vmin=-4, vmax=4)
if cross == 'BA2':
    for y, s in enumerate(strain_order):
        sp = S[s].split('.')[-1]
        if sp in pcr_markers.index:
            atp6 = pcr_markers.loc[sp, 'atp6']
            if not np.isnan(atp6):
                ax.scatter(0, y, color=pcr_marker_color[atp6], s=12)
            rnl = pcr_markers.loc[sp, 'rnl']
            if not np.isnan(rnl):
                ax.scatter(4, y, color=pcr_marker_color[rnl], s=12)

ax.set_yticks(range(len(strain_order)))
ax.set_yticklabels(strain_order)
ax.set_xticks(range(12))
ax.set_xticklabels([genes_alias[i] for i in genes_to_display], rotation=90) 

# growth data
ax = fig.add_subplot(gs[3])
growth_dat = gitter_pivot.loc[[f'{s}_P1' for s in strain_order]]

HM4 = ax.imshow(growth_dat, aspect='auto', interpolation='nearest', cmap='magma', vmin=0, vmax=38)
ax.set_yticks(range(len(strain_order)))
ax.set_yticklabels(strain_order)
ax.set_xticks(range(4))
ax.set_xticklabels([cond_alias[i] for i in growth_dat.columns], rotation=90)


ax_cbar1 = fig.add_axes([0.12, 0.1, 0.15, 0.02])
cb = plt.colorbar(HM1, cax=ax_cbar1, orientation='horizontal', ticks=np.linspace(-0.5,0.5,5), label='depth of coverage\ndeviation')
cb.outline.set_visible(False)

ax_cbar2 = fig.add_axes([0.4, 0.1, 0.15, 0.015])
cb = plt.colorbar(HM2, cax=ax_cbar2, orientation='horizontal', ticks=np.arange(0,5), label='called CN')
cb.outline.set_visible(False)

ax_cbar3 = fig.add_axes([0.67, 0.1, 0.15, 0.015])
cb = plt.colorbar(HM3, cax=ax_cbar3, orientation='horizontal', ticks=np.linspace(-4,4,5), label='log$_2$ normalized\ndepth of coverage')
cb.outline.set_visible(False)

ax_cbar4 = fig.add_axes([0.9, 0.1, 0.07, 0.015])
cb = plt.colorbar(HM4, cax=ax_cbar4, orientation='horizontal', ticks=np.linspace(0,38,3), label='growth (AUC)')
cb.outline.set_visible(False)

fig.text(0.02, 0.95, 'A', weight='bold', size=24)
fig.text(0.32, 0.95, 'B', weight='bold', size=24)
fig.text(0.62, 0.95, 'C', weight='bold', size=24)
fig.text(0.85, 0.95, 'D', weight='bold', size=24)

ax = fig.axes[5]
ax.set_xlim(1,4)

#plt.savefig(f'/home/mathieu/mhenault_landrylab/Publications/mito_ma/resubmission2_GRes/fig/depth_hm_nucCN_{cross}.svg', dpi=300)
#plt.show()
plt.close()

# %% [markdown]
## Fig S40

# %% [code]
fig = plt.figure(figsize=[7,7])
gs = plt.GridSpec(ncols=2, nrows=2, hspace=0.4, wspace=0.5, left=0.1, right=0.97, top=0.85, bottom=0.08)

for p, df in dpc_mean.loc[(~dpc_mean['cross'].isin(['BSc1','BSc2'])) &
                         (~dpc_mean['median_dp'].isna()) & 
                         (~dpc_mean['mean_ratio_dev'].isna())].groupby('passage'):
    if p == 1:
        ax = ax_idx = 0
        pa = 'initial timepoint'
    elif p == 35:
        ax = ax_idx = 1
        pa = 'final timepoint'
    else:
        continue
        
    ax = fig.add_subplot(gs[0, ax_idx])
    for cross, df1 in df.groupby('cross'):
        ax.plot(df1['mean_ratio_dev'].sort_values(), np.linspace(0,1,df1.shape[0]), color=cross_color[cross])
    ax.set_title(pa, size=14)
    ax.set_xlabel('mean ratio deviation')
    ax.set_ylabel('cumul')
    ax.set_xlim(0, 0.14)
    
    ax = fig.add_subplot(gs[1, ax_idx])
    df = df.sample(frac=1)
    
    ax.scatter(df['median_dp'], df['mean_ratio_dev'], c=df['cross'].apply(lambda x: cross_color[x]), s=12)
    
    lr = stats.linregress(df['median_dp'], df['mean_ratio_dev'])
    X = np.array([0,200])
    ax.plot(X, np.apply_along_axis(lambda x: x*lr.slope+lr.intercept, 0, X), color='black', zorder=1)
    ax.text(0.72, 0.78, f'r={lr.rvalue:.3f}\np={plot_pval_text(lr.pvalue)}', transform=ax.transAxes)
    
    ax.set_title(pa, size=14)
    ax.set_xlabel('median depth of coverage (X)')
    ax.set_ylabel('mean ratio deviation')
    

fig.text(0.03, 0.88, 'A', weight='bold', size=24)
fig.text(0.03, 0.42, 'B', weight='bold', size=24)

ax_legend = fig.add_axes([0.15,0.9,0.7,0.1])
ax_legend.axis('off')
legend_elms = [Line2D([0], [0], color=cross_color[c], label=c) for c in cross_order]
ax_legend.legend(handles=legend_elms, loc=2, bbox_to_anchor=(0,1), ncol=5, frameon=False)
    
sns.despine()
#plt.savefig('/home/mathieu/mhenault_landrylab/Publications/mito_ma/resubmission2_GRes/fig/ratio_dev_depth.svg', dpi=300)
#plt.show()
plt.close()

# %% [markdown]
## Build a table integrating all the variables of interest per line

# %% [code]
combined_drr = []
for cross, df in ma_strains.groupby('cross'):
    if cross in cross_order:
        for fn in df['filename'].values:
            
            sp = fn.split('.')[-1]
            s, p = sp.split('_')
            p = int(p.strip('P'))
            fnr = fn
            
            # correct fn for clones
            if ma_strains.loc[fn, 'mt_haplotype'] == 'clone':
                fn_list = ma_strains.loc[(ma_strains['strain']==s) & (ma_strains['mt_haplotype']!='clone'), 'filename']
                if fn_list.shape[0] == 1:
                    fnr = fn_list.iloc[0]
                elif fn == 'HI.4802.001.N708---N506.B40_P1':
                    fnr = 'NS.1250.002.N704---N507.B40_P35'
                elif fn == 'NS.1249.002.N712---N504.B55_P10':
                    fnr = 'NS.1250.002.N705---N508.B55_P35'
     
            deletion = np.nan
            recomb = np.nan
            respiration = np.nan
            mito_inst = np.nan
            nuc_inst = np.nan
            mean_ratio_dev = np.nan
            
            # deletion
            if fn in STRAIN_ORDER_DELETION[cross][True]:
                deletion = False
            elif False in STRAIN_ORDER_DELETION[cross]:
                if fn in STRAIN_ORDER_DELETION[cross][False]:
                    deletion = True
            
            # recomb
            if fnr in REC['strain'].values:
                if REC.set_index('strain').loc[fnr, 'count'] > 0:
                    recomb = True
                else:
                    recomb = False
            
            #respiration
            if sp in GDR.index:
                respiration = GDR.loc[sp, 'resp']

            #mito instability
            if (respiration == True and deletion == True) or (respiration == False and deletion == False):
                mito_inst = True
            elif (respiration == True and deletion == False) or (respiration == False and deletion == True):
                mito_inst = False
                
            #nuc instability
            if cross not in ['BSc1', 'BSc2']:
                if sp in dpc_mean['filename'].values:
                    if dpc_mean.set_index('filename').loc[sp, 'high_ratio_dev'] == True:
                        nuc_inst = True
                    if dpc_mean.set_index('filename').loc[sp, 'high_ratio_dev'] == False:
                        nuc_inst = False
                    mean_ratio_dev = dpc_mean.set_index('filename').loc[sp, 'mean_ratio_dev']
            combined_drr.append([s, p, cross, deletion, recomb, respiration, mito_inst, nuc_inst, mean_ratio_dev])
            
combined_drr = pd.DataFrame(combined_drr, columns=['strain','passage','cross','deletion','recomb','respiration','mito_inst','nuc_inst', 'mean_ratio_dev']).reset_index(drop=True)

# %% [code]
# lines with changes in deletion status
deletion_change = []
for (s, cross), df in combined_drr.set_index('passage').loc[[1,35]].groupby(['strain','cross']):
    if df.shape[0] == 2:
        if df.loc[1, 'deletion'] == True and df.loc[35, 'deletion'] == False:
            deletion_change.append([s, cross, 'gain'])
        elif df.loc[1, 'deletion'] == False and df.loc[35, 'deletion'] == True:
            deletion_change.append([s, cross, 'loss'])
        else:
            deletion_change.append([s, cross, 'no_change'])
deletion_change = pd.DataFrame(deletion_change, columns=['strain','cross','change'])
deletion_change['cross_order'] = deletion_change['cross'].apply(lambda x: cross_order[x])
deletion_change['strain_order'] = deletion_change['strain'].apply(lambda x: int(x[1:]))
#add filter for haplotype consistensy
deletion_change['consistent'] = consistent_hap.set_index('strain').loc[deletion_change['strain'], 'consistent'].values

# %% [markdown]
## Fig S34

# %% [code]
ma_strains_sp_index = ma_strains.set_index('strain_passage')
dc = deletion_change.loc[deletion_change['change'].isin(['gain', 'loss'])]
Y_dict = dict(zip(dc.sort_values(by=['cross_order','strain_order'])['strain'], np.arange(0,dc.shape[0])*3))
consistent_deletion = ['J15','J26','J29','J54','J64','L5','L46','B54','E34','E37','C87','F15']

fig = plt.figure(figsize=[8,10])
gs = plt.GridSpec(nrows=2, ncols=1, height_ratios=[14,1], hspace=0.08,
                 left=0.06, right=0.91, top=0.99, bottom=0.05)
xlim = (-500,ref_genome_length+500)

ax = fig.add_subplot(gs[0])
for i in ['right','top','left']:
    ax.spines[i].set_visible(False)
alpha = 0.3

for (s, cross, change), df in dc.groupby(['strain', 'cross', 'change']):
    sp1 = f'{s}_P1'
    sp35 = f'{s}_P35'

    for sp, off in zip([sp1, sp35], [0,1]):
        S = ma_strains_sp_index.loc[sp, 'filename']
        tracts = TRACTS.loc[TRACTS['strain']==S]
    
        y = Y_dict[s]+off
        
        for i in tracts.index:
            (start, end, parent) = tracts.loc[i, ['start','end','parent']]
            c = parents_color[parent.split('.')[-1]]
            if start < end:
                fa = FancyArrow(start, y, end-start, 0, width=0.8, head_length=0, head_width=0, color=c, lw=0, alpha=alpha)
                ax.add_patch(fa)
            elif start > end:
                fa1 = FancyArrow(0, y, end, 0, width=0.8, head_length=0, head_width=0, color=c, lw=0, alpha=alpha)
                fa2 = FancyArrow(start, y, ref_genome_length-start, 0, width=0.8, head_length=0, head_width=0, color=c, lw=0, alpha=alpha)
                ax.add_patch(fa1)
                ax.add_patch(fa2)
        
        ax.text(-700, y, sp.split('_')[1], ha='right', va='center', size=6)
        if sp == sp1:
            ax.text(-3e3, y+0.25, s, ha='right', va='center', size=8)
        
        #add support markers
        markers = MARKERS[cross]
        vcf_melt = VCF_MELT[cross]
        vcf_melt = vcf_melt.loc[(vcf_melt['vma_uid'].isin(markers)) &
                            (vcf_melt['higher']==True) &
                            (vcf_melt['parent'].isin(parents_dict[cross])) &
                            (vcf_melt['filename']==S)]
        
        for parent, df1 in vcf_melt.groupby('parent'):
            c = parents_color[parent.split('.')[-1]]
            ax.scatter(df1['POS'], np.repeat(y, df1.shape[0]), s=5, color=c)
            
    y = Y_dict[s]+0.5
    ax.text(ref_genome_length+5e3, y, change, size=8, ha='left', va='center')
    if s in consistent_deletion:
        ax.text(ref_genome_length+750, y-0.35, '*', size=18, ha='left', va='center')
    
    if change == 'loss':
        (y1, y2) = (y-0.5, y+0.5)
    elif change == 'gain':
        (y1, y2) = (y+0.5, y-0.5)
    ax.scatter(ref_genome_length+3e3, y1, marker='s', color='k', edgecolor=(1,1,1,0), lw=1, s=24, clip_on=False)
    ax.scatter(ref_genome_length+3e3, y2, marker='s', color='w', edgecolor='k', lw=1, s=24, clip_on=False)
        
        
ax.set_xlim(xlim)
ax.set_xticks(np.arange(0, 81e3, 1e4))
ax.set_xticklabels(range(0,90,10))
ax.text(ref_genome_length+3e3, -1, 'complete', rotation=90, ha='center', va='top')
ax.set_yticks([])
ax.set_ylim(-1, 105)

# plot GFF entries
ax = fig.add_subplot(gs[1])
for i in ['left','right','top']:
    ax.spines[i].set_visible(False)
ax.arrow(0, 1, ref_genome_length, 0, color='0.9', lw=0, width=1, head_length=0, head_width=0, zorder=-1)

legend_elms = []
for at in annot_type_order:
    if at != 'gc_cluster':
        l = annot_type_alias[at]
        fc = annot_type_color[at]
        lw = 0
        if fc == 'white':
            lw = 1
        legend_elms.append(Line2D([0], [0], color='w', marker='s', ms=9, mfc=fc, mec='black', mew=lw, label=l))
ax.legend(handles=legend_elms, ncol=6, loc=9, bbox_to_anchor=(0.5, 1.1), frameon=False,
         borderpad=0.0, labelspacing=0.2, handlelength=0.3, columnspacing=1.5)

for i in artificial_genome_feat.loc[~artificial_genome_feat['annot_type'].isin(['gc_cluster','other'])].\
sort_values(by='annot_plot_order', ascending=False).index:
    start, end, color, w, z = artificial_genome_feat.loc[i, [3,4,'annot_color','annot_width','annot_plot_order']]
    ax.arrow(start, 1, end-start, 0, color=color, lw=0, width=w, head_length=0, head_width=0, zorder=z)
ax.set_xlim(xlim)
ax.set_ylim(0.45, 3)
ax.set_xticks(np.arange(0, 81e3, 1e4))
ax.set_xticklabels(range(0,90,10))
ax.set_xlabel('kb')
ax.set_yticks([])

#plt.savefig('/home/mathieu/mhenault_landrylab/Publications/mito_ma/resubmission2_GRes/fig/deletion_change.svg', dpi=300)
#plt.show()
plt.close()

# %% [markdown]
## Fig S41

# %% [code]
fig, axes = plt.subplots(nrows=2, figsize=[7,7],
                        gridspec_kw=dict(hspace=0.3, left=0.15, bottom=0.07, right=0.96, top=0.93))

ax = axes[0]

dat = combined_drr.value_counts(['cross','passage','mito_inst']).rename('count').reset_index().pivot_table(index=['cross','passage'], columns='mito_inst', values='count').fillna(0).reset_index()
dat['perc'] = (dat[True]/(dat[True]+dat[False])*100)
dat['timepoint'] = dat['passage'].replace({1:'initial', 35:'final'})

sns.barplot(x='cross', y='perc', hue='timepoint', palette={'initial':'0.8', 'final':'k'}, 
            order=cross_order, data=dat, ax=ax)

for (cross, tp), df in dat.groupby(['cross','timepoint']):
    x = cross_order[cross]
    if tp == 'initial':
        x -= 0.2
    elif tp == 'final':
        x += 0.2
    
    y, c = df[['perc', True]].iloc[0].values
    if y > 0:
        ax.text(x, y, f'{c:.0f}', ha='center', va='bottom', size=10)

ax.set_ylabel('% lines with mtDNA instability')
ax.set_xlabel('')
ax.legend(loc=2, bbox_to_anchor=(0.1, 1), frameon=False, title='timepoint')

ax = axes[1]
mwu_mito_nuc_inst = []
for cross, df in combined_drr.loc[(~combined_drr['cross'].isin(['BSc1','BSc2'])) &
                                  ~(combined_drr['mean_ratio_dev']).isna()].groupby('cross'):

    u = df.loc[df['mito_inst']==False, 'mean_ratio_dev']
    v = df.loc[df['mito_inst']==True, 'mean_ratio_dev']
    
    if u.shape[0] > 0 and v.shape[0] > 0:
        U, pval = stats.mannwhitneyu(u, v)
        mwu_mito_nuc_inst.append([cross, U, pval])

mwu_mito_nuc_inst = pd.DataFrame(mwu_mito_nuc_inst, columns=['cross', 'U', 'pval'])
mwu_mito_nuc_inst['pval_corr'] = multipletests(mwu_mito_nuc_inst['pval'], method='fdr_bh')[1]
mwu_mito_nuc_inst.set_index('cross', inplace=True)

dat = combined_drr.copy().rename({'mito_inst':'mtDNA instability'})
sns.violinplot(x='cross', y='mean_ratio_dev', hue='mito_inst', order=['CC1','CC2','CC3','BB1','BB2','BC1','BC2','BA1','BA2'],
               palette={False:'white', True:'grey'}, scale='width', cut=0, data=dat, ax=ax)
for cross in mwu_mito_nuc_inst.loc[mwu_mito_nuc_inst['pval_corr']<=0.05].index:
    x = cross_order[cross]
    ax.plot([x-0.3, x+0.3], np.repeat(0.14, 2), color='k')
    ax.text(x, 0.14, plot_pval_symbol(mwu_mito_nuc_inst.loc[cross, 'pval_corr']), va='bottom', ha='center')

ax.set_ylabel('mean coverage depth deviation')
ax.set_ylim(0, 0.155)
ax.set_xlabel('')
ax.legend(loc=2, bbox_to_anchor=(0.1, 1), frameon=False, title='mtDNA instability')
    
fig.text(0.02, 0.94, 'A', size=24, weight='bold')
fig.text(0.02, 0.47, 'B', size=24, weight='bold')
    
sns.despine()

#plt.savefig('/home/mathieu/mhenault_landrylab/Publications/mito_ma/resubmission2_GRes/fig/mito_nuc_instability.svg', dpi=300)
#plt.show()
plt.close()

# %% [markdown]
## Fisher exact tests for pairwise combinations of variables

# %% [code]
fe_combined = []
for v1, v2 in itertools.combinations(['deletion','recomb','respiration','mito_inst','nuc_inst'], 2):
    name = f'{v1}*{v2}'
    for (cross, p), df in combined_drr.groupby(['cross','passage']):
        F = df.value_counts([v1,v2]).rename('count').reset_index().pivot_table(index=v1, columns=v2, values='count', aggfunc=lambda x: x)
        
        if F.shape == (2,2):
            odds, pval = stats.fisher_exact(F.fillna(1))
            fe_combined.append([cross, p, name, odds, pval])
        else:
            fe_combined.append([cross, p, name, np.nan, np.nan])

fe_combined = pd.DataFrame(fe_combined, columns=['cross','passage','test','odds_ratio','pval'])

fe_combined.loc[~fe_combined['pval'].isna(), 'pval_corr'] = multipletests(fe_combined.loc[~fe_combined['pval'].isna(), 'pval'], method='fdr_bh')[1]
fe_combined['log_pval'] = -1*np.log10(fe_combined['pval_corr'])
fe_combined['log_or'] = np.log10(fe_combined['odds_ratio'])

# %% [code]
var_alias = {'deletion':'mtDNA\ndeletion','respiration':'respiration','recomb':'mtDNA\nrecombination','mito_inst':'mtDNA\ninstability','nuc_inst':'aneuploidy\ninstability'}
var_alias_short = {'deletion':'mtDNA del','respiration':'resp','recomb':'mtDNA rec','mito_inst':'mtDNA inst','nuc_inst':'aneup inst'}
var_order_dict = {j:i for i,j in enumerate(['deletion','recomb','respiration','mito_inst','nuc_inst'])}
test_order = ['deletion*recomb',
              'deletion*respiration',
              'deletion*mito_inst',
              'deletion*nuc_inst',
              'recomb*respiration',
             'recomb*mito_inst',
             'recomb*nuc_inst',
             'respiration*mito_inst',
             'respiration*nuc_inst',
             'mito_inst*nuc_inst']
test_order_dict = {j:i for i,j in enumerate(test_order)}
cmap_test = {j:i for i,j in enumerate([t for t in test_order if t in fe_combined.loc[fe_combined['pval_corr']<=0.05, 'test'].values])}

# %% [markdown]
## Fig 6AC

# %% [code]
for p, p_alias in zip([1, 35], ['initial timepoint', 'final timepoint']):
    
    fig= plt.figure(figsize=[4,5])
    gs = plt.GridSpec(ncols=1, nrows=2, height_ratios=[3,2], hspace=0.1, left=0.3, right=0.77, top=0.94, bottom=0.08)
    
    dat = fe_combined.loc[fe_combined['passage']==p].pivot_table(index='cross', columns='test', values='log_or', dropna=False)

    ax = fig.add_subplot(gs[0])
    cbar_ax = fig.add_axes([0.81, 0.5, 0.03, 0.25])
    sns.heatmap(dat.loc[cross_order, test_order], cmap='BrBG', ax=ax, center=0, vmin=-2.6, vmax=2.6, # center=-1*np.log10(0.05), vmax=9,
               cbar_ax=cbar_ax, cbar_kws=dict(label='log$_{10}$ odds ratio'))

    for (test,x), (cross, y) in itertools.product(zip(test_order, range(10)), cross_order.items()):
        pval, odds = fe_combined.set_index(['passage','cross','test']).loc[(p, cross, test), ['pval_corr','log_or']]
        if abs(odds) >= 1.5:
            c = 'white'
        else:
            c = 'black'
        ax.text(x+0.5, y+0.5, plot_pval_symbol(pval), color=c, ha='center', va='center')

    
    ax.set_xticklabels([])
    ax.set_xlabel('')
    ax.set_yticklabels(cross_order, rotation=0)
    ax.set_ylabel('')
    ax.set_title(p_alias, size=14)
    
    for s in ['top', 'bottom', 'right', 'left']:
        ax.spines[s].set_visible(True)
        
    #plot variables
    ax = fig.add_subplot(gs[1])    

    for i,j in itertools.product(range(11), range(5)):
        ax.scatter(i-0.5, j, marker='o', s=8**2, c='white', edgecolors='0.6', zorder=0)
    for t in test_order:
        v1, v2 = t.split('*')

        ax.plot(np.repeat(test_order_dict[t], 2)+0.5, [var_order_dict[v] for v in (v1, v2)], c='k', lw=2,
                marker='o', ms=8, mfc='white', mec='black', mew=2, zorder=1)
    
    ax.set_xticks([])
    ax.set_xlim(0, 10)
    ax.set_yticks(range(5))
    ax.set_yticklabels([var_alias[v] for v in var_order_dict])
    ax.invert_yaxis()
    for s in ['top', 'bottom', 'right']:
        ax.spines[s].set_visible(False)
        
    if p == 1:
        fig.text(0.03, 0.94, 'A', size=24, fontweight='bold')
        fig.text(0.93, 0.94, 'B', size=24, fontweight='bold')
    elif p == 35:
        fig.text(0.03, 0.94, 'C', size=24, fontweight='bold')
        fig.text(0.93, 0.94, 'D', size=24, fontweight='bold')
    
    
    #plt.savefig(f'/home/mathieu/mhenault_landrylab/Publications/mito_ma/resubmission2_GRes/fig/Fig6A_P{p:.0f}.svg', dpi=300)
    #plt.show()
    plt.close()

# %% [markdown]
## Fig 6BD

# %% [code]
for p in [1, 35]:
    
    
    fig= plt.figure(figsize=[3,5])
    gs = plt.GridSpec(ncols=3, nrows=5, hspace=1.7, wspace=1.7, left=0.15, right=0.93, top=0.94, bottom=0.09)

    Dat = fe_combined.loc[(fe_combined['passage']==p) & (fe_combined['pval_corr']<=0.05)].copy()
    Dat['cross_order'] = Dat['cross'].apply(lambda x: cross_order[x])
    Dat['v1'] = Dat['test'].apply(lambda x: x.split('*')[0])
    Dat['v1_order'] = Dat['v1'].apply(lambda x: var_order_dict[x])
    Dat['v2'] = Dat['test'].apply(lambda x: x.split('*')[1])
    Dat['v2_order'] = Dat['v2'].apply(lambda x: var_order_dict[x])
    Dat = Dat.sort_values(by=['v1_order','v2_order','cross_order']).reset_index(drop=True)
    
    if p == 1:
        Dat = pd.concat([Dat, fe_combined.loc[[56,58,60,40,42,31]]])
    if p == 35:
        Dat = pd.concat([Dat, fe_combined.loc[[61,41,43]]])
    Dat['v1'] = Dat['test'].apply(lambda x: x.split('*')[0])
    Dat['v2'] = Dat['test'].apply(lambda x: x.split('*')[1])
    
    for i, (ax_row, ax_col) in zip(Dat.index, itertools.product(range(6), range(3))):
        test, v1, v2, cross = Dat.loc[i, ['test','v1','v2','cross']]

        ax = fig.add_subplot(gs[(ax_row, ax_col)])

        dat = combined_drr.loc[(combined_drr['cross']==cross) & (combined_drr['passage']==p)].value_counts([v1, v2]).rename('count').reset_index()
        dat[v1] = dat[v1].replace({True:'T', False:'F'})
        dat[v2] = dat[v2].replace({True:'T', False:'F'})
        dat = dat.pivot_table(index=v1, columns=v2, values='count', aggfunc=lambda x: x).fillna(0)
        for b in ['T','F']:
            if b not in dat.columns:
                dat[b] = 0
            if b not in dat.index:
                dat.loc[b] = 0

        cmap = LinearSegmentedColormap.from_list('custom', ['white', sns.color_palette()[cmap_test[test]]], N=256)
        sns.heatmap(dat.loc[['F','T'], ['T','F']], cmap=cmap, annot=True, annot_kws={'size':9}, fmt='.0f', cbar=False, ax=ax, vmin=0,
                   linewidths=0.5, linecolor='k')

        ax.set_title(cross, size=10, fontweight='bold')
        ax.set_xticks([0.5, 1.5])
        ax.set_yticks([0.5, 1.5])
        ax.set_xticklabels(['T','F'], size=9, rotation=0)
        ax.set_yticklabels(['F','T'], size=9, rotation=0)

        ax.set_ylabel(var_alias_short[v1], size=9, labelpad=1)
        ax.set_xlabel(var_alias_short[v2], size=9, labelpad=1)
        
    if p == 1:
        R = [Rectangle((0.03, 0.01), 0.94, 0.39, fc='0.85', lw=0, transform=fig.transFigure, zorder=-1)]
        
    if p == 35:
        R = [Rectangle((0.03, 0.01), 0.94, 0.195, fc='0.85', lw=0, transform=fig.transFigure, zorder=-1)]
    fig.patches.extend(R)

    #plt.savefig(f'/home/mathieu/mhenault_landrylab/Publications/mito_ma/resubmission2_GRes/fig/Fig6B_P{p:.0f}.svg', dpi=300)
    #plt.show()
    plt.close()

# %% [markdown]
## Combine growth data with haplotype categories (parental, rec)

# %% [code]
gdrec = []
for cross, df in ma_strains.loc[ma_strains['identity_filter']].groupby('cross'):
    if cross in cross_order:
        for fn in df['filename'].values:
            
            sp = fn.split('.')[-1]
            s, p = sp.split('_')
            p = int(p.strip('P'))
            fnr = fn
            
            # correct fn for clones
            if ma_strains.loc[fn, 'mt_haplotype'] == 'clone':
                fn_list = ma_strains.loc[(ma_strains['strain']==s) & (ma_strains['mt_haplotype']!='clone'), 'filename']
                if fn_list.shape[0] == 1:
                    fnr = fn_list.iloc[0]
                elif fn == 'HI.4802.001.N708---N506.B40_P1':
                    fnr = 'NS.1250.002.N704---N507.B40_P35'
                elif fn == 'NS.1249.002.N712---N504.B55_P10':
                    fnr = 'NS.1250.002.N705---N508.B55_P35'
     
            deletion = np.nan
            recomb = np.nan
            parent = np.nan
            ypd_25 = np.nan
            ypd_37 = np.nan
            ypeg_25 = np.nan
            ypeg_37 = np.nan
            resp = np.nan
            fid = np.nan
            
            # deletion
            if fn in STRAIN_ORDER_DELETION[cross][True]:
                deletion = False
            elif False in STRAIN_ORDER_DELETION[cross]:
                if fn in STRAIN_ORDER_DELETION[cross][False]:
                    deletion = True
            
            # recomb
            if fnr in REC['strain'].values:
                if REC.set_index('strain').loc[fnr, 'count'] > 0:
                    recomb = True
                else:
                    recomb = False
            #major parent
            if recomb == False:
                parent = TRACTS.set_index('strain').loc[fnr, 'parent'].split('.')[-1]
            elif recomb == True:
                parent = 'rec'
            
            #respiration
            if sp in GDR.index:
                ypd_25, ypd_37, ypeg_25, ypeg_37, resp, fid = GDR.loc[sp, ['YPD_25','YPD_37','YPEG_25','YPEG_37','resp','identity_filter']]
                
            gdrec.append([fn, cross, s, p, deletion, recomb, parent, ypd_25, ypd_37, ypeg_25, ypeg_37, resp, fid])

gdrec = pd.DataFrame(gdrec, columns=['filename','cross','strain','passage','deletion','recomb','parent','YPD_25','YPD_37','YPEG_25','YPEG_37','resp','identity_filter'])

# %% [markdown]
## Mann-Whitney U tests between categories of haplotypes

# %% [code]
mwu_growth_rec = []
for cross, df in gdrec.loc[gdrec['resp']==True].groupby(['cross']):
    for cond in cond_order:
        df1 = df.loc[(~df[cond].isna()) & (~df['parent'].isna())]
        p1, p2 = [p.split('.')[-1] for p in parents_dict[cross]]
        order = [p1, 'rec', p2]
        order = [p for p in order if p in df1['parent'].values]
        for p1, p2 in itertools.combinations(order, 2):
            u = df1.loc[df1['parent']==p1, cond]
            v = df1.loc[df1['parent']==p2, cond]
            U, pval = stats.mannwhitneyu(u,v)
            diff = u.median()-v.median()
            mwu_growth_rec.append([cross, cond, p1, p2, U, pval, diff])
mwu_growth_rec = pd.DataFrame(mwu_growth_rec, columns=['cross','cond','p1','p2','U','pval','diff'])
mwu_growth_rec.loc[~mwu_growth_rec['pval'].isna(), 'pval_corr'] = \
multipletests(mwu_growth_rec.loc[~mwu_growth_rec['pval'].isna(), 'pval'], method='fdr_bh')[1]
mwu_growth_rec['log_pval_corr'] = mwu_growth_rec['pval_corr'].apply(lambda x: -1*np.log10(x))
mwu_growth_rec['diff_abs'] = mwu_growth_rec['diff'].abs()

# %% [markdown]
## Fig S35

# %% [code]
fig, axes = plt.subplots(ncols=4, nrows=3, 
                         figsize=[9,9], gridspec_kw=dict(top=0.97, right=0.97, bottom=0.08, left=0.06,
                                                          hspace=0.5, wspace=0.25))
parents_color_rec = parents_color
parents_color_rec['rec'] = 'white'
cross_ax_dict = dict(zip(cross_order, itertools.product(range(3), range(4))))

for cross, df in gdrec.loc[gdrec['resp']==True].groupby('cross'):
    
    p1, p2 = [p.split('.')[-1] for p in parents_dict[cross]]
    order = [p1, 'rec', p2]
    order = [p for p in order if p in df['parent'].values]
    c = [parents_color_rec[x] for x in order]
    df = pd.melt(df, id_vars=['cross', 'parent'], value_vars=cond_order, var_name='cond', value_name='AUC')
    y_max = df.groupby('cond')['AUC'].max().to_dict()
    
    ax_idx = cross_ax_dict[cross]
    ax = axes[ax_idx]
    
    sns.violinplot(x='cond', y='AUC', hue='parent', data=df, scale='width', cut=0, ax=ax,
                   order=cond_order, hue_order=order, palette=c, linewidth=1)
    ax.legend_.remove()
    
    ax.text(0.5, 1, cross, transform=ax.transAxes, size=14, ha='center')
    if ax_idx[1] != 0:
        ax.set_ylabel('')
    ax.set_xlabel('')
    ax.set_xticklabels([cond_alias[c] for c in cond_order], rotation=30, ha='right')
    
    ax.set_ylim(15,40)
    legend_elms = [Line2D([0], [0], color='w', marker='s', mfc=parents_color_rec[x], 
                          ms=9, mec='k', mew=1, label=x) for x in order]
    ax.legend(handles=legend_elms, loc=3, bbox_to_anchor=[0,-0.02], 
             handlelength=0.3, labelspacing=0.3, borderaxespad=0.2)
    
    for i, cond in enumerate(cond_order):
        df1 = mwu_growth_rec.loc[(mwu_growth_rec['cross']==cross) & (mwu_growth_rec['cond']==cond)].set_index(['p1','p2'])
        
        offset_y = y_max[cond] + 2
        for (p1, x1), (p2, x2) in itertools.combinations(zip(order, range(len(order))), 2):
            p = plot_pval_symbol(df1.loc[(p1, p2), 'pval_corr'])
            if p != None:
                X = np.array([x1, x2])
                X = (X-np.mean(X))*0.5+i
                ax.plot(X, np.repeat(offset_y, 2), lw=1, color='k', clip_on=False)
                ax.text(np.mean(X), offset_y, p, ha='center', size=10)
                offset_y += 3
                
ax = axes[2,3]
ax.axis('off')
    
sns.despine()
#plt.savefig('/home/mathieu/mhenault_landrylab/Publications/mito_ma/resubmission2_GRes/fig/fitness_mt_hap.svg', dpi=300)
#plt.show()
plt.close()

# %% [markdown]
## Fig S36

# %% [code]
fig, ax = plt.subplots(figsize=[6,4], gridspec_kw=dict(right=0.75, left=0.1, top=0.96, bottom=0.14))
dat = mwu_growth_rec.copy()
dat['condition'] = dat['cond'].apply(lambda x: cond_alias[x])
sns.scatterplot(x='diff_abs', y='log_pval_corr', hue='cross', hue_order=cross_order, 
                palette=cross_color, style='condition', data=dat, ax=ax)
ax.set_ylim(-0.03, 0.7)
ax.set_xlim(-0.03, 1.5)
ax.legend(loc=2, bbox_to_anchor=(1,1), ncol=1, frameon=False)
ax.margins(0.1)

ax.set_xlabel('median growth difference')
ax.set_ylabel('-log$_{10}$ FDR-corrected p-value')
sns.despine(ax=ax)

ax = fig.add_axes([0.2, 0.6, 0.25 ,0.3])
sns.scatterplot(x='diff_abs', y='log_pval_corr', hue='cross', hue_order=cross_order, 
                palette=cross_color, style='condition', data=dat, ax=ax, legend=False)
ax.margins(0.1)
ax.set_xlabel('')
ax.set_ylabel('')

#plt.savefig('/home/mathieu/mhenault_landrylab/Publications/mito_ma/resubmission2_GRes/fig/growth_diff_effect_size.svg', dpi=300)
#plt.show()
plt.close()

# %% [markdown]
## Table S2

# %% [code]
ma_strains_export = ma_strains.loc[ma_strains['mt_haplotype']!='none'].copy().reset_index(drop=True)
cdrr_idx = combined_drr.set_index(['strain','passage']).sort_index()

for (s,p,clone), df in ma_strains_export.groupby(['strain','passage','mt_haplotype']):
    if (s,p) in cdrr_idx.index:

        ma_strains_export.loc[df.index, 'respiration'] = cdrr_idx.loc[(s,p), 'respiration'].iloc[0]
        #if clone != 'clone':
        ma_strains_export.loc[df.index, 'recomb'] = cdrr_idx.loc[(s,p), 'recomb'].iloc[0]
        ma_strains_export.loc[df.index, 'deletion'] = cdrr_idx.loc[(s,p), 'deletion'].iloc[0]
        ma_strains_export.loc[df.index, 'mito_inst'] = cdrr_idx.loc[(s,p), 'mito_inst'].iloc[0]
        ma_strains_export.loc[df.index, 'nuc_inst'] = cdrr_idx.loc[(s,p), 'nuc_inst'].iloc[0]

ma_strains_export['cross_order'] = ma_strains_export['cross'].apply(lambda x: cross_order[x])
ma_strains_export['strain_num'] = ma_strains_export['strain'].apply(lambda x: int(x[1:]))

for p, df in ma_strains_export.groupby('passage'):
    if p == 1:
        ma_strains_export.loc[df.index, 'timepoint'] = 'initial'
    elif p == 35:
        ma_strains_export.loc[df.index, 'timepoint'] = 'final'

ma_strains_export = ma_strains_export.sort_values(by=['cross_order','strain_num','passage']).replace({True:'True', False:'False'})

# %% [code]
#export for S2
#ma_strains_export[['cross','strain','passage','timepoint','filename','mt_haplotype','recomb','respiration','deletion','mito_inst','nuc_inst']]\
#.to_csv('/home/mathieu/mhenault_landrylab/Publications/mito_ma/resubmission2_GRes/Supplemental_Table_S2.csv')

# %% [markdown]
# Mobility of introns in COB and COX1
## Identify introns to check

# %% [code]
agfi = artificial_genome_feat.set_index('Name')

introns_to_check = []
for cross in cross_order:

    p1, p2 = [p.split('.')[-1] for p in parents_dict[cross]]
    gff = GFF.loc[GFF['strain'].isin([p1,p2])]
    vcf_melt = VCF_MELT[cross]
    markers = MARKERS[cross]
    vcf_melt = vcf_melt.loc[(vcf_melt['marker']==True) &
                            (vcf_melt['vma_uid'].isin(markers))]
    marker_pos = vcf_melt.groupby('vma_uid').apply(lambda x: x['POS'].iloc[0])

    for intron, df in gff.loc[(gff['annot_type']=='intron') & 
                         (gff['Name']!='ai2')].groupby('Name'):

        absent = np.nan
        for p in (p1, p2):
            if p not in df['strain'].values:
                absent = p
            
        if not pd.isna(absent):
            check_mb = [False, False]
            bp1, bp2 = agfi.loc[intron, [3,4]].astype(int)

            #define adjacent markers
            mb1 = marker_pos.loc[marker_pos<bp1]
            mb2 = marker_pos.loc[marker_pos>bp2]
            if mb1.shape[0] > 0 and mb2.shape[0] > 0:
                mb1 = (mb1-bp1).sort_values().index[-1]
                mb2 = (mb2-bp2).sort_values().index[0]

                for bi, mb in enumerate([mb1, mb2]):
                    annot_mb = gft[marker_pos[mb]]

                    at, name = artificial_genome_feat.loc[[i[2] for i in list(annot_mb)]].sort_values(by='annot_plot_order').iloc[-1][['annot_type', 'Name']]
                    #if at in ['exon','rna_exon','gene','rna', 'other', 'gc_cluster']:
                    if at not in ['intron', 'orf']:
                        check_mb[bi] = True
                    else:
                        if set(gff.loc[gff['Name']==name, 'strain']) == set([p1, p2]):
                            check_mb[bi] = True

                introns_to_check.append([cross, intron, bp1, bp2, mb1, mb2, absent, all(check_mb)])
introns_to_check = pd.DataFrame(introns_to_check, columns=['cross', 'intron', 'bp1', 'bp2', 'mb1', 'mb2', 'absent', 'check'])

# %% [markdown]
## Extract coverage depth profiles around intron-exon junctions

# %% [code]
wdw = 50
intron_profiles = []
for cross, df in introns_to_check.loc[introns_to_check['check']==True].groupby('cross'):
    
    p1, p2 = parents_dict[cross]
    P1, P2 = [p.split('.')[-1] for p in (p1, p2)]
    
    #gff = GFF.loc[GFF['strain'].isin([P1,P2])]
    vcf_melt = VCF_MELT[cross]
    vcf_melt = vcf_melt.loc[vcf_melt['marker']].set_index(['filename','vma_uid'])
    
    S = ma_strains.loc[(ma_strains['cross']==cross) & (ma_strains['mthap_filter']), 'filename']
    
    for i in df.index:
        intron, bp1, bp2, mb1, mb2, absent = introns_to_check.loc[i, ['intron', 'bp1', 'bp2', 'mb1', 'mb2', 'absent']].values

        for (bp, mb) in [(bp1, mb1), (bp2, mb2)]:
            
            window = range(bp-wdw, bp+wdw)
            parental_profiles = {}
            pp_check = True
            for p in parents_dict[cross]:
                depth = DEPTH[p].set_index('pos')
                pprofile = depth.loc[window, 'depth']

                pprofile = pprofile/np.quantile(pprofile, 0.8)
                if pprofile.isna().sum() != 0:
                    pp_check = False
                parental_profiles[p] = pprofile
            
            if pp_check == True:
            
                for s in S:

                    # get parent of adjacent marker
                    pmb = np.nan
                    if (s,mb) in vcf_melt.index:
                        pmb = vcf_melt.loc[(s,mb), 'parent']

                    #analyze profiles
                    depth = DEPTH[s]
                    profile = depth.loc[window, 'depth']
                    #profile = profile/profile.mean()
                    profile = profile/np.quantile(profile, 0.8)
                    if profile.isna().sum() == 0:
                        profile_r = {}
                        profile_d = {}
                        for p, pprofile in parental_profiles.items():
                            r, pval = stats.pearsonr(profile, pprofile)
                            profile_r[p] = r
                            d = euclidean(profile, pprofile)
                            profile_d[p] = d

                        intron_profiles.append([cross, intron, s, bp, mb, pmb, profile_r[p1], profile_r[p2], profile_d[p1], profile_d[p2]])

intron_profiles = pd.DataFrame(intron_profiles, columns=['cross', 'intron', 'filename', 'bp', 'mb', 'pmb', 'pearsonr_p1', 'pearsonr_p2', 'euclid_p1', 'euclid_p2'])

# %% [code]
#make calls based on pearson correlations and euclidean distance
intron_profiles['call'] = np.nan
for cross, df in intron_profiles.groupby('cross'):
    p1, p2 = parents_dict[cross]
    df.loc[(df['pearsonr_p1']>df['pearsonr_p2']) & (df['euclid_p1']<df['euclid_p2']), 'call'] = p1
    df.loc[(df['pearsonr_p2']>df['pearsonr_p1']) & (df['euclid_p2']<df['euclid_p1']), 'call'] = p2
    intron_profiles.loc[df.index, 'call'] = df['call']

# %% [code]
#classify introns which have all information
intron_profiles['both_sides'] = False
for (intron, s), df in intron_profiles.loc[(~intron_profiles['call'].isna()) &
                                           (~intron_profiles['pmb'].isna())].groupby(['intron','filename']):
    if df.shape[0] == 2:
        intron_profiles.loc[df.index, 'both_sides'] = True

# %% [markdown]
## Call introns based on junction calls

# %% [code]
intron_calls = []

for (cross, intron, s), df in intron_profiles.loc[intron_profiles['both_sides']].sort_values(by='bp').groupby(['cross','intron','filename']):
    pmb1, pmb2 = df['pmb']
    call1, call2 = df['call']
    bp1, bp2 = df['bp']
    absent = introns_to_check.set_index(['cross','intron']).loc[(cross,intron), 'absent']
    
    rec_bp = np.nan
    rec_bp_side = np.nan
    rec_pmb = np.nan
    
    if call1 == call2:
        if pmb1 == pmb2:
            if pmb1 == call1:
                c = 'wt'
            if pmb1 != call1:
                if call1.split('.')[-1] == absent:
                    c = 'excision'
                elif call1.split('.')[-1] != absent:
                    c = 'mobility'
        else:
            if call1.split('.')[-1] == absent:
                c = 'rec'
            elif call1.split('.')[-1] != absent:
                c = 'rec_intron'
                if call1 == pmb1:
                    rec_bp = bp2
                    rec_pmb = pmb2
                    rec_bp_side = 'right'
                elif call2 == pmb2:
                    rec_bp = bp1
                    rec_pmb = pmb1
                    rec_bp_side = 'left'

    else:
        c = 'call_inconsistent'
    
    intron_calls.append([cross, intron, s, pmb1, pmb2, call1, call2, c, rec_bp, rec_pmb, rec_bp_side])
    
intron_calls = pd.DataFrame(intron_calls, columns=['cross', 'intron', 'filename', 'pmb1', 'pmb2', 'call1', 'call2', 'class', 'rec_bp', 'rec_pmb', 'rec_bp_side'])

# %% [code]
# intron polymorphisms consistent with rec tracts, after visual examination
intron_call_rec_consistent = [['HI.4803.001.N706---N504.I11_P35', 'bi2'],
                              ['HI.4803.001.N706---N504.I11_P35', 'ai1'],
                              ['HI.4803.001.N709---N503.I47_P35', 'ai1'],
                              ['HI.4803.001.N711---N506.I33_P35', 'bi2'],
                              ['HI.4803.002.N701---N504.I38_P35', 'ai1'],
                              ['HI.4803.002.N707---N505.I23_P35', 'ai1'],
                              ['HI.4803.002.N707---N517.I40_P35', 'ai1'],
                              ['HI.4803.002.N702---N517.H46_P35', 'ai1'],
                              ['HI.4803.002.N706---N507.A87_P35', 'bi3'],
                              ['HI.4803.002.N711---N505.A3_P35', 'bi3'],
                              ['HI.4803.002.N703---N504.A70_P35', 'bi3'],
                              ['HI.4802.003.N712---N505.C25_P35', 'bi2'],
                              ['HI.4803.001.N705---N517.C84_P35', 'omega'],
                              ['HI.4803.001.N711---N507.C3_P35', 'ai1'],
                              ['HI.4803.003.N706---N506.F56_P35', 'bi4']]

# %% [markdown]
## Fig S18

# %% [code]
fig, axes = plt.subplots(ncols=2, nrows=2, figsize=[7,5])

for ((cross, intron), df), ax_idx in zip(intron_calls.loc[intron_calls['class']=='mobility'].groupby(['cross','intron']),
                                        itertools.product(range(2), range(2))):
    df = df.set_index('filename', drop=False)
    itc = introns_to_check.loc[(introns_to_check['intron']==intron) & (introns_to_check['cross']==cross)].iloc[0]
    bp1, bp2 = itc[['bp1', 'bp2']].values
    
    S = STRAIN_ORDER_MARKERS[cross] + list(parents_dict[cross])
    
    ax = axes[ax_idx]
    
    for Window, bp in zip([np.arange(-100,0), np.arange(0,100)], [bp1, bp2]):
        window = range(bp-50, bp+50)
        
        xticklabels = np.repeat([bp1, bp2], 3) + np.array([-30, 0, 30, -30, 0, 30])
        Xticks = np.array([-80, -50, -20, 20, 50, 80])
        
        
        for s in S:
            if s in parents_dict[cross]:
                c = parents_color[s.split('.')[-1]]
                lw = 2
                z = 2
                alpha = 1
            elif s in df['filename'].values:
                c = 'k'
                lw = 2
                z = 1
                alpha = 1
                
            else:
                c = 'k'
                lw = 0.5
                z = 0
                alpha = 0.5

            depth = DEPTH[s].set_index('pos')
            bp1_profile = depth.loc[window]

            ax.plot(Window, bp1_profile['depth']/np.quantile(bp1_profile['depth'], 0.8), color=c, lw=lw, zorder=z, alpha=alpha)
    
    fa = FancyArrow(0, 1.11, 0, -1.3, fc='white', width=3, head_length=0, head_width=0, lw=0, clip_on=False, zorder=3)
    ax.add_patch(fa)
    
    for x in (-50, 50):
        ax.axvline(x, ls='--', c='k', alpha=0.3, lw=3, zorder=-1)
    
    ax.set_ylim(-0.1, 1.1)
    ax.set_ylabel('rel depth of cov')
    
    ax.set_xticks(Xticks)
    ax.set_xticklabels(xticklabels, rotation=45, ha='right')
    
    ax.set_title(f'{cross} {pres_abs_poly_alias[intron]}', size=14)
    
    #ax_idx += 1
    
plt.tight_layout()
sns.despine()
#plt.savefig('/home/mathieu/mhenault_landrylab/Publications/mito_ma/resubmission2_GRes/fig/mobility_intron_junctions.svg', dpi=300)
#plt.show()
plt.close()

# %% [markdown]
## Fig S19

# %% [code]
fig = plt.figure(figsize=[8,10])
gs = plt.GridSpec(ncols=1, nrows=2, height_ratios=[14,1], hspace=0, left=0.12, top=0.99, right=0.97, bottom=0.05)

xlim = (-500,ref_genome_length+500)

ax = fig.add_subplot(gs[0])
y = 0
for cross in cross_order:

    if cross in intron_calls.loc[(intron_calls['class']=='rec_intron'), 'cross'].values:
        
        markers = MARKERS[cross]
        vcf_melt = VCF_MELT[cross]
        vcf_melt = vcf_melt.loc[(vcf_melt['marker']) & (vcf_melt['higher'])]
        
        ic = intron_calls.loc[(intron_calls['class']=='rec_intron') & (intron_calls['cross']==cross)]
        
        for s, df in ic.groupby('filename'):
            tracts = TRACTS.loc[TRACTS['strain']==s]
            if tracts.shape[0]>0:
                alpha = 0.3
                for i in tracts.index:
                    (start, end, parent) = tracts.loc[i, ['start','end','parent']]

                    c = parents_color[parent.split('.')[-1]]
                    if start < end:
                        fa = FancyArrow(start, y, end-start, 0, width=0.3, head_length=0, head_width=0, color=c, lw=0, alpha=alpha)
                        ax.add_patch(fa)
                    elif start > end:
                        fa1 = FancyArrow(0, y, end, 0, width=0.3, head_length=0, head_width=0, color=c, lw=0, alpha=alpha)
                        fa2 = FancyArrow(start, y, ref_genome_length-start, 0, width=0.3, head_length=0, head_width=0, color=c, lw=0, alpha=alpha)
                        ax.add_patch(fa1)
                        ax.add_patch(fa2)

                vcf_sub = vcf_melt.set_index('filename').loc[s]
                for parent, df1 in vcf_sub.groupby('parent'):
                    c = parents_color[parent.split('.')[-1]]
                    ax.scatter(df1['POS'], np.repeat(y, df1.shape[0]), s=6, color=c)
            
            ax.text(-500, y, f'{ma_strains.loc[s, "mt_haplotype"]} ({ma_strains.loc[s, "cross"]})', ha='right', va='center', size=9)
            
            for i in ic.loc[ic['filename']==s].index:
                intron, bp, pmb, bp_side = ic.loc[i, ['intron', 'rec_bp', 'rec_pmb', 'rec_bp_side']]
                c = parents_color[pmb.split('.')[-1]]
                if bp_side == 'right':
                    dx = 4e3
                elif bp_side == 'left':
                    dx = -4e3
                #ya = y+0.25
                ya = y+0.3
                if (s == 'HI.4803.003.N706---N506.F56_P35' and intron == 'bi5') or\
                (s == 'NS.1250.002.N706---N507.D28_P35' and intron == 'bi1beta') or\
                (s == 'HI.4803.001.N701---N506.D73_P35' and intron == 'bi1beta'):
                    #ya += 0.25
                    ya += 0.3
                fc = 'white'
                c = parents_color[pmb.split('.')[-1]]
                if [s, intron] in intron_call_rec_consistent:
                    fc = parents_color[pmb.split('.')[-1]]
                    c = 'white'
                fa = FancyArrow(bp, ya, dx, 0, width=0.15, head_length=500, head_width=0.3, fc=fc, lw=0.5, ec=c)
                ax.add_patch(fa)
                ax.text(bp, ya, pres_abs_poly_alias[intron], ha=bp_side, va='center', size=7)
            
            y += 1

ax.set_ylim(-1,y)
ax.set_xlim(xlim)
ax.axis('off')
            
# plot GFF entries
ax = fig.add_subplot(gs[1])

for i in ['left','right','top']:
    ax.spines[i].set_visible(False)
ax.arrow(0, 1, ref_genome_length, 0, color='0.9', lw=0, width=1, head_length=0, head_width=0, zorder=-1)

legend_elms = []
for at in annot_type_order:
    if at != 'gc_cluster':
        l = annot_type_alias[at]
        fc = annot_type_color[at]
        lw = 0
        if fc == 'white':
            lw = 1
        legend_elms.append(Line2D([0], [0], color='w', marker='s', ms=9, mfc=fc, mec='black', mew=lw, label=l))
ax.legend(handles=legend_elms, ncol=6, loc=9, bbox_to_anchor=(0.5, 1.1), frameon=False,
         borderpad=0.0, labelspacing=0.2, handlelength=0.3, columnspacing=1.5)

for i in artificial_genome_feat.loc[~artificial_genome_feat['annot_type'].isin(['gc_cluster','other'])].\
sort_values(by='annot_plot_order', ascending=False).index:
    start, end, color, w, z = artificial_genome_feat.loc[i, [3,4,'annot_color','annot_width','annot_plot_order']]
    ax.arrow(start, 1, end-start, 0, color=color, lw=0, width=w, head_length=0, head_width=0, zorder=z)
ax.set_xlim(xlim)
ax.set_ylim(0.45, 3)
ax.set_xticks(np.arange(0, 81e3, 1e4))
ax.set_xticklabels(range(0,90,10))
ax.set_xlabel('kb')
ax.set_yticks([])


#plt.savefig('/home/mathieu/mhenault_landrylab/Publications/mito_ma/resubmission2_GRes/fig/rec_intron.svg', dpi=300)
#plt.show()
plt.close()

# %% [markdown]
# de novo mutations
## Identify candidate de novo variants

# %% [code]
PRIV_CAND = {}
min_dp = 5
min_ratio = 0.8

lines_fn_to_include = {cross:ma_strains.loc[(ma_strains['cross']==cross) & (ma_strains['identity_filter']), 'filename'].values for cross in cross_order}

for cross, vcf_melt in VCF_MELT.items():
    private_candidates = []
    for (v, gt), df in vcf_melt.loc[(vcf_melt['filename'].isin(lines_fn_to_include[cross])) &
                                   (vcf_melt['dp']>=min_dp) & 
                                   (vcf_melt['ratio']>=min_ratio)].groupby(['var_uid','gt']):
        if len(set(df['line'].values)) == 1:
            fn = set(df['filename'])
            private_candidates.extend(list(df.index))
    private_candidates = vcf_melt.loc[private_candidates].copy().reset_index(drop=True)
    
    # check if the allele is in any of the parents
    print(f'{cross} checking parental variants...')
    vcf_melt_merge = []
    for cross2, vcf_melt2 in VCF_MELT.items():
        #if cross2 != cross:
        vcf_melt2 = VCF_MELT[cross2]
        vcf_melt2 = vcf_melt2.loc[(vcf_melt2['filename'].isin(parents_dict[cross2])) &
                                  (~vcf_melt2['gt'].isna()) &
                                  (vcf_melt2['dp']>=min_dp) &
                                  (vcf_melt2['ratio']>=min_ratio)]
        vcf_melt_merge.append(vcf_melt2)
    vcf_melt_merge = pd.concat(vcf_melt_merge)
    
    private_candidates['parent_filter'] = True
    for (v, s, gt, POS, REF, ALT), df in private_candidates.groupby(['var_uid','line','gt','POS','REF','ALT']):
        sub = vcf_melt_merge.loc[(vcf_melt_merge['POS']==POS) &
                                 (vcf_melt_merge['REF']==REF) &
                                 (vcf_melt_merge['ALT']==ALT) &
                                 (vcf_melt_merge['gt']==gt)]
        if sub.shape[0] > 0:
            private_candidates.loc[df.index, 'parent_filter'] = False
    
    # check if the non-parent allele is not in any other line
    print(f'{cross} checking lines variants...')
    vcf_melt_merge = []
    for cross2, vcf_melt2 in VCF_MELT.items():
        if cross2 != cross:
            vcf_melt2 = VCF_MELT[cross2]
            vcf_melt2 = vcf_melt2.loc[(vcf_melt2['filename'].isin(lines_fn_to_include[cross2])) & 
                                      (~vcf_melt2['gt'].isna()) & 
                                      (vcf_melt2['dp']>=min_dp) &
                                      (vcf_melt2['ratio']>=min_ratio)]
            vcf_melt_merge.append(vcf_melt2)
    vcf_melt_merge = pd.concat(vcf_melt_merge)
    
    private_candidates['lines_filter'] = private_candidates['parent_filter']
    for (v, s, gt, POS, REF, ALT), df in private_candidates.loc[private_candidates['lines_filter']].groupby(['var_uid','line','gt','POS','REF','ALT']):
        sub = vcf_melt_merge.loc[(vcf_melt_merge['POS']==POS) &
                                 (vcf_melt_merge['REF']==REF) &
                                 (vcf_melt_merge['ALT']==ALT) &
                                 (vcf_melt_merge['gt']==gt)]
        if sub.shape[0] > 0:
            private_candidates.loc[df.index, 'lines_filter'] = False
    
    PRIV_CAND[cross] = private_candidates

# %% [code]
for cross, priv_cand in PRIV_CAND.items():
    priv_cand['cross'] = cross
    PRIV_CAND[cross] = priv_cand

PRIV_CAND = pd.concat(PRIV_CAND.values()).reset_index(drop=True)

# %% [code]
# get proportion of lines with defined genotypes for each variant

for (cross, v), df in PRIV_CAND.groupby(['cross','var_uid']):
    vcf_melt = VCF_MELT[cross]
    vcf_melt = vcf_melt.loc[(vcf_melt['filename'].isin(lines_fn_to_include[cross])) &
                            (vcf_melt['var_uid']==v)]

    PRIV_CAND.loc[df.index, 'missing_ratio'] = vcf_melt['gt'].isna().sum()/vcf_melt.shape[0]

# %% [code]
PRIV_CAND['snv'] = (PRIV_CAND['REF'].apply(lambda x: len(x)==1)) & (PRIV_CAND['ALT'].apply(lambda x: len(x)==1))
PRIV_CAND['var_len'] = PRIV_CAND.apply(lambda x: np.abs(len(x['REF'])-len(x['ALT'])), axis=1)

# %% [code]
#filter out variants with more than one alt
PRIV_CAND['single_alt'] = True
for (cross, pos, ref), c in PRIV_CAND.value_counts(['cross','POS','REF']).items():
    if c > 1:
        PRIV_CAND.loc[(PRIV_CAND['cross']==cross) & (PRIV_CAND['POS']==pos) & (PRIV_CAND['REF']==ref), 'single_alt'] = False

# %% [code]
PRIV_CAND['true_snv'] = ((PRIV_CAND['parent_filter']) &
                         (PRIV_CAND['lines_filter']) &
                         (PRIV_CAND['snv']) &
                         (PRIV_CAND['single_alt']) &
                         (PRIV_CAND['gt']==1))

# %% [code]
# inspect the supporting reads for each variant
for (cross, v), df in PRIV_CAND.loc[PRIV_CAND['true_snv']].groupby(['cross','var_uid']):
    
    fn, POS, REF, ALT, gt = df.iloc[0].loc[['filename','POS','REF','ALT','gt']]
    if gt == 1:
        SEQ = ALT
    elif gt == 0:
        SEQ = REF
    
    count_valid = 0
    
    samfile = pysam.AlignmentFile(f'/mnt/HDD3/mito_ma/bam/{fn}.mt.rmdup.rg.bam', 'rb')
    POS -= 1
    al = len(SEQ)
    supporting_reads = []
    for x in samfile.fetch('mt_art', POS, POS+al):
        ap_dict = {j:i for (i,j) in x.aligned_pairs}
        #only consider reads that entirely span the variant length
        if (POS in ap_dict) and (POS+al in ap_dict):
            if (ap_dict[POS] != None) and (ap_dict[POS+al] != None):
                rl = x.reference_length
                a_seq = x.query_sequence[ap_dict[POS]:ap_dict[POS]+al]
                if a_seq == SEQ and rl < 161 and rl > 141:
                    count_valid += 1
                    
    PRIV_CAND.loc[df.index, 'valid_support_reads'] = count_valid

# %% [code]
#perform final filtering; if any candidate is found elsewhere with high enough relative support, discard
for (cross, fn, POS, REF, ALT), df in PRIV_CAND.loc[PRIV_CAND['true_snv']].groupby(['cross','filename','POS','REF','ALT']):
    other_support = []
    for cross2 in cross_order:
        vcf_melt = VCF_MELT[cross2]
        vcf_melt = vcf_melt.loc[(vcf_melt['filename']!=fn) & 
                                (vcf_melt['POS']==POS) &
                                (vcf_melt['REF']==REF) &
                                (vcf_melt['ALT']==ALT)]
        if vcf_melt.shape[0] > 0:
            other_support.extend(vcf_melt['ao'])
    

    if len(other_support) > 0:
        other_support = np.array(other_support)
        other_sum = np.nansum(other_support)
        other_n = other_support.shape[0]
        
    else:
        other_sum = 0
        other_n = 0
    
    PRIV_CAND.loc[df.index, 'other_sum'] = other_sum
    PRIV_CAND.loc[df.index, 'other_n'] = other_n

# %% [code]
display = ['cross', 'line', 'passage', '#CHROM', 'POS', 'REF', 'ALT', 'dp', 'ro', 'ao', 'missing_ratio', 'valid_support_reads','other_sum']
best_cand_snv = PRIV_CAND.loc[PRIV_CAND['true_snv']]
best_cand_snv = best_cand_snv.loc[(best_cand_snv['ao'] >= (best_cand_snv['other_sum']*2)) &
                                  (best_cand_snv['valid_support_reads'] >= 2), display].reset_index(drop=True)
best_cand_snv['annot'] = best_cand_snv['POS'].apply(lambda x: artificial_genome_feat.loc[pos_func_annot[x], 'Name'])
best_cand_snv['annot_type'] = best_cand_snv['POS'].apply(lambda x: artificial_genome_feat.loc[pos_func_annot[x], 'annot_type'])

# %% [markdown]
## Table S5

# %% [code]
#export candidate de novo snv
#best_cand_snv.to_csv('/mnt/HDD3/mito_ma/results/best_cand_snv.csv')

# %% [markdown]
# Nanopore assemblies of lines
## Parse nanopore libraries metadata

# %% [code]
assemblies_lines = pd.read_table('/mnt/HDD3/mito_nanopore/mugsy/assemblies_lines.txt', header=None)
assemblies_lines = pd.concat([assemblies_lines,
                              assemblies_lines.apply(lambda x: pd.Series(x[0].split('.'), index=['strain','k','p','l']), axis=1)], axis=1)
assemblies_lines.index = assemblies_lines[0].values
assemblies_lines['cross'] = ma_strains.groupby('strain')['cross'].apply(lambda x: x.iloc[0]).loc[assemblies_lines['strain']].values

assemblies_lines['cross_order'] = assemblies_lines['cross'].apply(lambda x: cross_order[x])

for s, df in assemblies_lines.groupby('strain'):
    
    assemblies_lines.loc[df.index, 'identity_filter'] = ma_strains.set_index(['strain','passage']).loc[(s, 35), 'identity_filter'].iloc[0]

# %% [markdown]
## Import assemblies

# %% [code]
CNS_LINES = {}

for s in assemblies_lines.index:
    CNS_LINES[s] = {seq.id: seq for seq in SeqIO.parse(f'/mnt/HDD3/mito_nanopore/assemblies_lines/{s}/consensus.fasta', 'fasta')}

# %% [code]
#import atp6-splitted assemblies
CNS_LINES_ATP6 = {}

for s in assemblies_lines.index:
    path = f'/mnt/HDD3/mito_nanopore/assemblies_lines/{s}/consensus.atp6.fasta'
    if os.path.exists(path):
        CNS_LINES_ATP6[s] = {seq.id: seq for seq in SeqIO.parse(f'/mnt/HDD3/mito_nanopore/assemblies_lines/{s}/consensus.atp6.fasta', 'fasta')}
    else:
        CNS_LINES_ATP6[s] = CNS_LINES[s]

# %% [code]
for s in assemblies_lines.index:
    n_tigs = len(CNS_LINES[s])
    len_tigs = sum([len(tig.seq) for tig in CNS_LINES[s].values()])
    assemblies_lines.loc[s, 'n_tigs'] = n_tigs
    assemblies_lines.loc[s, 'len_tigs'] = len_tigs

# %% [markdown]
## Split contigs at ATP6

# %% [code]
idx = 0
with ProgressBar(max_value=assemblies_lines.shape[0]) as bar:
    for s in assemblies_lines.index:
        path_tab = f'/mnt/HDD3/mito_nanopore/assemblies_lines/{s}/tblastn_atp6.consensus.tab'
        if getsize(path_tab) > 0:
            tblastn = pd.read_csv(path_tab, sep='\t', header=None)
            assembly = {seq.id:seq for seq in SeqIO.parse(f'/mnt/HDD3/mito_nanopore/assemblies_lines/{s}/consensus.fasta', 'fasta')}
            start, end = tblastn[[8,9]].iloc[0].astype(int)
            if start < end:
                b = start-1
            else:
                b = start
            c = tblastn[1].iloc[0]
            c0 = SeqRecord.SeqRecord(assembly[c].seq[:b], id=f'{c}.0', description='')
            c1 = SeqRecord.SeqRecord(assembly[c].seq[b:], id=f'{c}.1', description='')
            del assembly[c]
            assembly[f'{c}.0'] = c0
            assembly[f'{c}.1'] = c1
            #SeqIO.write(assembly.values(), f'/mnt/HDD3/mito_nanopore/assemblies_lines/{s}/consensus.atp6.fasta', 'fasta')
        idx += 1
        bar.update(idx)

# %% [markdown]
## Import contig alignments

# %% [code]
coords_lines = {}
for s in assemblies_lines.index:
    coords = pd.read_csv(f'/mnt/HDD3/mito_nanopore/mummer_lines/{s}.artificial_genome.coords', sep='\t', skiprows=4, header=None)
    coords['assembly'] = s
    coords['strain'] = s.split('.')[0]
    coords_lines[s] = coords

# %% [markdown]
## Extract assemblies with deletions

# %% [code]
line_order_deletion = {}
lod_idx = 0
for cross in cross_order:
    df = assemblies_lines.loc[assemblies_lines['cross']==cross]
    for s, df1 in df.groupby('strain'):
        if (df1['len_tigs'].mean() < 60000):
        
            line_order_deletion[s] = lod_idx
            lod_idx += 1

# %% [markdown]
## Fig S37

# %% [code]
fig = plt.figure(figsize=[8,8])
gs = plt.GridSpec(nrows=2, ncols=1, height_ratios=[4,1], hspace=0.1,
                 left=0.05, right=0.93, top=0.97, bottom=0.06)
xlim = (-500,ref_genome_length+500)

ax = fig.add_subplot(gs[0])
for i in ['right','top']:
    ax.spines[i].set_visible(False)
    
for s, df in assemblies_lines.groupby('strain'):
    if s in line_order_deletion:
        
        y = line_order_deletion[s]
        y_offset = 0
        for sa in df.index:
            c = cross_color[assemblies_lines.loc[sa, 'cross']]
            coords = coords_lines[sa]
            #coords = coords.loc[coords[4] >= 1000]

            for i in coords.index:
                ax.plot(coords.loc[i, [0,1]], np.repeat(y+y_offset, 2), c, lw=1.5)
                
            y_offset += 0.05

ax.set_ylim(-0.25, len(line_order_deletion)-0.25)
ax.set_yticks(np.arange(len(line_order_deletion))+0.25)
ax.set_yticklabels(pd.Series(line_order_deletion).sort_values().index)
ax.set_xlim(xlim)
ax.set_xticks(np.arange(0, 81e3, 1e4))
ax.set_xticklabels(range(0,90,10))

ax.xaxis.set_minor_locator(MultipleLocator(1000))

ax.grid(axis='x', which='minor', lw=0.5)
ax.grid(axis='x', which='major', lw=1.5)

for cross, dy in zip(['CC1', 'BC2', 'BA1', 'BA2', 'BSc1', 'BSc2'],
                     [(0,0), (1,1), (2,3), (4,6), (7,8), (9,14)]):
    
    y = np.mean(dy)+0.25
    ax.arrow(8.35e4, y, 5e3, 0, color=cross_color[cross], lw=0, width=(dy[1]-dy[0])+0.9,
             head_length=0, head_width=0, clip_on=False)
    ax.text(8.6e4, y, cross, color='white', ha='center', va='center', size=10)

# plot GFF entries

ax = fig.add_subplot(gs[1])
xlim = (-500, ref_genome_length+500)
ax.arrow(0, 1, ref_genome_length, 0, color='0.9', width=1, head_length=0, head_width=0, zorder=-1)

for i in artificial_genome_feat.loc[~artificial_genome_feat['annot_type'].isin(['gc_cluster','other'])].sort_values(by='annot_plot_order', ascending=False).index:
    start, end, color, w, z = artificial_genome_feat.loc[i, [3,4,'annot_color','annot_width','annot_plot_order']]
    ax.arrow(start, 1, end-start, 0, color=color, width=w, head_length=0, head_width=0, zorder=z)

#create linked labels
pap_sub = [a for a in pres_abs_poly if a in artificial_genome_feat['Name'].values]
for x, a in zip(np.linspace(-4e3, ref_genome_length+6e3, len(pap_sub)),
                pap_sub):

    fc = at_color_hm[at_dict[a]]
    ec = 'white'
    tc = 'white'
    if fc == 'white':
        tc = 'black'
        ec = 'black'
    if fc == 'limegreen':
        tc = 'black'

    t = ax.text(x, 6, pres_abs_poly_alias[a], rotation=90, ha='center', va='top', color=tc, size=7.5,
                 bbox=dict(ec=ec, fc=fc, lw=0.5, boxstyle='square,pad=0.12'), zorder=1)
    pos = artificial_genome_feat.loc[artificial_genome_feat['Name']==a].iloc[0,3]
    ax.plot([pos, x, x], [1.5, 3, 6], lw=0.5, color='black', zorder=-1, clip_on=False)

ax.set_xticks(np.arange(0,83e3,10e3))
ax.set_xticklabels(np.arange(0,83,10))
ax.set_xlabel('kb')
ax.xaxis.set_minor_locator(MultipleLocator(1000))
ax.set_ylim(0.5, 6)
ax.set_yticks([])
ax.set_xlim(xlim)
for i in ['left','right','top']:
    ax.spines[i].set_visible(False)

#plt.savefig(f'/home/mathieu/mhenault_landrylab/Publications/mito_ma/resubmission2_GRes/fig/deletion_lr.svg', dpi=300)
#plt.show()
plt.close()

# %% [markdown]
# petite frequency

# %% [code]
petite_count = pd.read_excel('/home/mathieu/mhenault_landrylab/Experiments/screen_glycerol/petite_test/2022_07_01/counts.xlsx', sheet_name='Sheet1')
petite_count = pd.melt(petite_count.loc[~petite_count['large'].isna()], id_vars=['strain', 'image', 'colony', 'medium'], var_name='size', value_name='count')
petite_hap = petite_count.pivot_table(index='strain', columns='size', values='count', aggfunc=np.mean)
petite_hap['freq_petite'] = (petite_hap['small']/(petite_hap['small']+petite_hap['large']))*100

# %% [code]
#extract frequencies of non-respiring lines at all timepoints
petite_cross = GDR.loc[(GDR['identity_filter']!=False)].value_counts(['Cross','resp']).rename('count').reset_index()\
.pivot_table(index='Cross', columns='resp', values='count', aggfunc=lambda x: x)
petite_cross['freq_resp_loss'] = petite_cross[False]/(petite_cross[True] + petite_cross[False])*100

for cross, parents in parents_dict.items():
    parents = [p.split('.')[-1] for p in parents]
    mean_petite = petite_hap.loc[parents, 'freq_petite'].mean()
    petite_cross.loc[cross, 'freq_petite'] = mean_petite

# %% [code]
petite_stock = pd.read_excel('/home/mathieu/mhenault_landrylab/Experiments/screen_glycerol/petite_test/2022_07_10/counts.xlsx',
                             sheet_name='Sheet1').set_index('strain')
petite_stock['freq_petite'] = (petite_stock['small']/(petite_stock['small']+petite_stock['large']))*100

# %% [code]
inh_petite = gdrec.loc[(gdrec['resp']==False) & (gdrec['passage']==1)].value_counts(['cross','parent'], sort=False).rename('count').reset_index()
inh_petite['total'] = gdrec.loc[(~gdrec['resp'].isna()) & (gdrec['passage']==1)].value_counts('cross', sort=False).rename('count')\
.loc[inh_petite['cross']].values
inh_petite['frac_inh_petite'] = inh_petite['count']/inh_petite['total']*100
inh_petite = inh_petite.loc[inh_petite['parent']!='rec']
inh_petite['freq_petite'] = petite_stock.loc[inh_petite['parent'].values, 'freq_petite'].values

# %% [markdown]
## Fig S33

# %% [code]
fig = plt.figure(figsize=[8,7])
gs = plt.GridSpec(nrows=2, ncols=2, wspace=0.5, hspace=0.4,
                 bottom=0.12, left=0.1, right=0.85, top=0.96)

# Spontaneous petite frequency
ax = fig.add_subplot(gs[0,0])
fig.text(0.02, 0.95, 'A', weight='bold', size=24)
ax.bar(range(13), petite_hap.loc[parents_order.index, 'freq_petite'], color=[parents_color[p] for p in parents_order.index])
ax.set_xticks(range(13))
ax.set_xticklabels(parents_order.index, size=8, ha='right', rotation=45)
ax.set_ylabel('petite rate (%)')

ax = fig.add_subplot(gs[0,1])
fig.text(0.45, 0.95, 'B', weight='bold', size=24)
ax.scatter(petite_cross['freq_petite'], petite_cross['freq_resp_loss'], color=[cross_color[cross] for cross in petite_cross.index])
lr = stats.linregress(petite_cross['freq_petite'], petite_cross['freq_resp_loss'])
X = [0.4, 2.7]
ax.plot(X, [x*lr.slope+lr.intercept for x in X], c='k', zorder=0)
ax.text(0.1, 0.5, f'r={lr.rvalue:.2f}\np={plot_pval_text(lr.pvalue)}', transform=ax.transAxes)
ax.set_xlabel('mean parental petite rate (%)')
ax.set_ylabel('% lines with respiration loss')

# fraction of petite cells in stock cultures

ax = fig.add_subplot(gs[1,0])
fig.text(0.02, 0.45, 'C', weight='bold', size=24)
ax.bar(range(13), petite_stock.loc[parents_order.index, 'freq_petite'], color=[parents_color[p] for p in parents_order.index])
ax.set_xticks(range(13))
ax.set_xticklabels(parents_order.index, size=8, ha='right', rotation=45)
ax.set_ylabel('petite fraction in stock (%)')

ax = fig.add_subplot(gs[1,1])
fig.text(0.42, 0.45, 'D', weight='bold', size=24)
ax.scatter(inh_petite['freq_petite'], inh_petite['frac_inh_petite'], color=[cross_color[c] for c in inh_petite['cross'].values])
lr = stats.linregress(inh_petite['freq_petite'], inh_petite['frac_inh_petite'])
X = [0.8, 9]
ax.plot(X, [x*lr.slope+lr.intercept for x in X], c='k', zorder=0)
ax.text(0.6, 0.5, f'r={lr.rvalue:.2f}\np={plot_pval_text(lr.pvalue)}', transform=ax.transAxes)
ax.set_xlabel('petite fraction in stock (%)')
ax.set_ylabel('% non-recombinant\ndeleted mtDNA haplotypes')

ax = fig.add_subplot(gs[:, 1])
ax.axis('off')
legend_elms = [Line2D([0], [0], color='white', marker='o', ms=8, mfc=cross_color[cross], label=cross) for cross in cross_order]
ax.legend(handles=legend_elms, ncol=1, loc=3, bbox_to_anchor=[1.02,0.3])

sns.despine()
#plt.savefig('/home/mathieu/mhenault_landrylab/Publications/mito_ma/resubmission2_GRes/fig/petite_freq.svg', dpi=300)
#plt.show()
plt.close()
```
