## Supplemental Code for "Hybridization drives mitochondrial DNA degeneration and metabolic shift in a species with biparental mitochondrial inheritance": nanopore_assemblies.ipynb

```
# %% [code]
import pandas as pd
import numpy as np
from Bio import SeqIO
from Bio import SeqRecord
from Bio import Seq
from Bio.SeqUtils import GC
import os
import re
import copy
import itertools
from collections import Counter
import gzip
from matplotlib import pyplot as plt
from matplotlib.colors import rgb2hex
from matplotlib import cm
import seaborn as sns
import Levenshtein as lvs
import pickle as pkl
sns.set(style='ticks', font='DejaVu Sans')

# %% [code]
def is_non_zero_file(fpath):  
    return os.path.isfile(fpath) and os.path.getsize(fpath) > 250

# %% [markdown]
# Parse metadata

# %% [code]
nano_strains = pd.read_csv('/mnt/HDD3/mito_nanopore/script/nano_strains.csv', header=0, index_col=0)

# %% [code]
strain_order = nano_strains.loc[nano_strains['cross']=='P'].sort_values(by=['ref_name','strain'])['strain']

# %% [code]
#export library filenames for ncbi
filenames_list = []
for s in nano_strains.loc[nano_strains['cross']=='P'].index:
    paths = nano_strains.loc[s, 'path']
    d = re.search('(.+)(/2019\*)(.+)(\*\.fastq.gz)', paths)#.group(1)
    if d:
        parent_d = d.group(1)
        child_d = [i for i in os.listdir(parent_d) if i[:4]=='2019'][0]
        d = f'{parent_d}/{child_d}{d.group(3)}'
    else:
        d = re.search('(.+)(\*\.fastq.gz)', paths).group(1)
    files = [f for f in os.listdir(d) if f.split('.')[-1]=='gz']
    filenames_list.append('\t'.join([s]+files))
#with open('/home/mathieu/mhenault_landrylab/Experiments/MA_2019/ncbi_submissions/filenames_parents.tsv', 'w') as handle:
#    handle.write('\n'.join(filenames_list))

# %% [code]
parents_color = {'MSH-604':'red',
                   'UWOPS-91-202':'red',
                   'LL2012_021':'darkred',
                   'LL2012_028':'darkred',
                   'LL2011_004':'dodgerblue',
                   'LL2011_009':'dodgerblue',
                   'MSH-587-1':'midnightblue',
                   'LL2011_012':'midnightblue',
                   'LL2011_001':'midnightblue',
                   'YPS644':'limegreen',
                   'YPS744':'limegreen',
                   'LL2013_040':'dimgrey',
                   'LL2013_054':'dimgrey'}

# %% [markdown]
# MtDNA assembly

# %% [markdown]
## Generate a compound reference genome to bait mitochondrial reads

# %% [code]
# concat mt refs and append them to the ref genomes
mt_ref = ['/home/mathieu/mhenault_landrylab/Sequences/ref_genomes/CBS432_pacbio/CBS432.mt.genome.fa',
          '/home/mathieu/mhenault_landrylab/Sequences/ref_genomes/YPS138_pacbio/YPS138.mt.genome.fa',
          '/home/mathieu/mhenault_landrylab/Sequences/ref_genomes/S288C_pacbio/S288c.mt.genome.fa']
mt_ref_len = {}
MT = []
for path in mt_ref:
    with open(path, 'r') as fi:
        seq = SeqIO.read(fi, 'fasta').seq
        MT.append(str(seq))
    mt_ref_len[path] = len(seq)
        
MT = ('N'*100).join(MT)
MT = np.apply_along_axis(lambda x: ''.join(x), 1, np.array(list(MT)+list('N'*29)).reshape(-1, 60))
MT = '\n'.join(MT)

# %% [code]
# import raw reads baited from mapping to PacBio refs
RL = {}
for s in nano_strains.loc[nano_strains['cross']=='P'].index:
    with gzip.open(f'/mnt/HDD3/mito_nanopore/bait/mito_reads_{s}.fastq.gz', 'rt') as fi:
        RL[s] = [len(i.seq) for i in SeqIO.parse(fi, 'fastq')]

# %% [code]
read_stats = []
for s in RL:
    read_stats.append([s, len(RL[s]), np.median(RL[s])])
read_stats = pd.DataFrame(read_stats, columns=['strain','n','median'])
read_stats.index = read_stats['strain'].values

# %% [markdown]
## Optimize assembly parameters

# %% [code]
opt = []
for (s,k,p,l) in itertools.product(nano_strains.loc[nano_strains['cross']=='P'].index,
                                   [23,21,19,17,15,13,11],
                                   [8,5,2,1,0],
                                   [4096,2048,1024]):
    opt.append([f'{s}.{k}.{p}.{l}', s, k, p, l])
opt = pd.DataFrame(opt, columns=['dir','strain','k','p','l'])
opt['ref_mt'] = nano_strains.loc[opt['strain'], 'ref_mt'].apply(lambda x: x.replace('Users','home')).values
opt.index = opt['dir'].values

with open('/mnt/HDD3/mito_nanopore/wtdbg2/test/cns.txt') as fi:
    cns_list = fi.read().splitlines()
opt['cns'] = False
for d in opt.index:
    if f'{d}.cns.fa' in cns_list:
        # test if file is not empty
        with open(f'/mnt/HDD3/mito_nanopore/wtdbg2/test/{d}/{d}.cns.fa') as fi:
            seq = [i for i in SeqIO.parse(fi, 'fasta')]
            if len(seq) != 0:
                opt.loc[d, 'cns'] = True

#opt.to_csv('/mnt/HDD3/mito_nanopore/mummer/opt_mummer.csv')
opt['coords'] = opt['dir'].apply(lambda x: is_non_zero_file(f'/mnt/HDD3/mito_nanopore/mummer/{x}.coords'))

# %% [code]
plot = False
OPT = []
for i in opt.loc[opt['coords']].index:
    
    # parse assembly
    cns = {}
    d = opt.loc[i, 'dir']
    with open(f'/mnt/HDD3/mito_nanopore/wtdbg2/test/{d}/{d}.cns.fa') as fi:
        for tig in SeqIO.parse(fi, 'fasta'):
            cns[tig.id] = tig
    sizes = [len(tig.seq) for tig in cns.values()]
    assembly_size = sum(sizes)
    n_contigs = len(sizes)
    largest = max(sizes)
    
    if is_non_zero_file(f'/mnt/HDD3/mito_nanopore/mummer/{d}.coords'):
        
        delta = pd.read_csv(f'/mnt/HDD3/mito_nanopore/mummer/{d}.coords', sep='\t', skiprows=range(4), index_col=None, header=None)
        tig_order = delta.groupby(10).apply(lambda x: x[[0,1]].mean(axis=1).mean()).sort_values()
        tig_rank = pd.Series(range(tig_order.shape[0]), index=tig_order.index)
        tig_len = pd.Series(np.cumsum([0]+[len(cns[tig].seq) for tig in tig_order.index[:-1]]), index=tig_order.index)

        ref_len = mt_ref_len[opt.loc[i, 'ref_mt']]
        ref_cov = np.zeros(ref_len)
        for c1,c2 in delta[[0,1]].values:
            ref_cov[c1-1:c2] += 1

        OPT.append(list(opt.loc[i].values)+[delta[4].sum(), delta[5].sum(), delta[6].mean(), assembly_size, ref_len, n_contigs, largest, np.where(ref_cov==1, 1, 0).mean(), np.std(ref_cov)])

        if plot:
            
            delta['cns_start'] = delta[2]+tig_len.loc[delta[10]].values
            delta['cns_end'] = delta[3]+tig_len.loc[delta[10]].values
            
            fig, ax = plt.subplots(figsize=[7,7])

            for i in delta.index:
                ax.plot(delta.loc[i, [0,1]], delta.loc[i, ['cns_start','cns_end']], color='k')
            for i in tig_len.index:
                ax.axhline(tig_len.loc[i], lw=0.5, color='grey')
                ax.text(ax.axis()[1]-5e3, tig_len.loc[i]+1e3, s=i, ha='right')
            ax.set_title(d)
            ax.set_xlabel('YPS138')
            ax.set_ylabel(f'{d} cns')
            ax.margins(0)

            plt.tight_layout()
            #plt.savefig(f'/mnt/HDD3/mito_nanopore/mummer/fig/{d}_cns.png', dpi=300)
            plt.close()

OPT = pd.DataFrame(OPT, columns=['dir','strain','k','p','l','ref_mt','cns','coords','Rcov','Qcov','ident','Qlen', 'Rlen','n_contigs','largest','Rcov_1','Rcov_std']).astype({'k':int,'p':int,'l':int})
OPT['QCov'] = OPT['Qcov']/OPT['Qlen']
OPT['RCov'] = OPT['Rcov']/OPT['Rlen']
OPT['QRmean'] = OPT[['QCov','RCov']].mean(axis=1)

# %% [code]
# compute score that includes deviation from expected coverage and contiguity
OPT['contiguity'] = 1/OPT['largest']
OPT['dev'] = OPT[['QCov','Rcov_1']].apply(lambda x: np.sum(np.abs(x-1)), axis=1)
score_comp = OPT[['dev','contiguity']]
OPT['score'] = ((score_comp-score_comp.mean())/score_comp.std()).sum(axis=1)

# %% [code]
# add read stats
OPT['n'] = read_stats.loc[OPT['strain'], 'n'].values
OPT['median'] = read_stats.loc[OPT['strain'], 'median'].values

# %% [code]
# exploratory variables
OPT['dummy_k.p'] = OPT.apply(lambda x: f'{x["strain"]}.{x["k"]}.{x["p"]}', axis=1)
OPT['dummy_k.l'] = OPT.apply(lambda x: f'{x["strain"]}.{x["k"]}.{x["l"]}', axis=1)
OPT['dummy_l.p'] = OPT.apply(lambda x: f'{x["strain"]}.{x["l"]}.{x["p"]}', axis=1)
for s, df in OPT.sort_values(by='dev', ascending=False).groupby('strain'):
    OPT.loc[df.index, 'rank_dev'] = np.linspace(0,1,df.shape[0])

interactions = []
for interaction_size in [1,2,3]:
    for i in itertools.combinations(['k','p','l'], interaction_size):
        terms = ['median']+list(i)
        name = '*'.join(terms)
        d = OPT[terms]
        interactions.append(name)
        OPT[name] = ((d-d.mean())/d.std()).apply(lambda x: np.prod(x), axis=1)

# %% [code]
best_10 = OPT.sort_values(by='dev').groupby('strain').apply(lambda x: x.iloc[:10])

for f in ['k','p','l']:
    
    for q, ls in zip([0.05, 0.5], [':','-']):
    

        df = OPT.groupby(['strain','n',f])['score'].apply(lambda x: np.quantile(x, q)).reset_index()
        sns.lineplot(x=f, y='score', hue='n', palette='viridis', ls=ls, lw=0.5, legend=False, data=df)

    
    plt.show()
    plt.close()

# %% [code]
best = OPT.sort_values(by='score', ascending=True).groupby('strain').apply(lambda x: x.iloc[0])['dir']
#with open('/Volumes/Seagate/mito_nanopore/wtdbg2/best_assemblies.txt', 'w') as fo:
#    fo.write('\n'.join(best.values))

# %% [markdown]
## Circularize assembies at the start of ATP6

# %% [code]
breakpoint_atp6 = pd.read_csv('/Volumes/Seagate/mito_nanopore/circularization/breakpoint_atp6.csv', header=None, index_col=0)

CIRC = {}
CIRC_dup = {}
for d in best:
    # parse uncircularized contig
    to_circ = {}
    with open(f'/Volumes/Seagate/mito_nanopore/wtdbg2/test/{d}/{d}.cns.fa') as fi:
        for tig in SeqIO.parse(fi, 'fasta'):
            to_circ[tig.id] = tig
    # parse breakpoint at atp6
    s = d.split('.')[0]
    pos, strand = breakpoint_atp6.loc[s, [2,3]]
    # select 
    seq = to_circ['ctg1'].seq
    # circularize
    if strand == '+':
        pos -= 1
    print(s, len(seq[pos:]), len(seq[:pos]))
    Seq = seq[pos:] + seq[:pos]
    #reverse complement if needed
    if strand == '-':
        Seq = Seq.reverse_complement()
    # store
    sr = SeqRecord.SeqRecord(seq=Seq, id=f'mt.{s}', description='')
    
    CIRC[s] = sr

#for s in CIRC:
#    with open(f'/Volumes/Seagate/mito_nanopore/circularization/{s}.circ.fasta', 'w') as fo:
#        SeqIO.write(CIRC[s], fo, 'fasta')

# %% [code]
# compute levenshtein distance between windiws close to the breakpoint
bw = 20
step = 7
wdw = 500

for s in CIRC:

    seq = CIRC[s].seq
    # 2.5 kb on each side of the breakpoint
    pos, strand = breakpoint_atp6.loc[s, [2,3]]
    if strand == '+':
        # convert position of the breakpoint
        pos = len(CIRC[s].seq)-pos
    start, end = pos-wdw, pos+wdw

    distance = []
    bin_seq = {i+0.5*bw:str(seq[i:i+bw]) for i in np.arange(start, end-bw, step)}
    for (i1,b1), (i2,b2) in itertools.combinations(bin_seq.items(), 2):
        #similarity = (b1 == b2).mean()
        d = lvs.distance(b1, b2)
        distance.append([i1, i2, d])

    dat = pd.pivot_table(pd.DataFrame(distance), index=0, columns=1, values=2)
    fig, ax = plt.subplots(figsize=[10,10])
    sns.heatmap(dat, cmap='viridis', ax=ax)
    ax.set_title(f'{s} {pos}')

    ax.axvline(70, color='red', lw=0.5)

    plt.tight_layout()
    #plt.savefig(f'/Volumes/Seagate/mito_nanopore/circularization/distance/levenshtein.{bw}.{step}.{wdw}.{s}.png', dpi=300)
    plt.close()

# %% [code]
# remove tandem duplications at the artificial breakpoint site
remove_dup = pd.read_csv('/Volumes/Seagate/mito_nanopore/circularization/remove_dup.csv', header=None, index_col=0)

CIRC_RD = {}
for s, sr in CIRC.items():
    start, end = remove_dup.loc[s]
    seq1 = sr.seq[:start-1]
    seq2 = sr.seq[end:]
    
    nsr = SeqRecord.SeqRecord(seq=seq1+seq2, id='mt', description='')
    CIRC_RD[s] = nsr
    
    # export mt genomes alone
    #with open(f'/Volumes/Seagate/mito_nanopore/circularization/{s}.circ.rd.fasta', 'w') as fo:
    #    SeqIO.write(CIRC_RD[s], fo, 'fasta') 

# %% [code]
# analysis of coverage of baited reads on refs
DEPTH_BAIT = {}
for s in nano_strains.loc[nano_strains['cross']=='P'].index:
    depth = pd.read_csv(f'/Volumes/Seagate/mito_nanopore/minimap2/{s}.bait.concat_mt.sorted.depth', sep='\t', header=None)
    for ref, df in depth.groupby(0):
        depth.loc[df.index, 'bin'] = pd.cut(df[1], np.arange(0, df[1].max(), 1000))
    depth = depth.groupby([0,'bin'])[2].mean().reset_index()
    depth['pos'] = depth['bin'].apply(lambda x: np.mean([x.left,x.right]))
    DEPTH_BAIT[s] = depth

# %% [code]
for s, df in DEPTH_BAIT.items():
    fig, ax = plt.subplots()
    
    for ref, df1 in df.groupby(0):
        ax.plot(df1['pos'], df1[2])
    ax.set_title(s)
    
    #plt.show()
    plt.close()

# %% [markdown]
# Annotations processing

# %% [code]
# import polished assemblies
CORR = {}

for s in strain_order:
    with open(f'/mnt/HDD3/mito_nanopore/polishing/{s}/{s}.nanopolish.pilon.fasta') as handle:
        CORR[s] = SeqIO.read(handle, 'fasta')

# %% [code]
# import mfannot gff files
GFF = []

for s in nano_strains.loc[nano_strains['cross']=='P', 'strain']:
    
    gff = pd.read_csv(f'/mnt/HDD3/mito_nanopore/mfannot/{s}.mfannot.gff', sep='\t', skiprows=1, header=None)
    fields = gff[8].apply(lambda x: pd.Series(dict([i.split('=') for i in x.split(';')])))
    fields['strain'] = s
    
    GFF.append(pd.concat([gff, fields], axis=1))

GFF = pd.concat(GFF).reset_index(drop=True)

GFF['strain_order'] = pd.Series(range(13), index=strain_order).loc[GFF['strain']].values

GFF['annot_color'] = 'k'
GFF['annot_type'] = 'k'
GFF['annot_width'] = 1
GFF['annot_plot_order'] = 0

for f, df in GFF.groupby('gene'):
    if 'trn' in f:
        GFF.loc[df.index, 'annot_color'] = 'limegreen'
        GFF.loc[df.index, 'annot_type'] = 'tRNA'
        GFF.loc[df.index, 'annot_width'] = 1
        GFF.loc[df.index, 'annot_plot_order'] = 1
    if '-E' in f:
        GFF.loc[df.index, 'annot_color'] = 'blue'
        GFF.loc[df.index, 'annot_type'] = 'exon'
        GFF.loc[df.index, 'annot_width'] = 0.5
        GFF.loc[df.index, 'annot_plot_order'] = 3
    if '-I' in f:
        GFF.loc[df.index, 'annot_color'] = 'white'
        GFF.loc[df.index, 'annot_type'] = 'intron'
        GFF.loc[df.index, 'annot_width'] = 0.5
        GFF.loc[df.index, 'annot_plot_order'] = 4
    if 'orf' in f:
        GFF.loc[df.index, 'annot_color'] = 'red'
        GFF.loc[df.index, 'annot_type'] = 'orf'
        GFF.loc[df.index, 'annot_width'] = 0.3
        GFF.loc[df.index, 'annot_plot_order'] = 5
    if f in ['atp6', 'cob', 'atp9', 'rps3', 'cox3', 'cox2', 'rnpB', 'cox1', 'atp8']:
        GFF.loc[df.index, 'annot_color'] = 'dodgerblue'
        GFF.loc[df.index, 'annot_type'] = 'gene'
        GFF.loc[df.index, 'annot_width'] = 1
        GFF.loc[df.index, 'annot_plot_order'] = 2

# %% [code]
#re-import final gff
GFF = pd.read_csv('/mnt/HDD3/mito_nanopore/mfannot/gff_edit.csv', sep='\t')
GFF.columns = list(range(9)) + list(GFF.columns[9:])

# %% [markdown]
## Curation of MFannot annotations

# %% [code]
# export features by blocks; define anchors as well preserved genes
anchors = ['atp6','cob','atp9','rps3','cox2','cox3','rnpB','rns','cox1','atp8']

# %% [code]
# correct the atp6 initial positions
GFF.loc[GFF['gene']=='atp6', 3] = 1

# %% [code]
# export GFF for manual editing
#GFF.sort_values(by=['annot_type','strain',3,4]).to_csv('/mnt/HDD3/mito_nanopore/mfannot/gff.tsv', sep='\t')

# %% [code]
# import edited GFF
GFF = pd.read_csv('/mnt/HDD3/mito_nanopore/mfannot/gff_edit.csv', sep='\t')
GFF.columns = list(range(9)) + list(GFF.columns)[9:]
GFF = GFF.astype({3:int,4:int,'strain_order':int,'annot_width':float,'annot_plot_order':int})

# %% [code]
#export sequences of main genes
for s, df in GFF.loc[GFF['gene'].isin(anchors)].sort_values(by=3).groupby('strain'):
    
    # for each anchor, take the sequence of the anchor itself and the following block
    for i in range(df.shape[0]):
        a = df.iloc[i, 12]
        ca1, ca2 = df.iloc[i, 3], df.iloc[i, 4]
        if i == df.shape[0]-1:
            ci2 = len(CORR[s].seq)+1
        else:
            ci2 = df.iloc[i+1, 3]
        # add anchor
        
        anchor_id = f'{a}.anchor.{s}.{ca1-1}.{ca2}'
        inter_id = f'{a}.inter.{s}.{ca2-1}.{ci2}'
        
        #with open(f'/mnt/HDD3/mito_nanopore/muscle/fasta/{anchor_id}.fasta', 'w') as handle:
        #    SeqIO.write(SeqRecord.SeqRecord(CORR[s].seq[ca1-1:ca2], id=anchor_id, description=''), handle, 'fasta')
        #with open(f'/mnt/HDD3/mito_nanopore/muscle/fasta/{inter_id}.fasta', 'w') as handle:
        #    SeqIO.write(SeqRecord.SeqRecord(CORR[s].seq[ca2:ci2-1], id=inter_id, description=''), handle, 'fasta')

# %% [code]
# output list of sequence blocks
#with open('/mnt/HDD3/mito_nanopore/muscle/list_blocks.txt', 'w') as handle:
#    handle.write('\n'.join([f'{i}.anchor\n{i}.inter' for i in anchors]))

# %% [code]
# for cob and cox1, extract spliced cds
for a in ['cob', 'cox1']:
    for s, df in GFF.loc[(GFF['annot_type']=='exon') & (GFF['gene'].apply(lambda x: a in x))].sort_values(by=3).groupby('strain'):
        seq = []
        for i in df.index:
            start, end = df.loc[i, [3,4]]
            seq.append(str(CORR[s].seq[start-1:end]))
        seq = Seq.Seq(''.join(seq))
        name = f'{a}.spliced.{s}.{start}.{end}'
        with open(f'/mnt/HDD3/mito_nanopore/muscle/fasta/{name}.fasta', 'w') as handle:
            SeqIO.write(SeqRecord.SeqRecord(seq=seq, id=name, description=''), handle, 'fasta')
#with open('/mnt/HDD3/mito_nanopore/muscle/list_spliced.txt', 'w') as handle:
#    handle.write('\n'.join([f'{i}.spliced' for i in ['cob','cox1']]))

# %% [code]
# extract sequences from ref assemblies
REF_ASSEMBLIES = {}
for s in ['S288C','CBS432','CBS7400']:
    with open(f'/home/mathieu/mhenault_landrylab/Sequences/ref_genomes/mt_introns/{s}.mt.fasta') as handle:
        REF_ASSEMBLIES[s] = SeqIO.read(handle, 'fasta')

# %% [code]
# import gff annotations of reference mtDNAs
REF_GFF = []

for s in REF_ASSEMBLIES:
    
    gff = pd.read_csv(f'/home/mathieu/mhenault_landrylab/Sequences/ref_genomes/mt_introns/{s}.mt.gff3', sep='\t', skiprows=2, header=None)
    fields = gff[8].apply(lambda x: pd.Series(dict([i.split('=') for i in x.split(';')])))
    fields['strain'] = s
    
    REF_GFF.append(pd.concat([gff, fields], axis=1))

REF_GFF = pd.concat(REF_GFF).reset_index(drop=True)

# %% [code]
# output ref intron sequences
ref_introns = []
for i in REF_GFF.loc[REF_GFF[2]=='intron'].index:
    s, start, end, name = REF_GFF.loc[i, ['strain',3,4,'Note']]
    ref_introns.append(SeqRecord.SeqRecord(seq=REF_ASSEMBLIES[s].seq[start-1:end], id=name.replace(' ', '.'), description=''))
#with open('/mnt/HDD3/mito_nanopore/introns/ref_introns.fasta', 'w') as handle:
#    SeqIO.write(ref_introns, handle, 'fasta')
    
# output query intron sequences
query_introns = []
for i in GFF.loc[GFF['annot_type']=='intron'].index:
    s, start, end, name = GFF.loc[i, ['strain',3,4,'gene']]
    query_introns.append(SeqRecord.SeqRecord(seq=CORR[s].seq[start-1:end], id=f'{name}.{s}', description=''))
#with open('/mnt/HDD3/mito_nanopore/introns/query_introns.fasta', 'w') as handle:
#    SeqIO.write(query_introns, handle, 'fasta')

# %% [code]
# import intron blastn results
blastn_introns = pd.read_csv('/mnt/HDD3/mito_nanopore/introns/introns_blastn.tab', sep='\t', header=None)

bi = blastn_introns.sort_values(by=10).groupby(0).apply(lambda x: x.iloc[0]).reset_index(drop=True)
bi = pd.concat([bi, bi[0].apply(lambda x: pd.Series(x.split('.'), index=['intron','strain']))], axis=1)

# %% [code]
# export the sequences of curated intron annotations for MSA
# goal is to check intron boundaries to see if they agree well, and verify the extent of variation between strains 
introns = {}
for i, df in GFF.loc[GFF['annot_type']=='intron'].sort_values(by='strain_order').groupby('Name'):
    introns[i] = []
    for j in df.index:
        start, end, s = df.loc[j, [3,4,'strain']]
        introns[i].append(SeqRecord.SeqRecord(seq=CORR[s].seq[start-1:end], id=f'{i}.{s}', description=''))
#for i in introns:
#    with open(f'/mnt/HDD3/mito_nanopore/muscle/fasta/{i}.fasta', 'w') as handle:
#        SeqIO.write(introns[i], handle, 'fasta')
# output list of introns
#with open('/mnt/HDD3/mito_nanopore/muscle/list_introns.txt', 'w') as handle:
#    handle.write('\n'.join([i for i in introns]))

# %% [code]
#export orf sequences for alignment
orfs = {}
for o, df in GFF.loc[GFF['annot_type']=='orf'].sort_values(by='strain_order').groupby('Name'):
    if 'ai' not in o and 'bi' not in o:
        orfs[o] = []
        for s, df1 in df.groupby('strain'):
            start = df1[3].min()
            end = df1[4].max()
            orfs[o].append(SeqRecord.SeqRecord(seq=CORR[s].seq[start-1:end], id=f'{o}.{s}', description=''))
#for o in orfs:
#    with open(f'/mnt/HDD3/mito_nanopore/muscle/fasta/{o}.fasta', 'w') as handle:
#        SeqIO.write(orfs[o], handle, 'fasta')

# %% [code]
# export all features for search against assemblies
ref_feats = []
for (s, start, end), df in REF_GFF.loc[REF_GFF['gbkey']!='Src'].sort_values(by='gbkey').groupby(['strain',3,4]):

    name = f'{s}.' + '_'.join(np.unique(df.dropna(axis=1).iloc[:, 9:].values)).replace(' ', '.') + f'.idx{df.index[0]}'
    
    ref_feats.append(SeqRecord.SeqRecord(seq=REF_ASSEMBLIES[s].seq[start-1:end], id=name, description=''))
    
#with open('/mnt/HDD3/mito_nanopore/features/ref_features.fasta', 'w') as handle:
#    SeqIO.write(ref_feats, handle, 'fasta')

# %% [markdown]
# Creation of artificial reference genome

# %% [code]
# use the mtDNA with largest feature set as template (LL2012_028)
artificial_genome = copy.copy(CORR['LL2012_028'])
artificial_genome.description = ''

# coordinates of introns to complement in LL2012_028
ai3gamma = 56, 69566
bi5 = 755, 11224
bi4 = 752, 11174
bi3 = 12, 10924
intron_length = []

for i, intron in enumerate([ai3gamma, bi5, bi4, bi3]):
    s, c1, c2 = GFF.loc[intron[0], ['strain',3,4]]
    intron_length.append(c2-c1+1)
    pos = intron[1]
    print(artificial_genome.seq[pos-20:pos])
    artificial_genome.seq = artificial_genome.seq[:pos] + CORR[s].seq[c1-1:c2] + artificial_genome.seq[pos:]
    artificial_genome.id = 'mt_art'
    
# export artificial reference genome
#with open('/mnt/HDD3/mito_nanopore/artificial_genome/artificial_genome.fasta', 'w') as handle:
#    SeqIO.write(artificial_genome, handle, 'fasta')

# %% [markdown]
# Alignment of final mtDNA assemblies to S288c reference

# %% [code]
#import mummer coords files
COORDS_S288c= {}
for s in nano_strains.loc[nano_strains['cross']=='P', 'strain']:
    coords = pd.read_csv(f'/mnt/HDD3/mito_nanopore/mummer_S288c/{s}.coords', sep='\t', skiprows=4, header=None)
    COORDS_S288c[s] = coords

# %% [markdown]
## Fig S1

# %% [code]
fig = plt.figure(figsize=[8,8])
gs = plt.GridSpec(ncols=4, nrows=4)

parents_ax = dict(zip(parents_color, itertools.product(range(4), range(4))))

for s, coords in COORDS_S288c.items():
    
    ax = fig.add_subplot(gs[parents_ax[s]])
    
    for i in coords.index:
        strand, ident = coords.loc[i, [8,6]]
        c = parents_color[s]

        ax.plot(coords.loc[i, [0,1]], coords.loc[i, [2,3]], c=c, lw=3, label=s)
        
    ax.set_xlim([-2e3, 85e3])
    ax.set_xticks(np.arange(0,85e3,2e4))
    ax.set_xticklabels(np.arange(0,85,20), size=9)
    ax.set_xlabel('S288c')
    ax.set_ylim([-2e3, 85e3])
    ax.set_yticks(np.arange(0,85e3,2e4))
    ax.set_yticklabels(np.arange(0,85,20), size=9)
    ax.set_ylabel(f'{s}')
    
    ax.grid()

plt.tight_layout()
#plt.savefig('/home/mathieu/mhenault_landrylab/Publications/mito_ma/resubmission2_GRes/fig/assemblies.mt_S288c.svg', dpi=300)
#plt.show()
plt.close()
```
