## Supplemental Code for "Hybridization drives mitochondrial DNA degeneration and metabolic shift in a species with biparental mitochondrial inheritance": nj_trees.ipynb

```
# %% [code]
library(ape)

# %% [code]
#list gene alignment fasta files to parse
S = c('atp6.anchor.aln.fasta',
      'cob.spliced.aln.MSH-604.fasta',
      'atp9.anchor.aln.fasta',
      'rps3.anchor.aln.fasta',
      'cox2.anchor.aln.fasta',
      'cox3.anchor.aln.fasta',
      'cox1.spliced.aln.fasta',
      'atp8.anchor.aln.fasta',
      'ai1.aln.fasta',
      'ai3beta.aln.fasta',
      'ai3gamma.aln.fasta',
      'ai5alpha.aln.fasta',
      'ai5beta.aln.fasta',
      'ai5gamma.aln.fasta',
      'bi1alpha.aln.fasta',
      'bi1beta.aln.fasta',
      'bi2.aln.fasta',
      'bi3.aln.fasta',
      'ORF1.aln.fasta',
      'cox3_orf.aln.fasta',
      'atp6_orf.aln.fasta')

# define index to fetch the label for each filename in the list
LI = c(rep(3, 8), rep(2, 13))
# define scale bar length for each filename in the list
SC = c(rep(0.005, 8), c(0.0015, 0.05, 0.01, 0.005, 0.05, 0.005, 0.02, 0.02, 0.02, 0.05, 0.05, 0.01, 0.005))

# %% [code]
# compute, plot and export NJ trees
for(i in seq(1,21)){

    s = S[i]
    li = LI[i]
    sc = SC[i]
    
    seq <- read.dna(paste('/mnt/HDD3/mito_nanopore/muscle/aln/', s, sep=''), format='fasta')
    
    svg(paste('/mnt/HDD3/mito_nanopore/nj_trees/', s, '.svg', sep=''), 3,3)
    par(xpd=TRUE, mar=rep(1, 4))
    
    t <- nj(dist.dna(seq))
    L = strsplit(t$tip.label, '\\.')
    newL = c()
    for(l in L){
        newL = c(newL, paste('  ', l[li], '  ', sep=''))
    }
    t$tip.label <- newL
    plot.phylo(t, type='phylogram', cex=1)
    add.scale.bar(sc*2, 1)
    title(paste(i, '. ', strsplit(s, '\\.')[[1]][1], sep=''))

    dev.off()
}
```
