## Supplemental Code for "Hybridization drives mitochondrial DNA degeneration and metabolic shift in a species with biparental mitochondrial inheritance": README.md

# mito_ma: Scripts and data for the project on mtDNA evolution in yeast hybrid MA lines

Directory contents:

### nanopore_assemblies.ipynb
Notebook file for nanopore assembly and annotation of parental mtDNAs

### assembly_comparison.ipynb
Notebook file for comparison of nanopore mtDNA assemblies with previously published illumina mtDNA assemblies
lines.

### divergence.iynb
Notebook file for computation of parental evolutionary divergence (nuclear and mtDNA) of the MA crosses

### nj_trees.ipynb
Notebook file for neighbor-joining phylogenetic trees of individual mtDNA features

### mito_ma.ipynb
Notebook file for analysis of mtDNA recombination, mtDNA deletion, respiration, and aneuploidies in MA 

### make_liftover.py
Script to produce liftover tables from mugsy pairwise whole-genome alingments. Used to confirm marker variants and analyze long read assemblies from MA lines.

### mtDNA_assemblies/
Direcrtory containing mtDNA assemblies (.fasta), processed annotations (.gff), and raw MFannot outputs (.new, .new.sqn, .new.tbl) for the 13 parental strains of the MA crosses.
