## Supplemental Material for "Hybridization drives mitochondrial DNA degeneration and metabolic shift in a species with biparental mitochondrial inheritance"

Supplemental Figure S10. Paintings of mtDNA recombination tracts for independent haplotypes of the BC2 cross. ..19
Supplemental Figure S11. Paintings of mtDNA recombination tracts for independent haplotypes of the BA1 cross. ..20
Supplemental Figure S12. Paintings of mtDNA recombination tracts for independent haplotypes of the BA2 cross. ..21
Supplemental Figure S13. Paintings of mtDNA recombination tracts for independent haplotypes of the BSc1 cross. 22
Supplemental Figure S14. Paintings of mtDNA recombination tracts for independent haplotypes of the BSc2 cross. 23

### Supplemental Table S1. Parental strains of the MA experiment and mtDNA assembly statistics.

| Strain | Species | Lineage | Genotype | | |
|---|---|---|---|---|---|
| | | | MAT | *HO* (YDL277C) | *ADE2* (YOR128C) |
| LL2011_004 | *S. paradoxus* | *SpC* | α | *hoΔ::kanMX4* | *ade2Δ::hphNT1* |
| LL2011_009 | *S. paradoxus* | *SpC* | α | *hoΔ::natMX4* | *ade2Δ::hphNT1* |
| MSH-587-1 | *S. paradoxus* | *SpC* | a | *hoΔ::natMX4* | *ade2Δ::hphNT1* |
| LL2011_012 | *S. paradoxus* | *SpC* | a | *hoΔ::kanMX4* | *ade2Δ::hphNT1* |
| LL2011_001 | *S. paradoxus* | *SpC* | a | *hoΔ::kanMX4* | *ade2Δ::hphNT1* |
| MSH-604 | *S. paradoxus* | *SpB* | a | *hoΔ::natMX4* | *ade2Δ::hphNT1* |
| UWOPS-91-202 | *S. paradoxus* | *SpB* | a | *hoΔ::kanMX4* | *ade2Δ::hphNT1* |
| LL2012_028 | *S. paradoxus* | *SpB* | α | *hoΔ::kanMX4* | *ade2Δ::hphNT1* |
| LL2012_021 | *S. paradoxus* | *SpB* | α | *hoΔ::natMX4* | *ade2Δ::hphNT1* |
| YPS644 | *S. paradoxus* | *SpA* | α | *hoΔ::kanMX4* | *ade2Δ::hphNT1* |
| YPS744 | *S. paradoxus* | *SpA* | α | *hoΔ::natMX4* | *ade2Δ::hphNT1* |
| LL2013_040 | *S. cerevisiae* | *S. cerevisiae* | α | *hoΔ::kanMX4* | *ade2Δ::hphNT1* |
| LL2013_054 | *S. cerevisiae* | *S. cerevisiae* | α | *hoΔ::natMX4* | *ade2Δ::hphNT1* |

| Strain | Parent in crosses | Strain reference | MtDNA assembly | | |
|---|---|---|---|---|---|
| | | | Size (bp) | Number of contigs | Reference |
| LL2011_004 | CC1, BC1 | Charron et al. 2019 | 73 893 | 1 | This study |
| LL2011_009 | CC2, CC3, BC2 | Charron et al. 2019 | 73 862 | 1 | This study |
| MSH-587-1 | CC1 | Hénault et al. 2020 | 73 830 | 1 | This study |
| LL2011_012 | CC2 | Hénault et al. 2020 | 73 825 | 1 | This study |
| LL2011_001 | CC3 | Hénault et al. 2020 | 73 877 | 1 | This study |
| MSH-604 | BB1, BC1, BA1, BSc1 | Charron et al. 2019 | 70 885 | 1 | This study |
| UWOPS-91-202 | BB2, BC2, BA2, BSc2 | Charron et al. 2019 | 72 525 | 1 | This study |
| LL2012_028 | BB1 | Charron et al. 2019 | 76 828 | 1 | This study |
| LL2012_021 | BB2 | Charron et al. 2019 | 78 198 | 1 | This study |
| YPS644 | BA1 | Charron et al. 2019 | 71 268 | 1 | This study |
| YPS744 | BA2 | Charron et al. 2019 | 71 438 | 1 | This study |
| LL2013_040 | BSc1 | Charron et al. 2019 | 79 881 | 1 | This study |
| LL2013_054 | BSc2 | Charron et al. 2019 | 79 900 | 1 | This study |

**Supplemental Table S2. Summary of the sequenced MA lines.**
Supplemental_Table_S2.xslx

**Supplemental Table S3. Frequency of distinct mtDNA haplotype segregation between the initial and final timepoints of the MA experiment.**

| Cross | Same haplotype | Distinct haplotypes | % distinct haplotypes |
|---|---|---|---|
| CC1 | 46 | 1 | 2.13 |
| CC2 | 38 | 4 | 9.52 |
| CC3 | 42 | 3 | 6.67 |
| BB1 | 36 | 1 | 2.70 |
| BB2 | 34 | 3 | 8.11 |
| BC1 | 41 | 6 | 12.77 |
| BC2 | 42 | 4 | 8.70 |
| BA1 | 31 | 5 | 13.89 |
| BA2 | 26 | 13 | 33.33 |
| BSc1 | 33 | 4 | 10.81 |
| BSc2 | 31 | 8 | 20.51 |

**Supplemental Table S4. Main mtDNA deletion breakpoints identified by long-read assembly.**

| Cross | Line | Left breakpoint | | Right breakpoint | |
|---|---|---|---|---|---|
| | | Position (bp) | Annotation | Position (bp) | Annotation |
| CC1 | J54 | 51 584 | Other | 55 526 | Other |
| BC2 | D12 | 53 281 | Other | 2009 | Other |
| BA1 | B54 | 16 236 | Other | 38 917 | Other |
| BA1 | B67 | 48 974 | Other | 68 470 | GC cluster |
| BA2 | E25 | 55 734 | Other | 65 346 | Other |
| BA2 | E34 | 56 333 | RNA exon | 79 051 | Intron |
| BA2 | E84 | 22 332 | Other | 65 346 | Other |
| BSc1 | C26 | 22 174 | Other | 26 734 | Other |
| BSc1 | C31 | 60 749 | Other | 68 747 | Other |
| BSc2 | F15 | 60 749 | Other | 68 469 | GC cluster |
| BSc2 | F21 | 22 924 | Other | 39 206 | Other |
| BSc2 | F3 | 49 838 | Exon | 63 855 | RNA exon |
| BSc2 | F48 | 23 771 | Other | 26 735 | Other |
| BSc2 | F79 | 16 482 | Other | 49 984 | Exon |
| BSc2 | F8 | 32 174 | GC cluster | 46 271 | Other |

**Supplemental Table S5. Candidate de novo mutations in MA lines mtDNAs.**

| Cross | Line | Time-point | Position | Ref. allele | Alt. allele | Total reads | Ref. allele support | Alt. allele support | Ratio of missing data | High confidence support | Support elsewhere in dataset | Annotation |
|---|---|---|---|---|---|---|---|---|---|---|---|---|
| BB2 | H30 | final | 23 040 | T | A | 82 | 14 | 68 | 0.333333 | 15 | 0 | intergenic |
| BC2 | D68 | initial | 22 466 | A | T | 14 | 0 | 14 | 0.384615 | 13 | 6 | intergenic |
| BA2 | E15 | final | 53 459 | G | A | 26 | 2 | 24 | 0.368421 | 19 | 11 | intergenic |
| BA2 | E34 | final | 62 019 | G | T | 871 | 136 | 735 | 0.118421 | 3 | 18 | intergenic |
| BSc1 | C78 | initial | 24 590 | A | T | 108 | 1 | 107 | 0.094595 | 6 | 8 | intergenic |
| BSc1 | C26 | final | 16 760 | G | A | 19 | 2 | 17 | 0.391892 | 9 | 1 | intergenic |

### Supplemental Table S6. Primer sequences and PCR cycles.

| Locus | Primer name | Primer sequence | Amplicon size | TM | Elongation time | Reference |
|---|---|---|---|---|---|---|
| *ATP6* | CLOP80-C3 | GGTTCAAGATGATTAATTTCACAAG | 255 bp | 58°C | 20s | Leducq et al. 2017 |
| | CLOP80-D3 | ACCAGCAGGTACGAATAATGA | | | | |
| 21S rRNA | CLOP80-A3 | TGAGGTCCCGCATGAATGAC | 716 bp (*SpB*), 669 bp (*SpA*) | 58°C | 48s | Leducq et al. 2017 |
| | CLOP80-B3 | ACGTACTTGTTTCACTCGTTTGT | | | | |
| *MAT* | CLO1-42 | ACTCCACTTCAAGTAAGAGTTTG | 404 bp (MATα), 544 bp (MATa) | 54°C | 35s | Huxley et al. 1990 |
| | CLO1-43 | GCACGGAATATGGGACTACTTCG | | | | |
| | CLO1-44 | AGTCACATCAAGATCGTTTATGG | | | | |

| Step | Temperature | Duration | Repeats |
|---|---|---|---|
| 1 | 94°C | 3min | 1X |
| 2 | 94°C | 30s | 35X |
| | TM | 30s | |
| | 72°C | Elongation | |
| 3 | 72°C | 10min | 1X |

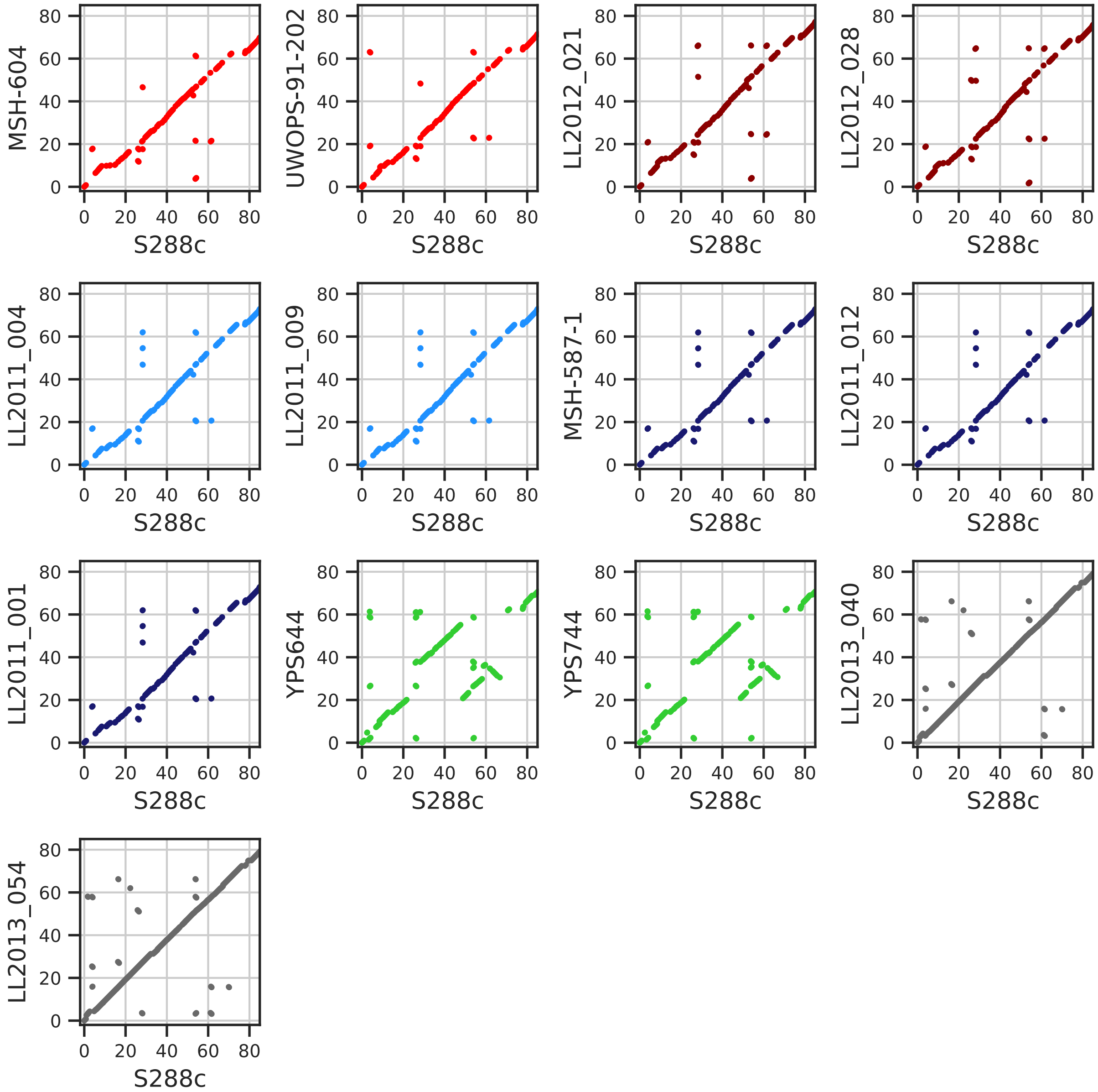

**Supplemental Figure S1. Synteny comparison of the parental mtDNA assemblies.** Coordinates show the assembly position against the *S. cerevisiae* S288c reference mtDNA (Yue et al. 2017) in kilobases. Dot plots are colored by lineage/species (red: *SpB*, blue: *SpC*, green: *SpA*, black: *S. cerevisiae*).

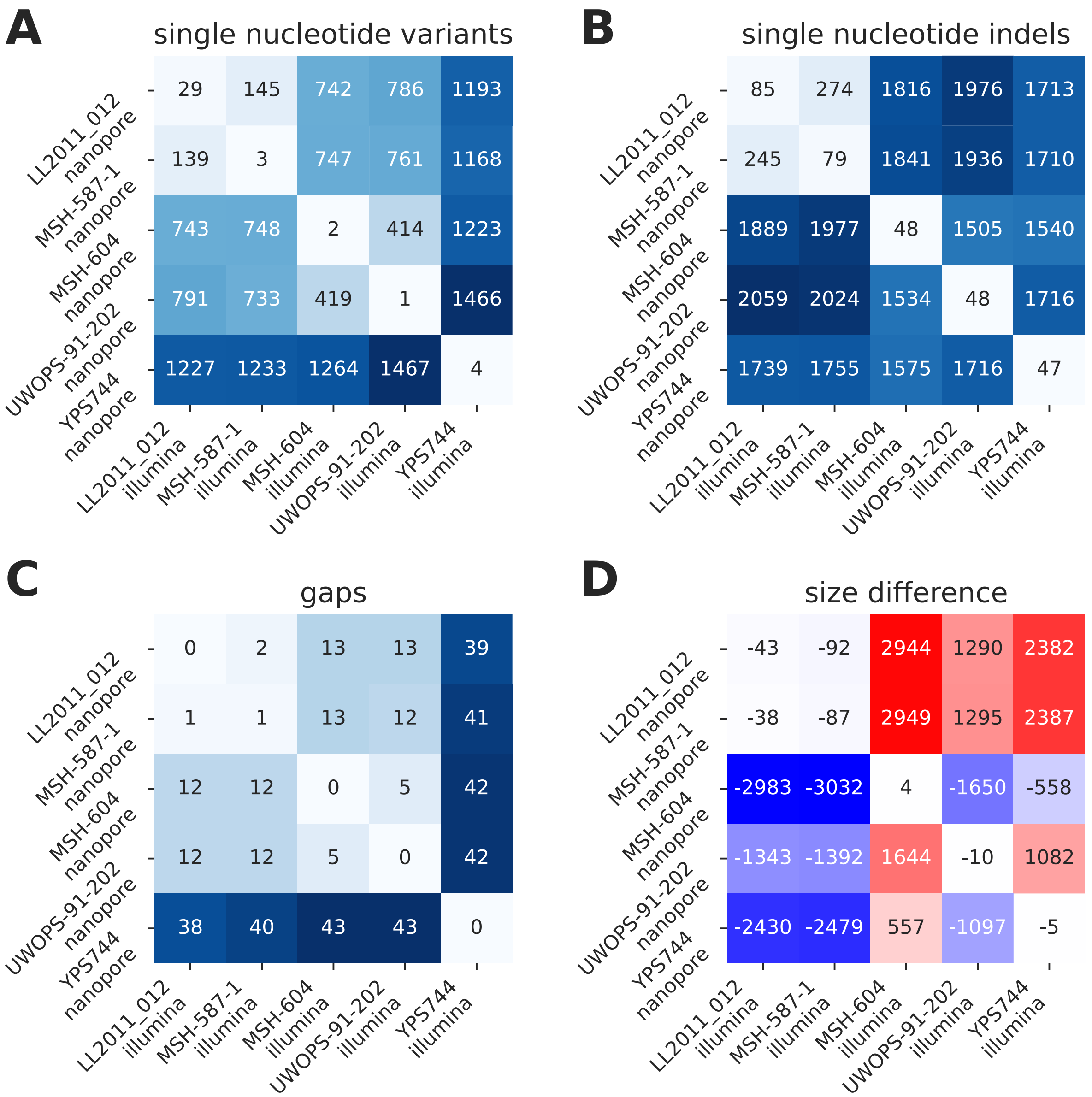

**Supplemental Figure S2. Sequence comparisons of parental mtDNA assemblies.** Sequence comparisons between five mtDNAs assembled from Illumina short reads by Leducq et al. 2017, and the mtDNAs assemblies for the corresponding strains from the current study. Counts of sequence differences are reported for single nucleotide variants *(A)*, single nucleotide indels *(B)* and gaps *(C)*. The difference in total assembly size is reported in *(D)*, with each value representing the difference (in bp) between the nanopore assembly (this study) minus the Illumina assembly (Leducq et al. 2017). For each strain, the nanopore and Illumina assemblies are entirely colinear.

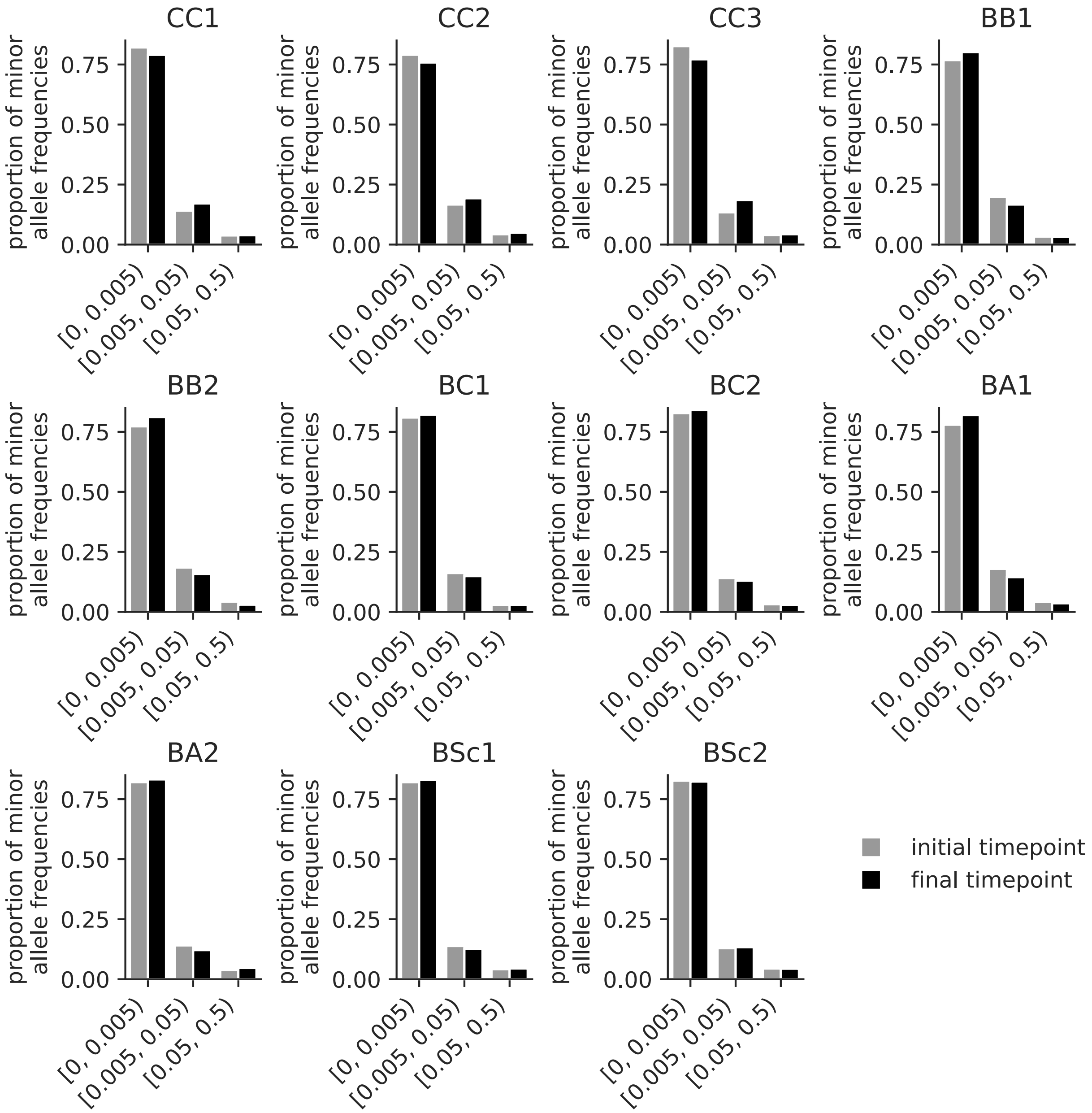

**Supplemental Figure S3. Minor allele frequency distributions of mtDNA variants.**
Variants in mtDNAs of MA lines sampled at the initial (grey) and final (black) timepoints are shown.

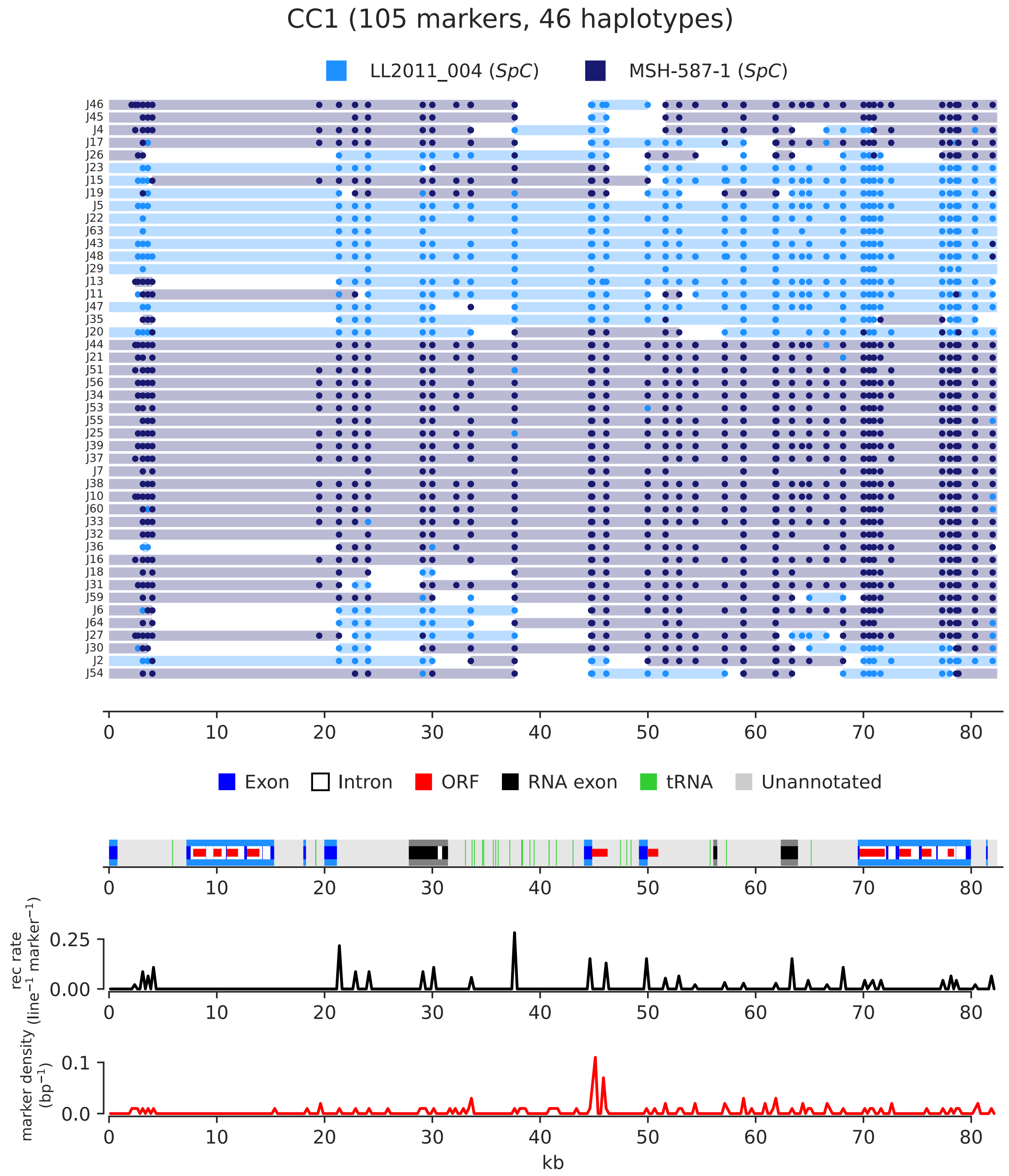

**Supplemental Figure S4. Paintings of mtDNA recombination tracts for independent haplotypes of the CC1 cross.**

Individual lines are stacked horizontally and tracts are colored by parental ancestry. Marker variants supporting each tract are shown as dots. Annotation summary is shown at the bottom. Recombination rate and marker density in 100-bp non-overlapping windows are shown at the bottom in black and red, respectively. Genome coordinates are in kb.

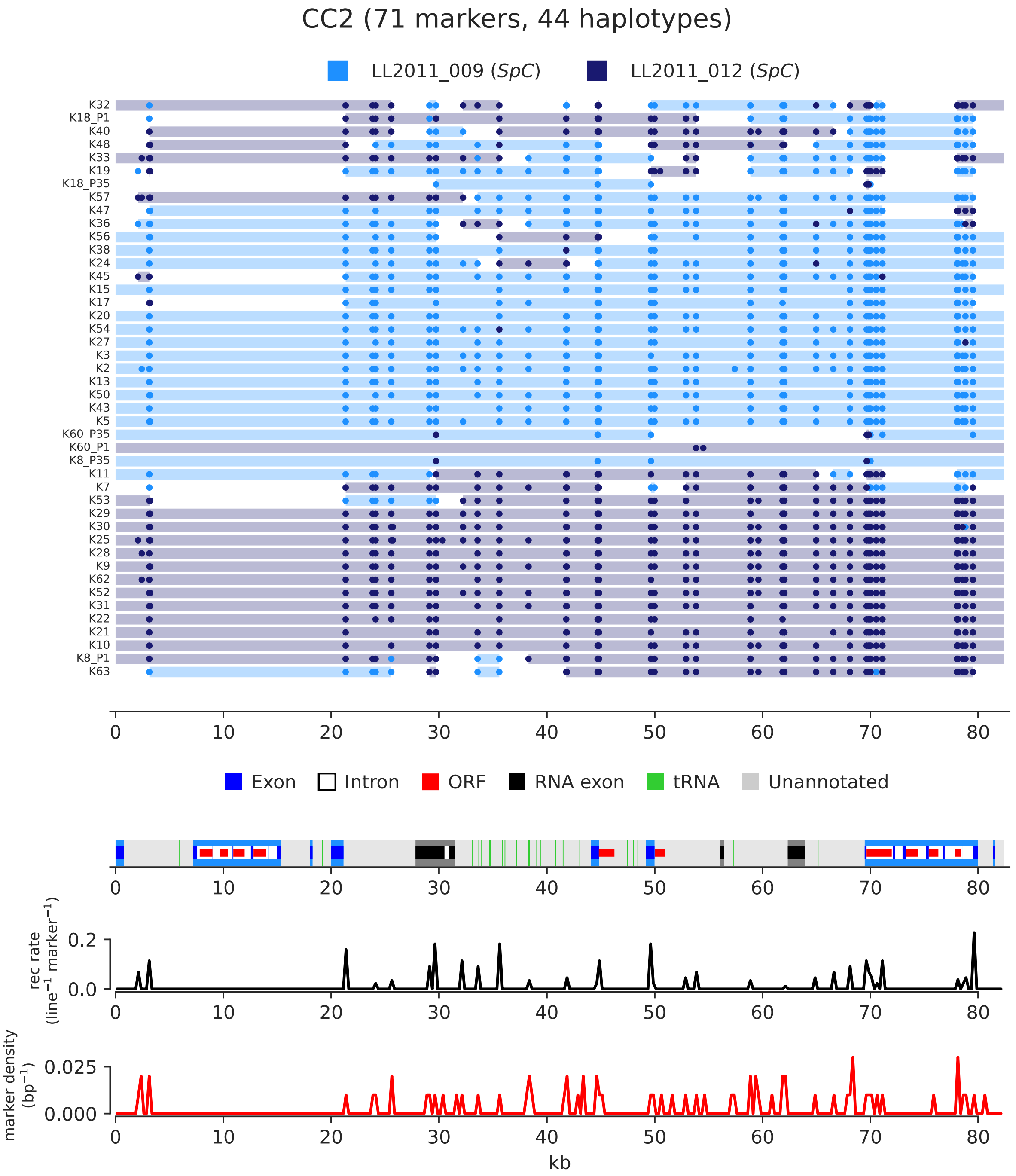

**Supplemental Figure S5. Paintings of mtDNA recombination tracts for independent haplotypes of the CC2 cross.**

Individual lines are stacked horizontally and tracts are colored by parental ancestry. Marker variants supporting each tract are shown as dots. Independent haplotypes for the same line at the initial (P1) and final (P35) timepoints are shown. Annotation summary is shown at the bottom. Recombination rate and marker density in 100-bp non-overlapping windows are shown at the bottom in black and red, respectively. Genome coordinates are in kb.

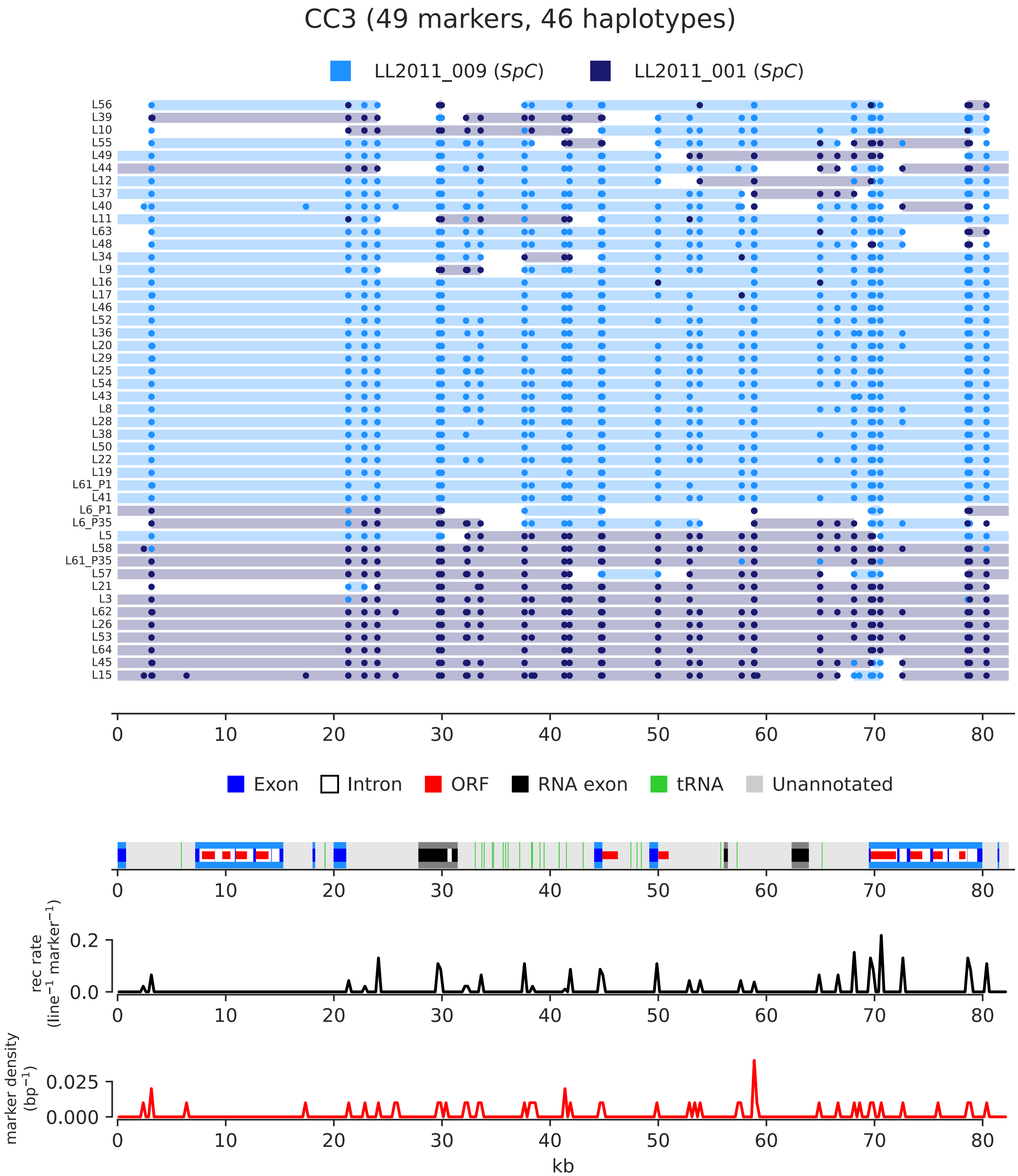

**Supplemental Figure S6. Paintings of mtDNA recombination tracts for independent haplotypes of the CC3 cross.**

Individual lines are stacked horizontally and tracts are colored by parental ancestry. Marker variants supporting each tract are shown as dots. Independent haplotypes for the same line at the initial (P1) and final (P35) timepoints are shown. Annotation summary is shown at the bottom. Recombination rate and marker density in 100-bp non-overlapping windows are shown at the bottom in black and red, respectively. Genome coordinates are in kb.

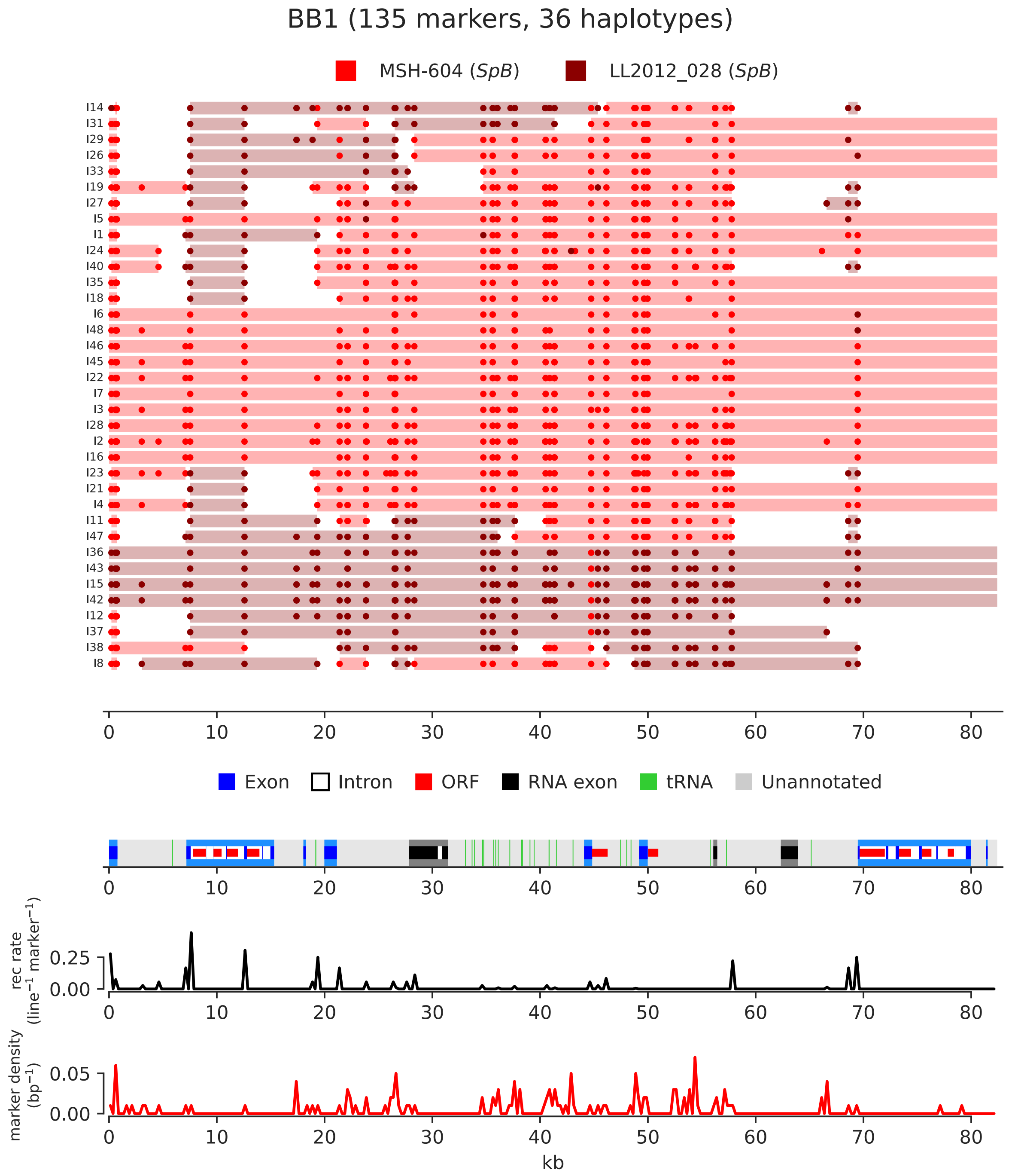

**Supplemental Figure S7. Paintings of mtDNA recombination tracts for independent haplotypes of the BB1 cross.**

Individual lines are stacked horizontally and tracts are colored by parental ancestry. Marker variants supporting each tract are shown as dots. Annotation summary is shown at the bottom. Recombination rate and marker density in 100-bp non-overlapping windows are shown at the bottom in black and red, respectively. Genome coordinates are in kb.

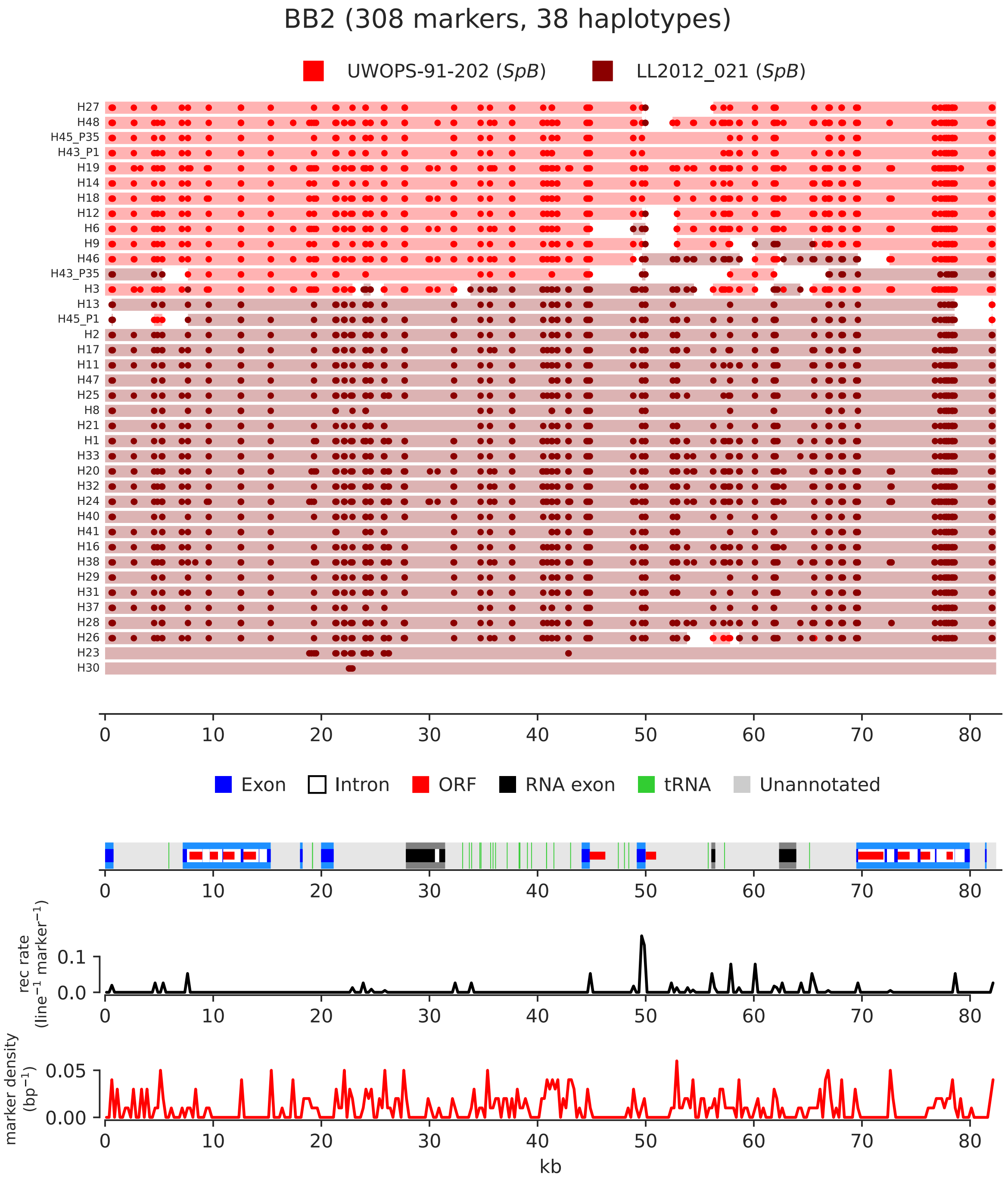

**Supplemental Figure S8. Paintings of mtDNA recombination tracts for independent haplotypes of the BB2 cross.**
Individual lines are stacked horizontally and tracts are colored by parental ancestry. Marker variants supporting each tract are shown as dots. Independent haplotypes for the same line at the initial (P1) and final (P35) timepoints are shown. Annotation summary is shown at the bottom. Recombination rate and marker density in 100-bp non-overlapping windows are shown at the bottom in black and red, respectively. Genome coordinates are in kb.

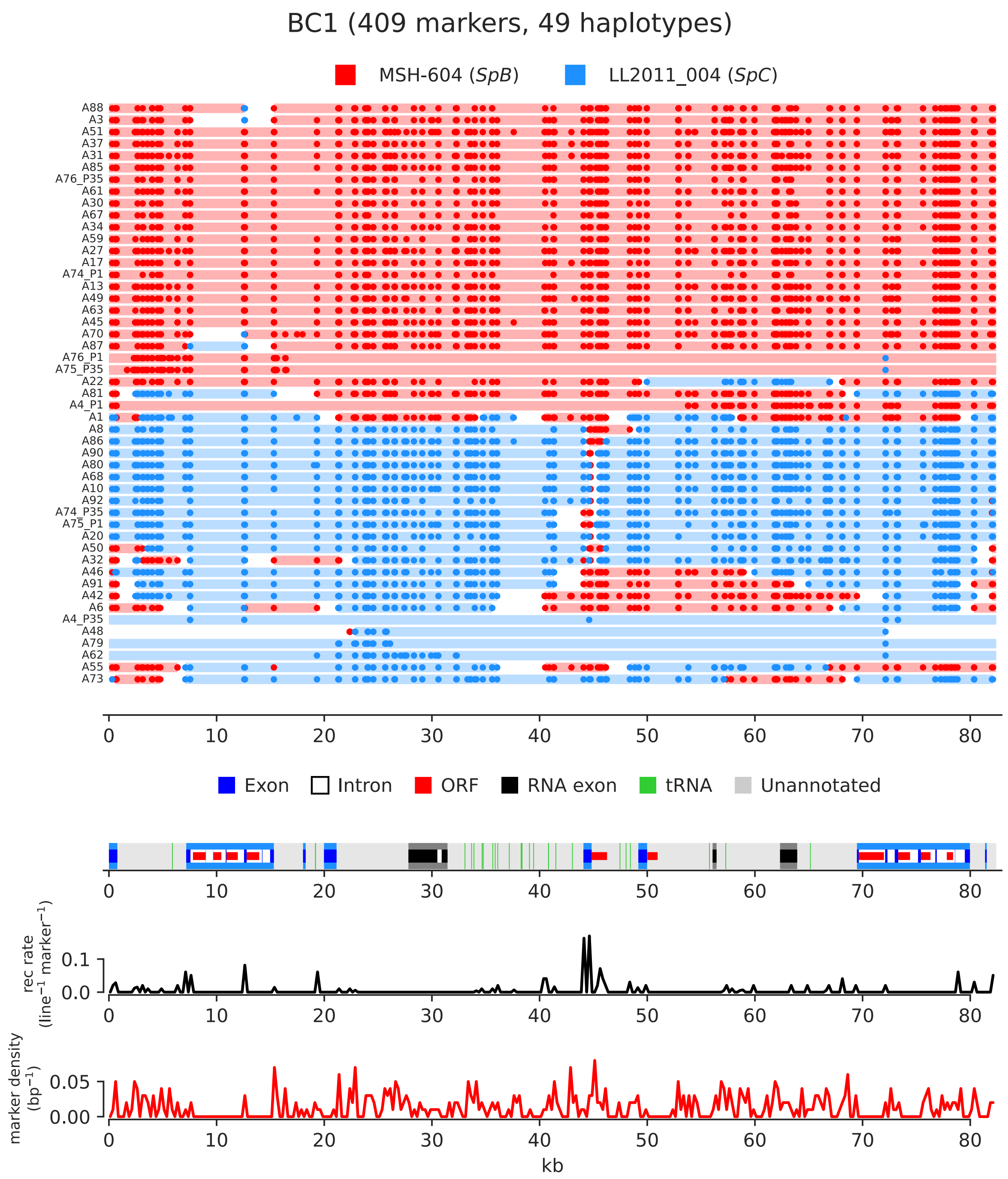

**Supplemental Figure S9. Paintings of mtDNA recombination tracts for independent haplotypes of the BC1 cross.**

Individual lines are stacked horizontally and tracts are colored by parental ancestry. Marker variants supporting each tract are shown as dots. Independent haplotypes for the same line at the initial (P1) and final (P35) timepoints are shown. Annotation summary is shown at the bottom. Recombination rate and marker density in 100-bp non-overlapping windows are shown at the bottom in black and red, respectively. Genome coordinates are in kb.

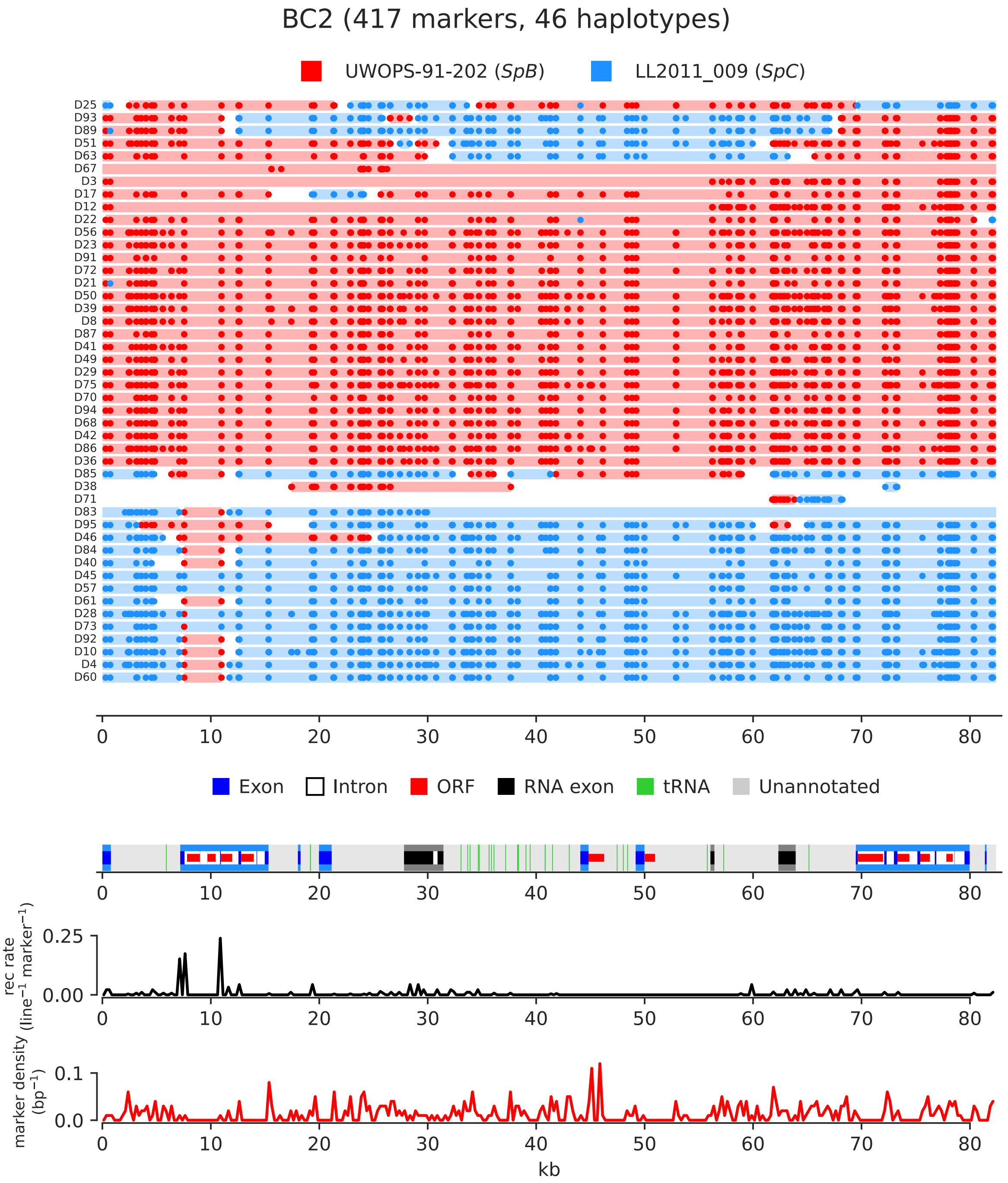

**Supplemental Figure S10. Paintings of mtDNA recombination tracts for independent haplotypes of the BC2 cross.**

Individual lines are stacked horizontally and tracts are colored by parental ancestry. Marker variants supporting each tract are shown as dots. Annotation summary is shown at the bottom. Recombination rate and marker density in 100-bp non-overlapping windows are shown at the bottom in black and red, respectively. Genome coordinates are in kb.

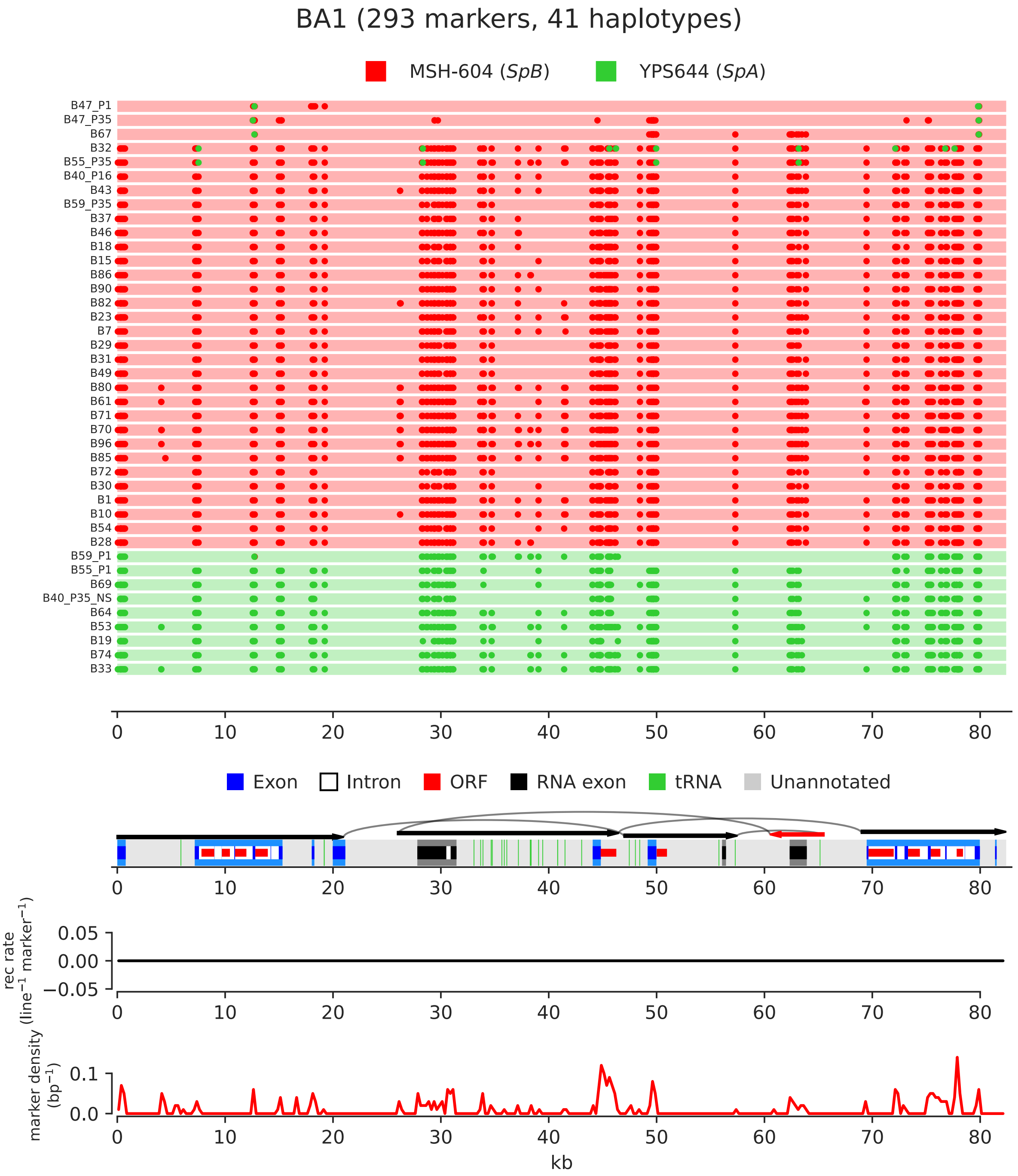

**Supplemental Figure S11. Paintings of mtDNA recombination tracts for independent haplotypes of the BA1 cross.**
Individual lines are stacked horizontally and tracts are colored by parental ancestry. Marker variants supporting each tract are shown as dots. Independent haplotypes for the same line at the initial (P1), median (P16) and final (P35) timepoints are shown. Annotation summary is shown at the bottom. Recombination rate and marker density in 100-bp non-overlapping windows are shown at the bottom in black and red, respectively. Genome coordinates are in kb. Arrows and arcs show the rearrangements common to the *SpA* mtDNAs in comparison to *S. cerevisiae*, with red arrows representing inverted segments.

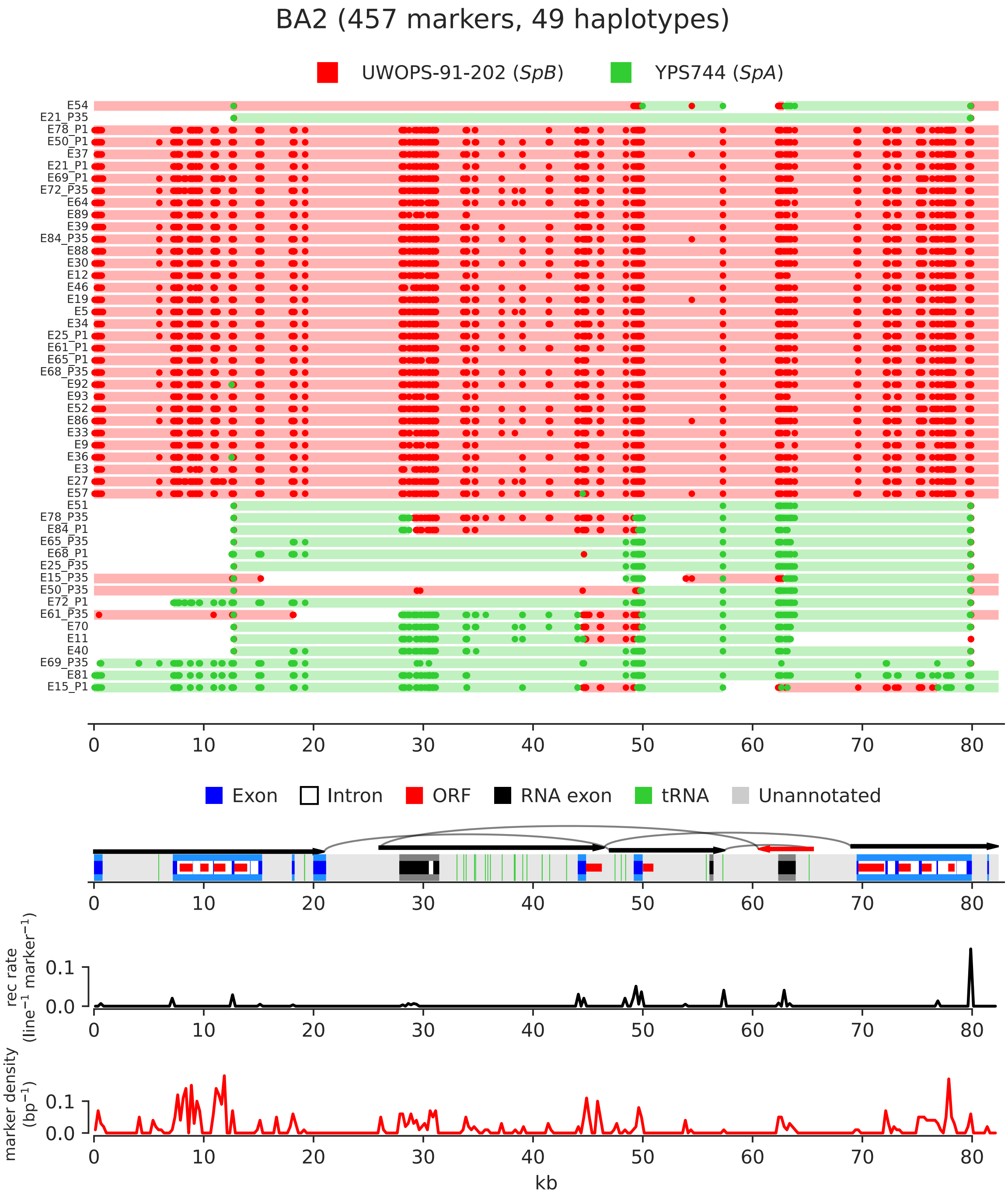

**Supplemental Figure S12. Paintings of mtDNA recombination tracts for independent haplotypes of the BA2 cross.**

Individual lines are stacked horizontally and tracts are colored by parental ancestry. Marker variants supporting each tract are shown as dots. Independent haplotypes for the same line at the initial (P1) and final (P35) timepoints are shown. Annotation summary is shown at the bottom. Recombination rate and marker density in 100-bp non-overlapping windows are shown at the bottom in black and red, respectively. Genome coordinates are in kb. Arrows and arcs show the rearrangements common to the *SpA* mtDNAs in comparison to *S. cerevisiae*, with red arrows representing inverted segments.

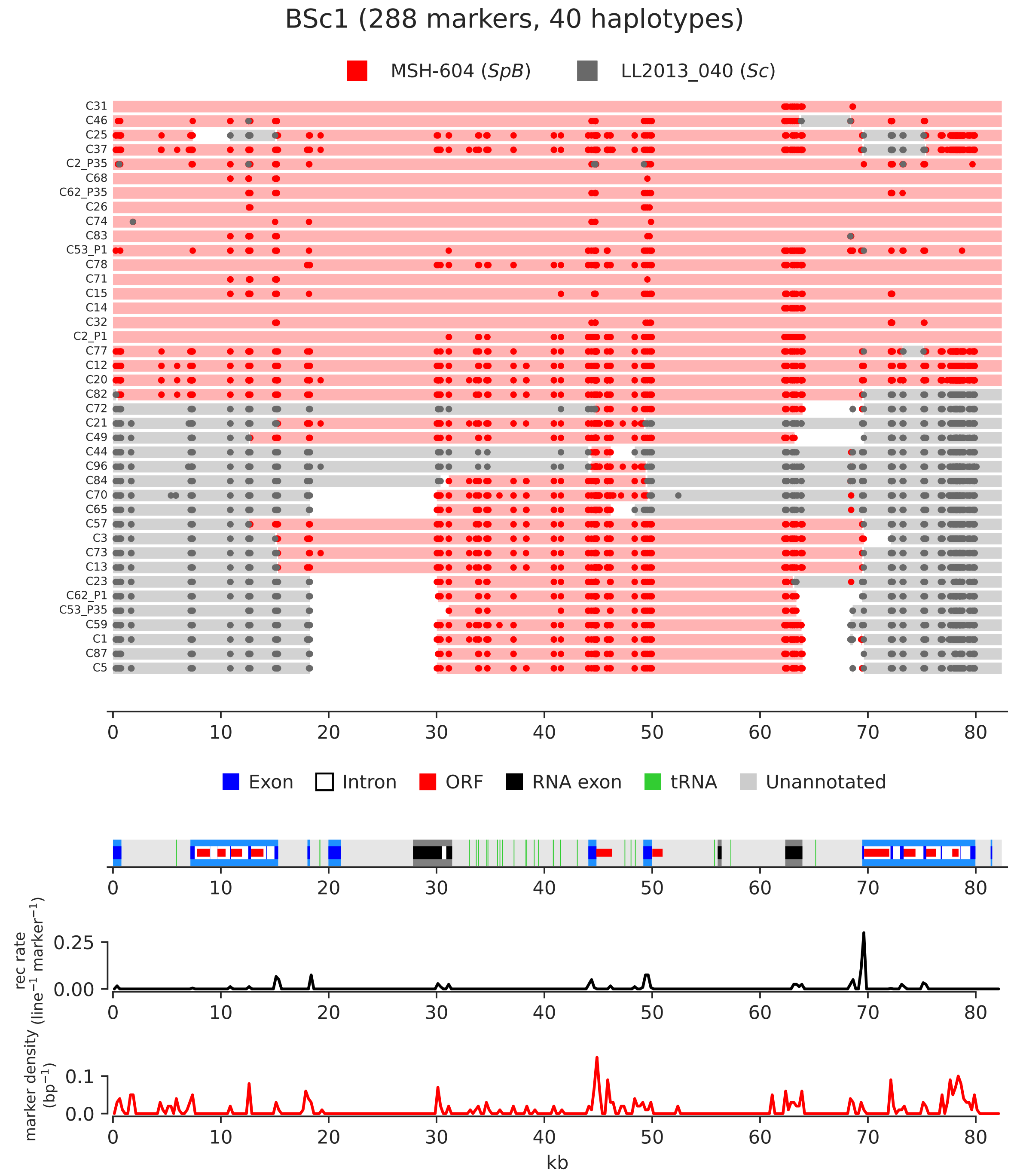

**Supplemental Figure S13. Paintings of mtDNA recombination tracts for independent haplotypes of the BSc1 cross.**
Individual lines are stacked horizontally and tracts are colored by parental ancestry. Marker variants supporting each tract are shown as dots. Independent haplotypes for the same line at the initial (P1) and final (P35) timepoints are shown. Annotation summary is shown at the bottom. Recombination rate and marker density in 100-bp non-overlapping windows are shown at the bottom in black and red, respectively. Genome coordinates are in kb.

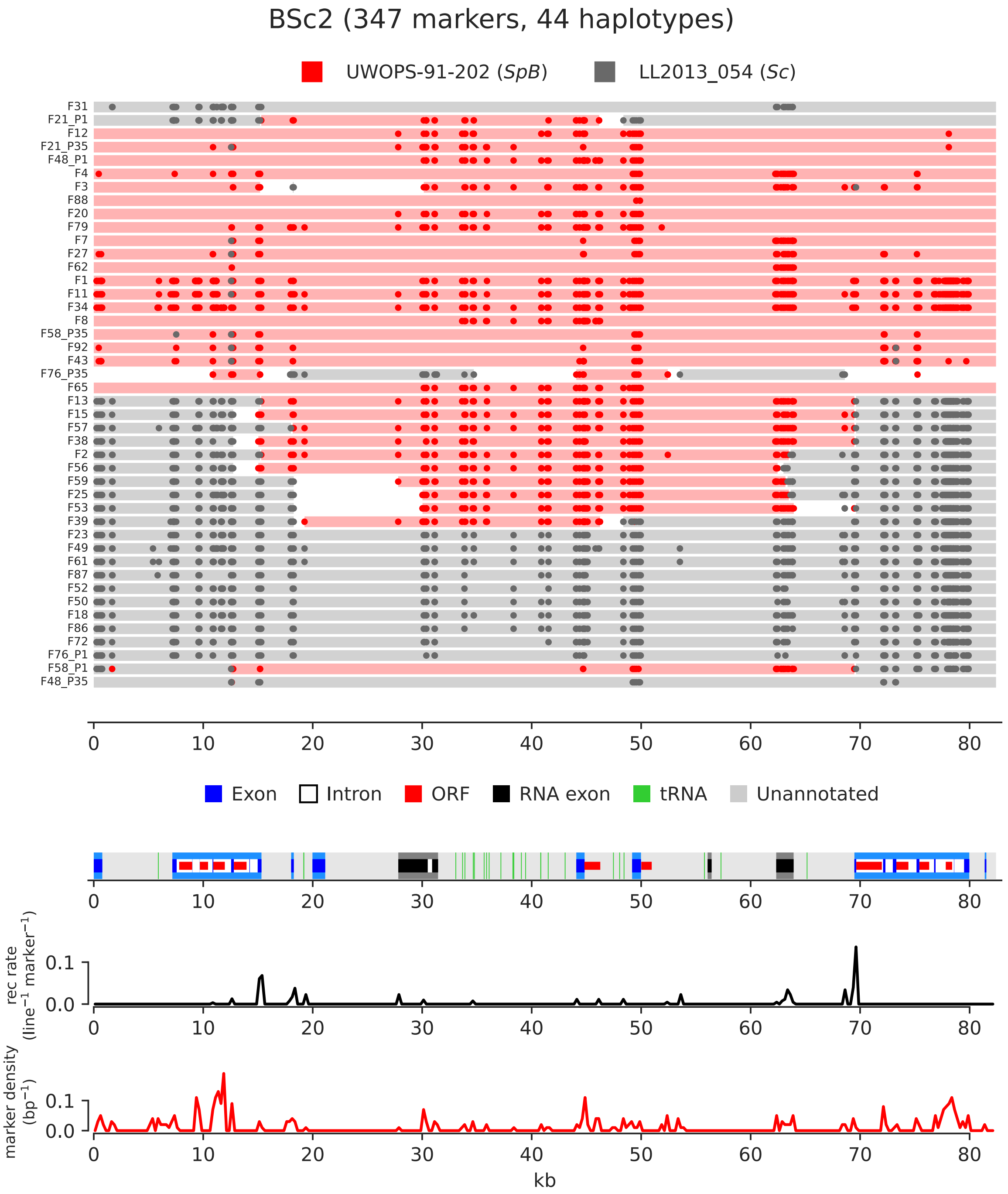

**Supplemental Figure S14. Paintings of mtDNA recombination tracts for independent haplotypes of the BSc2 cross.**

Individual lines are stacked horizontally and tracts are colored by parental ancestry. Marker variants supporting each tract are shown as dots. Independent haplotypes for the same line at the initial (P1) and final (P35) timepoints are shown. Annotation summary is shown at the bottom. Recombination rate and marker density in 100-bp non-overlapping windows are shown at the bottom in black and red, respectively. Genome coordinates are in kb.

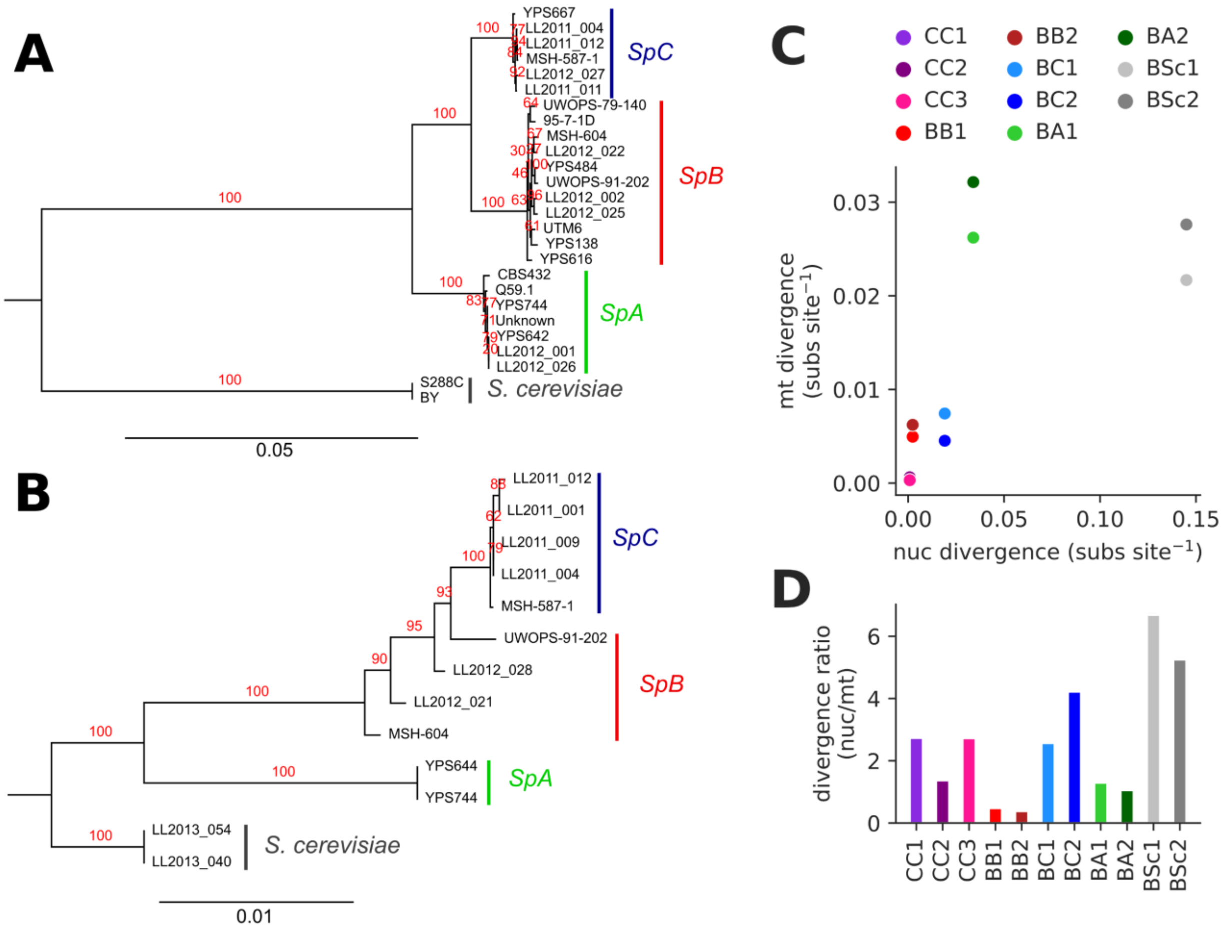

**Supplemental Figure S15. Evolutionary divergence between parental strains.**
*(A)* Maximum likelihood phylogenetic tree based on concatenated DNA sequences of 81 randomly chosen nuclear-encoded protein-coding genes for 26 strains of *S. paradoxus* and *S. cerevisiae* (Eberlein et al. 2017). The scale is in substitutions per site. Values in red indicate branch support percent from 200 bootstraps. *(B)* Maximum likelihood phylogenetic tree based on concatenated DNA sequences of the eight mtDNA-encoded protein-coding genes (excluding introns) for the 13 parental strains. The scale is in substitutions per site. Values in red indicate branch support percent from 200 bootstraps. *(C)* Nuclear and mitochondrial evolutionary divergence estimates for each MA cross. Nuclear divergence values were computed from the phylogenetic tree in *(A)*, taking the average of all pairwise distances within or between the corresponding parental clade(s). Mitochondrial divergence values were computed from the phylogenetic tree in *(B)*, taking the distances between the corresponding parental strains. *(D)* Ratio of nuclear and mitochondrial divergence for each cross.

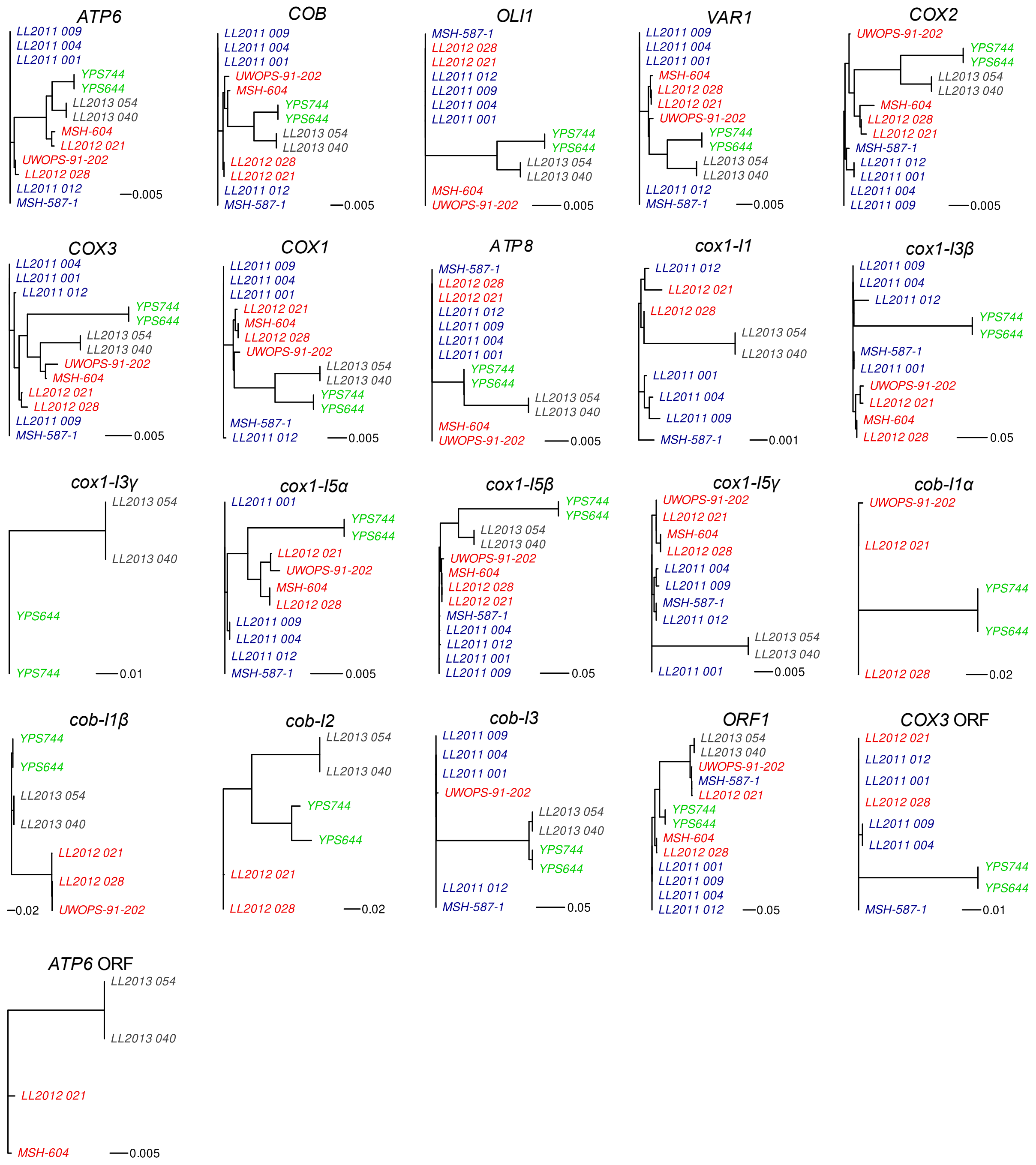

**Supplemental Figure S16. Phylogenetic trees of individual mtDNA genes.**
Neighbor-joining phylogenies for the main protein-coding genes (excluding introns), introns and ORFs. Scale bars are in substitutions per site.

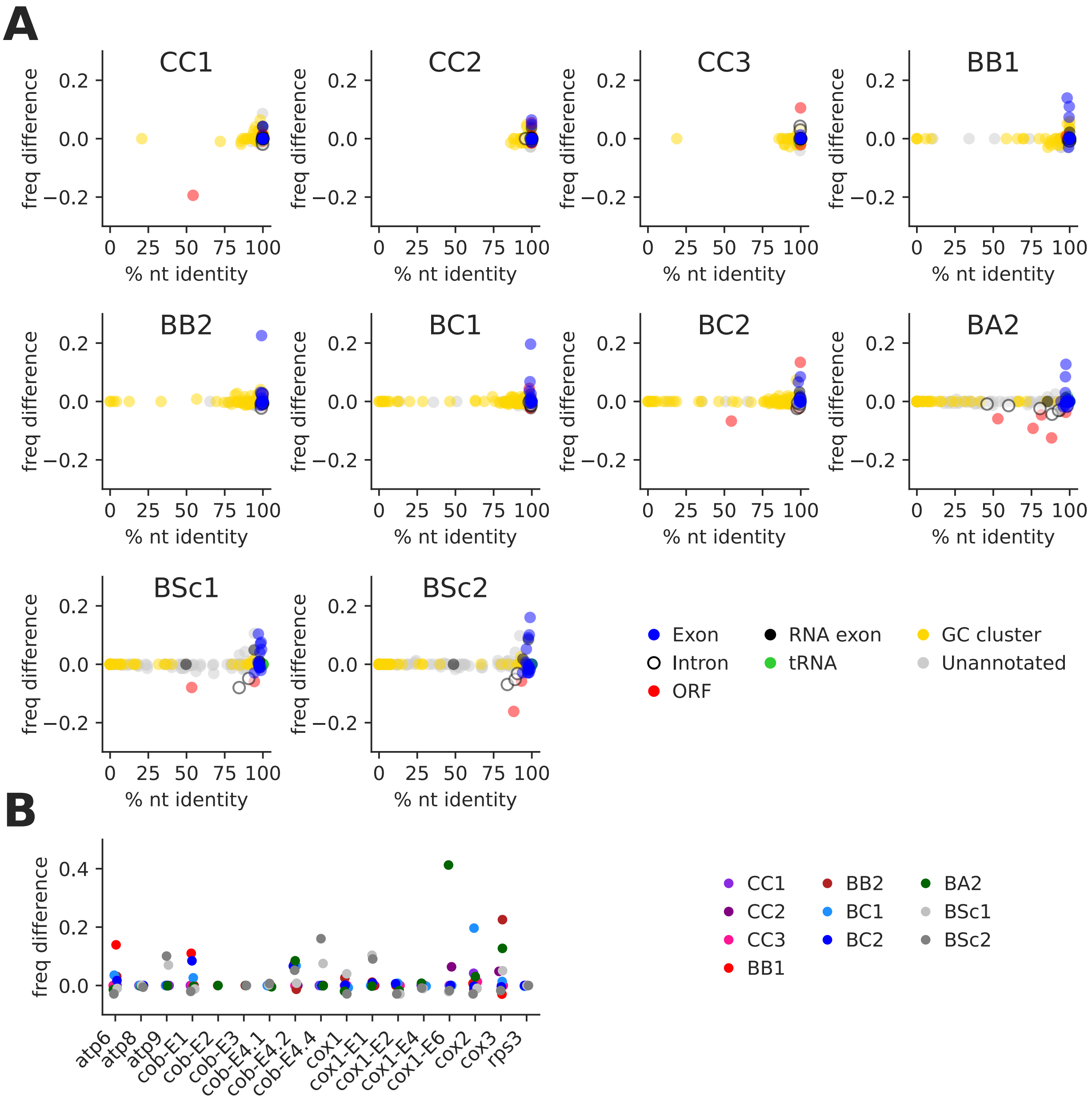

**Supplemental Figure S17. Overrepresentation of mtDNA annotations in recombination junctions.**

*(A)* Overrepresentation of individual features in recombination junctions (calculated as the difference in frequency between breakpoints and markers for each feature) against nucleotide identity between parental alleles. Unannotated regions were split into 500 bp windows. *(B)* Overrepresentation of protein-coding exons in recombination junctions.

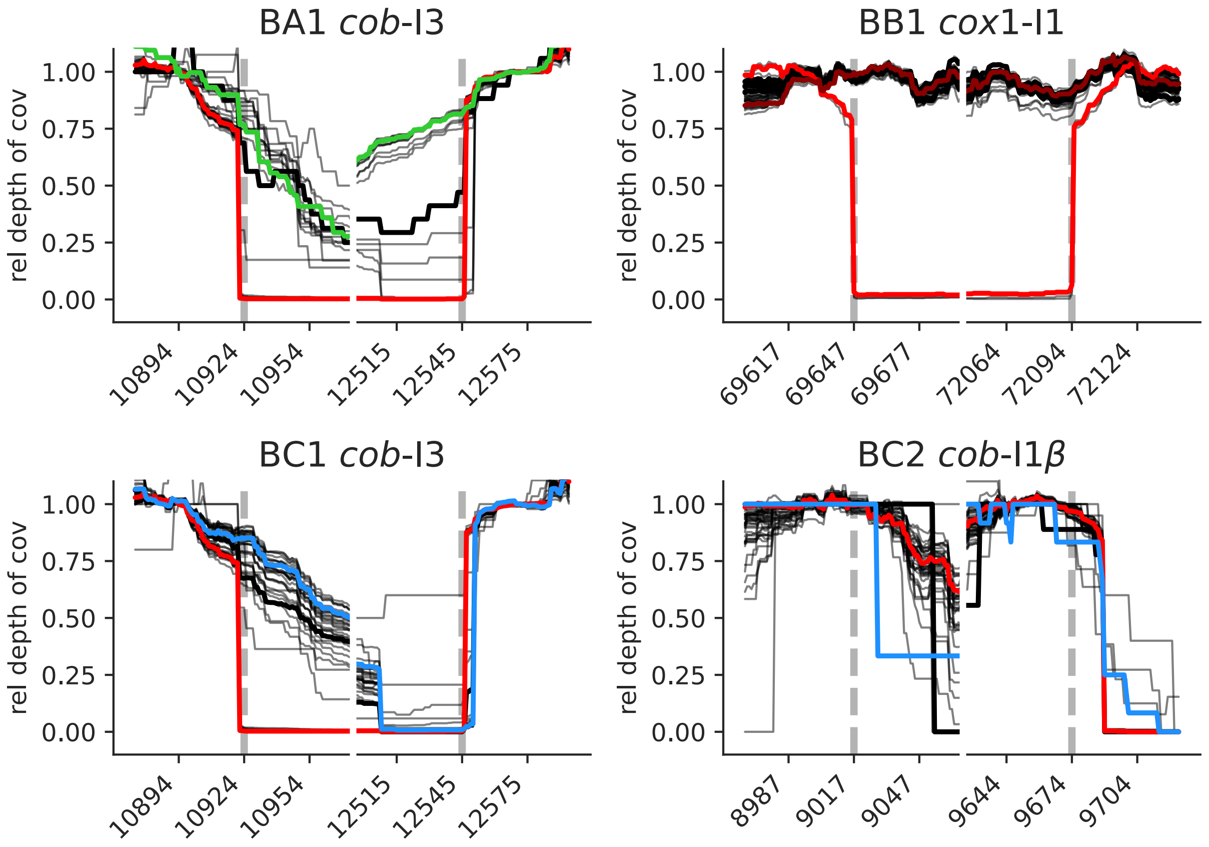

**Supplemental Figure S18. Depth of coverage profiles at putative intron mobilization sites.** Relative depth of coverage surrounding intron junctions for which putative mobility events were inferred. Vertical dotted lines indicate intron junctions. Thin lines represent the profiles of all lines of the corresponding cross. Thick colored lines represent the profiles of the parental strains (according to the color code in Fig. 1). Thick black lines represent lines for which putative mobility events were inferred. Coordinates are in bp. The putative mobility of *cob*-I3 in one BA1 line and *cob*-I1β in one BC2 line were classified as false positives after visual examination.

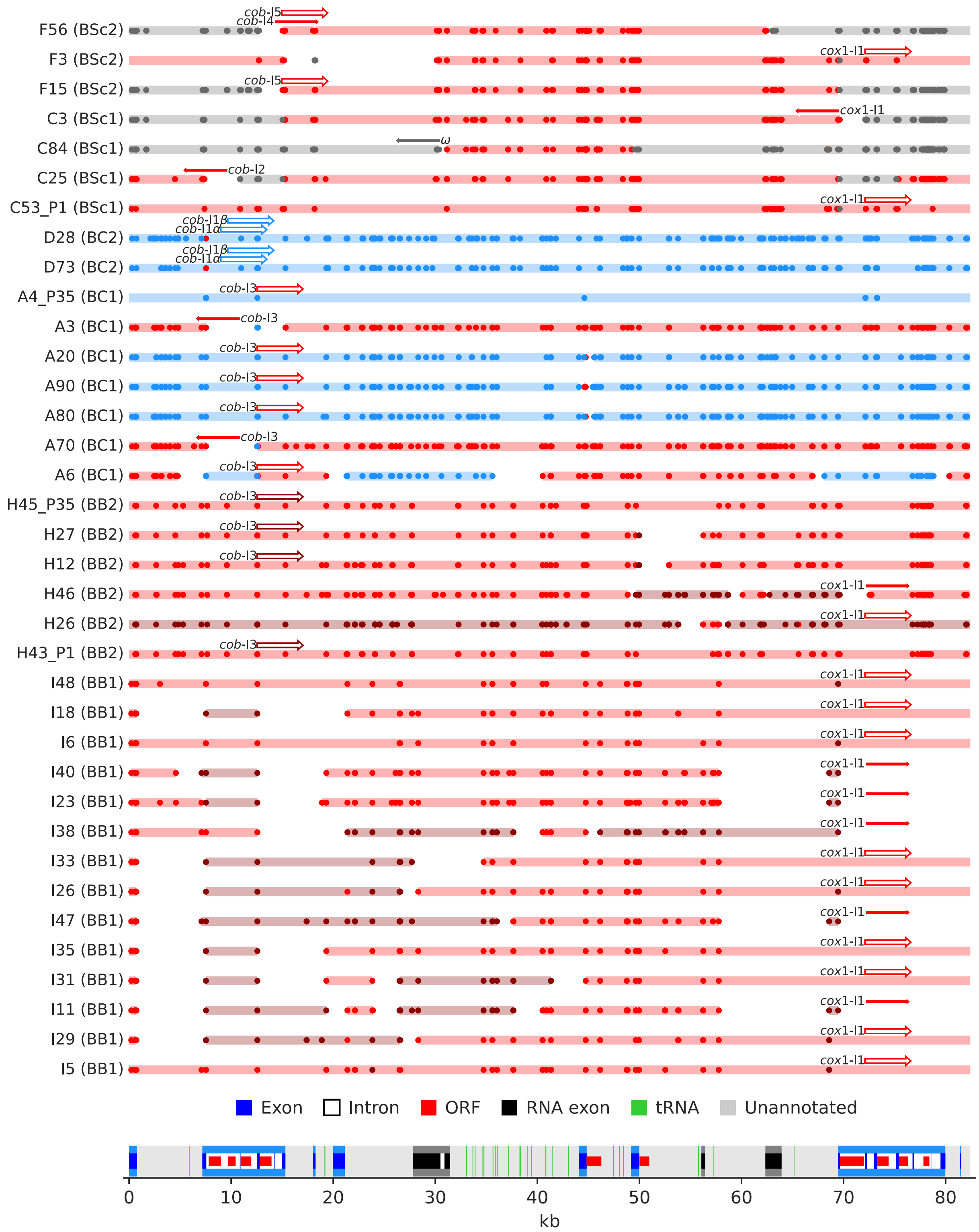

**Supplemental Figure S19. Putative recombination-associated mobilization events of polymorphic introns.**

Independent haplotypes from lines at the initial (P1) or final (P35) timepoints with putative recombination-associated intron mobilization are shown. Putative intron mobilization events are represented with arrows, with arrow color corresponding to the parental mtDNA that is natively devoid of the intron (see Fig. 1). Full and empty arrows represent putative mobilization events that are consistent and inconsistent with recombination tracts, respectively. Arrows point in the direction of the initiation of a recombination tract that would be consistent with the mobilization event. Recombination tracts and supporting marker variants are shown as in Supplemental Fig. S4-14. Annotation summary is shown at the bottom. Genome coordinates are in kb.

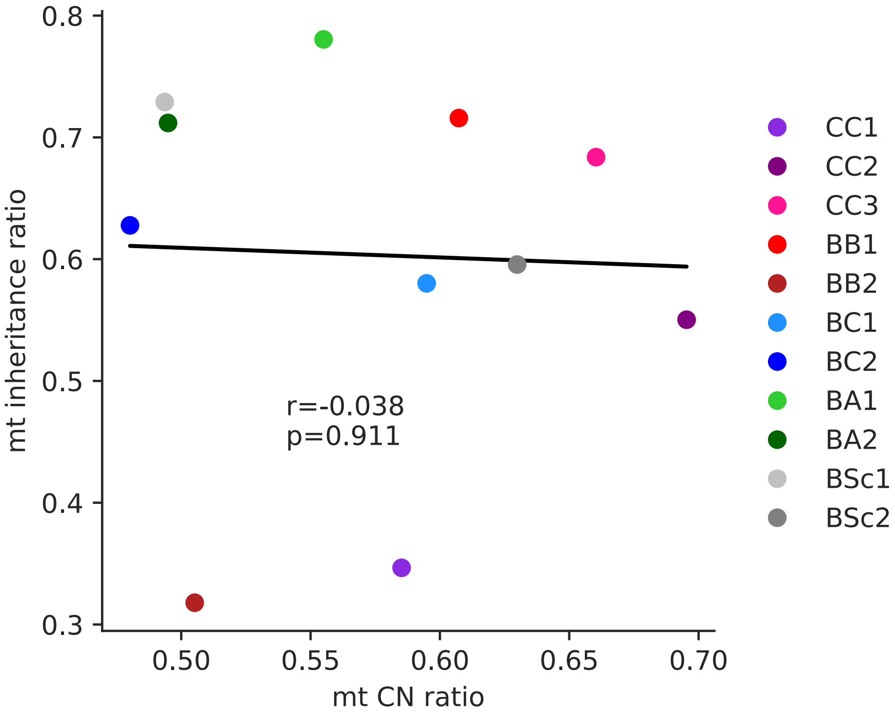

**Supplemental Figure S20. MtDNA inheritance ratios and parental mtDNA abundances.** Genome-wide inheritance ratio is shown against the ratio of mtDNA copy numbers for both parents, inferred by the depth of coverage of nanopore libraries on mtDNA assemblies relative to the nuclear genome. P-value is shown for Pearson correlation.

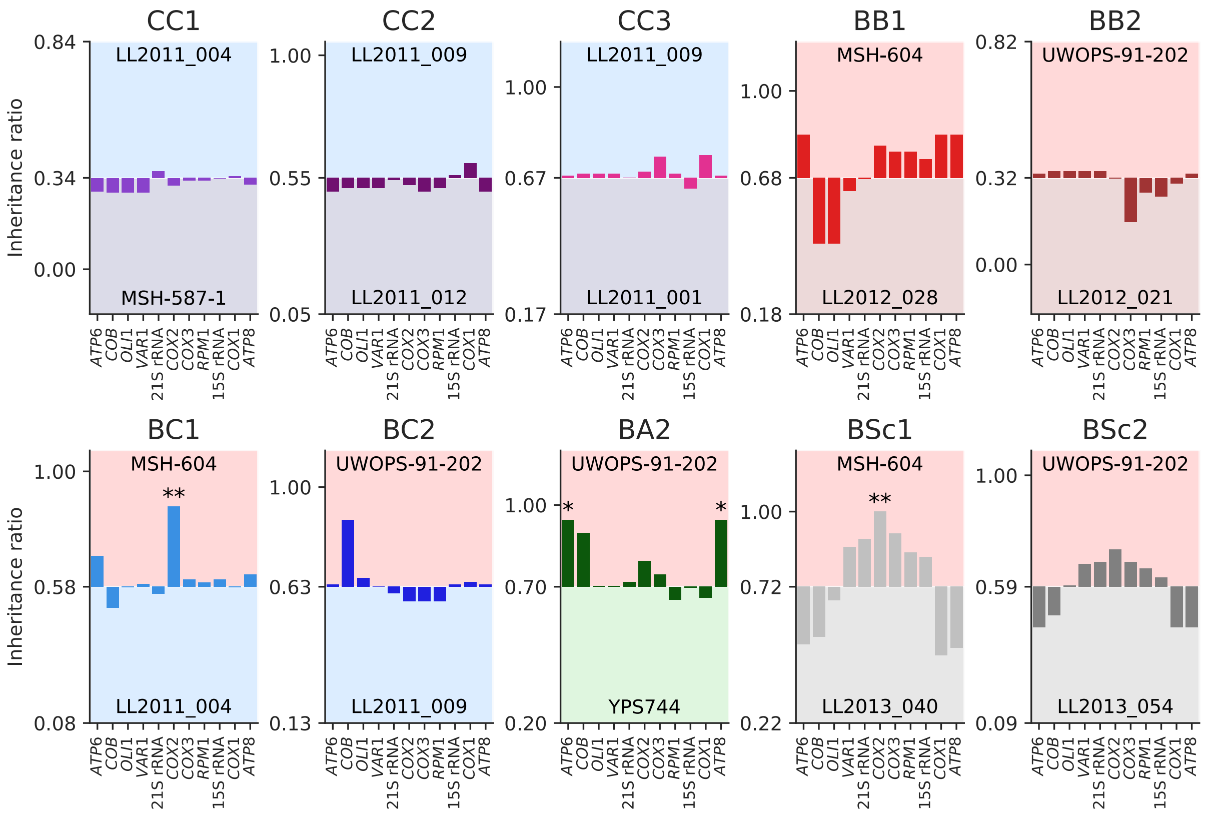

**Supplemental Figure S21. Inheritance ratios of mtDNA features among crosses.**
Deviation from the mtDNA-wide ratio for each cross is shown, ratios of 1 being complete inheritance from the upper parent. FDR-corrected p-values for binomial tests are shown. *: p≤0.05, **: p≤0.01.

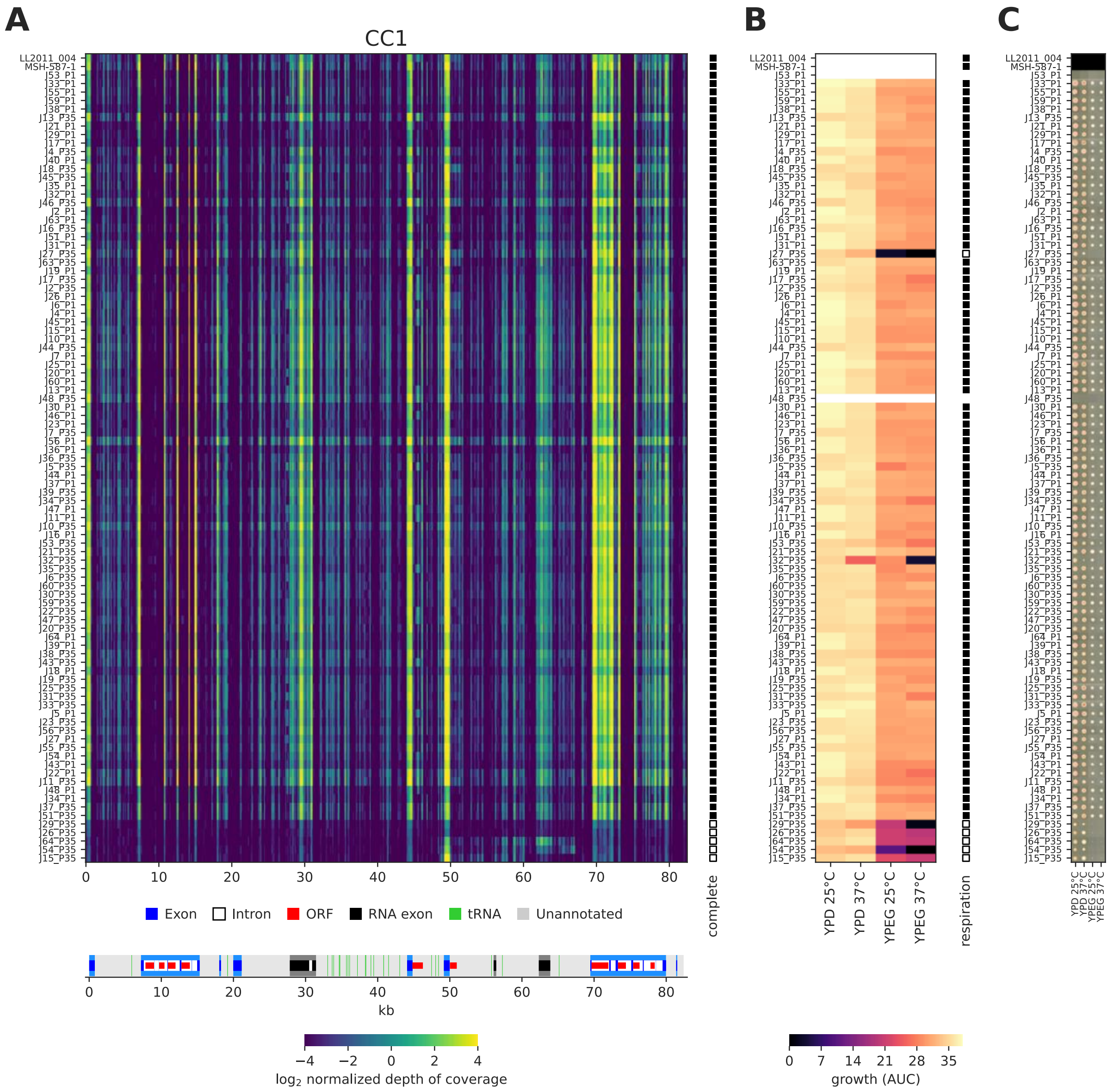

**Supplemental Figure S22. MtDNA deletions and respiration for CC1 lines.**
(*A*) Depth of coverage of short reads along the mtDNA sequence for haplotypes of the CC1 cross at the initial (P1) and final (P35) timepoints. Individual MA lines are stacked horizontally, with the parental strains at the top. Full and empty squares at the right of the heatmap show haplotype classification as complete or deleted, respectively. Annotation summary is shown at the bottom. Genome coordinates are in kb. (*B*) Growth AUC for corresponding lines. Full and empty squares at the right of the heatmap show line classification as respiring or non-respiring, respectively. (*C*) Raw colony images after four days of incubation for corresponding lines.

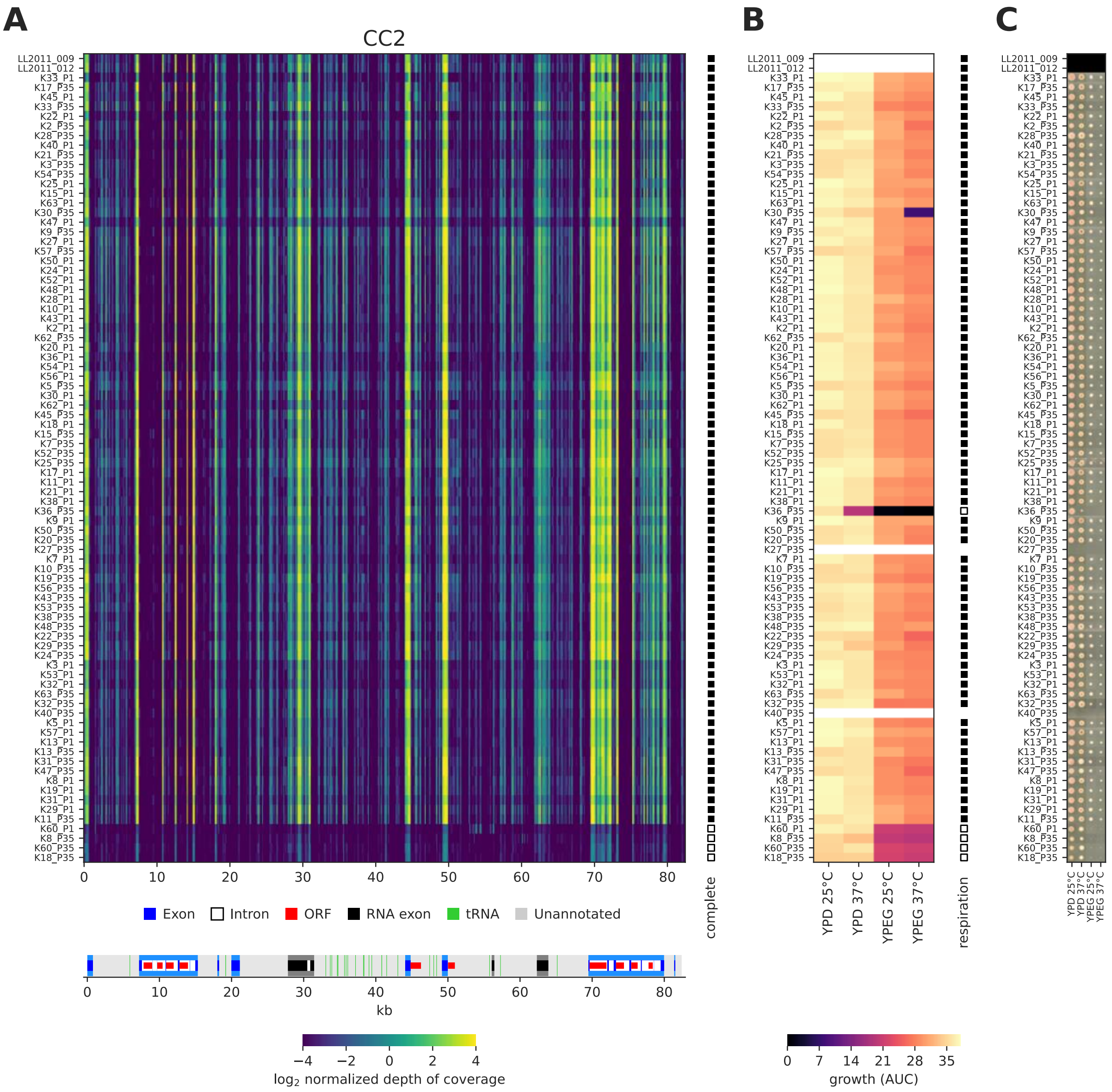

**Supplemental Figure S23. MtDNA deletions and respiration for CC2 lines.**
(*A*) Depth of coverage of short reads along the mtDNA sequence for haplotypes of the CC2 cross at the initial (P1) and final (P35) timepoints. Individual MA lines are stacked horizontally, with the parental strains at the top. Full and empty squares at the right of the heatmap show haplotype classification as complete or deleted, respectively. Annotation summary is shown at the bottom. Genome coordinates are in kb. (*B*) Growth AUC for corresponding lines. Full and empty squares at the right of the heatmap show line classification as respiring or non-respiring, respectively. (*C*) Raw colony images after four days of incubation for corresponding lines.

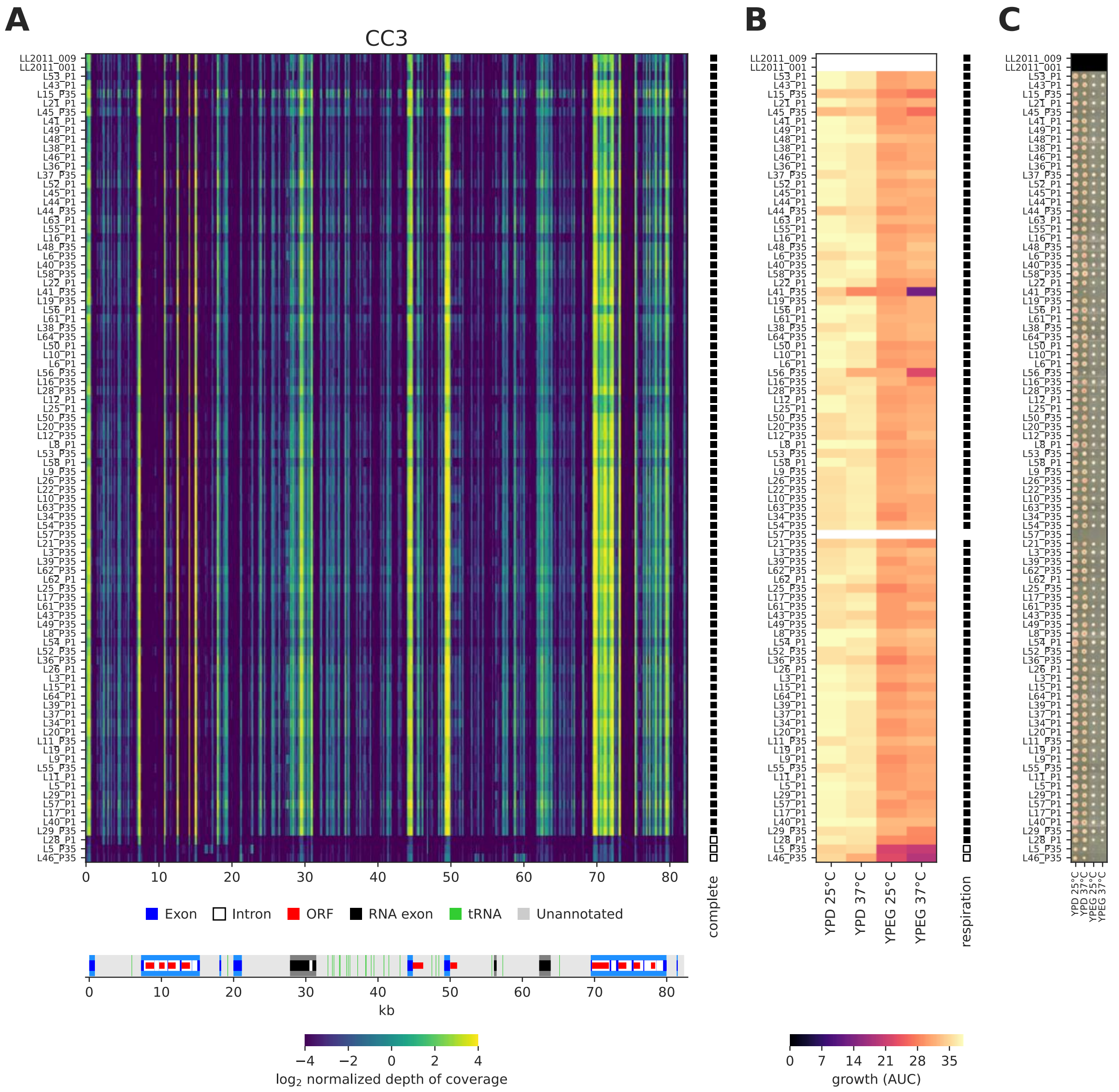

**Supplemental Figure S24. MtDNA deletions and respiration for CC3 lines.**
(*A*) Depth of coverage of short reads along the mtDNA sequence for haplotypes of the CC3 cross at the initial (P1) and final (P35) timepoints. Individual MA lines are stacked horizontally, with the parental strains at the top. Full and empty squares at the right of the heatmap show haplotype classification as complete or deleted, respectively. Annotation summary is shown at the bottom. Genome coordinates are in kb. (*B*) Growth AUC for corresponding lines. Full and empty squares at the right of the heatmap show line classification as respiring or non-respiring, respectively. (*C*) Raw colony images after four days of incubation for corresponding lines.

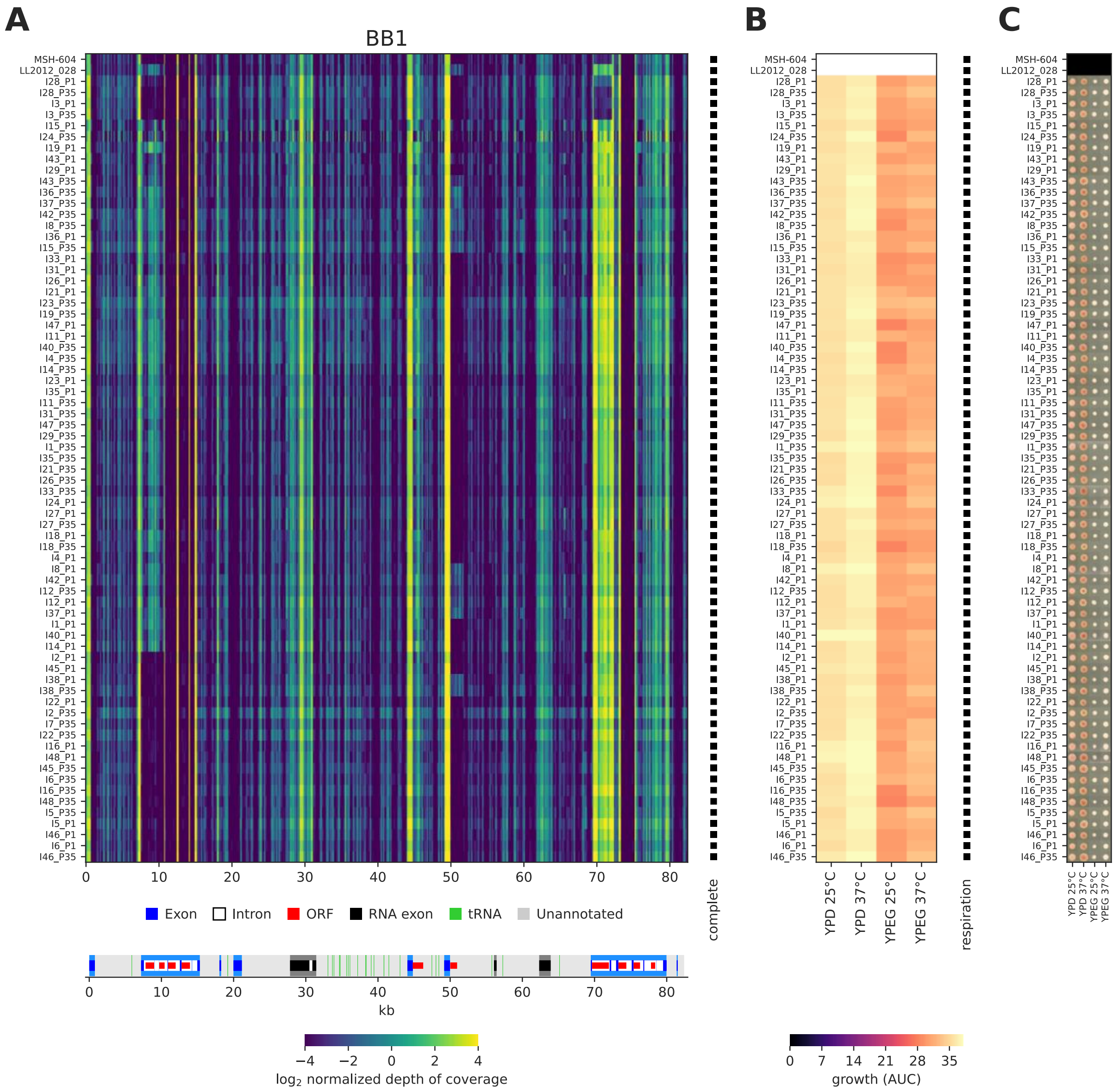

**Supplemental Figure S25. MtDNA deletions and respiration for BB1 lines.**
(*A*) Depth of coverage of short reads along the mtDNA sequence for haplotypes of the BB1 cross at the initial (P1) and final (P35) timepoints. Individual MA lines are stacked horizontally, with the parental strains at the top. Full and empty squares at the right of the heatmap show haplotype classification as complete or deleted, respectively. Annotation summary is shown at the bottom. Genome coordinates are in kb. (*B*) Growth AUC for corresponding lines. Full and empty squares at the right of the heatmap show line classification as respiring or non-respiring, respectively. (*C*) Raw colony images after four days of incubation for corresponding lines.

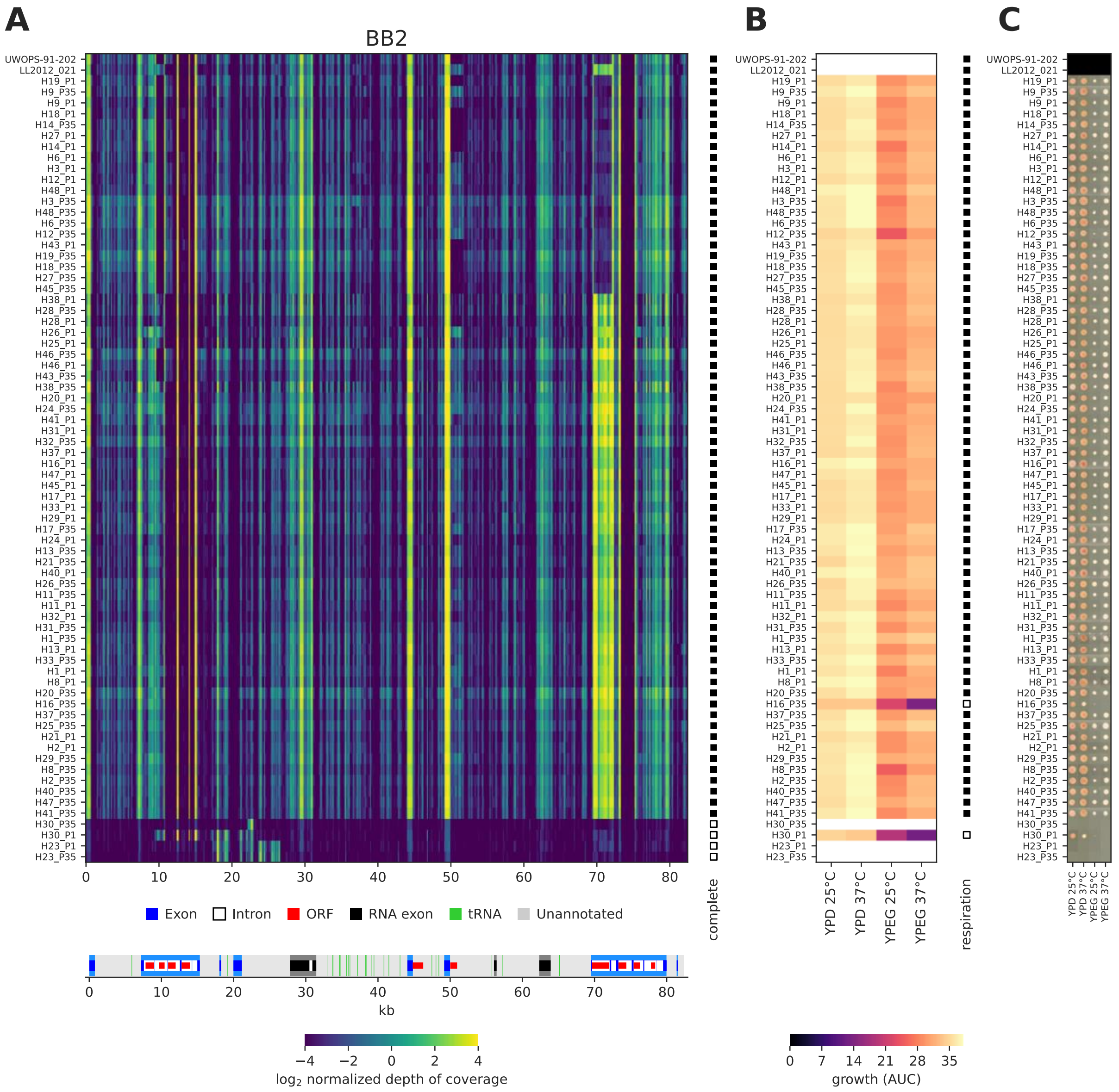

**Supplemental Figure S26. MtDNA deletions and respiration for BB2 lines.**
(*A*) Depth of coverage of short reads along the mtDNA sequence for haplotypes of the BB2 cross at the initial (P1) and final (P35) timepoints. Individual MA lines are stacked horizontally, with the parental strains at the top. Full and empty squares at the right of the heatmap show haplotype classification as complete or deleted, respectively. Annotation summary is shown at the bottom. Genome coordinates are in kb. (*B*) Growth AUC for corresponding lines. Full and empty squares at the right of the heatmap show line classification as respiring or non-respiring, respectively. (*C*) Raw colony images after four days of incubation for corresponding lines.

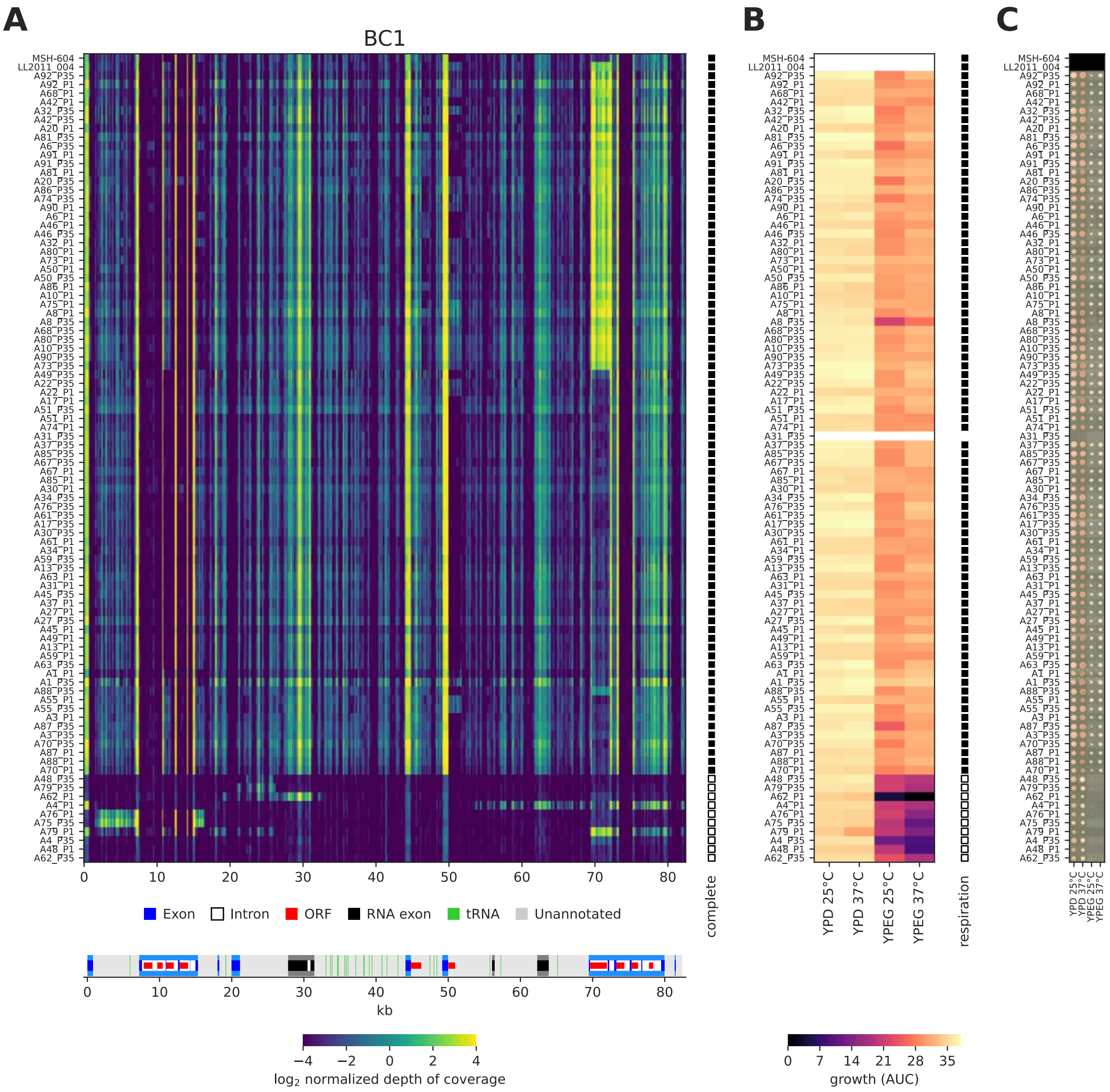

**Supplemental Figure S27. MtDNA deletions and respiration for BC1 lines.**
(*A*) Depth of coverage of short reads along the mtDNA sequence for haplotypes of the BC1 cross at the initial (P1) and final (P35) timepoints. Individual MA lines are stacked horizontally, with the parental strains at the top. Full and empty squares at the right of the heatmap show haplotype classification as complete or deleted, respectively. Annotation summary is shown at the bottom. Genome coordinates are in kb. (*B*) Growth AUC for corresponding lines. Full and empty squares at the right of the heatmap show line classification as respiring or non-respiring, respectively. (*C*) Raw colony images after four days of incubation for corresponding lines.

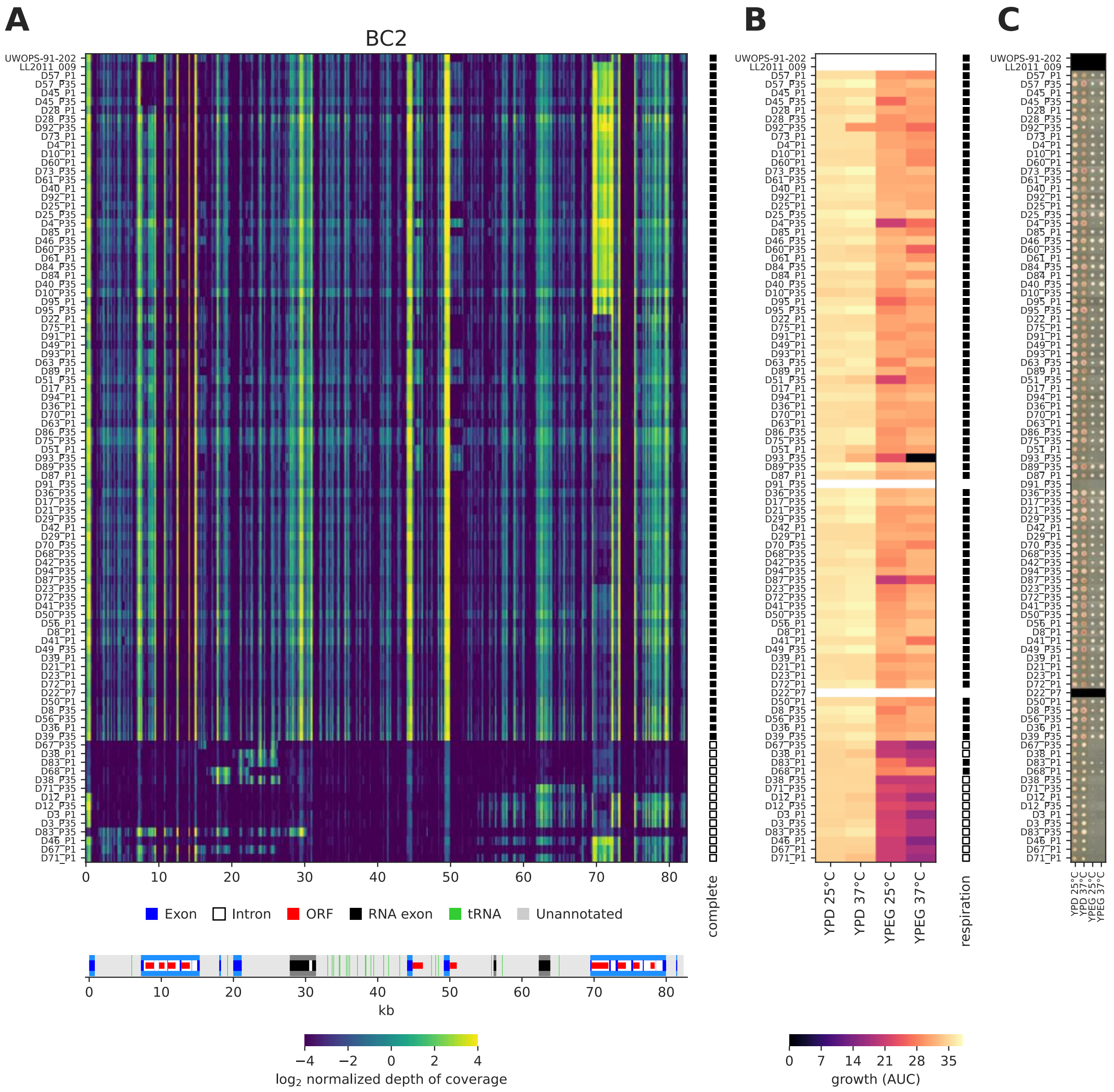

**Supplemental Figure S28. MtDNA deletions and respiration for BC2 lines.**
(*A*) Depth of coverage of short reads along the mtDNA sequence for haplotypes of the BC2 cross at the initial (P1) and final (P35) timepoints. Individual MA lines are stacked horizontally, with the parental strains at the top. Full and empty squares at the right of the heatmap show haplotype classification as complete or deleted, respectively. Annotation summary is shown at the bottom. Genome coordinates are in kb. (*B*) Growth AUC for corresponding lines. Full and empty squares at the right of the heatmap show line classification as respiring or non-respiring, respectively. (*C*) Raw colony images after four days of incubation for corresponding lines.

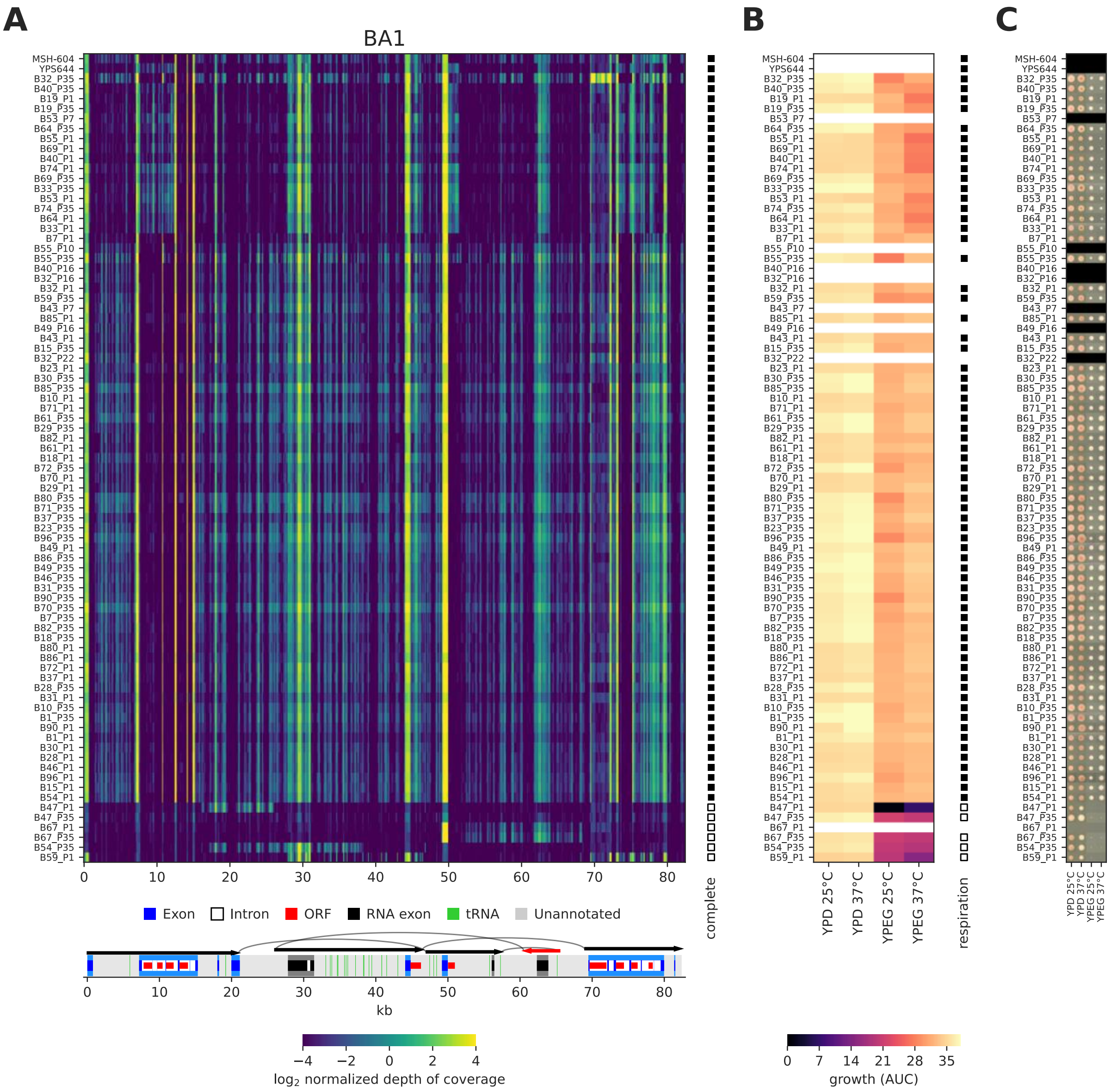

**Supplemental Figure S29. MtDNA deletions and respiration for BA1 lines.**
(*A*) Depth of coverage of short reads along the mtDNA sequence for haplotypes of the BA1 cross at the initial (P1) and final (P35) timepoints. Individual MA lines are stacked horizontally, with the parental strains at the top. Full and empty squares at the right of the heatmap show haplotype classification as complete or deleted, respectively. Annotation summary is shown at the bottom. Genome coordinates are in kb. (*B*) Growth AUC for corresponding lines. Full and empty squares at the right of the heatmap show line classification as respiring or non-respiring, respectively. (*C*) Raw colony images after four days of incubation for corresponding lines.

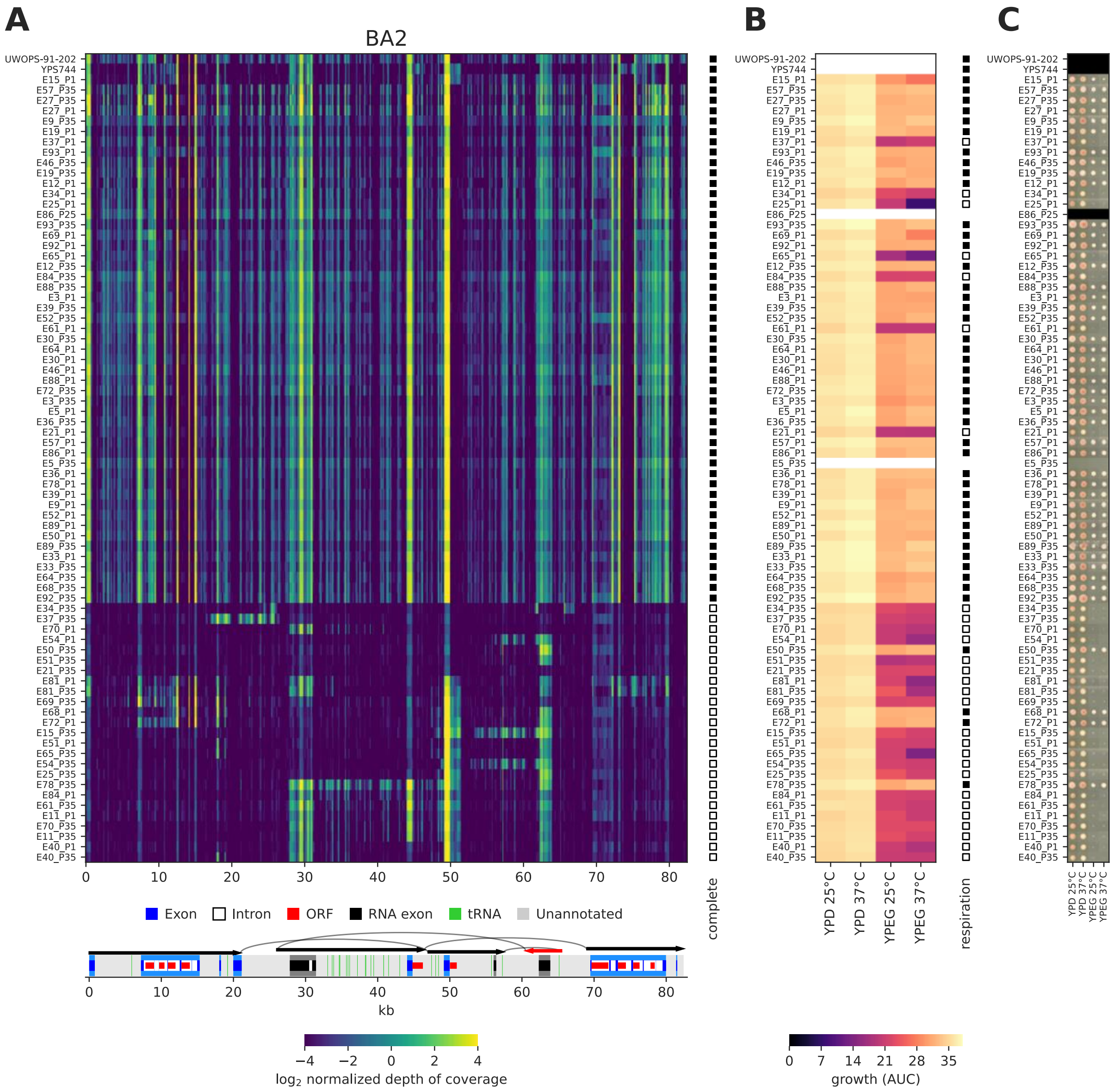

**Supplemental Figure S30. MtDNA deletions and respiration for BA2 lines.**
(*A*) Depth of coverage of short reads along the mtDNA sequence for haplotypes of the BA2 cross at the initial (P1) and final (P35) timepoints. Individual MA lines are stacked horizontally, with the parental strains at the top. Full and empty squares at the right of the heatmap show haplotype classification as complete or deleted, respectively. Annotation summary is shown at the bottom. Genome coordinates are in kb. (*B*) Growth AUC for corresponding lines. Full and empty squares at the right of the heatmap show line classification as respiring or non-respiring, respectively. (*C*) Raw colony images after four days of incubation for corresponding lines.

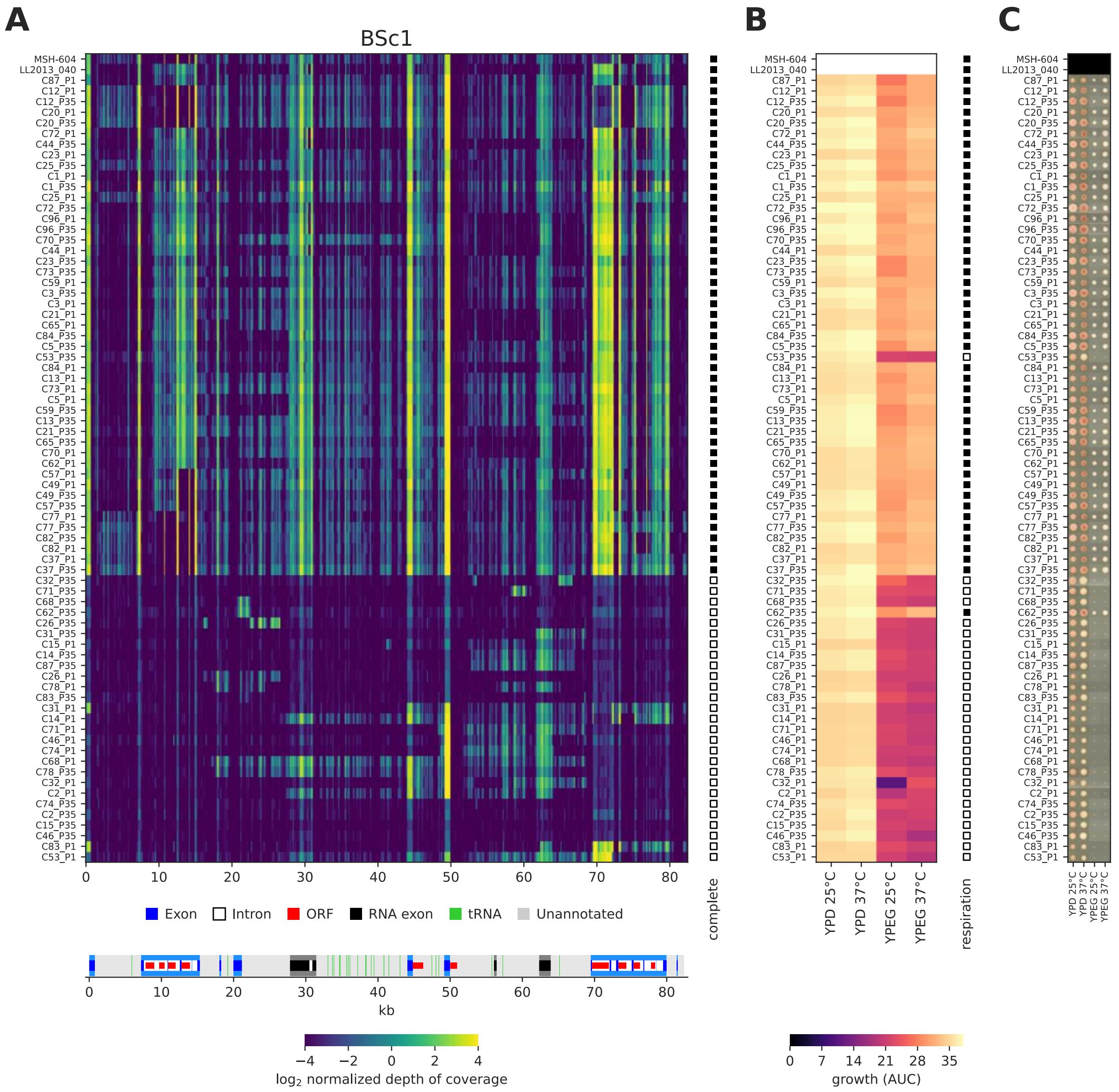

**Supplemental Figure S31. MtDNA deletions and respiration for BSc1 lines.**
(*A*) Depth of coverage of short reads along the mtDNA sequence for haplotypes of the BSc1 cross at the initial (P1) and final (P35) timepoints. Individual MA lines are stacked horizontally, with the parental strains at the top. Full and empty squares at the right of the heatmap show haplotype classification as complete or deleted, respectively. Annotation summary is shown at the bottom. Genome coordinates are in kb. (*B*) Growth AUC for corresponding lines. Full and empty squares at the right of the heatmap show line classification as respiring or non-respiring, respectively. (*C*) Raw colony images after four days of incubation for corresponding lines.

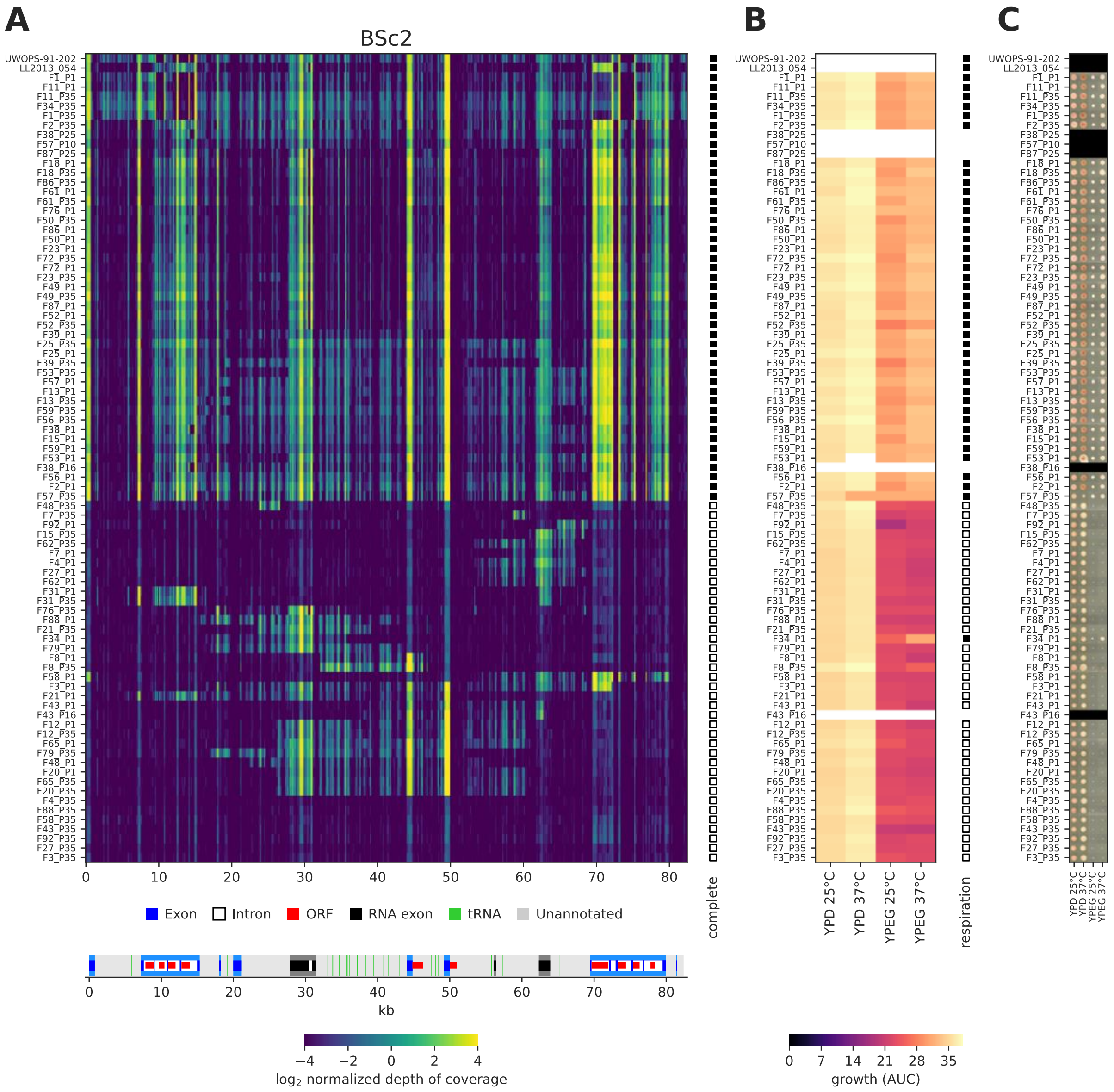

**Supplemental Figure S32. MtDNA deletions and respiration for BSc2 lines.**
(*A*) Depth of coverage of short reads along the mtDNA sequence for haplotypes of the BSc2 cross at the initial (P1) and final (P35) timepoints. Individual MA lines are stacked horizontally, with the parental strains at the top. Full and empty squares at the right of the heatmap show haplotype classification as complete or deleted, respectively. Annotation summary is shown at the bottom. Genome coordinates are in kb. (*B*) Growth AUC for corresponding lines. Full and empty squares at the right of the heatmap show line classification as respiring or non-respiring, respectively. (*C*) Raw colony images after four days of incubation for corresponding lines.

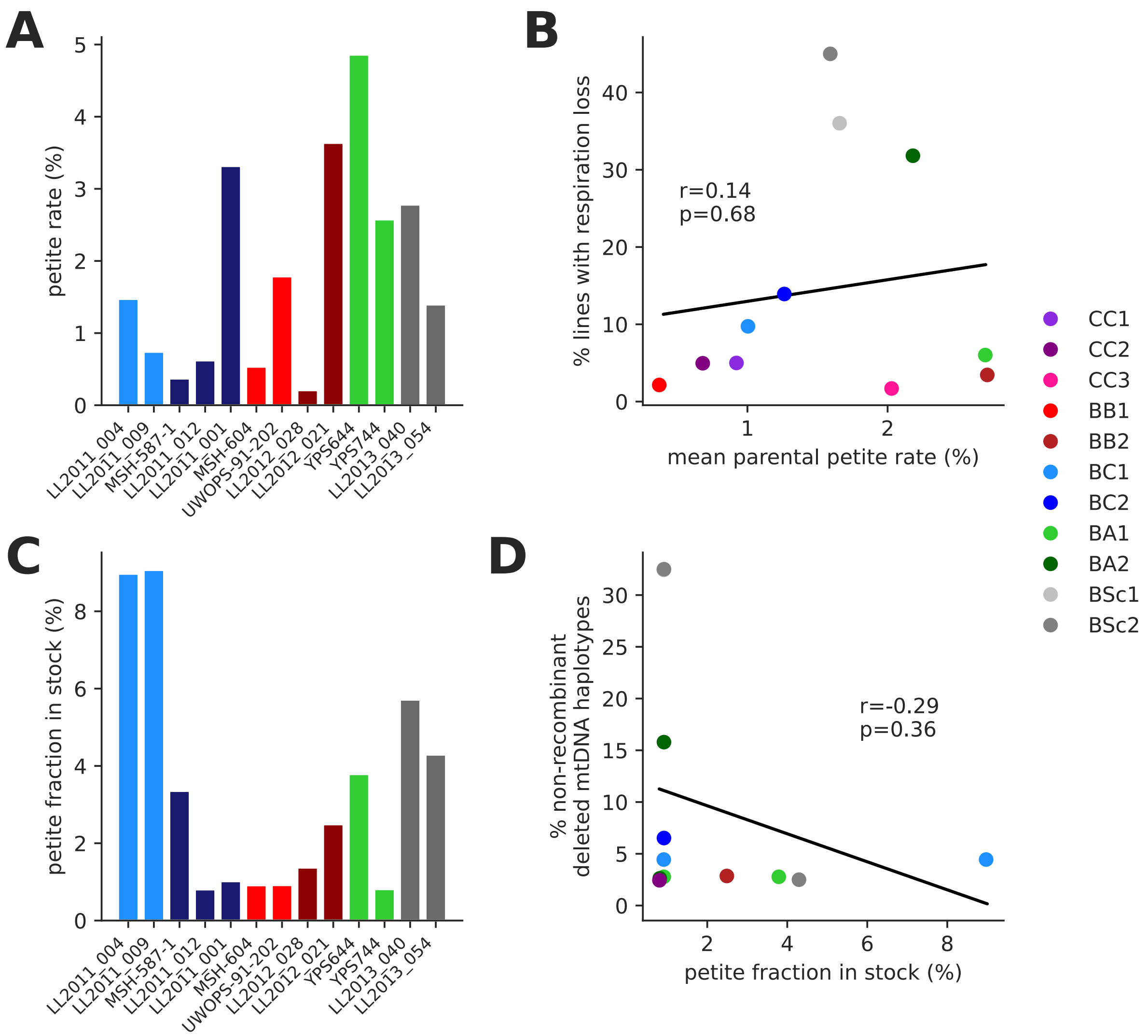

**Supplemental Figure S33. Frequency of loss of respiration in parental strains.**
*(A)* Rate of spontaneous loss of respiration in parental strains, measured as the frequency of petite colonies from cell resuspensions of large single colonies. *(B)* Rate of respiration loss among MA lines as a function of average parental rate of spontaneous petite formation. *(C)* Fraction of non-respiring cells in the stock cultures of the parental strains, measured as the frequency of petite colonies from cell resuspensions of stock patches. *(D)* Fraction of lines at the initial timepoint of the MA experiment which harbored parental, non-recombinant mtDNA haplotypes with large deletions, as a function of petite fraction in the corresponding parental stock culture.

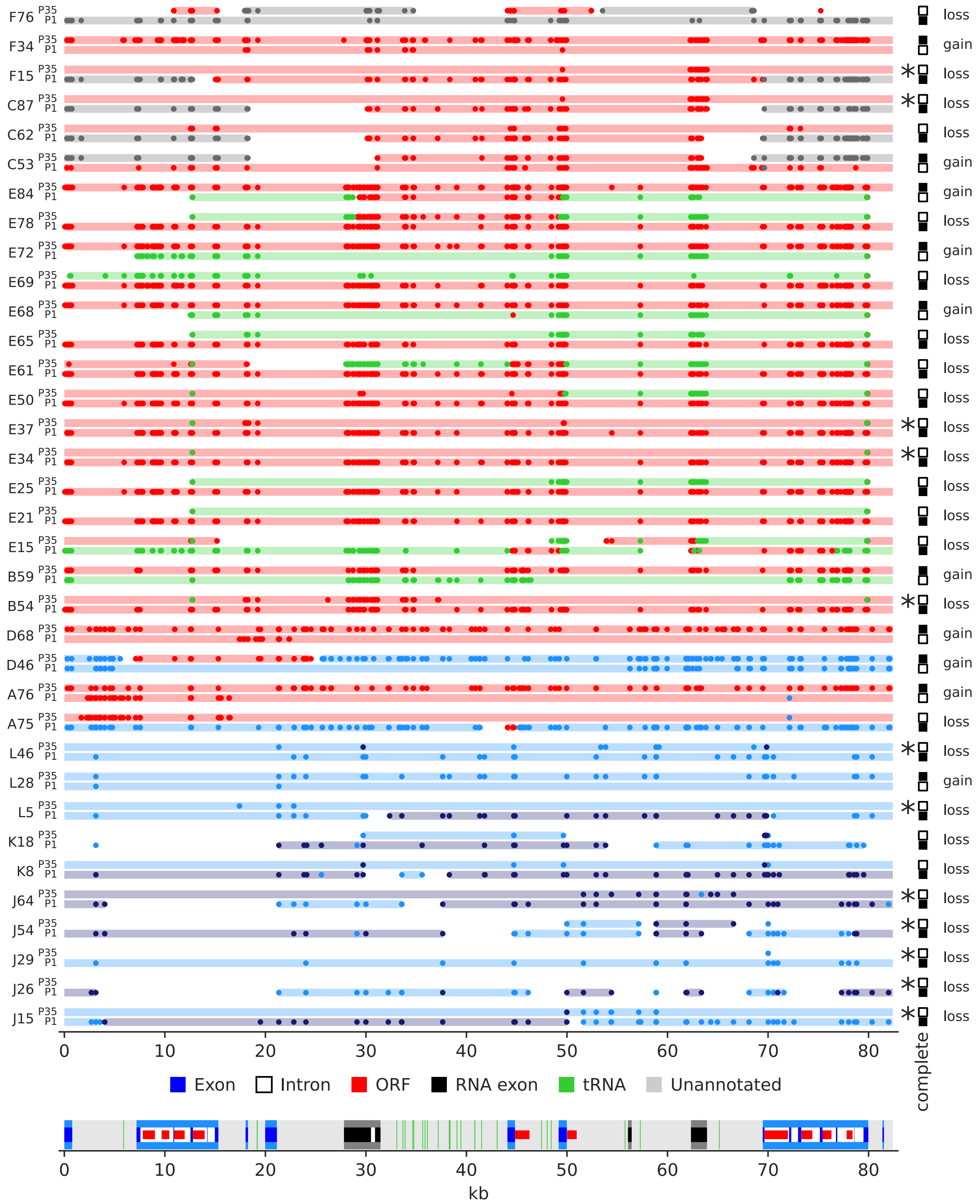

**Supplemental Figure S34. MA lines that experienced changes in mtDNA integrity.** Haplotypes at the initial (P1) and final (P35) timepoints are shown. Full and empty squares at the right show haplotype classification as complete or deleted, respectively, with the direction of the change (loss or gain of a complete mtDNA) indicated. Lines with losses of complete mtDNAs that are consistent with deletions occurring during evolution are shown by asterisks. Recombination tracts and supporting marker variants are shown as in Supplemental Fig. S4-14. Annotation summary is shown at the bottom. Genome coordinates are in kb.

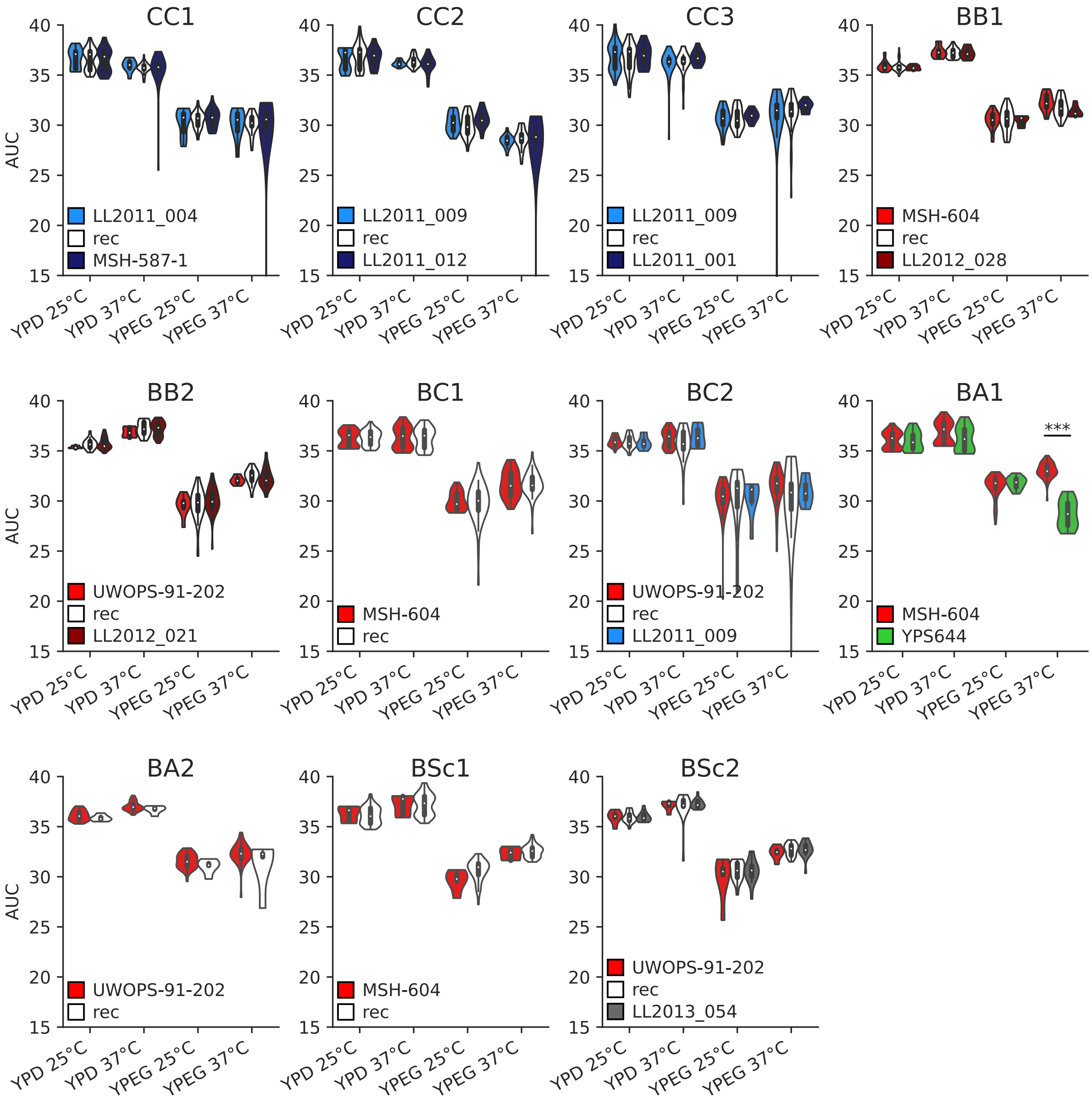

**Supplemental Figure S35. Effect of mtDNA haplotype on the growth of respiring lines.** Area under growth curve (AUC) is shown as a function of mtDNA haplotype category on media containing fermentable (glucose, YPD) or non-fermentable (glycerol and ethanol, YPEG) at 25°C or 37°C. FDR-corrected p-values are shown for Mann-Whitney *U* tests (***: p≤0.001).

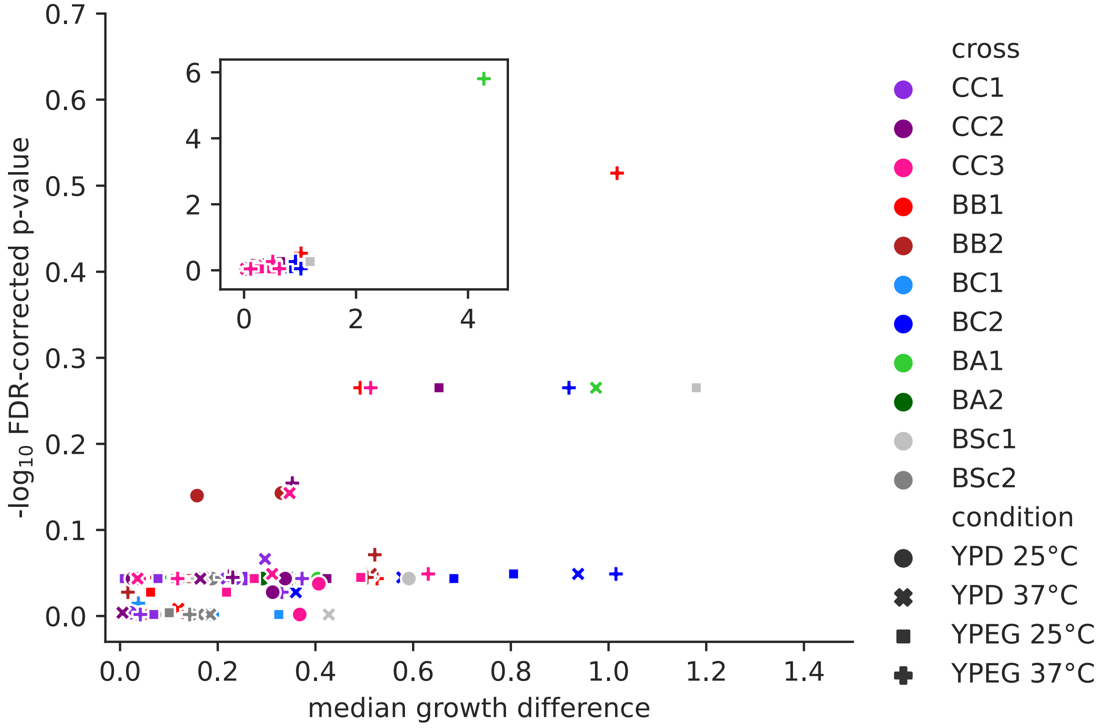

**Supplemental Figure S36. Effect size and significance levels for comparisons of growth AUC between parental and recombinant mtDNA haplotypes for respiring lines.**
The y axis shows FDR-corrected p-values for Mann-Whitney *U* tests and the x axis shows the absolute difference between median AUC values. The inset shows all the data, including the only significant comparison between *SpB* and *SpA* haplotypes in BA1 on YPEG at 37°C.

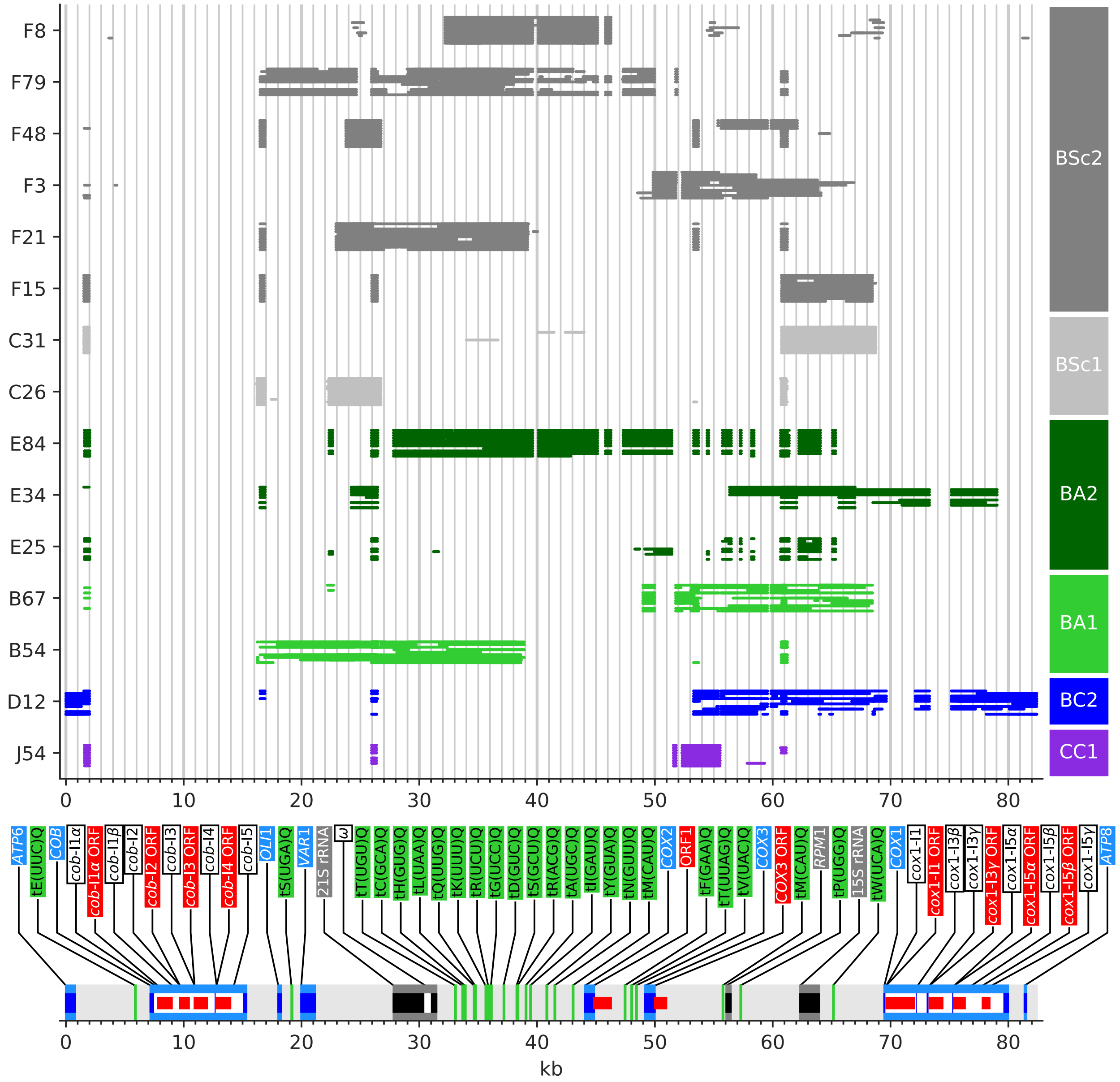

**Supplemental Figure S37. Deletion breakpoints from long-read mtDNA assemblies.**
For each MA line on the y axis, stacked bars display the alignment of distinct assemblies produced from the grid search of assembly algorithm parameters (see Methods). Coordinates and annotation feature types of left and right deletion breakpoints for the major segment of each line are detailed in Supplemental Table S4. Annotation summary is shown at the bottom. Genome coordinates are in kb.

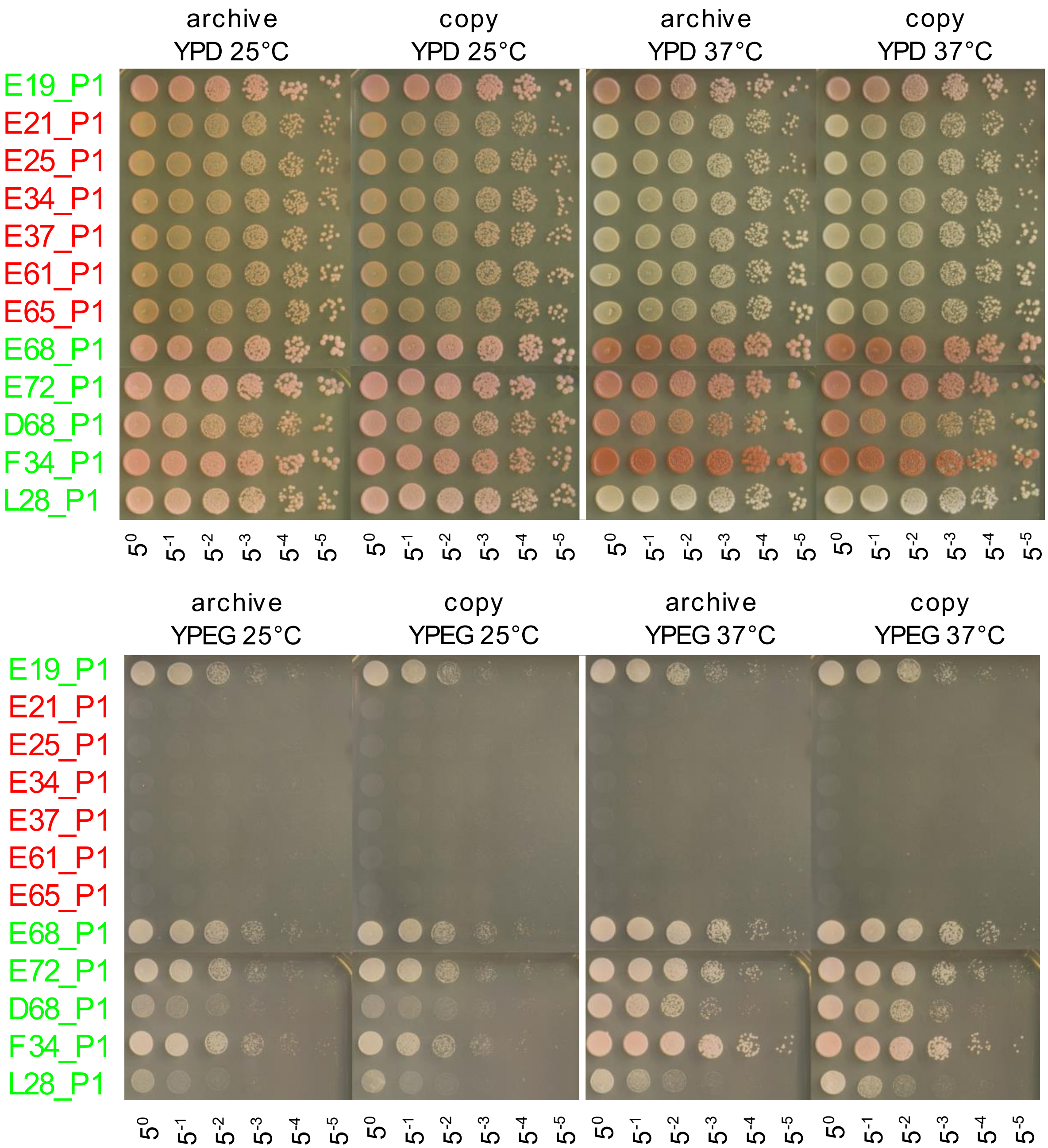

**Supplemental Figure S38. Growth phenotypes of lines with mtDNA instability.**
Lines marked in green and red were phenotyped as respiring and non-respiring in the main screen, respectively. Left panels show the original archive stocks, and right panels show the copies of stocks from which the main screen was performed. Serial dilutions are marked below spots, $5^0$ corresponding to 1 $OD_{600}$ $ml^{-1}$. All lines were sampled at the initial timepoint (P1).

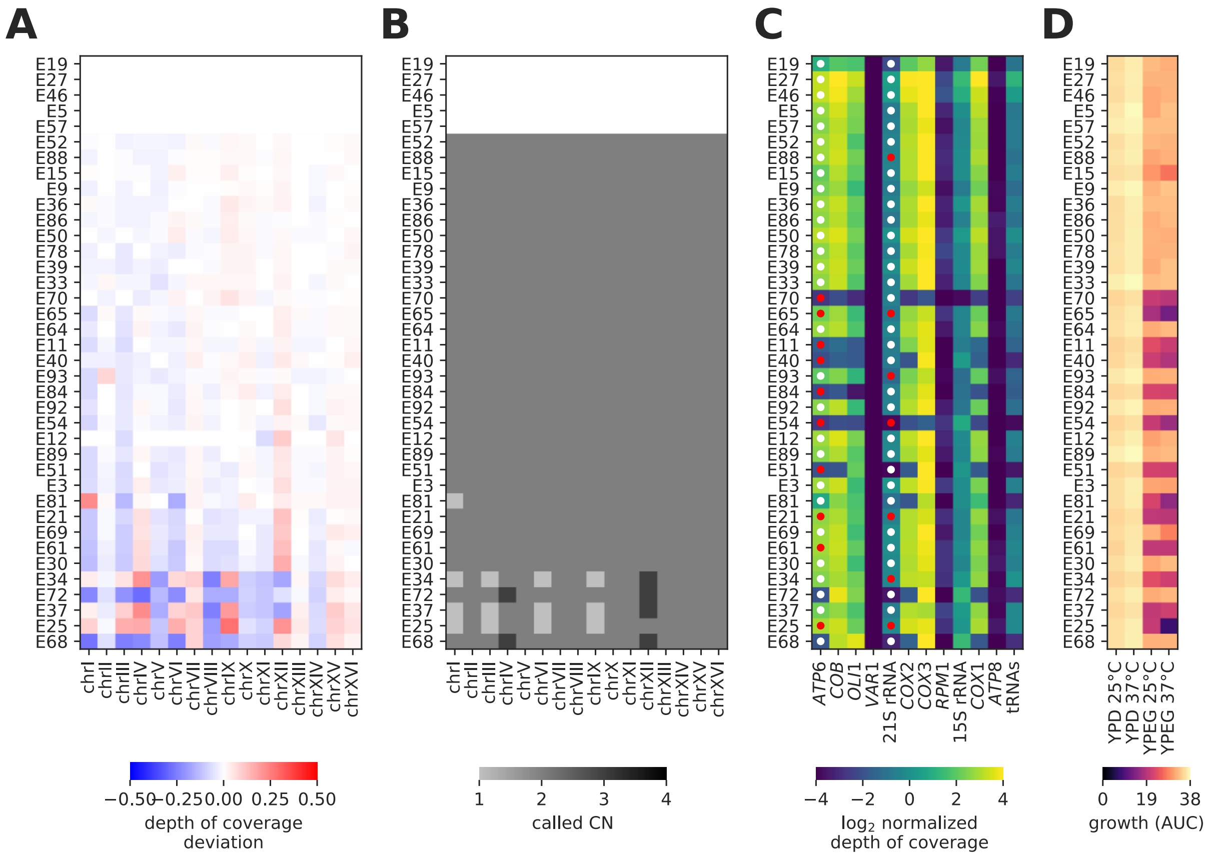

**Supplemental Figure S39. Aneuploidy instability and mtDNA instability in BA2 lines at the initial timepoint of the MA experiment.**

*(A)* Deviation in relative depth of coverage from per-chromosome ploidy calls. Rows represent individual MA lines at the initial timepoint of the MA experiment. Lines are sorted by absolute deviation averaged across all chromosomes. *(B)* Chromosome copy number calls from Fijarczyk et al. 2021. *(C)* Normalized depth of coverage grouped by mtDNA features. White and red dots indicate detected and undetected amplicon at the corresponding locus, respectively. *(D)* Growth AUC in the four conditions tested.

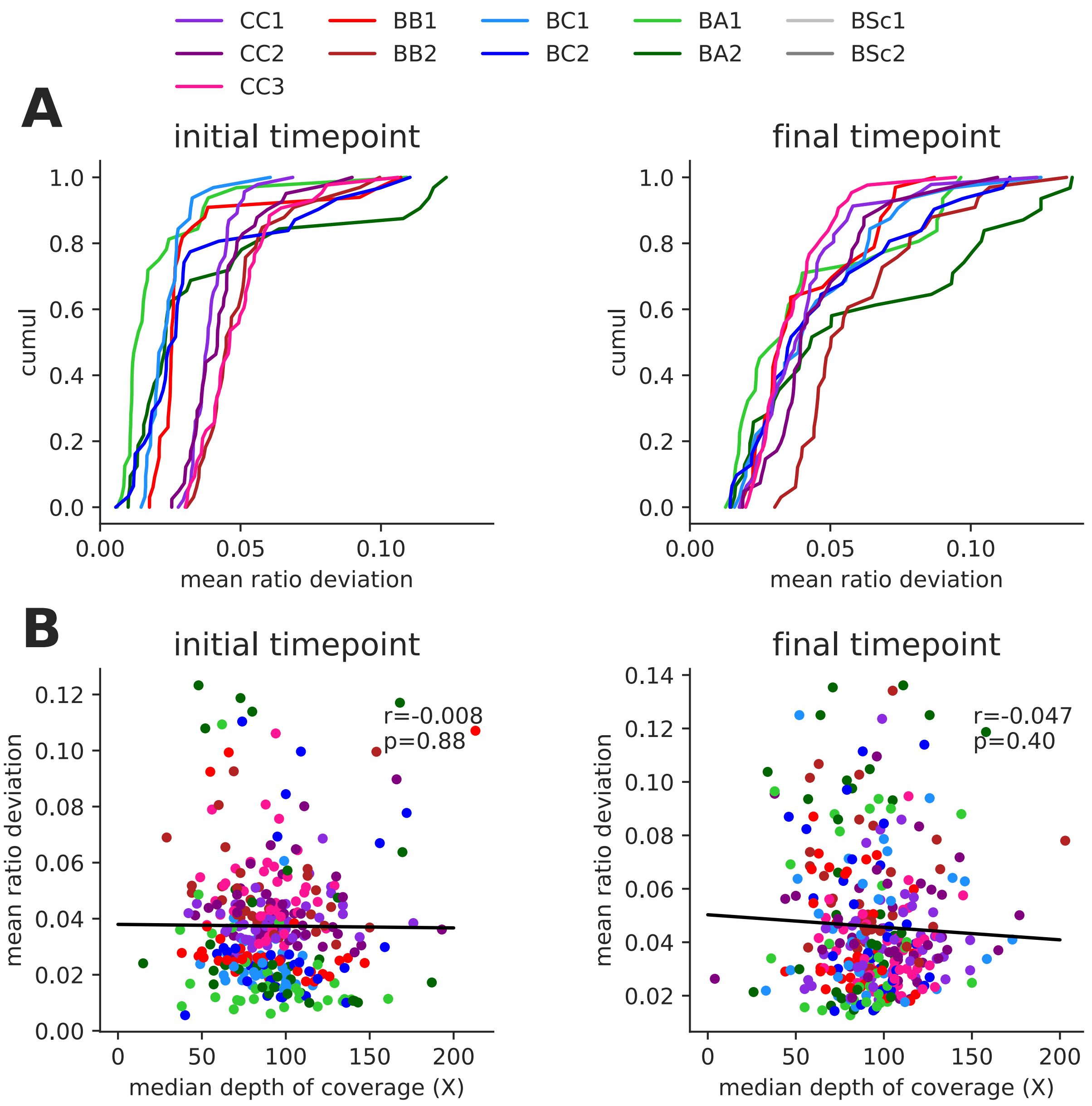

**Supplemental Figure S40. Quantification of aneuploidy instability from depth of coverage deviations on nuclear chromosomes.**

*(A)* Cumulative distributions of deviations in relative depth of coverage from per-chromosome ploidy calls, averaged across all chromosomes, for individual lines at the initial (left) and final (right) timepoints of the MA experiment. *(B)* Pearson correlation between average deviation and median depth of coverage for nuclear chromosomes.

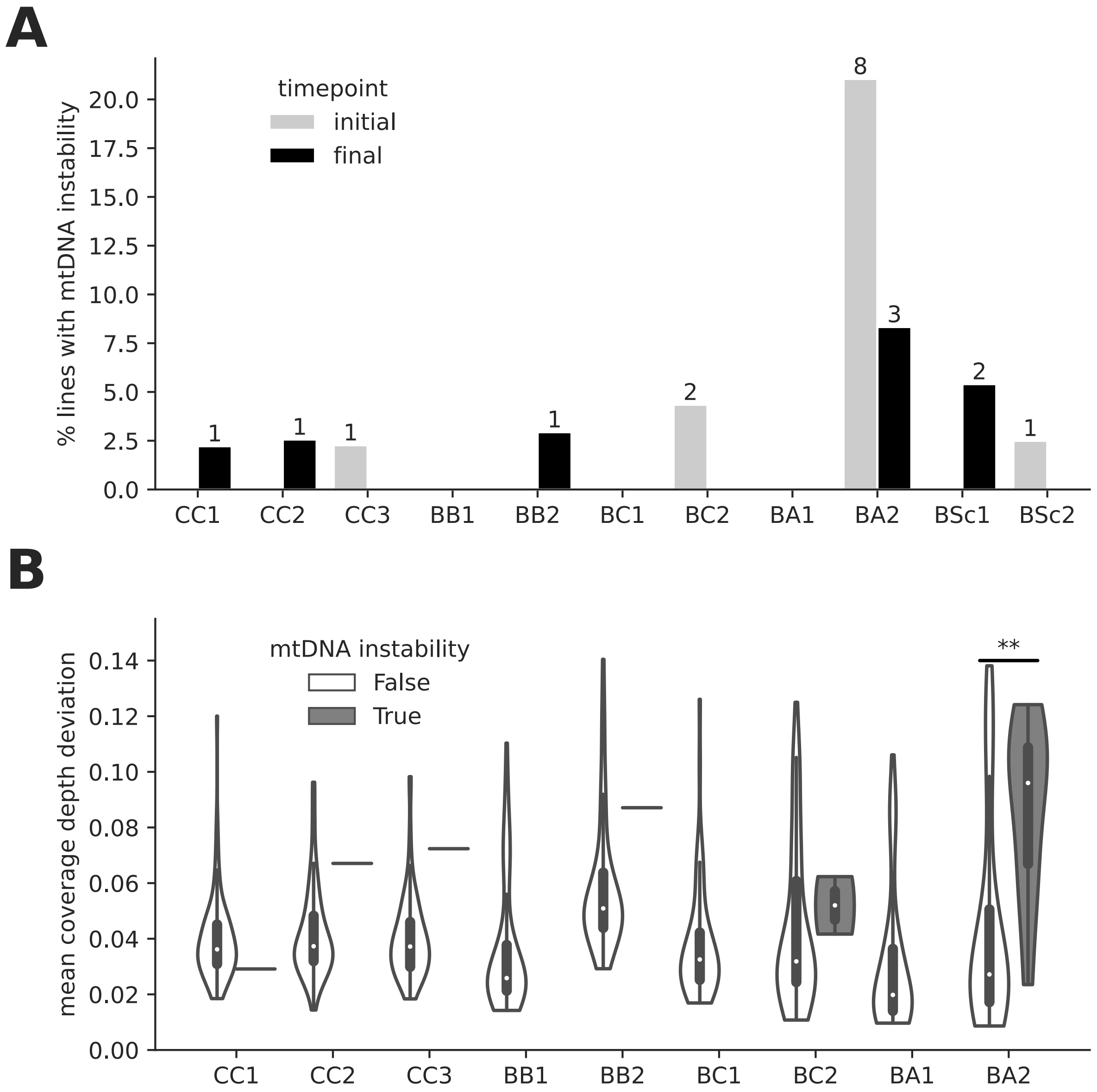

**Supplemental Figure S41. Associations between mtDNA instability and aneuploidy instability.**

*(A)* Percentage of lines at the initial and final timepoint with mtDNA instability. *(B)* Deviation in depth of coverage on nuclear chromosomes for lines grouped by mtDNA instability. FDR-corrected p-values for Mann-Whitney *U* test are shown (**: p≤0.01).
